# A cerebellar disinhibitory circuit supports synaptic plasticity

**DOI:** 10.1101/2023.09.15.557147

**Authors:** Changjoo Park, Jawon Gim, Sangkyu Bahn, Gyu Hyun Kim, Yoonseok Im, Sang-Hoon Lee, Kisuk Lee, Min-Soo Kim, Kea Joo Lee, Jinseop S. Kim

## Abstract

How does the cerebellum learn how to control motion? The cerebellar motor learning critically depends on the long-term depression of the synapses between granule cells and Purkinje cells, which encode motor commands and inhibitory modifications to motor outputs, respectively, for simultaneous granule cell inputs and climbing fibre inputs, the latter of which encode the error signals^1–3^. However, recent studies have revealed that inhibitory inputs to Purkinje cells may disrupt long-term depression^4–8^, and it is not clear how long-term depression can occur without disruption. In search of a clue, we investigated the synaptic connectivity among the neurons reconstructed from serial electron microscopy images of the cerebellar molecular layer^9,10^. We discovered synapses between climbing fibres and a subset of inhibitory interneurons, which synapse onto the remaining interneurons, which in turn synapse onto Purkinje cells. Such connectivity redefines the interneuron types, which have been defined morphologically or molecularly^11–13^. Together with climbing fibres to Purkinje cell connections, those cell types form a feedforward disinhibitory circuit^14^. We argued that this circuit secures long-term depression by suppressing inhibition whenever climbing fibre input is provided and long-term depression needs to occur^15^, and we validated the hypothesis through a computational model. This finding implies a general principle of circuit mechanism in which disinhibition supports synaptic plasticity^16,17^.

## Main

The cerebellum codes modulatory signals for motion and cognition^18–20^. External excitatory inputs enter the cerebellar molecular layer (CML) through parallel fibres (PFs) and climbing fibres (CFs), which are the axons of granule cells and axons of inferior olivary neurons, respectively. These inputs converge on Purkinje cells (PCs), which are the only output neurons of the CML^21,22^ (Fig. 1a). According to the cerebellar motor learning theory by Marr, Albus, and Ito^1–3^, the combinations of PF activities carry the efferent copy of the motor commands. As the CF inputs to PCs send teaching signals on the errors of motor commands, PF to PC synapses undergo long-term depression (LTD) upon simultaneous inputs from PFs and CFs. The consequent changes in PC outputs modulate motor output differently and correct motions (Fig. 1b).

**Figure 1.**
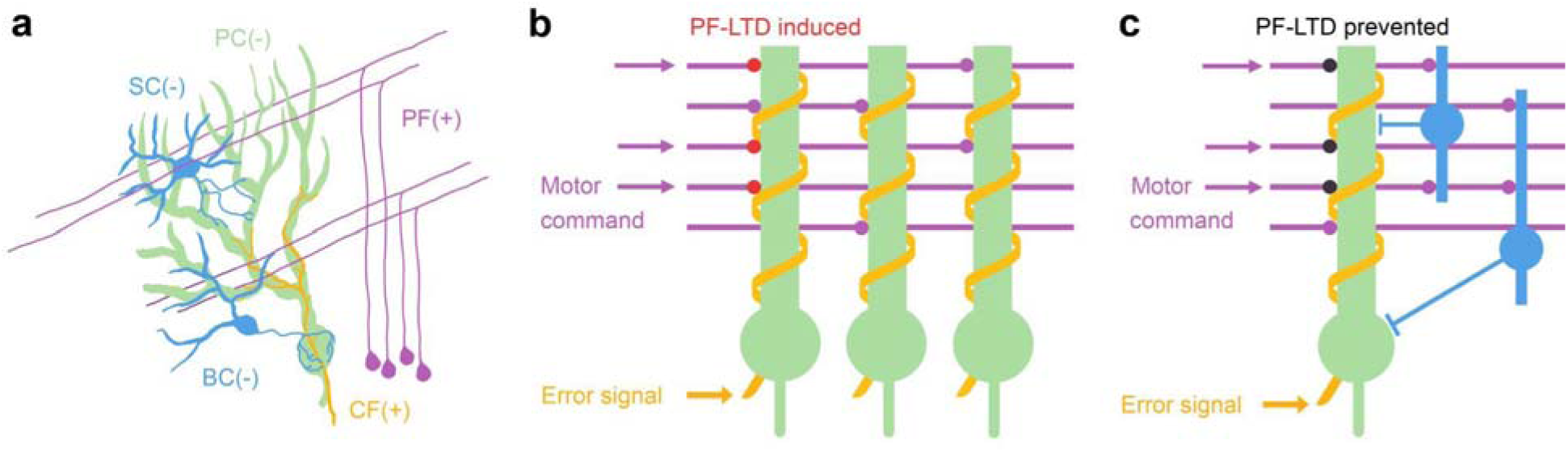
The Marr-Albus model of cerebellar motor learning. **a,** A unit of the CML microcircuits of excitatory PF (purple) and CF (orange) inputs and inhibitory IN (SC and BC, blue) inputs converging on PC (green). **b,** Illustration of the Marr-Albus-Ito model. Among the PF-PC synapses (purple), those encoding a particular motor command (purple arrows) that are coactivated with an error signal from the CF (orange arrow) undergo LTD (red dots). **c,** The IN inputs computationally contribute to the coordinated motions, however, the LTD of PF-PC synapses is disrupted if CF error signals coincide with IN inputs.

CML interneurons (INs), which are anatomically classified into stellate cells (SCs) and basket cells (BCs), have recently been recognised for their importance in cerebellar computation, but the understanding is still far from complete^23–28^. While INs are not involved in the Marr-Albus-Ito model, recent electrophysiological investigations have shown that synaptic plasticity is regulated by the intracellular calcium ion concentration ([Ca^2+^]_i_) of the PC^29–31^. LTD is induced when [Ca^2+^]_i_ is increased by excitatory inputs, and LTD is not induced or long-term potentiation (LTP) is induced when [Ca^2+^]_i_ is decreased by inhibitory inputs^5–8,32^. Moreover, molecular and chemical interventions of INs have been revealed to alter cerebellar learning as well^33–35^.

These facts about cerebellar synaptic plasticity pose a critical question. The INs, which can easily fire by receiving only a few PF inputs or fire spontaneously^36,37^, can provide inhibitory inputs to PC at any moment. Therefore, teaching signals of CFs are inclined to be suppressed by inhibition, which may disrupt LTD. Inhibition is necessary for cerebellar computation, but it prohibits normal synaptic plasticity (Fig. 1c). How can this paradox be resolved, and what is the exact role of inhibition in cerebellar learning?

In search of an answer, we studied the mouse CML by analysing the images from 3D serial block-face scanning EM^38^. We volumetrically reconstructed the neurons and semiautomatically found the synapses with pipelines using artificial intelligence (AI)^9,10,39^. The connectivity among the cell types was investigated to discover that CFs have synaptic connections to a subset of INs, which we call IN2s^40^, while CFs and INs have been known to communicate only by a non-synaptic mechanism^14,41^. We further discovered that IN2s have preferential connections to the remaining INs, which we call IN1s. IN1s have preferential connections to PCs with few connections to other INs. Such a disinhibition circuit^14^ supports cerebellar LTD because CF-triggered disinhibition suppresses the inhibition of INs to PCs whenever the CF teaching signal arrives at a PC and LTD is expected at the PF to PC synapses^15^. We validated this hypothetical mechanism through simulations using a computational model. The newly defined IN types based on connectivity may complement the anatomically defined types and agree with the molecularly defined types^13^. We argue that inhibition and disinhibition play critical roles in synaptic plasticity^16,42^.

## Results

### Reconstruction of EM images

The EM dataset was prepared from a small slice of lobule IV/V of the mouse cerebellar cortex by serial block-face scanning electron microscopy^38^ (Fig. 2a and Methods). The imaged area chiefly included the CML with a minor portion of the Purkinje cell layer (PCL). A total of 1,000 parasagittal EM images were obtained in 2×3 tiles each with 12 nm×12 nm×50 nm per voxel resolution, where one tile consisted of 5,000×5,000 voxels with 10∼20% overlap to adjacent tiles. We registered the images using TrakEM2 software with minor modifications (Methods). The registration resulted in a 3D image stack with an approximate size of 14,600×10,200×1,000 voxels (175 µm×122 µm×50 µm) (Fig. 2b and Supplementary Data 1).

**Figure 2.**
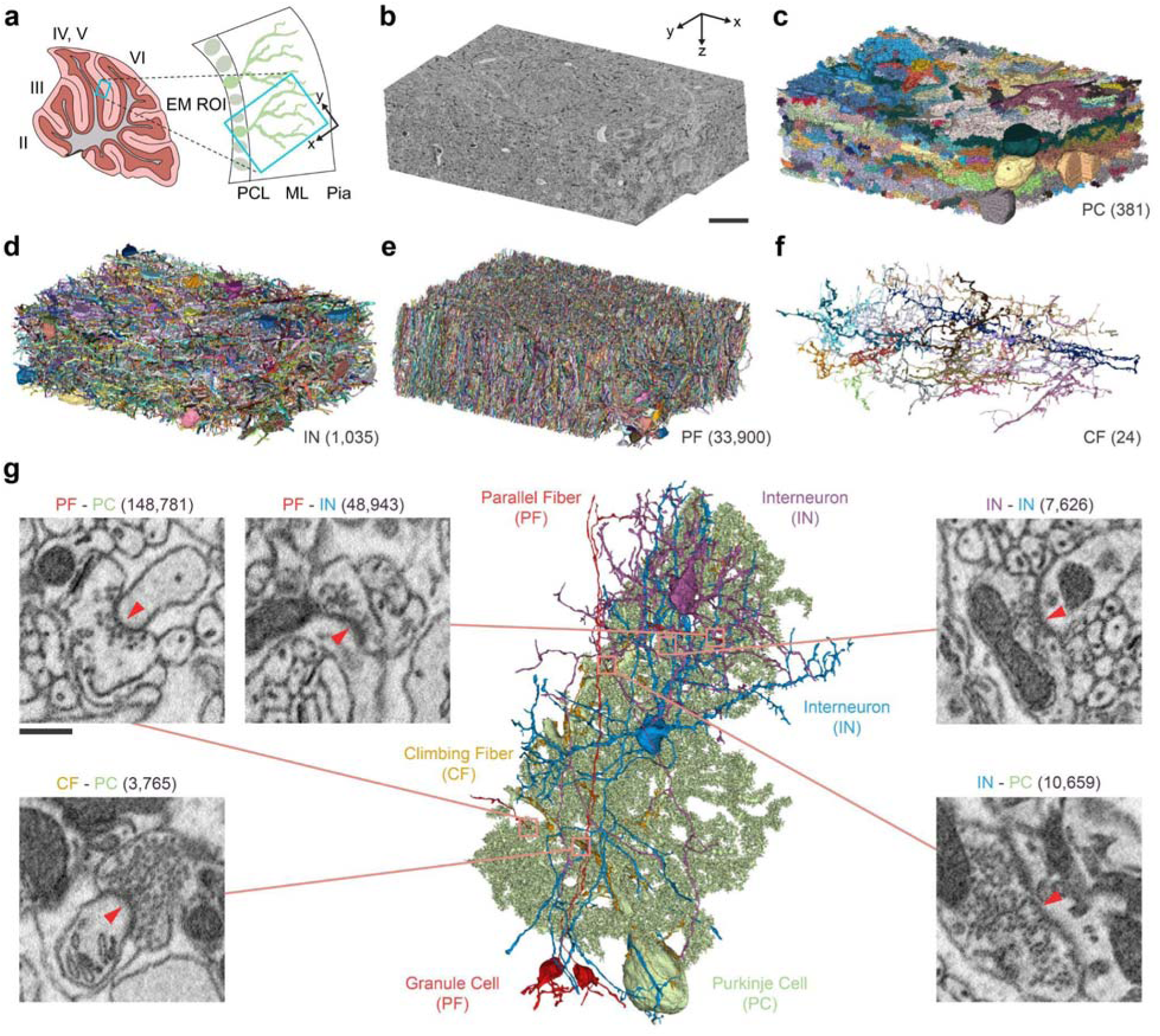
Reconstruction of the neural circuits in the CML. **a,** A midsagittal section of lobule IV/V of an adult mouse at P56 was sampled (cyan box), which mainly includes the molecular layer and partially the Purkinje cell layer. **b,** The 3D EM image volume after alignment was 175 _μ_m×122 ^μ^m×50 _μ_m. The *xyz*-axes are defined as shown. Scale bar: 25 _μ_m. **c–f,** The 3D rendering of reconstructed neurons grouped by cell type: PC, IN, PF and CF, respectively, with their numbers in parentheses. **g,** The examples of synapses are displayed by the pre- and postsynaptic neuronal types with the numbers in the parentheses, which were identified using a semiautomated method. The red triangles point from the postsynaptic to the presynaptic neurons on the EM images. Scale bar: 0.5 _μ_m.

We analysed the image volume using a semiautomated reconstruction pipeline, which is equivalent to what was used in a previous study^9^ comprising machine-learning segmentation of the image volume followed by human proofreading (Methods). The reconstruction resulted in 35,798 neurons or neuronal fragments (Fig. 2c–f and Supplementary Data 1). The reconstruction was estimated to cover 60.3% of the entire volume of the dataset (Methods).

Upon the completion of the reconstruction of a neuron or neuronal fragment, it was classified into one of the four cell or fibre types of the CML (Methods). The classification yielded 381 PCs, 24 CFs, 1,035 INs, and 33,900 PFs in total. We regard the granule cells and any of their axonal parts as PFs for brevity. We estimated that 99.6% of PCs, 97.7% of INs, 100% of CFs, and 76.9% of PFs were reconstructed in terms of the number of reconstructed cells over the estimated total number of cells in the dataset (Methods). In the following analyses, we used 10 PCs (Supplementary Data 2a) and 26 INs (Supplementary Data 3a) whose somata are intact or whose reconstructed volume is large (more than 5×10^8^ voxels for PCs and 6×10^7^ voxels for INs); 10 CFs (Supplementary Data 4a) innervating the 10 PCs; and all 33,900 PFs, called the main cells. The remaining cell fragments were used for supplementary analyses where necessary.

We also inspected the neural connectivity between reconstructed neurons by using the automated synapse detection method, which demonstrated over 95% accuracy for a small sample of the dataset^39^. Since accuracy varies with the cell types forming the synapses because of the heterogeneous number of synapse examples in the training set, we reviewed and proofread the results when required (Methods). It yielded a total of 219,774 synapses, comprising 148,781 PF-PC, 48,943 PF-IN, 3,765 CF-PC, 7,626 IN-IN, and 10,659 IN-PC synapses (Fig. 2g and Supplementary Data 2b, c and Supplementary Data 3b–d and Supplementary Data 4b). Compared to a recent study that showed the structured connectivity between the granule cells and PCs from EM reconstruction^10^, we mapped the circuits more densely in a smaller dataset.

### Cellular organisation around PCs

We were able to classify the CML cell types by using the classical descriptions of their morphologies as the criteria (Methods). This implies that the morphological descriptions established based on light microscopy are still generally correct when EM is applied to observe the same neurons. Then, what was newly found from the EM reconstruction? A closer inspection of the CFs revealed that the CFs arborise parasagittally on both normal faces of the flat PC dendrites and only occasionally cross the tangential side to reach the opposite face (Fig. 3a, b and Supplementary Data 1, 2a, 4a). This observation is contradictory to the conventional description that CFs helically wind around PC dendritic shafts^21,43^ (Fig. 3b, top).

**Figure 3.**
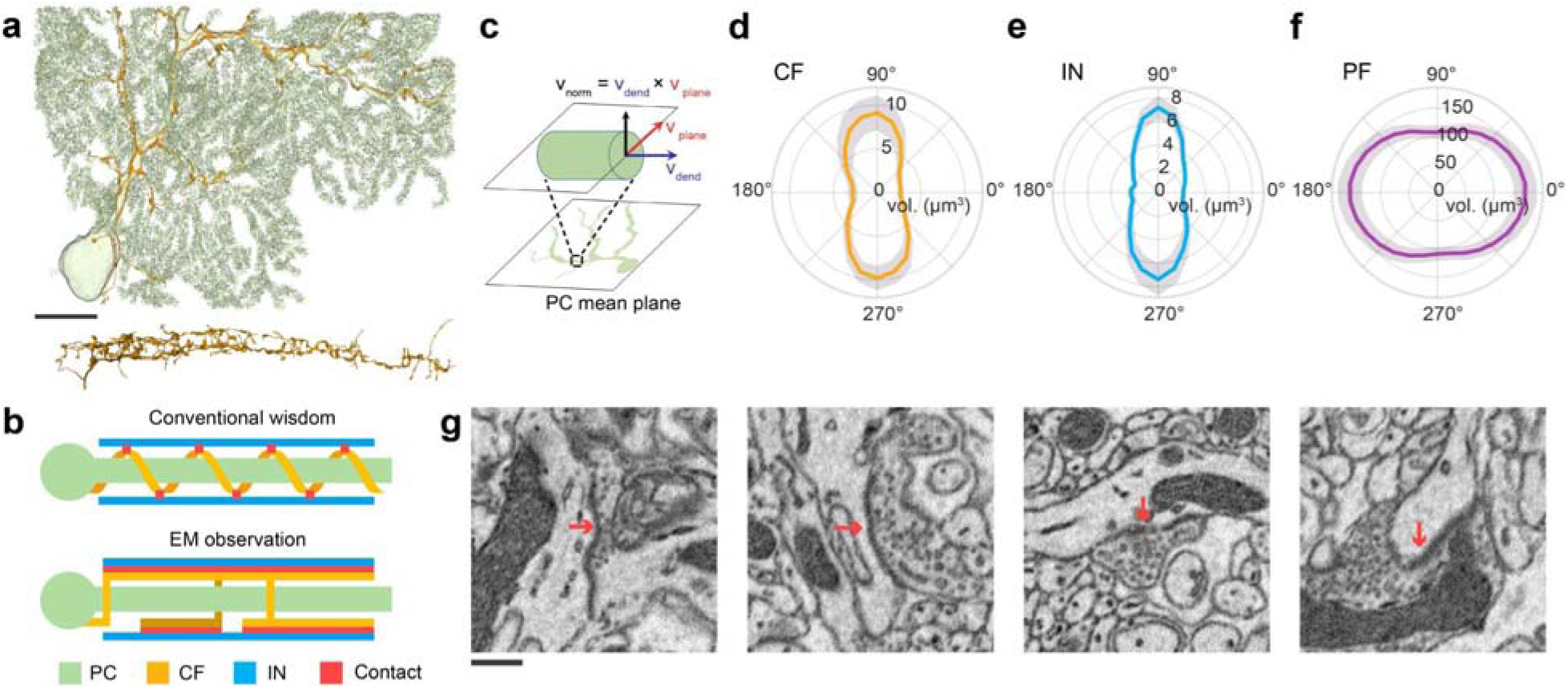
CFs arborise on the normal faces of PCs and synapse to INs. **a,** 3D renderings of a PC (green) and its presynaptic CF (orange) in the XY-view (top) and the same CF in the XZ-view (bottom). Scale bar: 25 µm. **b,** Diagrams of the XZ-view for a PC, a CF, and an IN, illustrating the possible locations of CF-IN contact (red) expected by conventional wisdom (top) versus the EM observation (bottom) for CF arborisation. **c,** Definition of the local axes in the dendritic cross section. ***v***_dend_: local tangential vector of the dendritic skeleton, ***v***_plane_: a vector in the mean plane of the PC dendrites, ***v***_norm_: the normal vector defined as ***v***_norm_ = ***v***_dend_×***v***_plane_. **d–f,** Polar plots of the angular distributions of CF, IN, and PF volume around the PC dendrites. The direction of ***v***_norm_ is defined to be 90°. The shade is the area of standard errors. **g,** EM sections showing four examples of CF to IN synapses. Scale bar: 0.5 µm.

The quantitative measurement of the CF volume distribution around the PC dendritic skeleton (Fig. 3c and Methods) confirmed this observation. The skeleton of a neuron is the topology-preserving tree graph whose nodes are located along the central axis of the neuronal processes. Indeed, the distribution was most dense near 90° and 270°, the orientation of normal faces (Fig. 3d and Supplementary Data 2f). The INs had a similar distribution, while PFs had the opposite distribution which was dense near 0° and 180° (Fig. 3e, f and Supplementary Data 2g, h). What would be the reason for this organisation? We suggested two hypotheses that are not mutually exclusive.

The first hypothesis was that the processes of different neuronal types efficiently divide and occupy the limited space around PC dendrites. The PFs were denser on the tangential sides than on the normal faces of PC dendrites (Fig. 3f) because PFs pass through the cleft of the flat PC dendrites. Then, the PC spines may also be denser on the tangential sides to effectively make connection with the PFs. Therefore, the CFs and INs were assumed to prefer the normal faces of PC dendrites to avoid the dense plexus of PFs and PC spines. We will further investigate this hypothesis in future work.

The second hypothesis, which we focused on in this study, was that the CFs arborise in a laminar manner to increase the chance of forming contacts with PCs or INs in the next laminae (Fig 3b, bottom). The increased contacts to PCs in the next laminae may result in establishing synapses to extra PCs, which were recently discovered to exist^44^; however, such synapse was not found in our dataset maybe because of the limited size (Fig. 4a). The enhancement of contact with the INs may increase the efficacy of glutamate spillover, which is the nonsynaptic communication between CFs and INs near CF boutons due to strong neurotransmitter release^41^. Nevertheless, a more interesting question would be whether the enhanced CF-IN contacts lead to CF-IN synapses.

**Figure 4.**
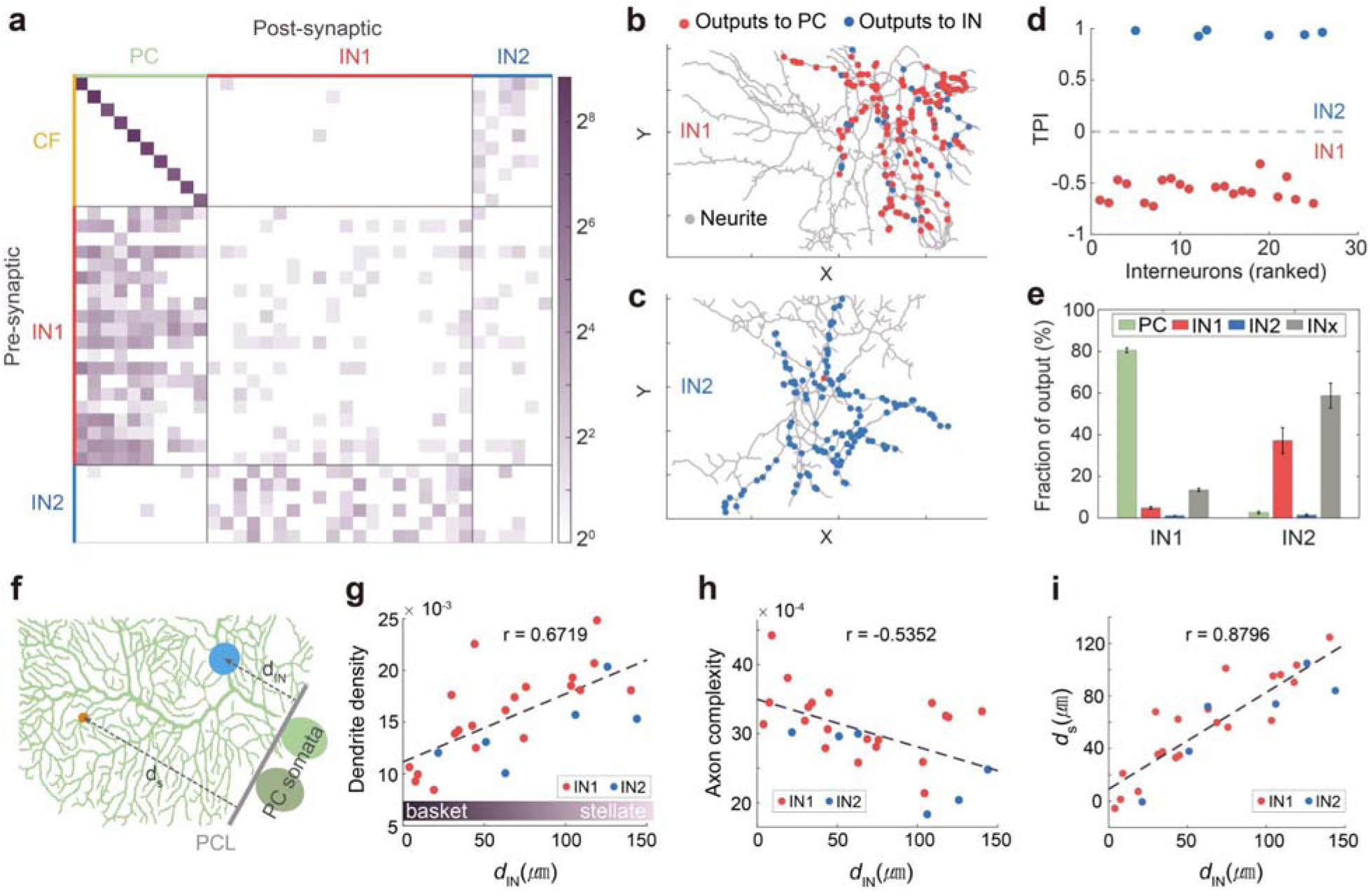
Connectivity-based vs. morphologically-defined IN types. **a,** Weighted adjacency matrix of the neurons where the colour intensity represents the number of synapses on the log scale. **b–c,** Visualisation of the IN skeleton and its synaptic output to PC (red) or IN (blue), demonstrating that the major synaptic target of IN1 is PCs and that of IN2 is INs. **d,** The target preference index (TPI) defines the two IN types. **e,** The target distribution of IN1 and IN2. The error bars are standard errors. INx indicates the IN fragments whose type was undetermined. **f,** Diagram illustrating the definitions of the PCL boundary, IN soma distance (*d*_IN_) and IN-PC synapse distance (*d*_s_). **g–i,** Dendrite density **(g)**, axon complexity **(h)**, and *d*_s_ **(i)** versus *d*_IN_. The red and blue dots indicate IN1 and IN2, respectively. The synapse distance is represented by the average distance of all IN-PC synapses.

### CFs synapse on INs selectively

Indeed, we discovered CF-IN synapses. They were verified by human experts through visual inspection of automatically detected synapses (Methods). Fifty-three CF-IN synapses were selected (Fig. 3g and Supplementary Data 3b, 4c, 5) out of the total 997 CF-IN contacts in the dataset whose volume is 958,470 µm^3^, yielding 5.5×10^-5^/ µm^3^ of volume synapse density. This type of synaptic connection was the rarest compared to other types (Extended Data Fig. 1a). The linear synapse density, the number of synapses per unit length of neural process, was also the minimum for CF-IN synapses when the input synapses along the IN dendrites and the output synapses along the CF axons were counted by type (Extended Data Fig. 1b, c). Note that the densities were slightly underestimated because the reconstruction of the neurons was not complete. The rareness may be the main reason that the CF-IN synapse has not been observed, and the limited evidence based on a few micrographs has not been widely appreciated^40^. Here, to our knowledge, we presented the first systematic evidence that CF-IN synapses indeed exist.

How does the newly discovered CF-IN connection fit into the CML microcircuit structure (Fig. 1a, c)? The connectivity among the main CFs, PCs, and INs was visualised by a weighted adjacency matrix, whose weight is the number of synapses (Fig. 4a). The one-to-one connection between CFs and PCs was obvious within the dataset. It also appeared that INs can be clustered into two groups, one which connects to PCs frequently (IN1) and the other which rarely does (IN2). The visualisation of the synaptic output on the IN arbours also supported the clustering (Fig. 4b, c and Supplementary Data 3c, e).

To clarify the definitions of IN1 and IN2, we introduced the target preference index (TPI): TPI = (*n*_IN_-*n*_PC_)/(*n*_IN_+*n*_PC_), where *n*_IN_ and *n*_PC_ are the number of output synapses of a given IN to INs and to PCs, respectively. As the TPI values showed a clear distinction between the two groups of INs, we defined IN1s as those with TPI<0 and IN2s as those with TPI>0 (Fig. 4d and Supplementary Data 3e). Among the 26 main INs, the ratio of IN1 to IN2 types was 3.3 to 1 (20 IN1s and 6 IN2s). Additionally, we calculated the TPI for large IN axon fragments to classify the 137 IN axonal fragments into 109 IN1s and 28 IN2s, which gave a 3.9 to 1 ratio (Extended Data Fig. 2a, b). The remaining small IN fragments could not be classified and were denoted as an unknown type, INx.

With the definition of IN1 and IN2, CF-IN connectivity was further translated from the adjacency matrix (Fig. 4a). The CF-IN connections are specific to IN2s, i.e., the IN2s are the IN type that makes the CF-IN synapses chiefly. All 6 IN2s but one were strongly innervated by multiple CFs (mean 9.6 and minimum 2 CF synapses per IN2). Since an IN2 without a CF connection was located at the corner of the dataset and its dendritic arbours were cut off by the boundary, we speculated that it may have had CF synapses outside the dataset. On the other hand, only 3 out of 20 IN1s were weakly innervated by a few CFs (mean 1.7 and maximum 3 CF synapses per IN1).

The proportions of output connections of IN1s exhibited 80.6% to PC and 19.4% to INs (4.8% to IN1s, 1.1% to IN2s, and 13.5% to INxs) for all the IN fragments whose type can be decided. For IN2s, the proportions of output connections were 97.4% to INs (37.2% to IN1s, 1.4% to IN2s, and 58.8% to INxs) and only 2.6% to PCs (Fig. 4e and Supplementary Data 3e). To summarise the CML connectivity, the CFs, which synapsed onto PCs in a one-to-one manner, also synapsed onto multiple IN2s, and the IN2s synapsed to multiple IN1s, each of which synapsed onto multiple PCs. Conversely, one PC was innervated by one CF and multiple IN1s, each of the IN1s by multiple IN2s, and then each of the IN2s by multiple CFs.

As a side note, we found that the angular distributions of both IN1 and IN2 exhibited a preference for the normal orientation of PCs (Extended Data Fig. 2c, d). This is contrary to the common anticipation based on the second hypothesis that IN2s would arborise on the normal faces of PCs and that IN1s would exhibit different distributions because only IN2s are innervated by CFs. This implies that the angular distribution of INs is not solely determined by the synaptic connectivity and that the first hypothesis, that different cell types would separate the space around PCs, may also be valid.

### IN classifications

We classified the CML INs based on connectivity. In contrast, it has been traditionally accepted that INs are clearly classified into BCs and SCs by anatomical cues^11^. It has also been debated that INs form a population with continuously varying morphology^12,45,46^. A recent study argued that BCs and SCs are clearly distinguished by axonal morphologies, while SCs have variability that appears as a continuum^47^. What is the relation between the connectivity-based types and the morphologically defined types, and which is the correct classification criterion?

To examine the relationship, we quantified the three anatomical traits of INs that have been widely accepted as the criteria for anatomical classification^11,12^ (Methods): the distance of the soma from the PCL boundary, the dendritic arborisation pattern, and the axonal arborisation and synaptic connection pattern to PCs. The distances of IN somata (*d*_IN_) and IN-PC synapses (*d*_s_) from the PCL boundary were measured (Fig. 4f). The dendrite density captures SC dendritic arbours occupying limited areas more effectively than BC dendritic arbours, while the axon complexity reflects the perisomatic axonal plexus, called the basket, of BCs^48^.

We plotted the dendrite density, axonal complexity, and the PCL distance of IN-PC synapses as functions of the PCL distance of IN somata for both IN1s and IN2s (Fig. 4g–i and Supplementary Data 2d, 3f–h). The PCL distance of somata and the other three measures exhibited strong correlations, which implies that the measures have similar capability in defining the cell types. Since IN somata are located roughly uniformly along the PCL distance, the choice of the 1/3 CML boundary for BCs versus SCs^11,12^ appears arbitrary. The other quantities in each plot on the *y*-axis also exhibited a roughly even distribution rather than a bimodal distribution, which is expected for classifiable data. These results support the claim that INs form a continuous spectrum^12,47^.

The data points of IN1s and IN2s on these plots appear irrelevant of the measures to categorise BCs and SCs, as seen from the random distribution of red and blue dots (Fig. 4g–i). Quantitatively, Mann–Whitney U tests on the three measures for BC versus SC classification revealed no difference in the distributions of IN1s and IN2s (Extended Data Fig. 3a, c, d), which supports the irrelevance between IN1 versus IN2 classification and BC versus SC classification. One exception was axon complexity (Extended Data Fig. 3b), which may be linked to the fact that distinction between BCs and SCs requires axonal signatures^47^. However, a more rigorous investigation is difficult since our dataset only marginally included the basket axons.

Logically, IN1 versus IN2 classification can be independent of BC versus SC classification, or the two classifications can be hierarchical. Whether two independent classification schemes can exist for the same neurons is uncertain. When the hierarchy is correct, BCs are unlikely to be related to IN2s because IN2s seldom innervate PCs and have no perisomatic basket. Then, the possibilities are either that IN1 and IN2 are subtypes of SC or that SC and BC are subtypes of IN1. The higher axon complexity of IN1s (Extended Data Fig. 3b) may be because some IN1s have baskets, while IN2s do not (Supplementary Data 3d). Therefore, we speculate that BC and SC might need to be redefined as subtypes of IN1. Then, the aforementioned study^47^ may have classified BCs correctly and have merged SC and IN2 into SC with high variability.

A recent study classified INs into molecular layer interneuron (MLI) 1-1, 1-2, and 2 by single-cell RNA sequencing^13^. The population ratio of MLI1s and MLI2s was 3.1:1, which is close to the 4:1 ratio of IN1s and IN2s (Extended Data Fig. 2b). The difference may be ascribed to the small sample size of this study. If the MLI1-IN1 and MLI2-IN2 relation is correct and the speculation that BC and SC are subtypes of IN1 is also correct, BC and SC may correspond to MLI1-1 and 1-2. Further investigation is required to unify these studies and to congruently define CML IN types by molecular, physiological, morphological, and connectional traits^49,50^.

### Disinhibition supports cerebellar motor learning

The novel synaptic connections between the newly defined cell types update the known wiring diagram and form a disinhibitory feedforward loop. The simplified circuit diagram could consist of four cells, one cell for each type (Fig. 5a). A similar disinhibitory interaction has recently been observed physiologically; however, it was explained by the consequence of glutamate spillover and nonspecific IN connectivity^14^.

**Figure 5.**
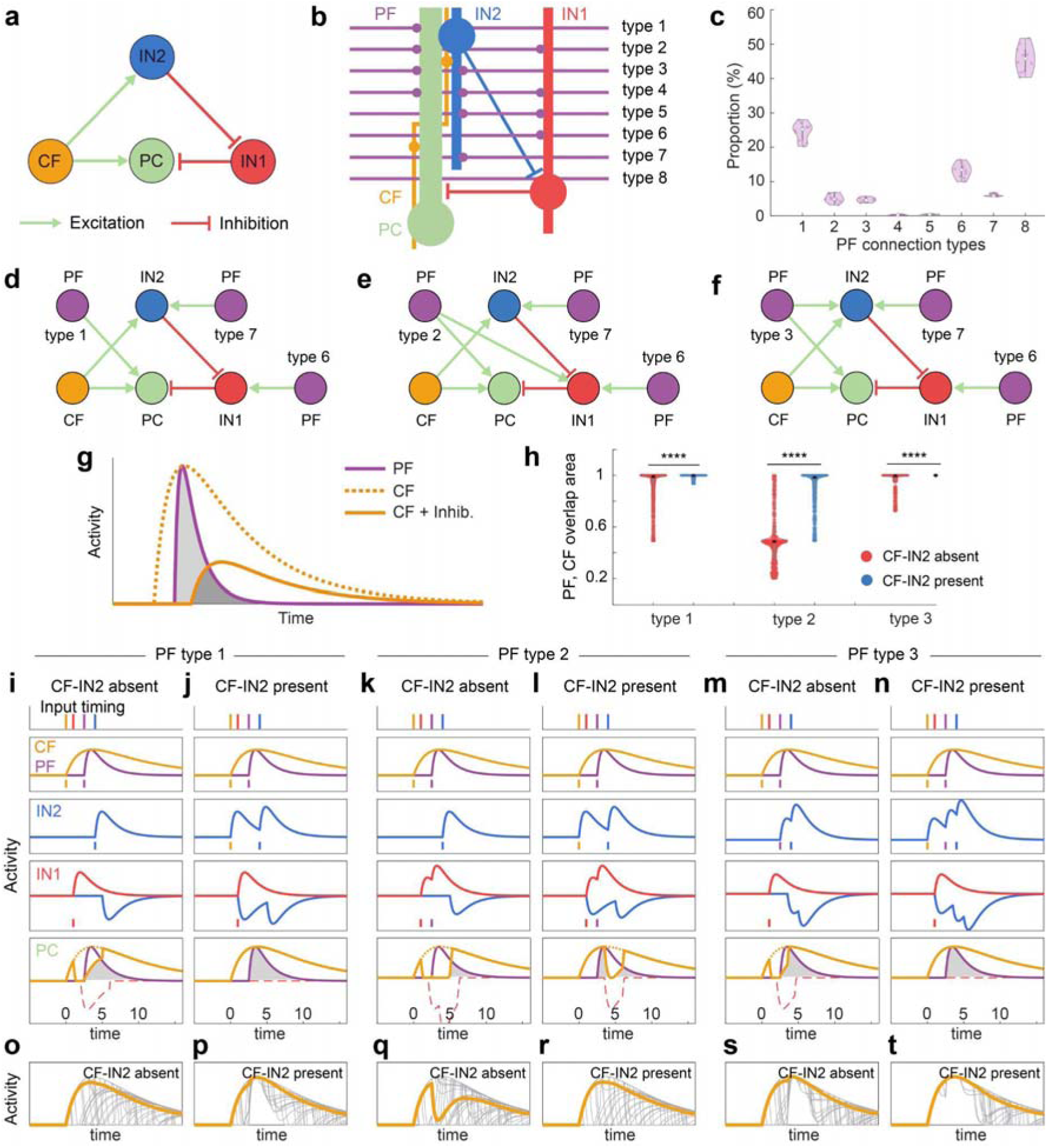
Disinhibition supports cerebellar motor learning. **a,** The minimal feedforward disinhibitory circuit. **b**, PF connection types in the dendritic field of a PC. **c**, The proportions of PFs belonging to each connection type. A dot in the violin plot represents data for one PC. **d–f**, Diagrams of disinhibitory circuit structures with the three PF connection types. **g**, The simultaneity of PF and CF inputs to PCs, which was measured by the overlapping area (grey shades) under PF and CF activity curves. The CF activity and the simultaneity decrease by inhibition. **h**, Distributions of the overlapping area for random PF inputs to INs with or without CF-IN2 connections, summarising the results from **(i–n)** below (*n*=10^4^, ****=p<10^-4^, Mann–Whitney U test). **i–n**, In one simulation run, CF and type-1 PF inputs at fixed timings and type-6 and 7 PF inputs at random timings (top) induce cascading excitatory and inhibitory activities on the IN2 (third), IN1 (fourth), and PC (bottom). The overlapping area under the CF and PF activity curves was calculated. The experiment was repeated with the same input timings for the three circuits **(d–f)** with **(j, l, n)** or without **(i, k, m)** the CF-IN2 connection. **o–t**, The net CF input to the PC after the cascade in each experiment (grey traces) and the mean (orange) for the case **(i–n)**, respectively.

In the circuit diagram, PFs, which are known to synapse onto INs and PCs to form feedforward inhibitory connections^23,24^, are missing. To add PF connectivity in the diagram correctly, we investigated the wiring specificity of PFs to PCs and INs and the relation between the feedforward inhibitory and disinhibitory circuits. In particular, we measured the proportions of PFs belonging to 8 different connection types determined by the connectivity of a PF to IN1s, IN2s, and PCs (Methods and Fig. 5b, c).

A total of 45.8% of PFs (type 8) passed through the dendritic field of a PC without any connection. For the rest, types 1, 6 and 7, which connect to only one postsynaptic type, were most common, comprising 43.8% of PFs. As types 2 and 3 are secondary and types 4 and 5 are negligible, only 10.5% of PFs connected to more than one type. Among all the PFs in the dendritic field of a PC, 34.5% (types 1–4) connected to the PC. Lastly, only 15.5% of the PFs that connected to PCs formed feedforward inhibitory connections (fraction of types 2, 4 over 1– 4), which appears minor compared to the physiological importance^23,24^.

We also directly counted for each PF connection type the number of different INs that a single PF connects to (Methods and Extended Data Fig. 4). The results demonstrated that any type of PFs routinely connected to the minimum possible number of INs for the type. These proportions indicate that PCs, IN1s, and IN2s can be considered to have independent PF inputs in the first-order approximation. Therefore, we summarised the circuit diagram into three representative structures, with one PF selected from the types 1–3 and two more PFs connected to IN1 and IN2, respectively (Fig. 5d–f).

What can be inferred on the functional role of the disinhibition of these circuits? We claim that disinhibition resolves the “paradox of inhibition” and supports cerebellar plasticity by ensuring the LTD of PF-PC synapses^15^. Whenever there is CF input to a PC, the CF-IN2-IN1 connection silences IN1-PC inhibition, and the simultaneity of CF and PF inputs is preserved. This hypothesis was tested by a numerical simulation, as described in the next section.

How can CF-IN synapses play such a critical role when the number is so small? The answer is that divergent connectivity of the disinhibition enables the universal action of CF-IN synapses. To quantify the divergence, we introduced the disinhibition coverage to measure the proportion of IN1s presynaptic to a PC are disynaptically connected by the PC-innervating CF via IN2s. The mean value was 81%, and similar results were obtained when the number of neurons in the definition was replaced by the total number of synapses or total synaptic area (Supplementary Data 2e).

### Computational model

We tested the hypothesis with a computational model simulating the neural activities of neurons in the three different circuit structures (Fig. 5d–f). The model quantified the chances of PF-PC LTD by estimating the amount of simultaneous input from the CF and PF to the PC. In the conceptual plot, PF and CF inputs were shown when inhibition is present or not, and the simultaneity of the inputs was measured by the overlap area under the curves (Fig. 5g). The overlap area was decreased by inhibition, which reflects that the simultaneity between CF and PF inputs decreases, as do the chances for PF-PC LTD.

The simulation was conducted as follows (Methods). All the neuronal activities in the circuits were initiated by the activities of one CF and three PFs, where the CF and type-1 PF activity onset timings are fixed and the type-6 and type-7 PF activity onset timings are randomly varied (Fig. 5i–n, top). The excitatory activities of PFs and CFs are described by double-exponential functions, which increase quickly and decrease slowly, modelling the firing rate or intracellular calcium concentration over time during a burst of spikes. The initiating CF and PF activities are cascaded down the circuits through synaptic transmission.

In general, the activity of the presynaptic neuron is assumed to become an input to the postsynaptic neuron as it is. Furthermore, the activity of a neuron is calculated from the summation of all the inputs where the excitatory and inhibitory inputs have positive and negative signs, respectively, and then the summation is rectified to remove the negative values and then shifted by a time constant.

For the PC, we are interested in the inputs to the PC rather than the activity. The CF and type-1 PF activities became the excitatory inputs to the PC with complete overlap (Fig. 5i–n, second). The complete overlap was changed by the inhibitory inputs. Type-7 and type-3 PF activities became excitatory inputs to the IN2, as does the CF activity when the CF-IN2 connection is present, which are summed to form the IN2 activity (Fig. 5i–n, third). Type-6 and type-2 PF activities became excitatory inputs to the IN1, while the IN2 activity became an inhibitory input, which together established the IN1 activity after rectification (Fig. 5i–n, fourth). IN1 activities became inhibitory input to the PC, which were subtracted from the direct CF excitatory inputs and then rectified to yield the net CF excitation. Finally, the overlap between the direct PF excitation and the net CF excitation could be calculated (Fig. 5i–n, bottom).

When the CF-IN2 synapse was absent, the time traces of the net CF input inhibited by IN1 exhibited a decrease compared to those of intact CF input (Fig. 5o, q, s). This led to a decrease in the overlap between CF and PF inputs to the PC (Fig. 5h, red dots). This indicates that the PF-PC LTD was disrupted by the inhibition. When CF-IN2 synapse was present, on the other hand, the decrease was only limited because inhibition was less functional due to disinhibition (Fig. 5p, r, t). This led to a smaller decrease in the overlap compared to when the CF-IN2 synapse was absent (Fig. 5h, blue dots).

The efficiency of disinhibition was the greatest for the circuit with type-3 PF because it has an additional disinhibitory connection and is the least for the circuit with type-2 PF because it provides extra inhibition (Fig. 5h). Despite the difference in efficacy, the results show that disinhibition supports PF-PC LTD in all three circuit structures and therefore in the entire cerebellar circuit, which consists of a combination of the three.

## Discussion

We raised a question on cerebellar motor learning theory that frequent inhibition may disrupt LTD at PF-PC synapses. In search of information to answer this question, we densely reconstructed the CML and mapped the microcircuits from the 3D EM images. We discovered CF-IN synapses, as CML neurons are organised to enhance the chances of making CF-IN contacts. We analysed the wiring specificity to redefine IN cell types into IN1, which primarily innervates PCs, and IN2, which primarily innervates IN1 and is innervated by CFs. Such wiring specificity yields the disinhibitory circuit structure that suppresses inhibition and supports the LTD, which we verified with a computational model.

Owing to the ostensive structural regularity and simplicity^15,21^, the intricateness and exquisiteness of the cerebellar circuits has only been recognised recently^10,44,45,51^. Advances in experimental technology and analysis tools to investigate neuronal organisation and connectivity, particularly the computational reconstruction of large-scale 3D EM image volumes, are revealing hidden mysteries. Further investigations on the same data shared by the published works^10,51^ may already lead to new discoveries. This work showed that studies on the structures provide insights into connectivity and studies on connectivity shed light on function.

With minor modifications to the computational model, various conditions could be simulated. For example, LTP induction by a [Ca^2+^]_i_ decrease^7,8^ could be simulated if CF excitatory input and inhibitory inputs to PCs are summed without rectification. The model has limitations as well due to many assumptions. We assumed that IN input inhibits CF input but does not inhibit PF input, to evaluate the impact of globally varying CF inputs on given PF inputs for local PF-PC synapses. However, point neurons of the model cannot simulate local variability or timing differences due to conduction speed^52^. While the activity of a presynaptic neuron was assumed to be transmitted to a postsynaptic neuron without any change, the validity of this assumption has yet to be theoretically supported. More detailed models that can explain the various physiologies of CML are expected in future studies.

The discoveries and proposed theory of this work could be cross-validated by molecular and cellular biological experiments. CF-IN synapses may be validated by labelling the IN2s through the injection of a transsynaptic marker to inferior olive neurons, for instance. CF-IN synapses do not necessarily deny CF-IN communication through glutamate spillover. Individual cases that have been suggested as evidence for glutamate spillover and for the nonexistence of CF-IN synapses need to be reexamined^28,41^. The proposed theory could be experimentally tested when genetic access to IN types are acquired. Disturbing IN1 alone or both INs would not disrupt LTD, as it keeps the CF and PF inputs intact^53^. Disturbing IN2 alone would disrupt LTD, as it destroys the disinhibitory pathway. A recent study showed the gating function of INs for cerebellar learning by optogenetically suppressing INs during the vestibulo-ocular reflex^15^.

Generally, LTP is induced by simultaneous presynaptic input and postsynaptic depolarisation in various brain regions^54,55^. Cerebellar LTD is the reverse since LTD is induced by simultaneous PF input and CF input which depolarises and fires a PC. It has been shown that LTP is disrupted when postsynaptic depolarisation is suppressed by inhibition^42,56,57^ and is intact when postsynaptic depolarisation is maintained by disinhibition^16,17^. Our findings suggest that inhibition and disinhibition are critical for both cerebellar LTD and LTP in various brain regions by the same physiological principle.

## Methods

### Experimental animal

Animal experiments were conducted in accordance with the National Institutes of Health Guidelines for the Care and Use of Laboratory Animals and were approved by the Institutional Animal Care and Use Committee (IACUC) of the Korea Brain Research Institute (IACUC-16-00017). An 8-week-old male C57BL/6J mouse (Jackson Laboratory) was maintained under alternating 12-12 hours of light-dark conditions, with an ambient temperature of 21±1°C and free access to food and water.

### Sample preparation & EM imaging

The brain of the mouse was perfused and fixed for 16 hours at 4°C in 0.15 M cacodylate buffer (pH 7.4) containing 2% paraformaldehyde and 2.5% glutaraldehyde. The brain was sliced into 200 μm-thick sagittal sections, which were used for the dissection of cerebellar lobule IV/V. We prepared the sample for serial block-face scanning electron microscopy as previously described^58^. The specimen was mounted on an aluminium rivet with Epon 812 resin (Electron Microscopy Sciences), which included 7% (weight/volume) carbon black powder to enhance conductivity^59^. The tissue was sectioned and imaged with a Merlin VP field emission scanning electron microscope (Carl Zeiss Microscopy LLC.), equipped with 3View2XP in-chamber ultramicrotome technology and a backscattered electron detector (Gatan, Inc.). We used the montage function of Digital Micrograph (ver. 2.31.734.0, Gatan Microscopy Suite) to capture 2×3 rectangular tiles. Each tile was scanned with the following parameters: 30 μm aperture, acceleration voltage of 2.5 kV, pixel dwell time of 3.0 μs, resolution of 5,000 by 5,000 pixels with 10∼20% overlap between tiles, and XY resolution of 12 nm with a Z-depth of 50 nm. A total of 1,020 section images (6,120 tiles) were acquired over 128 hours. The first 20 sections were discarded because of defects.

### Image registration

We registered the images using ImageJ with TrakEM2 plugin^60^ and custom MATLAB codes. Each of the 6 stacks was aligned by affine transformations. To compensate for the defective overall translation and rotation that commonly occur for many-sectioned stacks, the affine transform was adjusted by the amount of the polynomial fitting of each transform parameter. Six stacks were positioned by rigid transformations and then the images in the stacks were deformed by elastic transformations. The transformations were determined using both inter-tile and inter-section information. Finally, the tiles on a section were welded for the locations where tiles overlapped by selecting the pixel of a tile whose distance from the centre of the tile was the closest.

### Segmentation and proofreading

We used convolutional neural networks^61,62^ and modified watershed algorithm^9^ to segment the registered images into supervoxels. The training set was taken from seven locations which include various neural structures. The dense segmentation and EM images were divided into subvolumes for convenient human proofreading. The segmentation errors were proofread by human workers following the same procedure introduced in a previous work^9^, where two workers independently proofread the same neuronal fragment and the third worker made the final decision. Proofreading took approximately 5 years by 5 workers on average. To improve segmentation accuracy, the network was replaced once, and each network was retrained and watershed parameters were updated several times during the proofreading period.

### Cell type classification

We classified the reconstructed neurons and neuronal fragments into 4 types (PC, CF, PF, and IN) or undetermined by visually inspecting the morphology on the 3D renderings and cellular structures in the EM images (Supplementary Data 1–4, Fig. 2b–g)^39^. The classical description of the morphologies of the cell types was sufficient as the criterion of the classification for both cells with somata and fragmented processes without somata^11,21^.

PC dendrites that branch out from the thickest trunks to thinner arbours laminate parasagittally and have dense spines. IN dendrites are thinner than the thinnest arbours of PC dendrites, and they also roughly laminate parasagittally while radiating from the cell body on the laminae. The IN axons arborise in arbitrary directions with occasional parasagittal alignment and innervate PC and IN dendrites. Some IN axons have a prominent PC perisomatic basket structure. Granule cell axons are mostly straight PF fragments perpendicular to the laminating planes of PCs, while a small fraction of granule cell axons include both the parts for ascending axon and for PF, forming “T-shaped” arbours, all of which we referred to as PFs. CFs have numerous large presynaptic boutons, and they arborise on the surface of the dendrites of a PC.

The dendritic and axonal fragments were distinguishable by the presynaptic boutons with neurotransmitter vesicles. Fragmented processes that were too small to provide sufficient morphological cues were sorted as an undetermined type. Although the fragmented processes of granular layer cell types (Lugaro cells, unipolar brush cells, globular cells, and candelabrum cells) may have been misclassified into INs or undecidable, those types are minor^13,63,64^ and the impact on the analyses would be minimal.

### Estimation of reconstruction completeness

The volume reconstruction completeness, the number of voxels in the reconstructed segments over the number of voxels of the entire volume of the dataset, yielded *m*=0.6025. The non-reconstructed voxels mostly belonged to glia. We also estimated the reconstruction completeness of each cell type, i.e., the number of reconstructed cells over the estimated total number of cells in the dataset for each type, by random samplings. We took five subvolumes that were (6.1 µm×6.1 µm×6.4 µm) in size containing neuropil. For each subvolume, *N*=100 foreground voxels belonging to reconstructed cells and *N*=100 background voxels not belonging to reconstructed cells were randomly sampled. For the foreground voxels the cell types were already classified. For the background voxels, the cell type was speculated from the supervoxels in the subvolumes without proofreading. The reconstruction completeness for cell type *t* was approximated by *R*(*t*)=*mv*_f_(*t*)/(*mv*_f_(*t*)+(1-*m*)*v*_b_(*t*)), where *v*_f_(*t*) and *v*_b_(*t*) are the number of foreground and background voxels of the given cell type *t* among the sampled 5*N* voxels, respectively (therefore, Σ_t_*v*_f,b_(*t*)=5*N*), and *m* is the volume reconstruction completeness.

### Separation of neuronal processes and somata

We separated the volumetric representation of the main neurons into somata, dendrites, and axons when necessary, e.g., when calculating the angular volume distributions of only the IN dendrites along the PC dendrites (Fig. 3e). PC somata were separated in units of subvolumes by visually identifying the subvolumes that contained the somatic portion of the neuronal segment. For INs, because the size of the subvolumes was too large compared to the size of the somata, we used a (group of) bounding box(es) enclosing the soma. We determined the coordinates of bounding boxes of the somata by visual inspection and manual annotation. The axonal arbours of INs (which sometimes originated from a dendrite) were able to be identified by the presence of presynaptic boutons and neurotransmitter vesicles and separated by the units of subvolumes. A new reconstruction volume was created where each neuron was represented by three different segments for the somata, dendrites, and axon.

### Skeletonisation

For the PCs, the dendritic shaft skeletons were obtained through two steps to manage the frequent self-touches between dense spines. First, the dendritic spines were removed by 3D morphological operations. Erosion on the PC dendrites with a fixed structural element size allowed the removal of spine necks with few errors and occasionally removed the thinnest parts of the dendritic shafts, leaving only the spine heads and a few pieces of disconnected dendritic shafts. We then dilated the large dendritic shaft pieces, whose sizes were over ten times the spine head size, to remerge the split dendritic shaft pieces into one. Second, the skeletons were calculated from the spine-removed dendritic shafts by applying the TEASAR algorithm^65^. For the INs, direct application of the TEASAR algorithm to the soma-removed dendritic segments skeletonised both the shafts and spines. The branches for shaft skeletons and spine skeletons were distinguishable using a length threshold.

### Angular volume distribution

The volumetric amount of neural processes for different cell types around PCs was measured for the surrounding space of PC dendritic shafts. The surrounding space was a 2 µm-thick tube with varying radii, which was obtained by subtracting the spine-removed dendritic shaft from the dilation of the shaft by 2 µm. Each neuronal voxel within the surrounding space was projected to the closest PC dendritic skeleton point and the displacement vector from the projected point to the original voxel point was defined. The angle of the displacement vector was measured relative to the local axis defined at each skeleton point by considering the following vectors. First, the tangential vector of the skeleton was considered. Second, the XY grid on the dataset was set, the median *z*-coordinate of the PC dendritic voxels in each grid cell was calculated, and the mean plane of the PC dendritic plank was determined from the least square fit of the grid points. Then, the horizontal vector, which lies in the mean plane and perpendicular to the skeleton tangent vector, was considered. Third, the vertical vector was defined as the outer product of the two vectors. The displacement vector was projected onto the plane that is perpendicular to the tangential vector and included the closest point on the skeleton, and its relative angle was measured with respect to the horizontal vector and vertical vector, which were assumed to be the *x*- and *y*-axis. We counted the number of voxels of each cell type in each angular bin to define the angular distribution of each cell type around the shaft of the PC dendrite. These procedures were conducted for each PC, and the result of each PC was averaged over all the PCs.

### Semiautomated synapse detection

We searched the synapses using the automated method that was developed specifically for this dataset^39^ and proofread the results for certain cell types. The contact segments between the neuronal segments were extracted from the reconstruction volume for the pairs of cell types, PF-PC (*n*=464,491), PF-IN (*n*=341,771), CF-PC (*n*=7,564), CF-IN (*n*=3,702), IN-PC (*n*=42,786), and IN-IN (*n*=36,931), and were tested to determine whether they were synapses. For the calculations, the dataset was divided into subvolumes, parallelly processed, and then reassembled to reduce the computation time. The parameters were tuned to minimise the missing of the synapses (false negative errors) and to allow the inclusion of incorrect synapses (false positive errors) to visually review the predicted synapses and manually exclude incorrect ones.

We reviewed every IN-related synapse of the main neurons comprising the adjacency matrix (Fig. 4a) since the number of such synapses in the training set was small. A few incorrect CF-PC synapses to the next PCs were discarded. We initially did not include the CF-IN contacts as CF-IN synapses were thought not to exist. Later, we added those contacts and conducted the synapse detection procedures again. It yielded a total of 128 CF-IN synapses out of 997 contacts between the main neurons, however, the result was specifically unreliable because CF-IN synapses were not included in the training set. Among the predicted 128 synapses, 61 were obvious errors such as those without vesicles or postsynaptic densities and those on the IN axon. We regarded the remaining 67 as synapse candidates and 4 human experts systematically investigated them by visual inspection. Each human expert graded the confidence level of the candidates on a 0 to 3 scale, and only the synapse candidates exceeding the total score of 6 were accepted as correct synapses.

The size of a synapse was approximated from the surface area of the contact segment. Considering the anisotropy in the voxel resolution (12 nm×12 nm×50 nm), the area of the voxel faces normal to the *x*- and *y*-axes is 600 nm^2^, and normal to the *z*-axis is 144 nm^2^. We counted the respective number of faces of contacting voxels along the *x*-, *y*-, and *z*-axis and calculated the summation.

### Measures of neuronal anatomy

We measured the distances of the centres of IN somata and synapses from the boundary plane between the PCL and CML. The boundary plane was determined from the centres of the interface between the PC dendrite segment and soma segment by finding the mean plane of the centre points using least square fitting. The actual boundary in the cerebellar foliae is a curved and irregular surface, however, the plane approximation was valid within the dataset, which is much smaller than one folium.

The dendrite density was defined by the total dendritic skeleton length divided by the 2D convex hull area^48^. The skeleton length was calculated by the total summation of the distance between adjacent skeleton nodes. The skeleton included both dendritic shafts and spines. The convex hull area was calculated from the 2D projection of the dendrite. The axon complexity was defined by the total number of skeleton branching points divided by the axon skeleton length. Since the definitions included normalisation, they are expected to be consistent for whole cells and the cell fragments cut-off by the volume boundary.

### PF connectivity analysis

We divided the *n*=33,900 PFs into 8 groups depending on connectivity with the set of IN1s, IN2s, and a PC, with synaptic convergence towards the PC. The dendritic field of a PC was found by dilating the dendrite segment by a spherical structure element with a 160-voxel (∼1.92 µm) radius. All the PFs that intersected the dilated segment volume were considered to pass the dendritic field of the PC. The types of these PFs were uniquely determined in the following order.

We checked whether a given PF had any connection to the PC and counted the number of IN1s and IN2s that the PF connected to. A PF that had any number of connections to all three types was deemed type 4. Among the rest, a PF that had any number of connections to both IN1 and IN2 was deemed type 5. Note that such rules for types 4 and 5 can become ambiguous when the PF connects to multiple IN1s and IN2s; however, the cases were rare and thus negligible (Fig. 5c). The rules for the other types simply followed the connection pattern (Fig. 5b), e.g., a PF that connected to the PC but not to any of the INs was type 1, and a PF that did not connect to the PC nor any of IN2s but connected to any of the IN1s was type 6.

The number of PFs in each type was counted, and the proportions were normalised by the number of PFs in the dendritic field of the PC. The mean proportions were obtained by the average over the 10 main PCs. The numbers of IN1s and IN2s connected by the PF were also used to calculate the proportions of PFs according to the number of connected INs (Extended Data Fig. 4).

### Computational model

The computational model simulation was conducted using custom MATLAB code. The activities of CF and PFs, which initiated the activities of all the other neurons, were modelled by double-exponential functions, *f*(*t*) = [*e*^-*t*/*m*^-*e*^-*t*/*n*^]^+^, where *m* and *n* are type-specific parameters tuning the duration of the activity and the [⋯]^+^ symbol indicates the half-wave rectification. For CF, *m*=8 and *n*=2; for PF, *m*=2 and *n*=0.5 were chosen to reflect the sustained input from CFs by redundant synapses that make PC fire the complex spikes^66^. The unit of time was arbitrary.

Introducing the different timings of activation, the activities of CF and PFs were

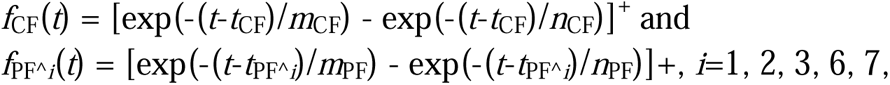

where *t*_CF_= 0 and *t*_PF^*i*_= 2.5 for types *i*=1, 2, 3 are fixed constants, while *t*_PF^*i*_= −5∼15 for types *i*=6, 7 are two different random numbers generated for every experiment of the simulation. The direct inputs of the activities *f*_CF_(*t*) and *f*_PF^*i*_(*t*) to PC without inhibition completely overlapped each other and the common area under the curves (Fig. 5i–n, second),

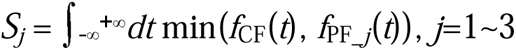

is 1.5. The subscript numbers denote the cases for circuit structures 1∼3.

The activities of postsynaptic INs were calculated by the summation of the presynaptic activities as excitatory or inhibitory inputs with a time delay, revealing that the firing of a neuron is a delayed output of the integration of the inputs. The time delay, □=1, was assumed to be identical in all neurons. For the case in which the CF-IN2 connection was present, the activities of the INs are given by

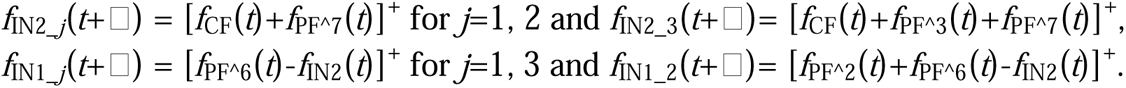

For the case in which the CF-IN2 connection was absent, the term *f*_CF_(*t*) was simply omitted.

Then, the inhibitory input from IN1s to PCs was subtracted from the CF to PC excitatory input, where the inputs are identical to the activities. Therefore, the net input from CFs to PCs was

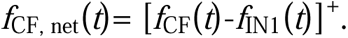

The overlap area was then *S* = ∫-∞^+∞^*dt* min(*f*_CF, net_(*t*), *f*_PF_(*t*)).

We repeated the simulation 10,000 times for random *t*_PF^6,7_ for each circuit structure. Using the same timing constants *t*_PF^6,7_, the simulations were repeated for the cases when the CF-IN2 connection was present and absent. The distributions of overlap areas were compared using Mann–Whitney U tests.

## Data availability

The 3D EM images, proofread segmentation, and neuronal 3D rendering dataset are publicly available at the BossDB (https://bossdb.org/) repository https://bossdb.org/project/park2023.

## Code availability

The codes and software used for this study are available in the following repository: https://github.com/cns-kim-lab/park_cerebellar_disinhibition/

## Acknowledgements

This research was supported by KBRI Basic Research Programs (16-BR-02, 23-BR-04-04) funded by the Korean Ministry of Science and ICT (MSICT), Basic Science Research Programs (2018R1A5A1060031) through the National Research Foundation of Korea (NRF) funded by the MSICT, and Basic Science Research Program (2019R1A6A1A10073079) through NRF funded by the Korean Ministry of Education. The EM images were acquired at the Advanced Neural Imaging Center of KBRI. We thank J. Shin, J. Kim, J. Yoon, H. Suh, D. Yoo, H. Kim, J. Kang, D. Cho, and C. Ham for proofreading and annotation. S. Yu, J. Park, S.-C. Yu helped verify the CF-IN synapses. We acknowledge discussions with J. C. Rah, J. Y. Kwon, and K. Yamamoto.

## Author contributions

C.P. conducted most of the scientific investigations including the computational simulation. J.G. segmented the images, using the network designed by K.L. and trained by K.L. and J.G. C.P. and J.G. classified the neurons and detected the synapses. J.G. and S.B. developed and managed the reconstruction pipeline software. S.B. registered the images and prototyped the skeletonisation that Y.I. completed. G.H.K. and S.H.L. prepared the sample and performed EM imaging of the sample under the supervision of K.J.L. Y.I. investigated the cellular organisation around the PC. C.P., S.Y., J.P., and S.-C.Y. proofread the synapses and verified the CF-IN synapses. J.G. trained and managed the proofreaders. M.S.K. advised on large data handling. C.P. and J.S.K wrote the manuscript with input from all the other authors. J.S.K. conceived the project and supervised the EM reconstruction and scientific investigation. K.J.L. and J.S.K. designed and led the overall study.

## Materials & Correspondence

Correspondence to <u>Jinseop S. Kim</u>.

## Additional information

Supplementary Information: This file contains full descriptions of Supplementary Data 1–5.

Supplementary Data 1: The open-source data repository https://bossdb.org/project/park2023 containing the 3D EM images, proofread neuronal segmentation, and the 3D rendering of neurons, which is provided together with the csv files for cell type and synapse annotations in a zip file.

Supplementary Data 2: Gallery of the main Purkinje cells in a PDF file, where each page contains the data of each cell for 3D renderings of the cell structure, input and output synapses and analysis results.

Supplementary Data 3: Gallery of the main interneurons in a PDF file, where each page contains the data of each cell for 3D renderings of the cell structure, input and output synapses and analysis results.

Supplementary Data 4: Gallery of the main climbing fibres in a pdf file, where each page contains the data of each cell for 3D renderings of the cell structure and input and output synapses.

Supplementary Data 5: Gallery of all the identified synapses between CFs and INs, showing 11 consecutive EM sections and the synapse scores given by experts.

**Extended Data Figure 1.**
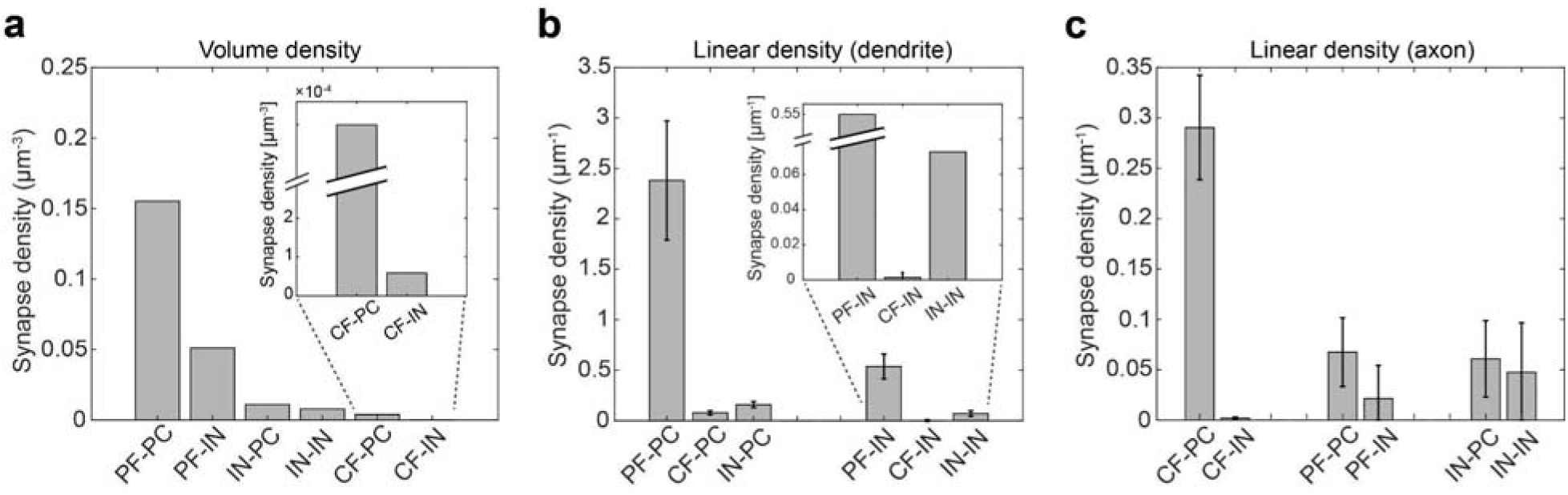
Volume and linear synapse density by synapse type. **a,** Volume synapse density by synapse type. **b,** Linear input synapse density along the PC and IN dendrites. **c,** Linear output synapse density along the axons of CFs, PFs, and INs.

**Extended Data Figure 2.**
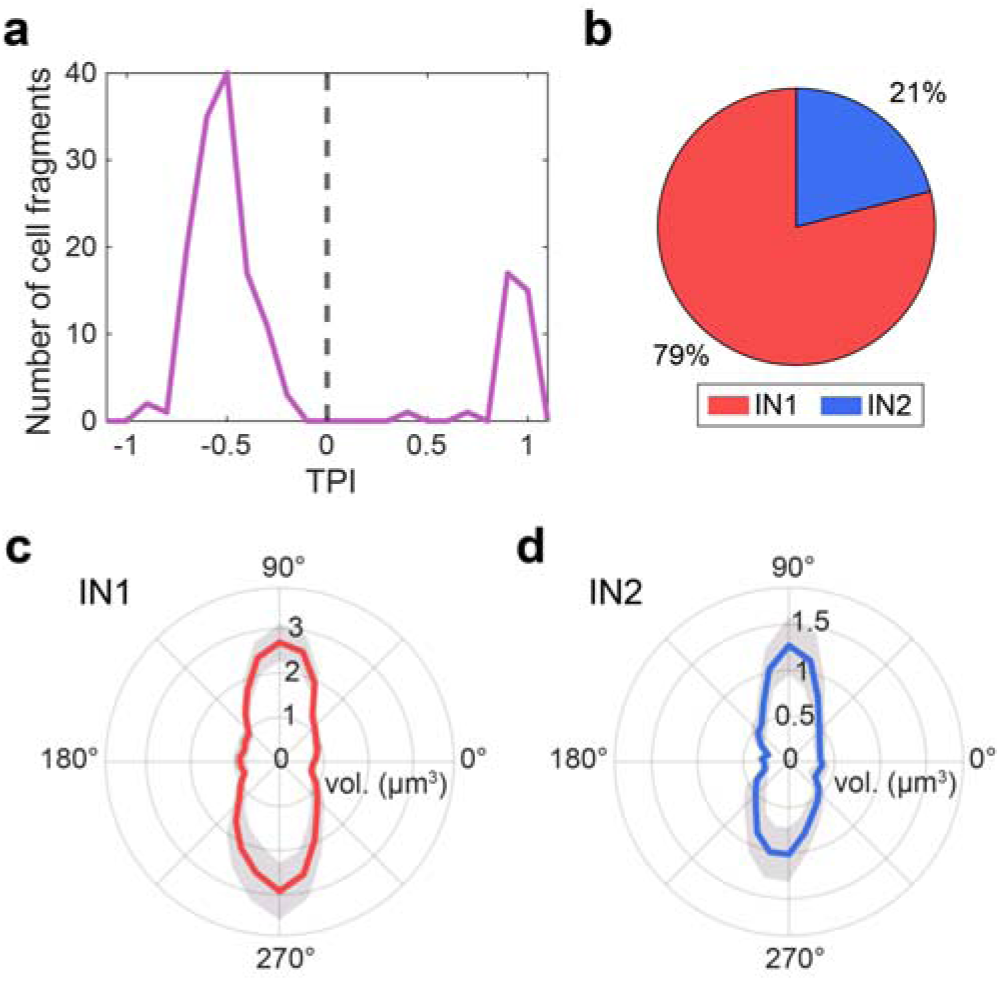
The proportion and angular volume distribution of IN types. **a,** The histogram of TPI of the IN fragments whose number of output synapses is greater than or equal to 30. **b,** The proportions of IN types. **c, d,** The angular volume distribution of the IN1 dendrites **(c)** and IN2 dendrites **(d)** around PC dendrites.

**Extended Data Figure 3.**
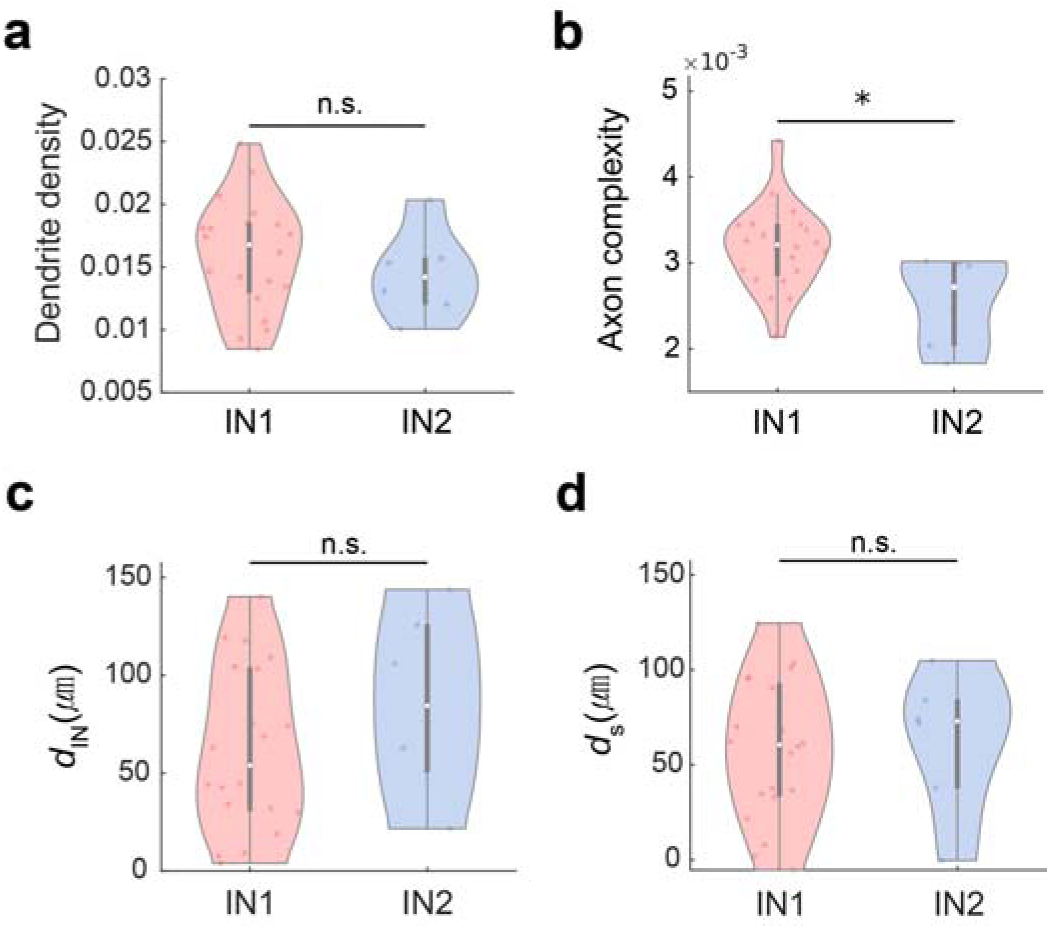
IN1 and IN2 comparisons. **a**, Violin plot showing the dendrite density of IN1(red, N=20) and IN2 (blue, N=6). p=0.447, Mann–Whitney U test. **b**, Axon complexity of IN1 and IN2. p=0.023. **c**, Soma distance from the PCL. p=0.260. **d**, Distance of outgoing synapses of INs to PCs measured from the PCL. p=0.563. For all data, the Mann–Whitney U test was used to assess significance.

**Extended Data Figure 4.**
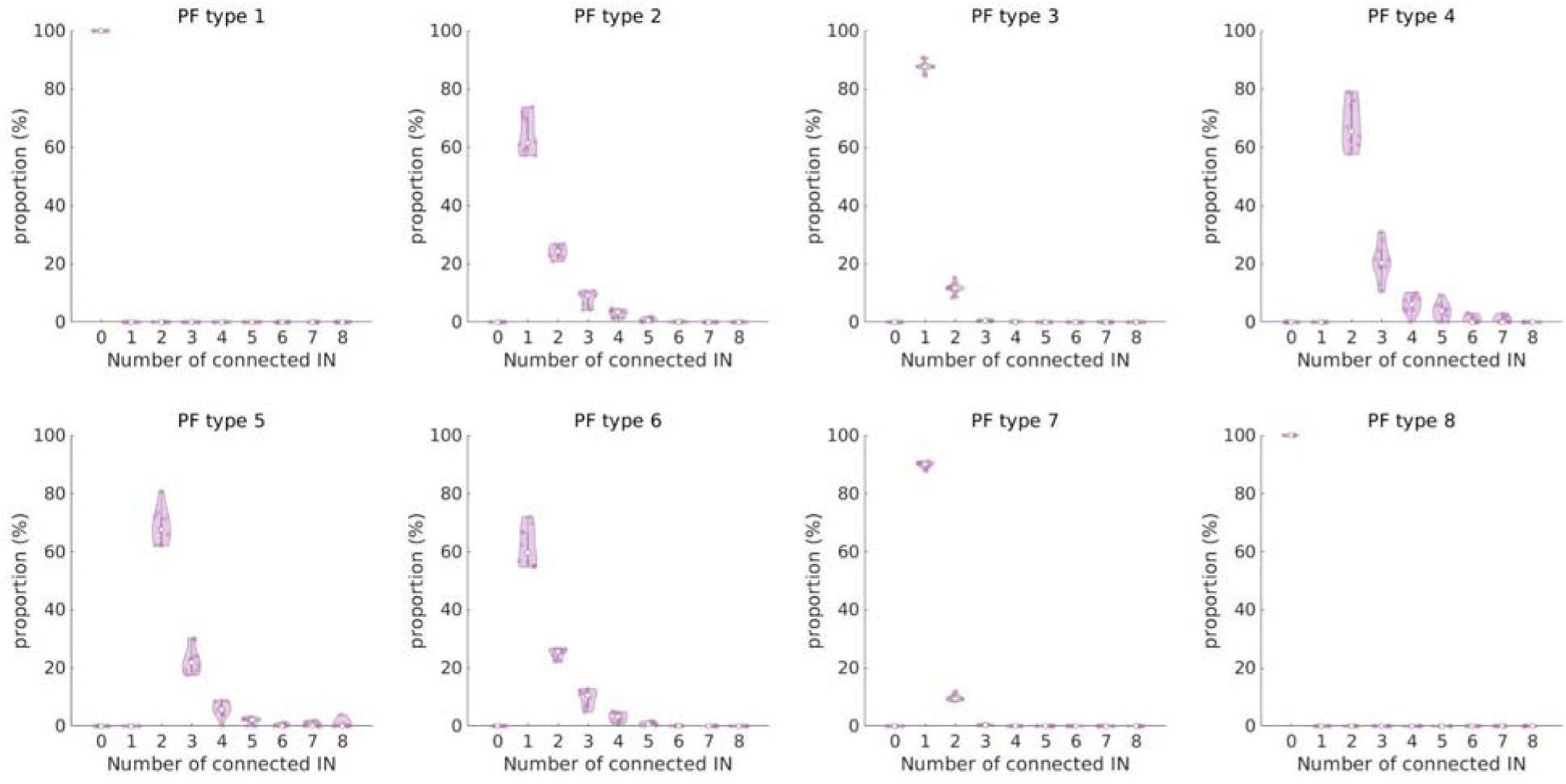
Proportion of PFs according to the number of connected INs. For the PFs belonging to each type, the number of connected INs to each PF was counted, and the proportions were measured.

