## Supplementary figures and images for "A cerebellar disinhibitory circuit supports synaptic plasticity"

### Supplementary Data 2

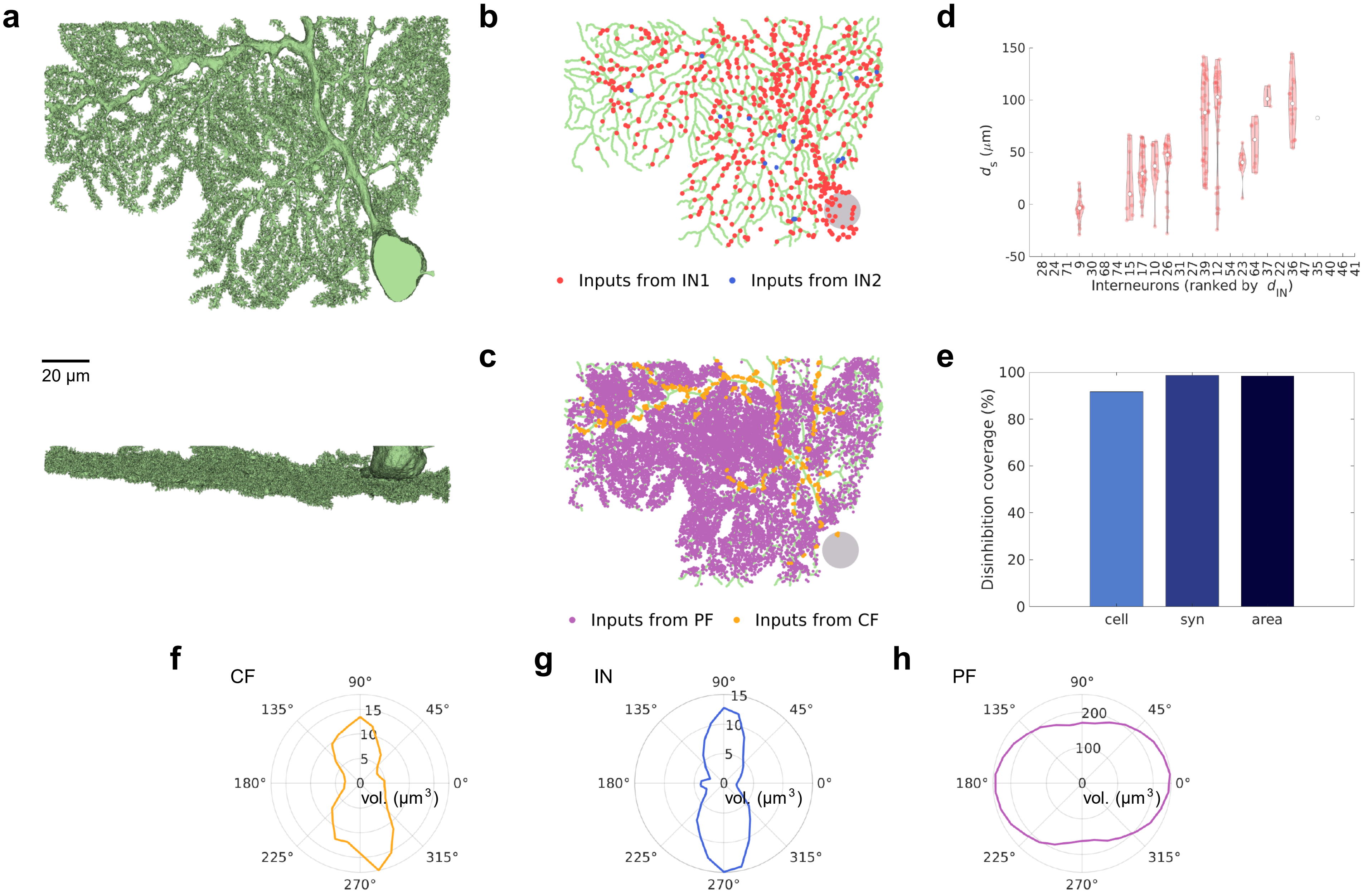

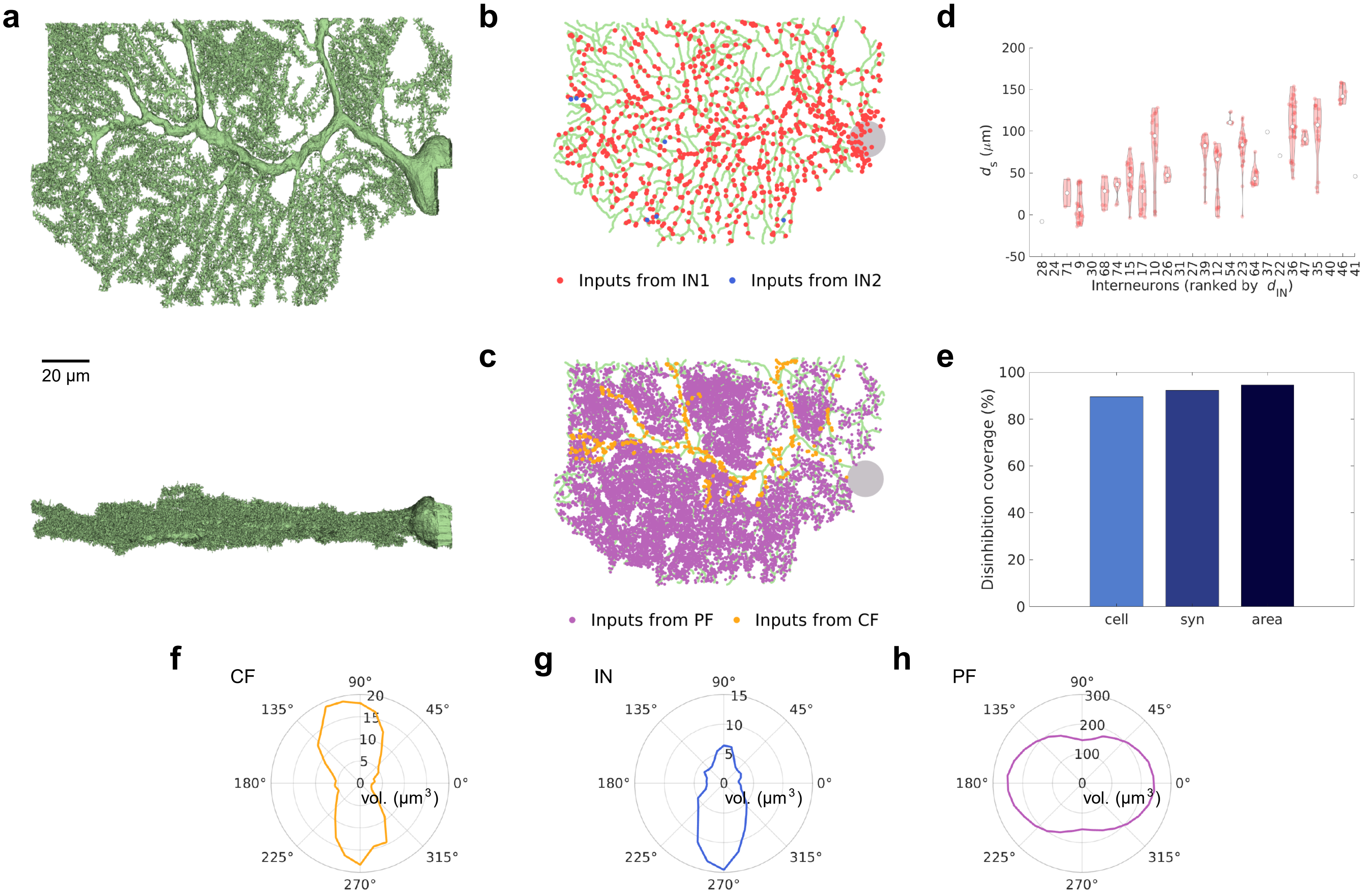

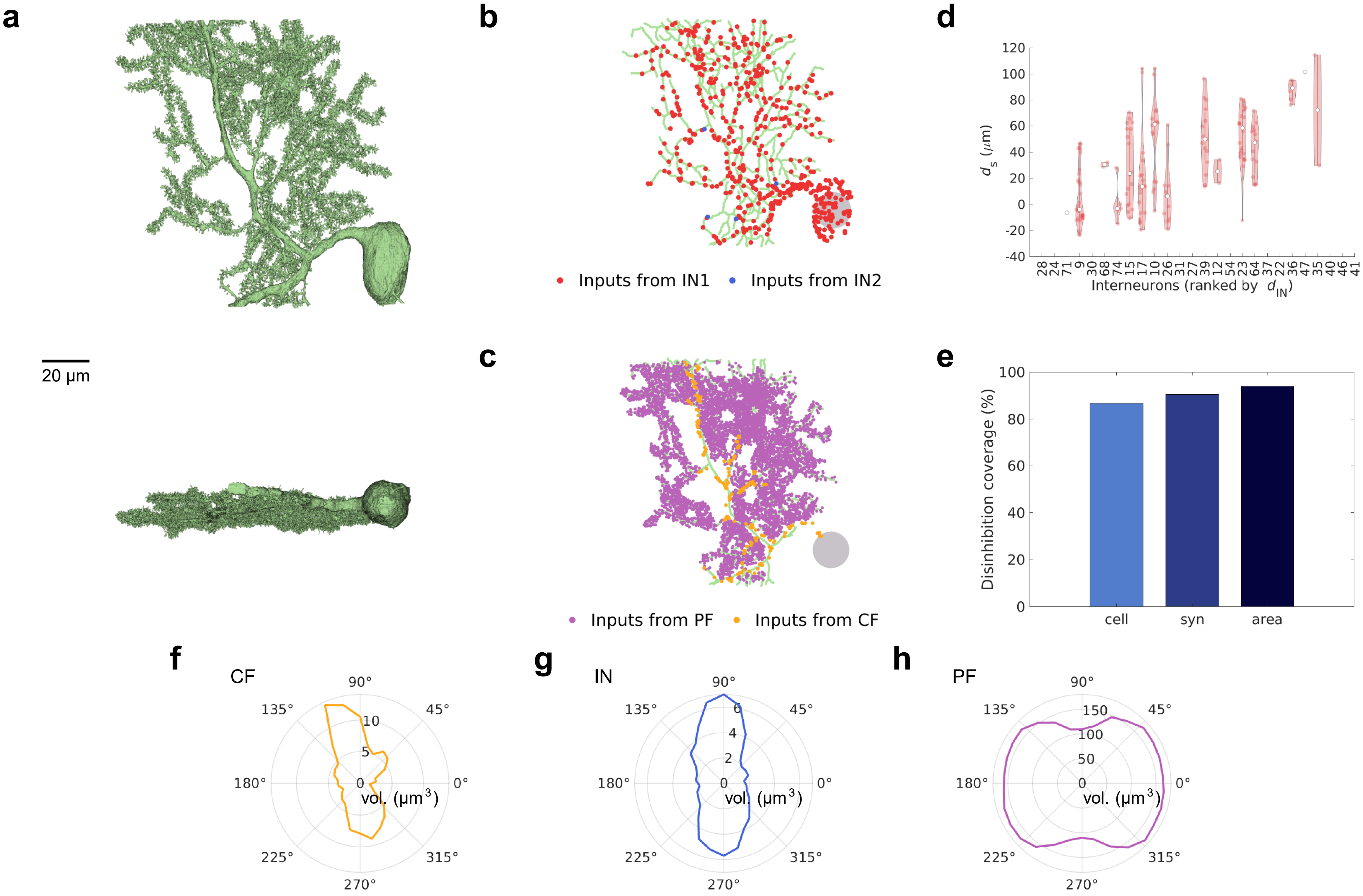

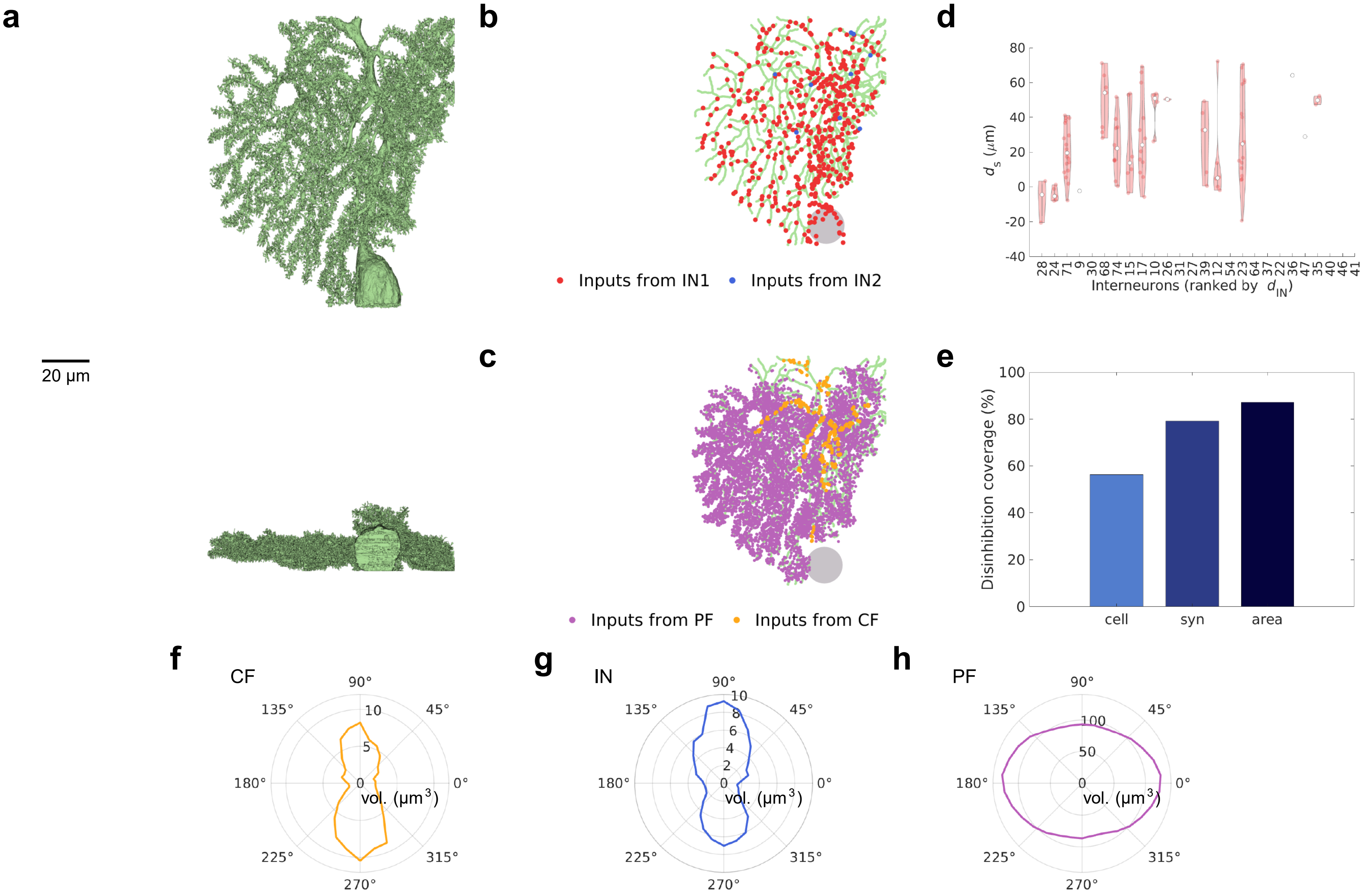

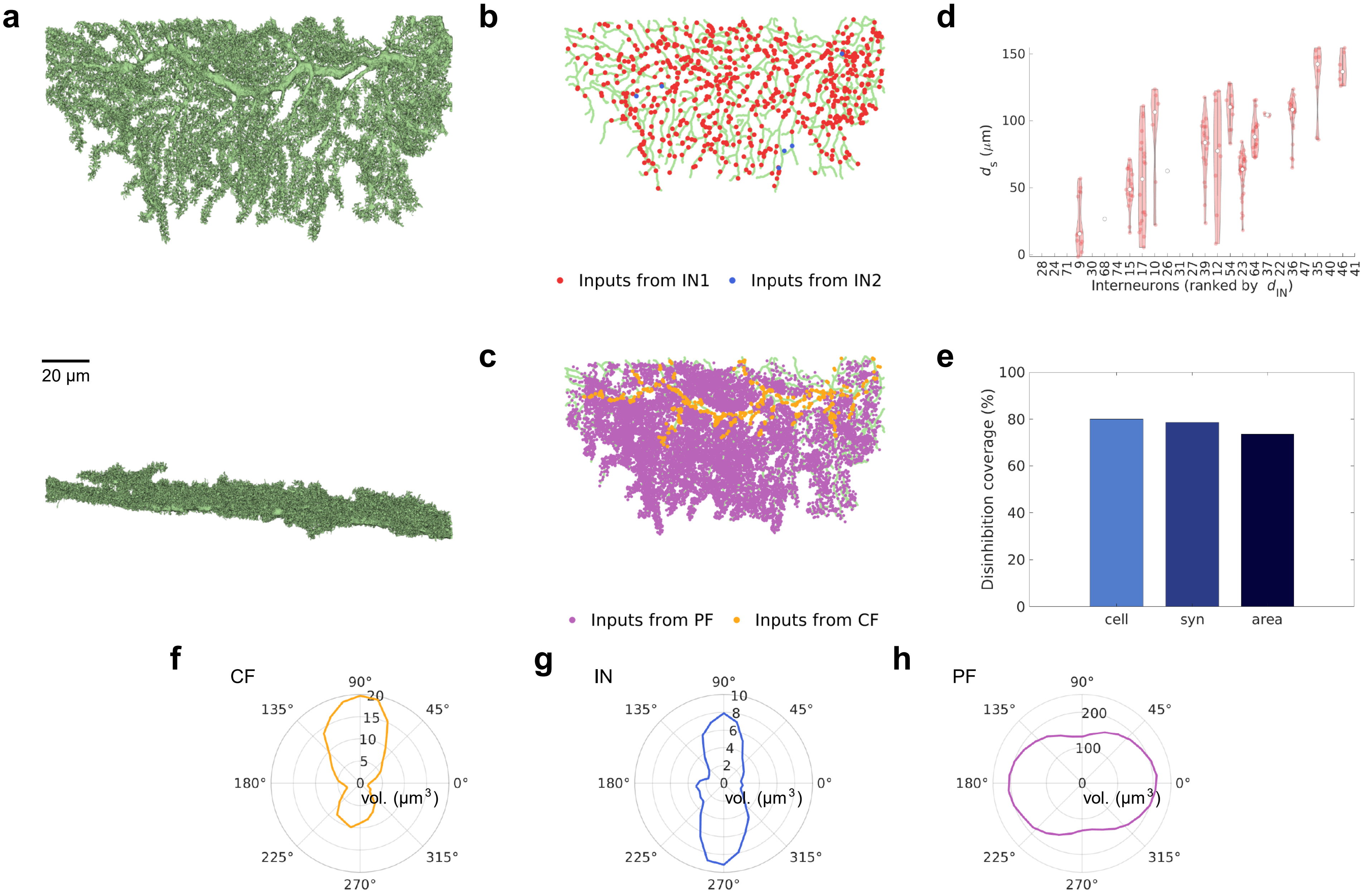

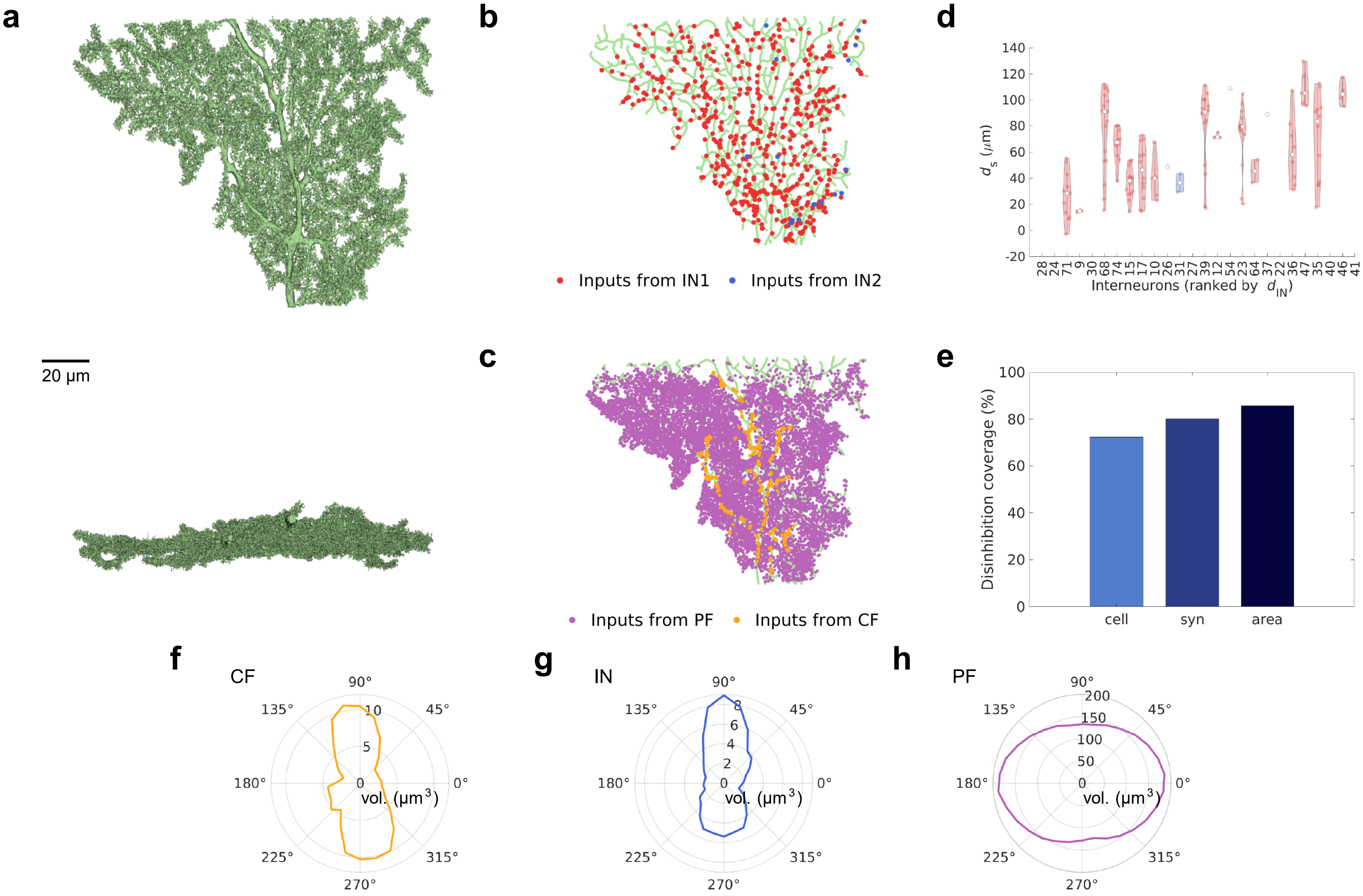

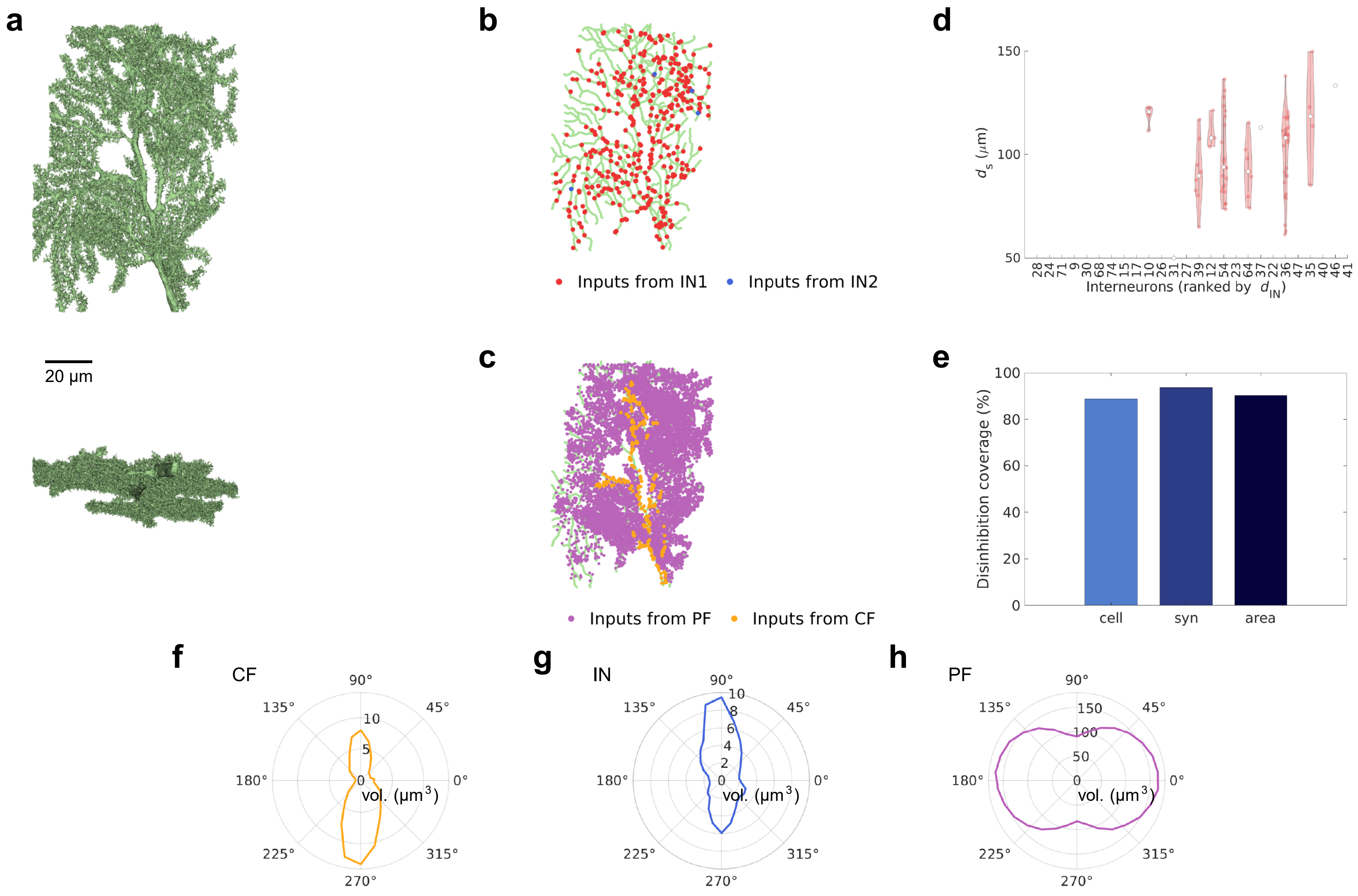

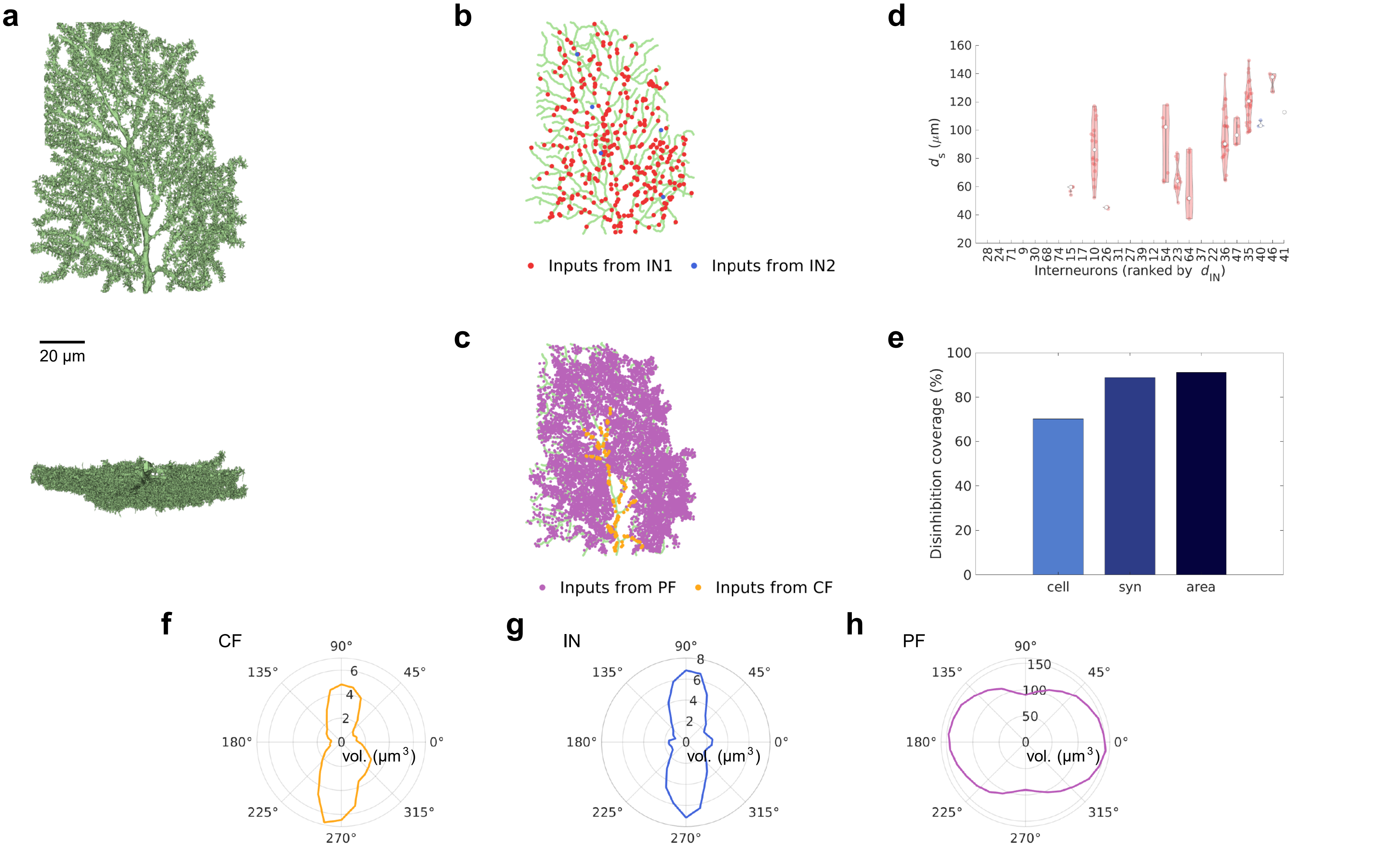

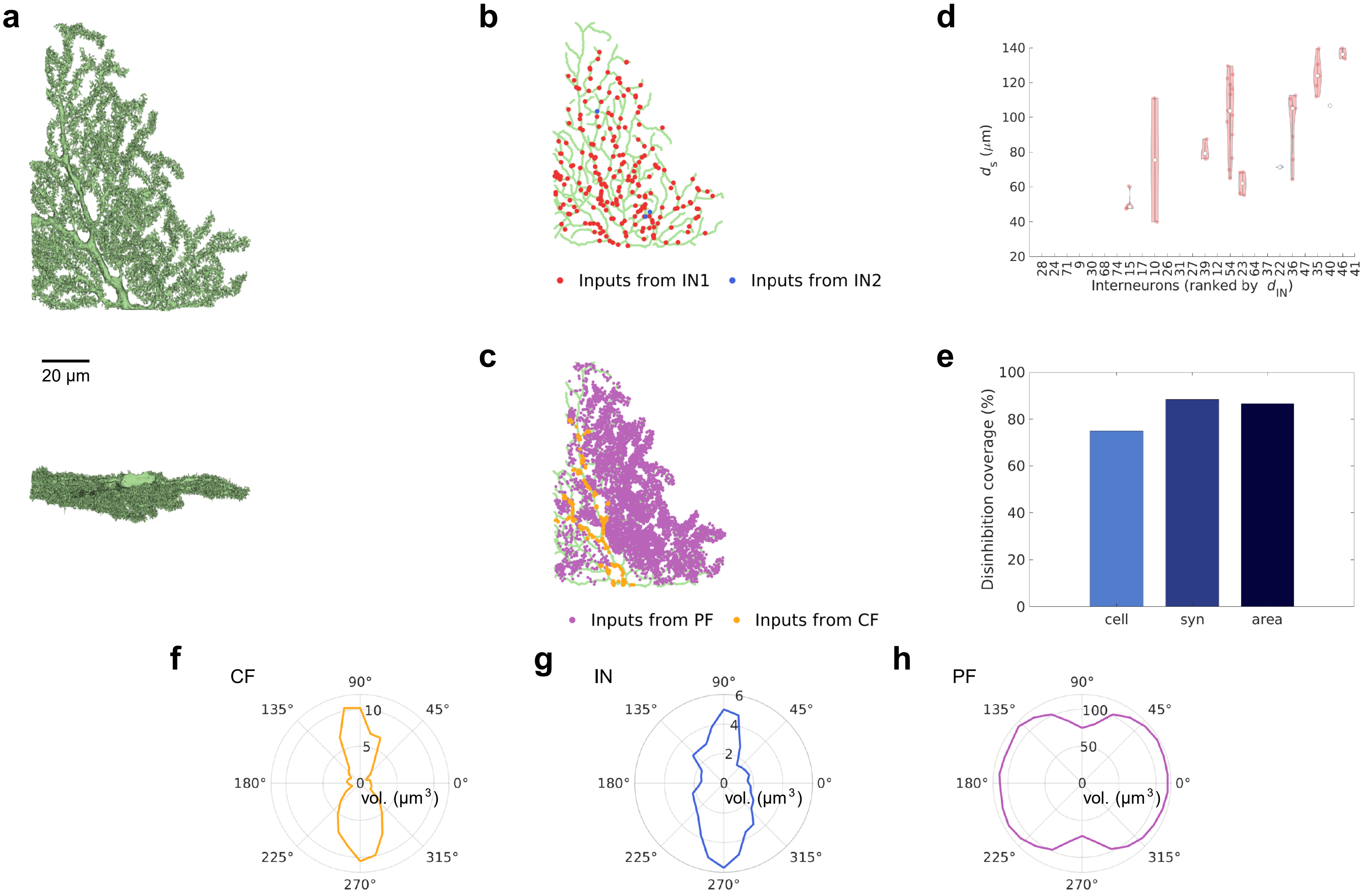

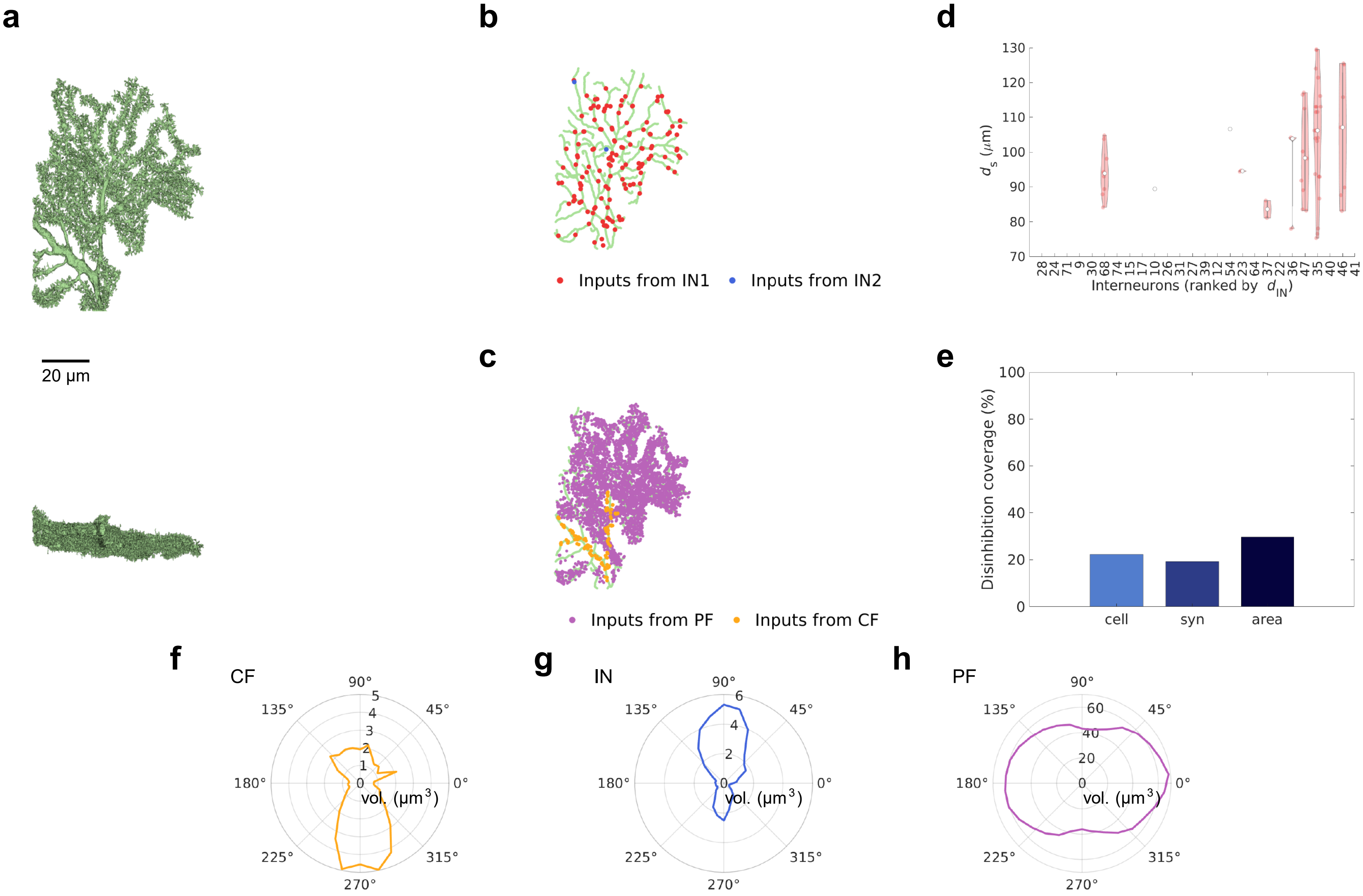

### Supplementary Data 4

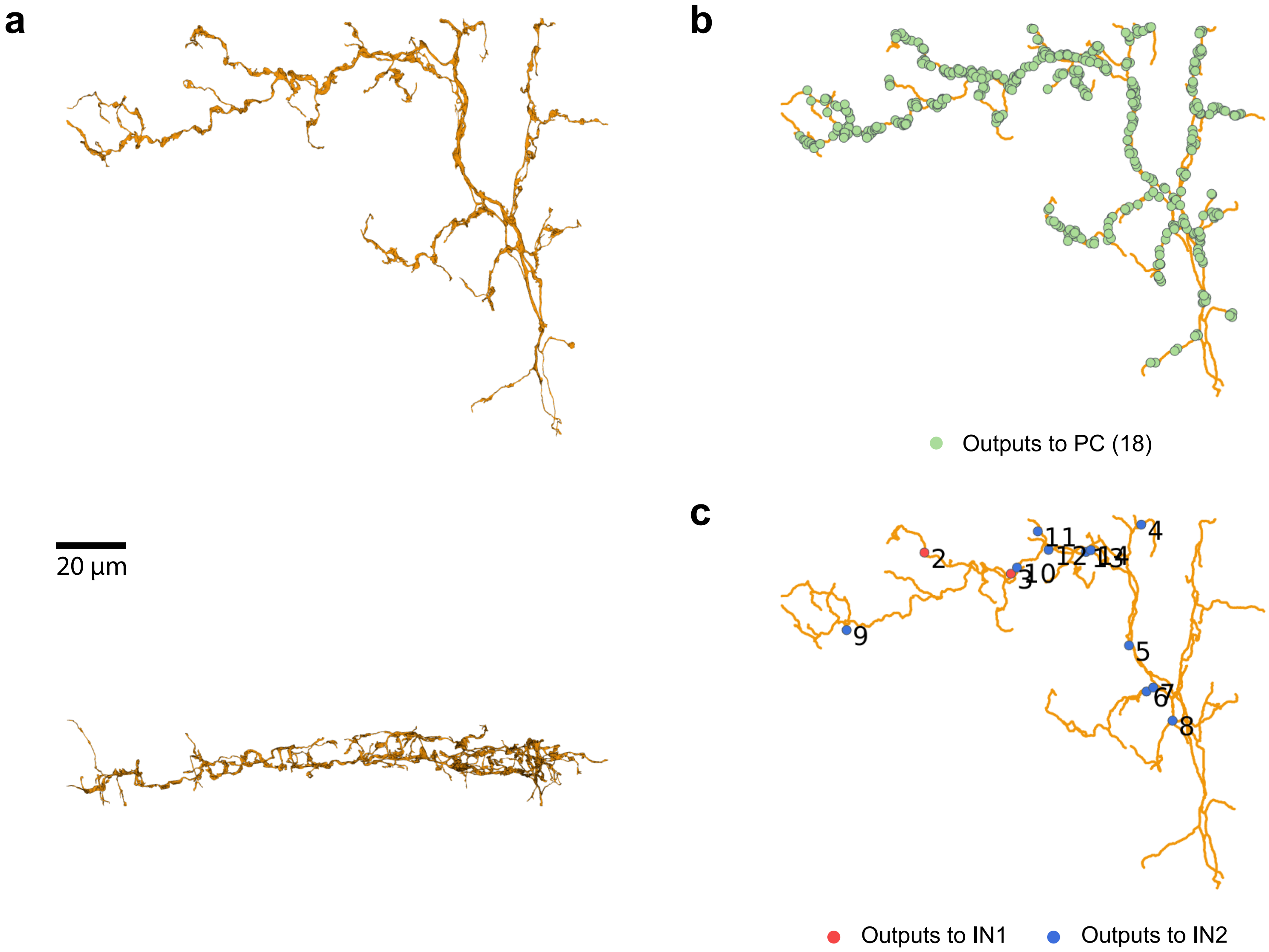

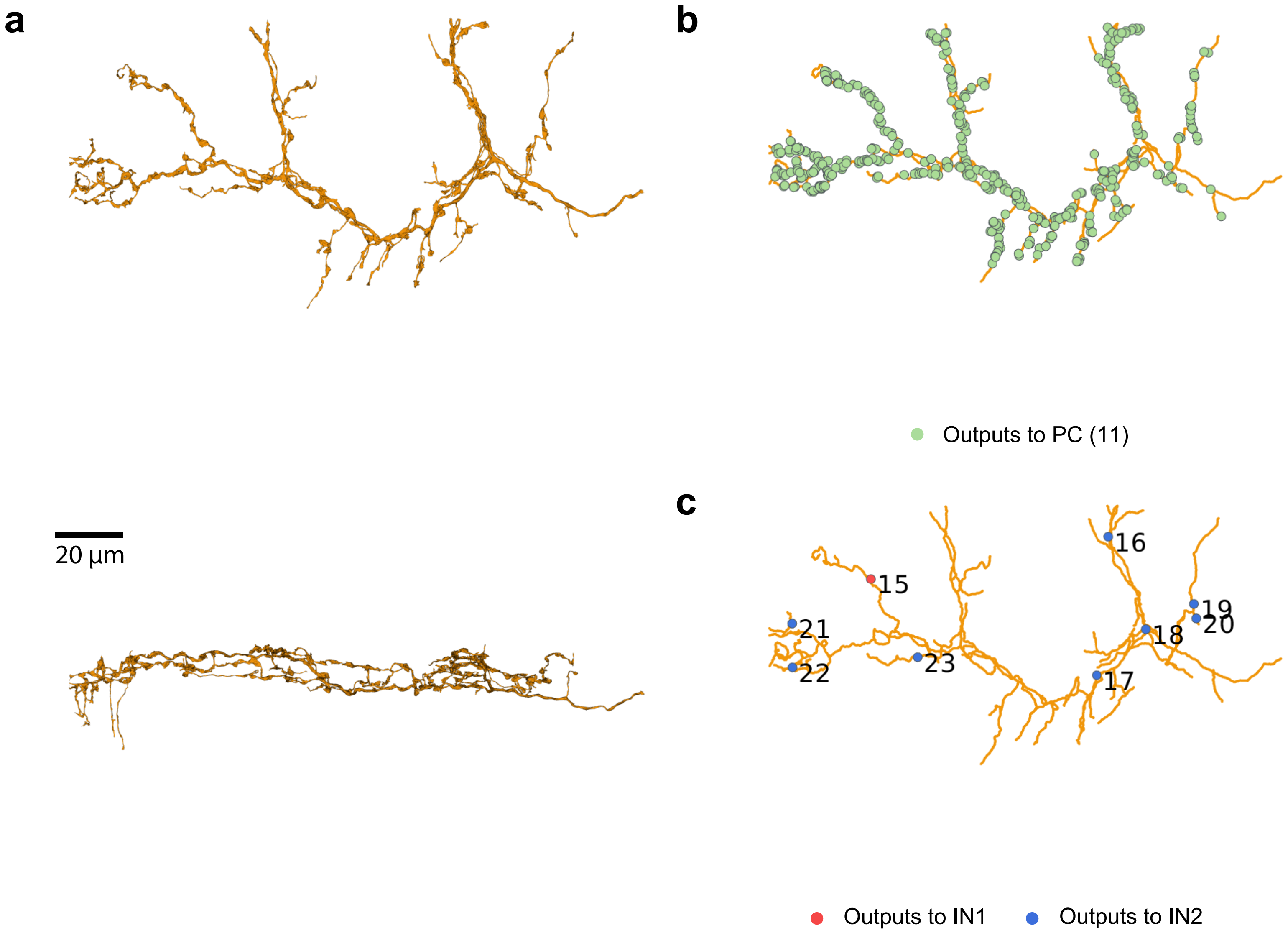

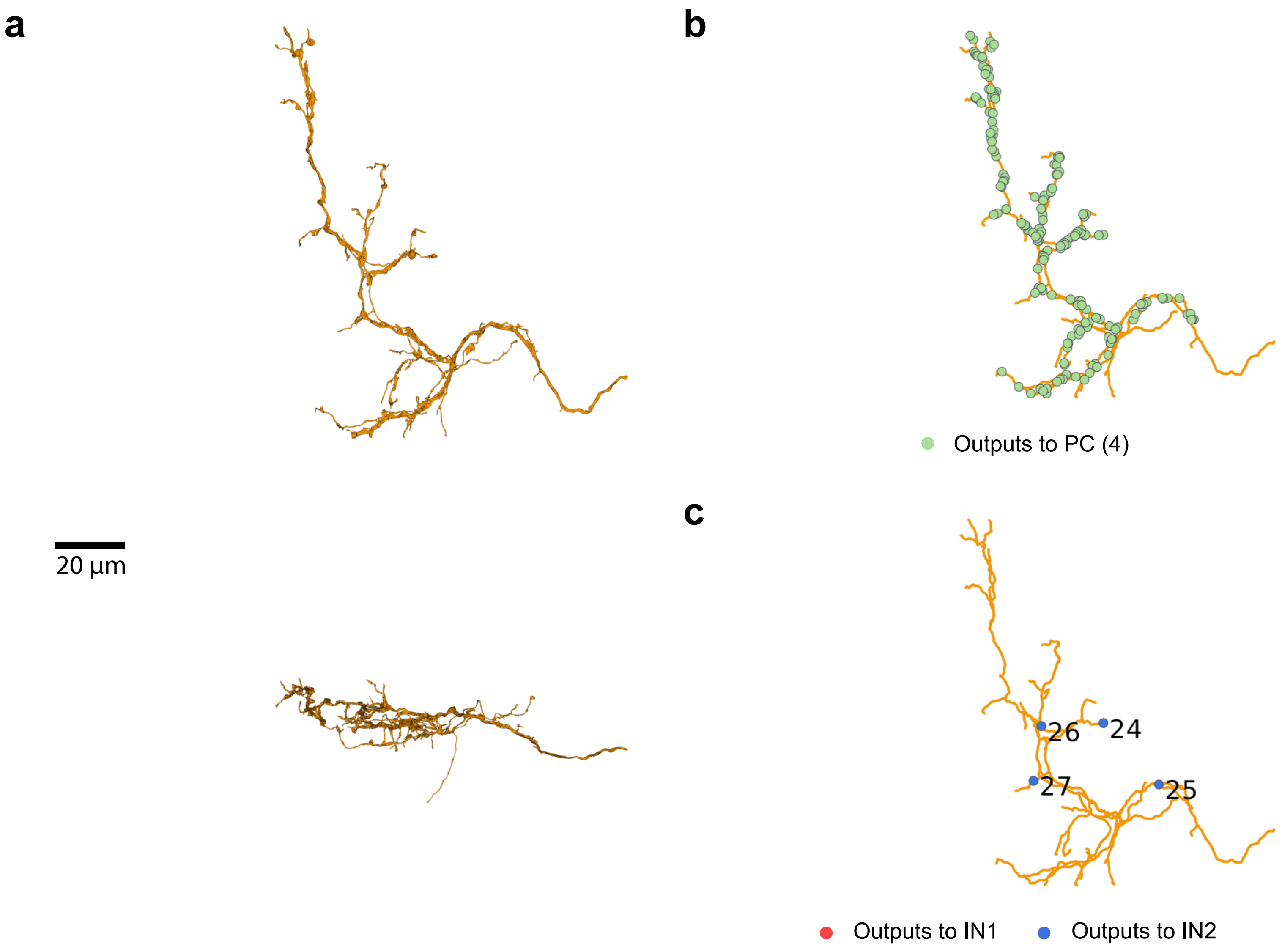

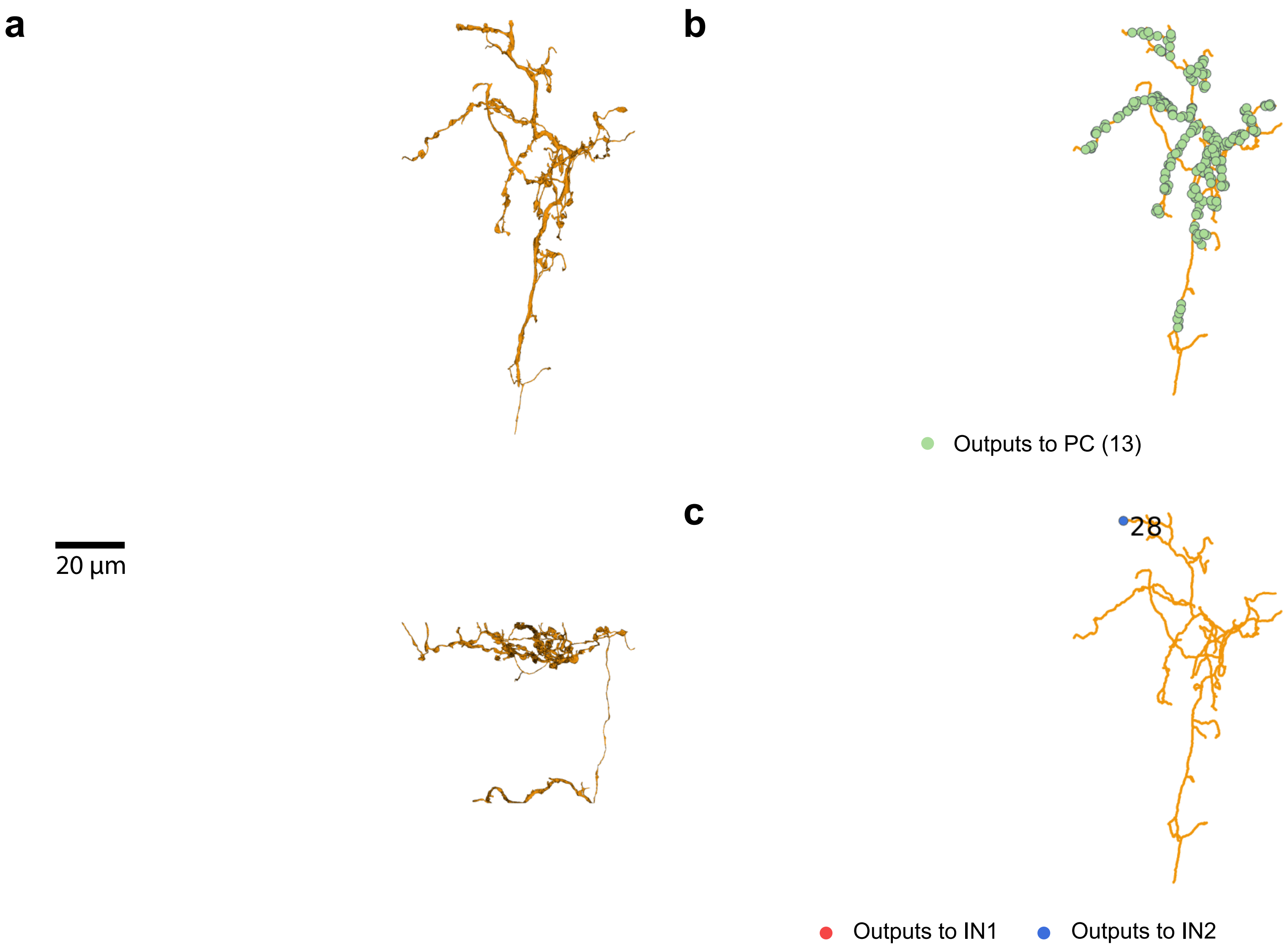


CF (681)

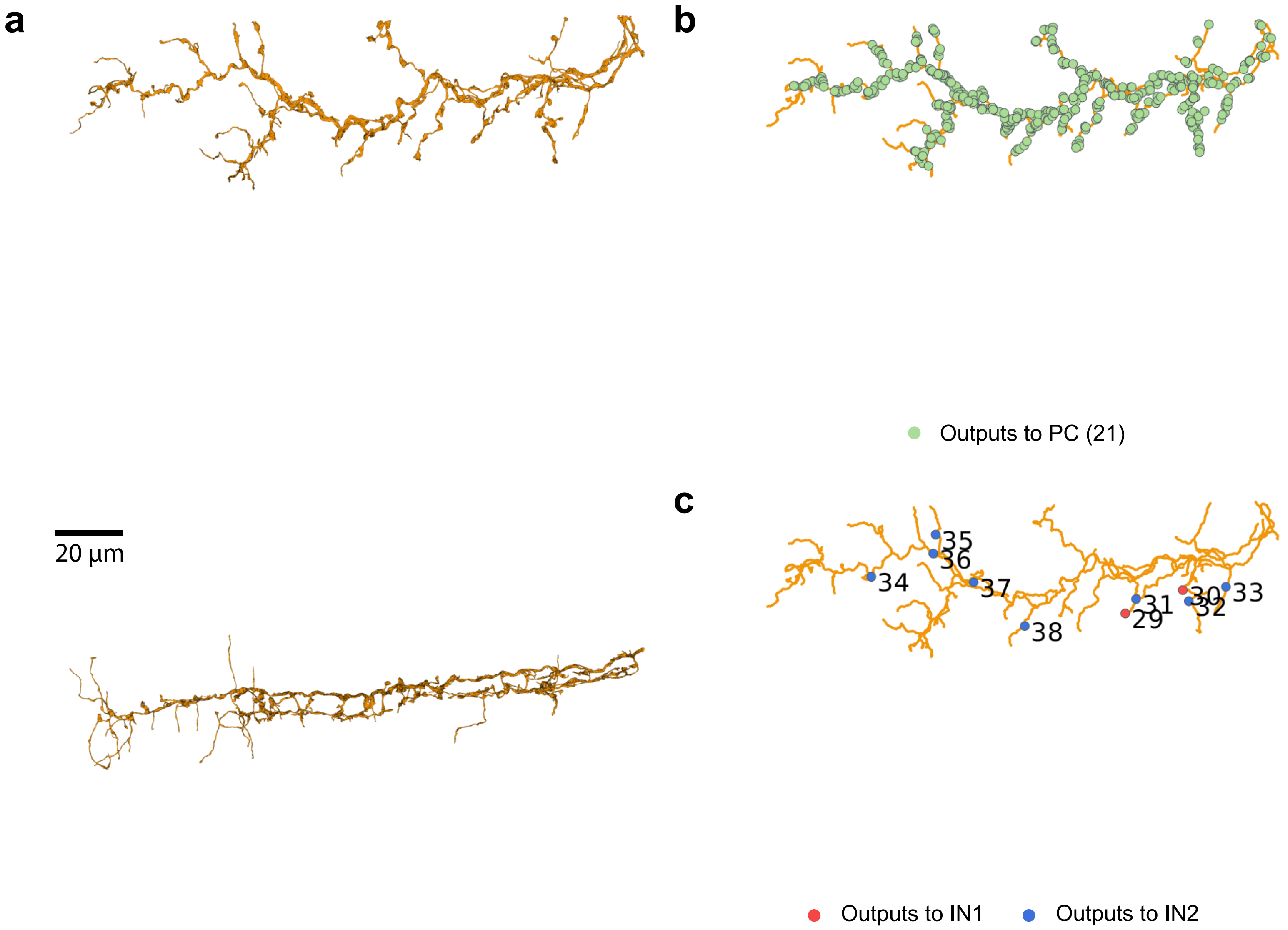

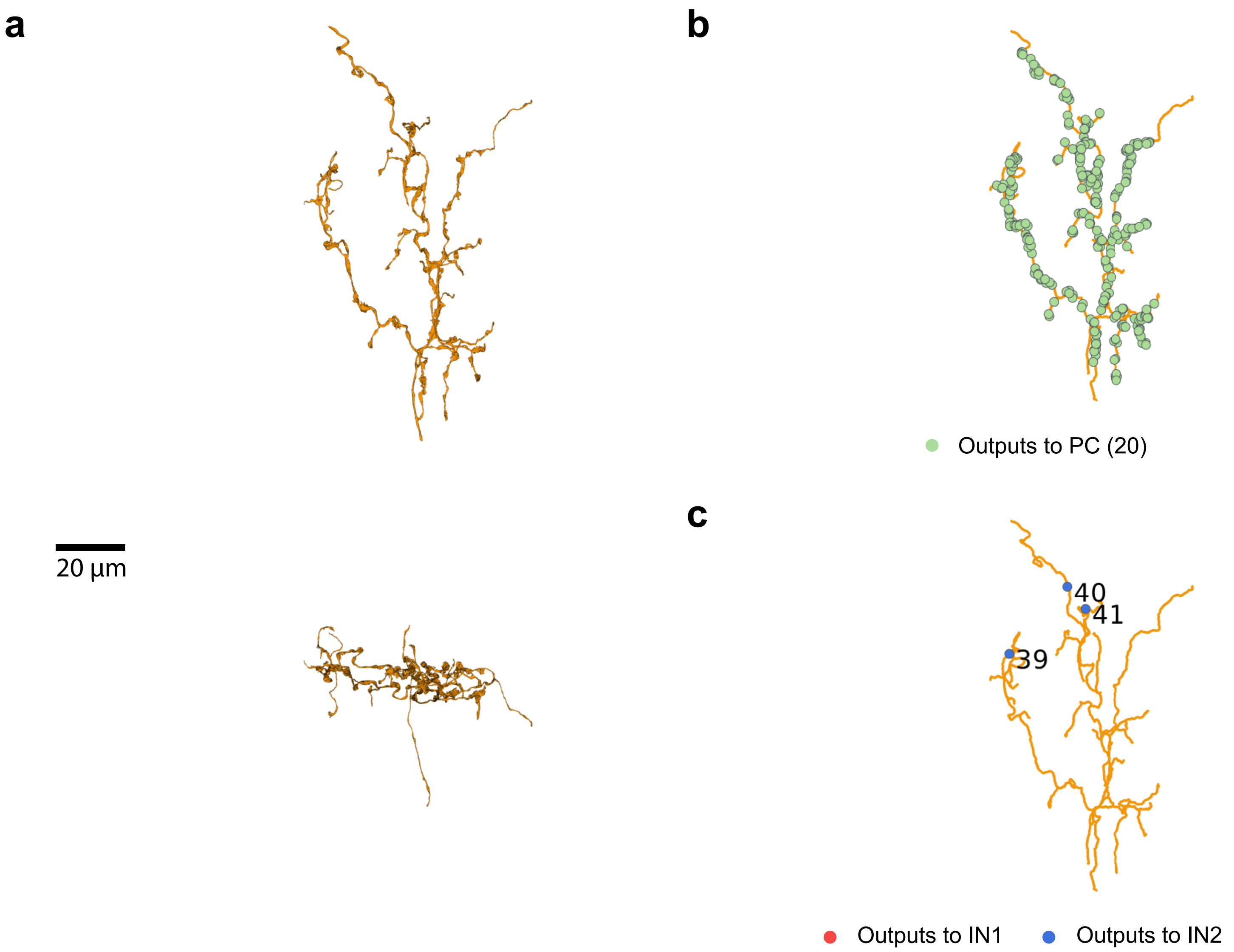

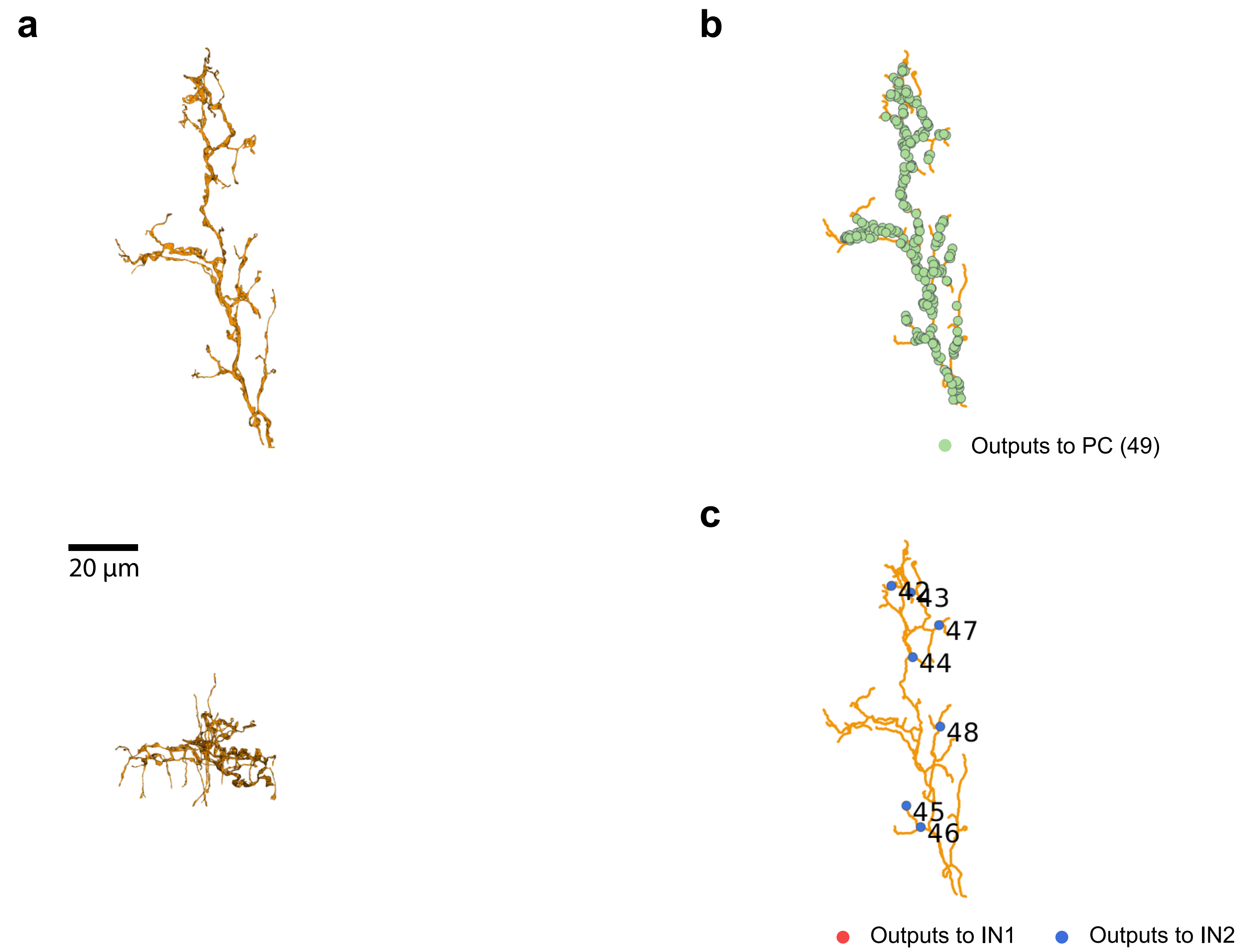

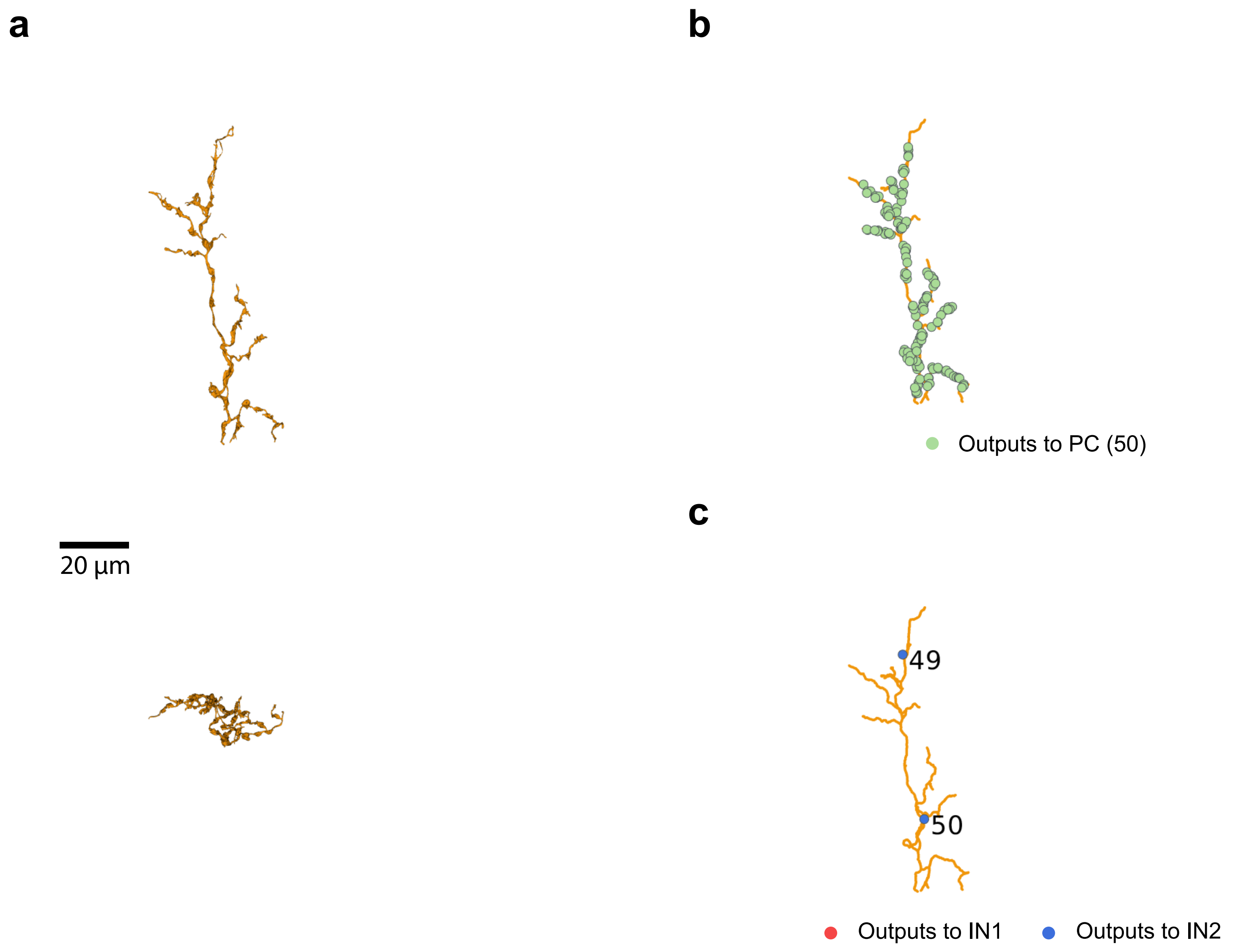


CF (931)

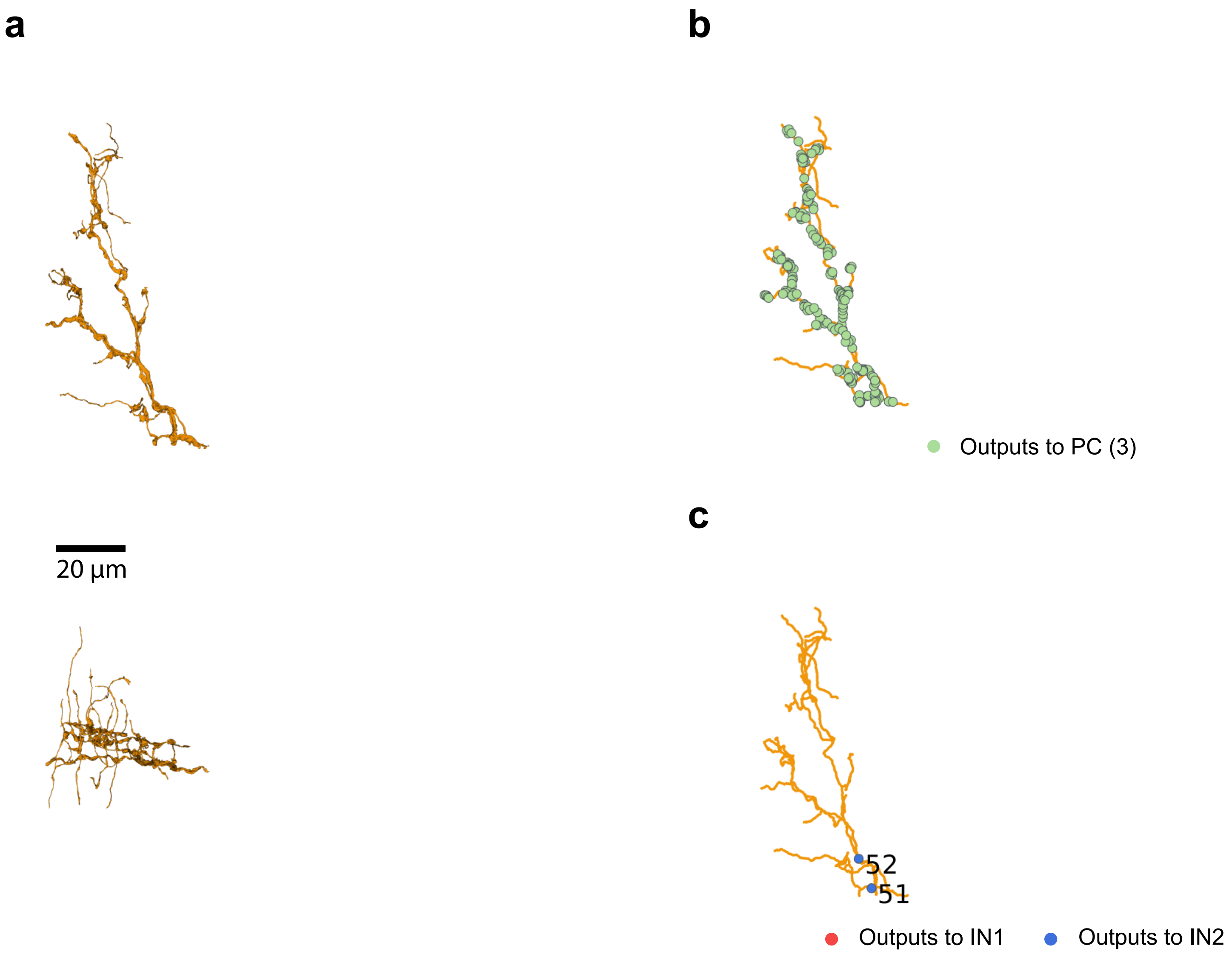


CF (166)

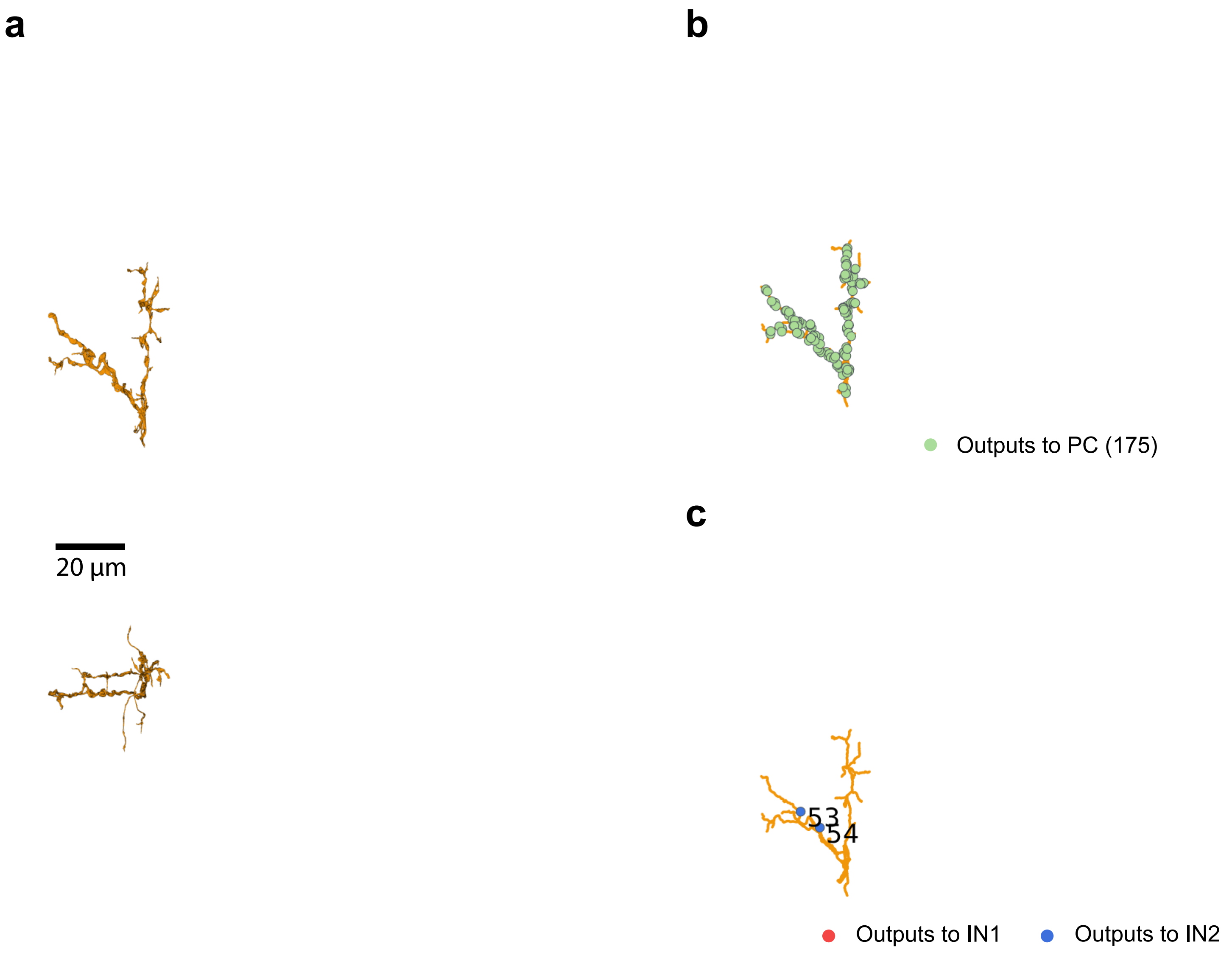


CF (953)
