## Supplementary Data 3 for "A cerebellar disinhibitory circuit supports synaptic plasticity"

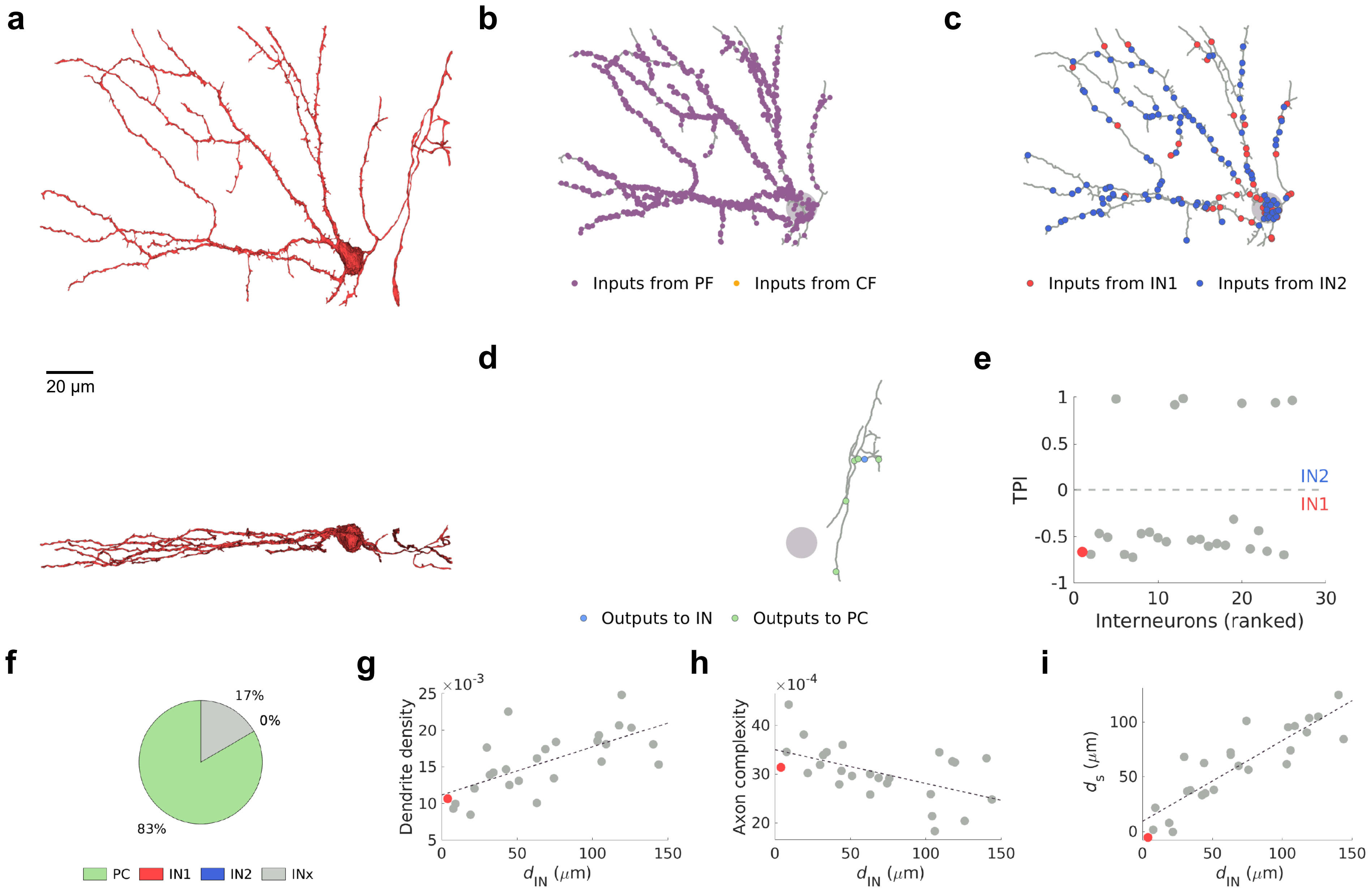
a
20 μm
b
Inputs from PF
Inputs from CF
c
Inputs from IN1
Inputs from IN2
d
Outputs to IN
Outputs to PC
e
TPI
IN2
IN1
Interneurons (ranked)
f
17%
0%
83%
PC
IN1
IN2
INx
g
Dendrite density
$\times 10^{-3}$
$d_{IN}$ ($\mu$m)
h
Axon complexity
$\times 10^{-4}$
$d_{IN}$ ($\mu$m)
i
$d_s$ ($\mu$m)
$d_{IN}$ ($\mu$m)

IN1 (28)

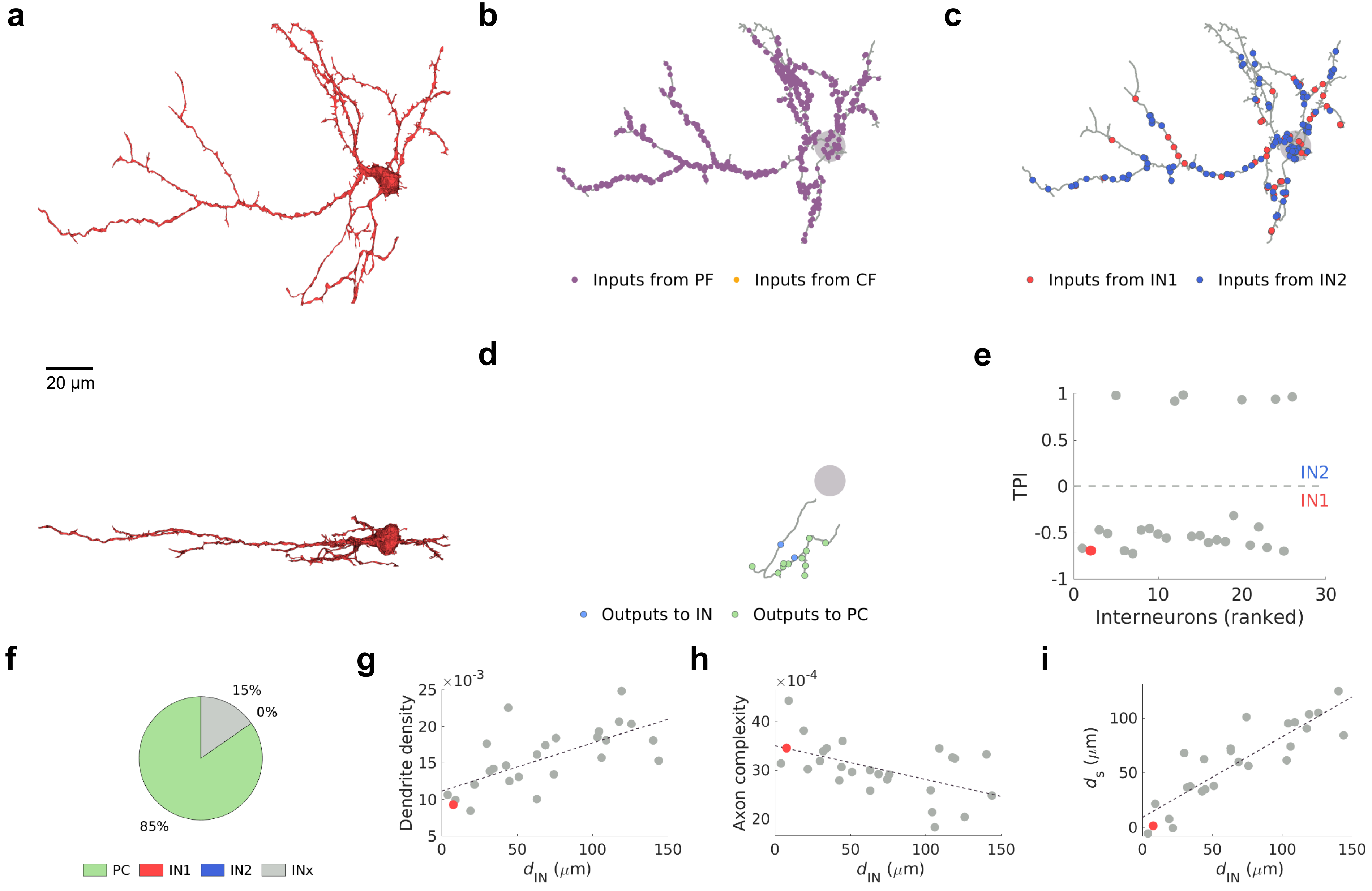
a
20 µm
b
Inputs from PF
Inputs from CF
c
Inputs from IN1
Inputs from IN2
d
Outputs to IN
Outputs to PC
e
TPI
IN2
IN1
Interneurons (ranked)
f
15%
0%
85%
PC
IN1
IN2
INx
g
Dendrite density
×10^-3
h
Axon complexity
×10^-4
i
d_s (µm)
d_IN (µm)

IN1 (24)

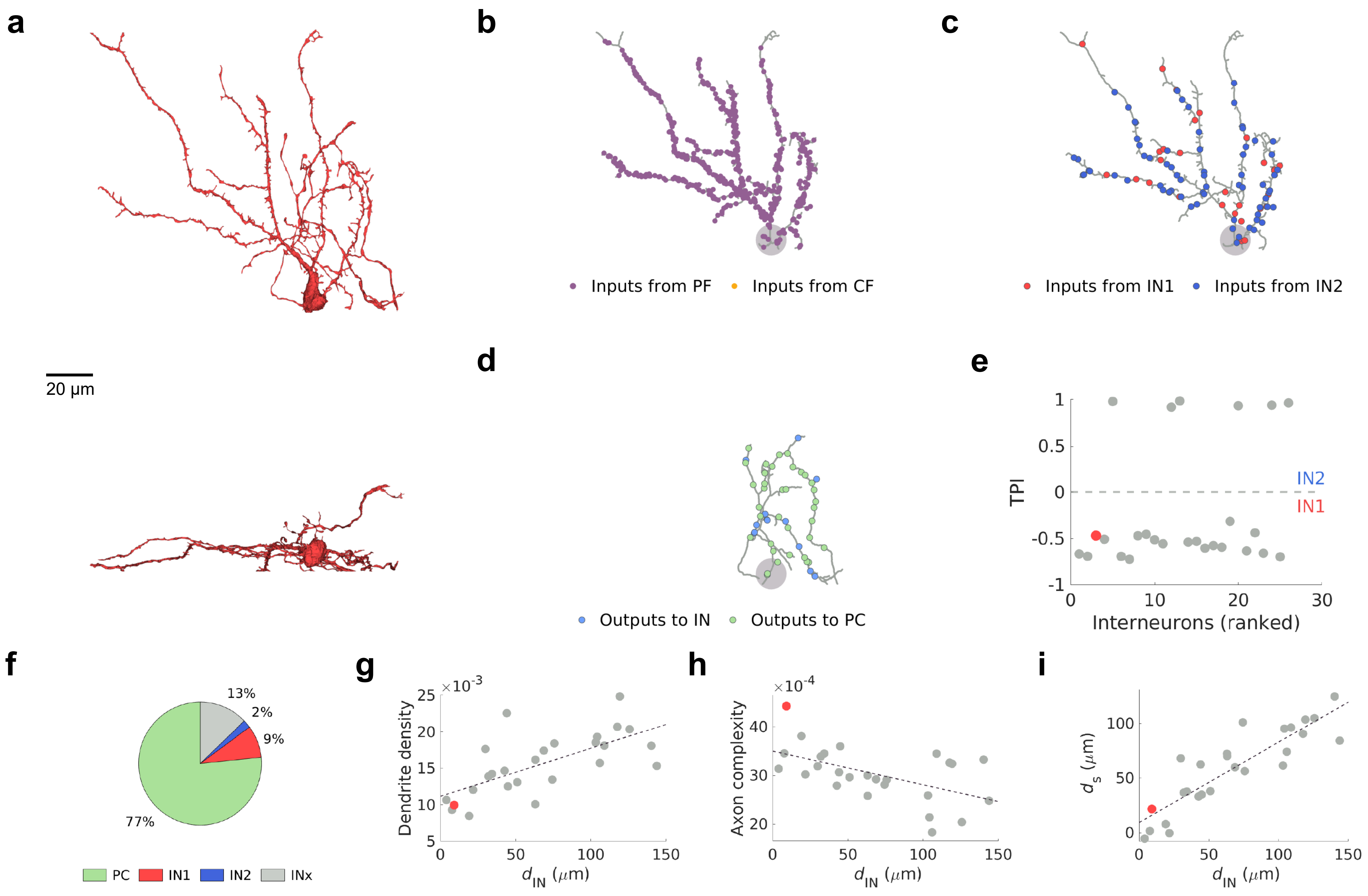
a
20 μm
b
Inputs from PF
Inputs from CF
c
Inputs from IN1
Inputs from IN2
d
Outputs to IN
Outputs to PC
e
TPI
IN2
IN1
Interneurons (ranked)
f
13%
2%
9%
77%
PC
IN1
IN2
INx
g
Dendrite density
×10-3
d_IN (μm)
h
Axon complexity
×10-4
d_IN (μm)
i
d_s (μm)
d_IN (μm)

IN1 (71)

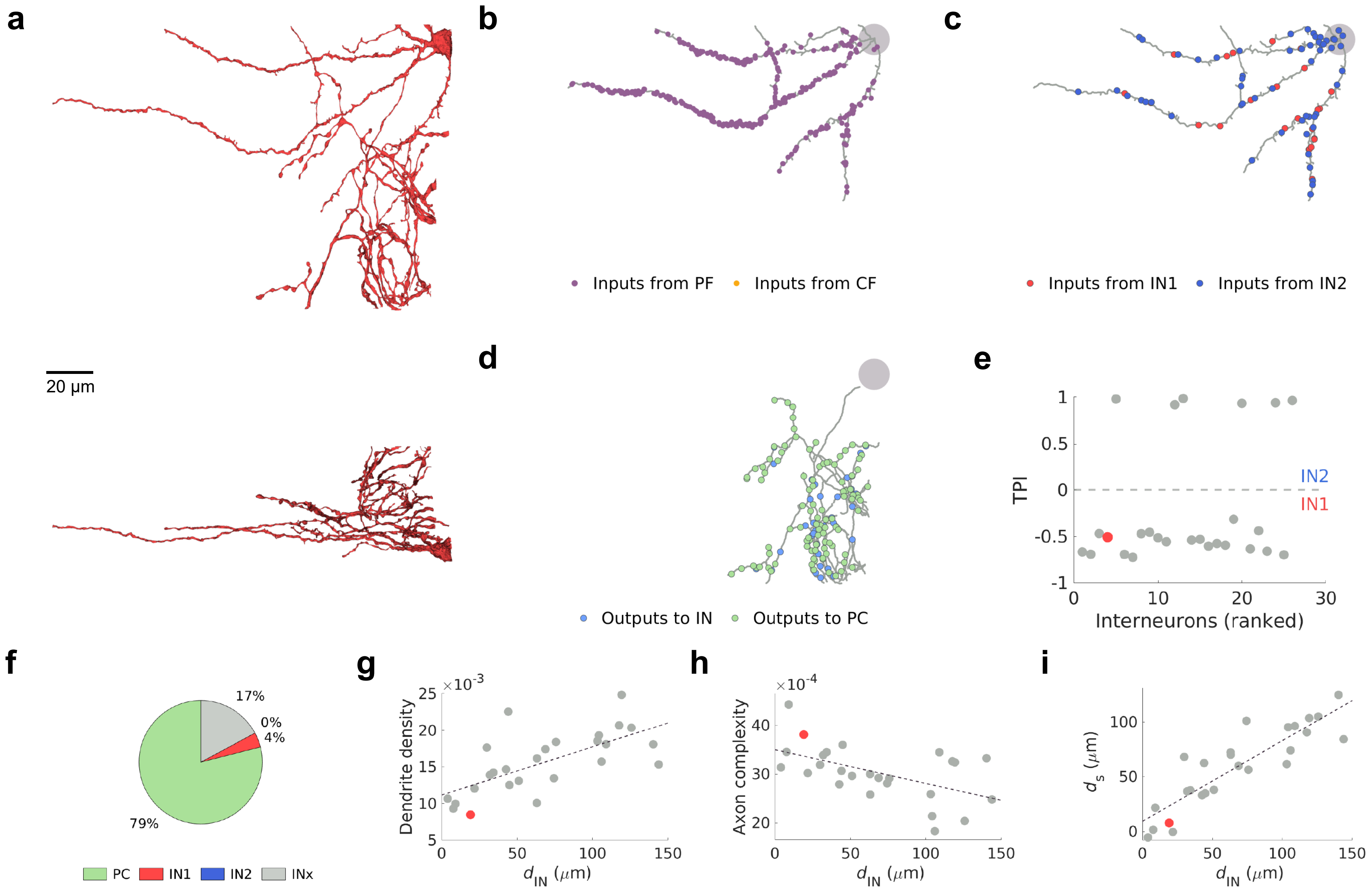
a
20 μm
b
Inputs from PF
Inputs from CF
c
Inputs from IN1
Inputs from IN2
d
Outputs to IN
Outputs to PC
e
TPI
IN2
IN1
Interneurons (ranked)
f
17%
0%
4%
79%
PC
IN1
IN2
INx
g
Dendrite density
×10^-3
d_IN (μm)
h
Axon complexity
×10^-4
d_IN (μm)
i
d_s (μm)
d_IN (μm)

IN1 (9)

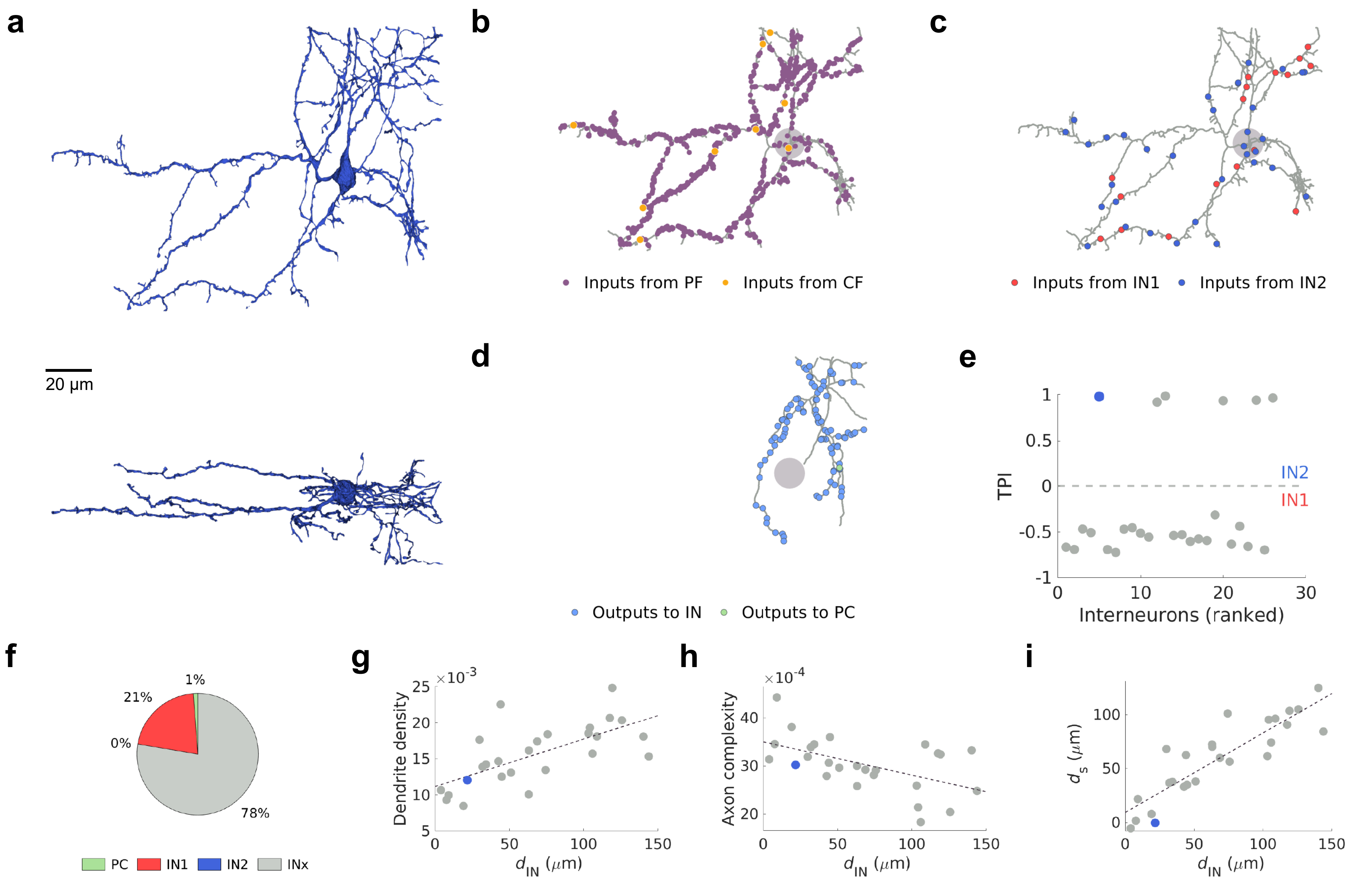
a
20 μm
b
Inputs from PF
Inputs from CF
c
Inputs from IN1
Inputs from IN2
d
Outputs to IN
Outputs to PC
e
TPI
IN2
IN1
Interneurons (ranked)
f
1%
21%
0%
78%
PC
IN1
IN2
INx
g
Dendrite density
×10-3
$d_{IN}$ (μm)
h
Axon complexity
×10-4
$d_{IN}$ (μm)
i
$d_s$ (μm)
$d_{IN}$ (μm)

IN2 (30)

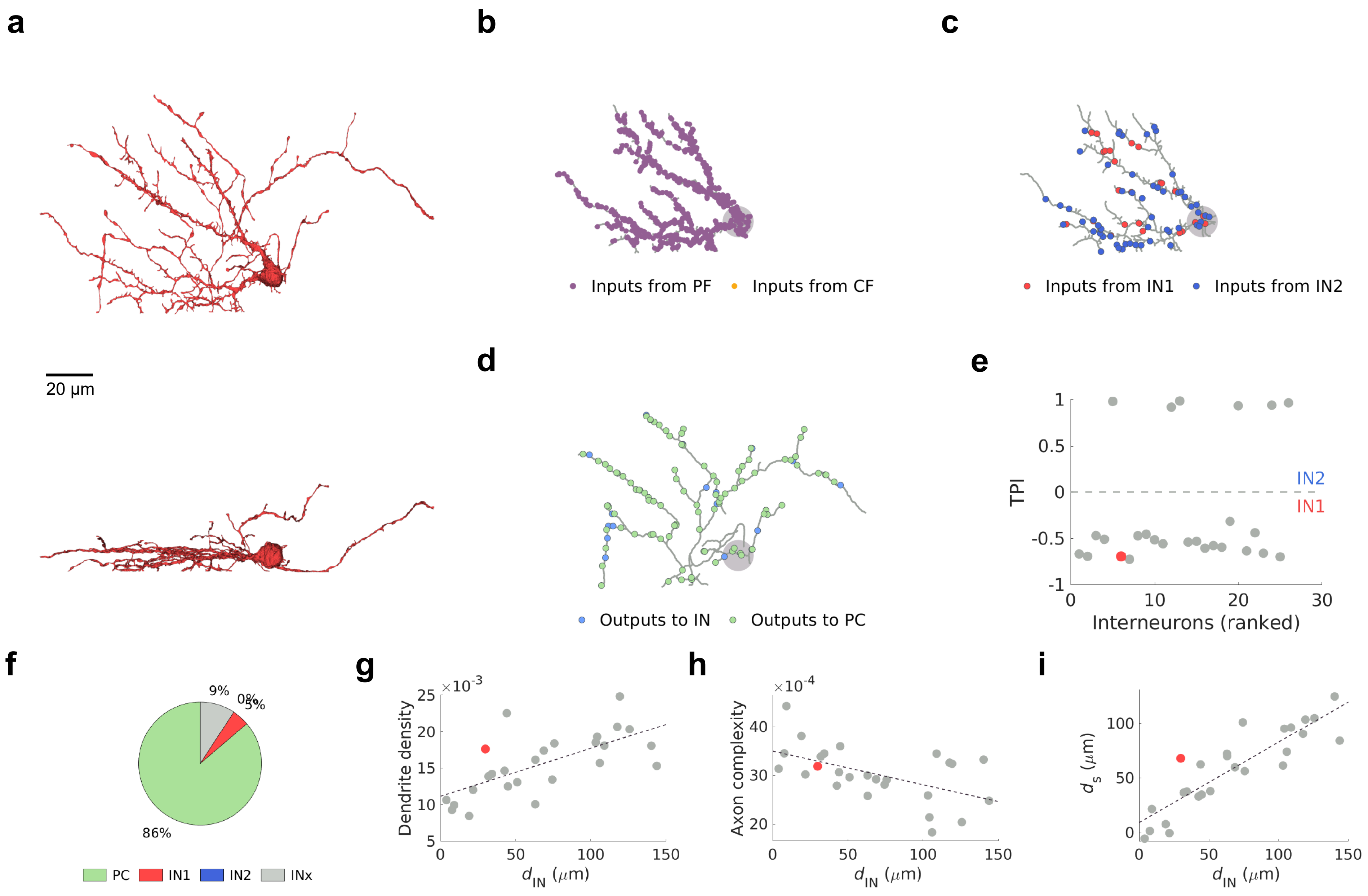
a
20 μm
b
Inputs from PF
Inputs from CF
c
Inputs from IN1
Inputs from IN2
d
Outputs to IN
Outputs to PC
e
TPI
IN2
IN1
Interneurons (ranked)
f
9%
0%
5%
86%
PC
IN1
IN2
INx
g
Dendrite density
×10^-3
$d_{IN}$ (μm)
h
Axon complexity
×10^-4
$d_{IN}$ (μm)
i
$d_s$ (μm)
$d_{IN}$ (μm)

IN1 (68)

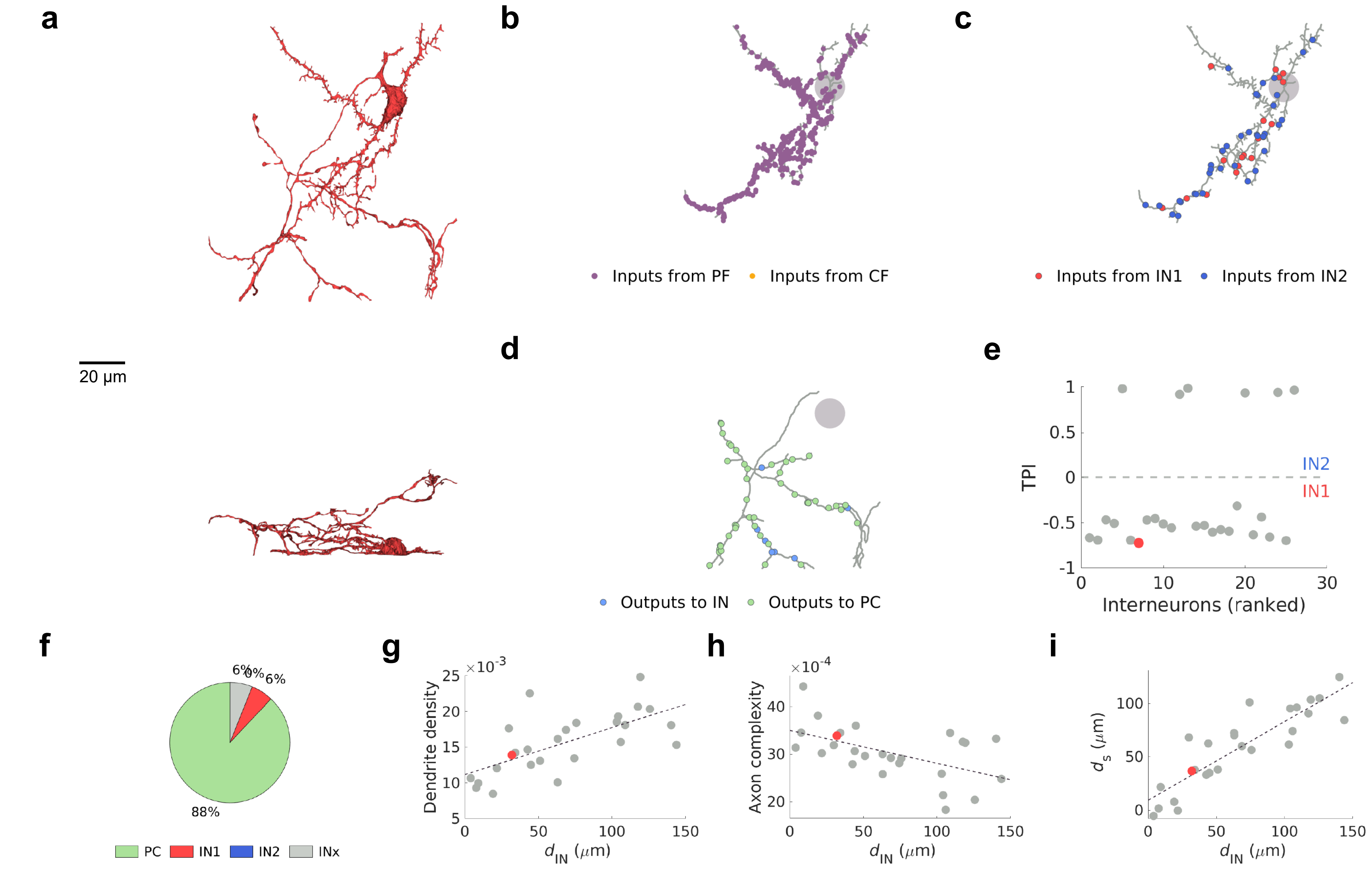
a
b
c
d
e
f
g
h
i
20 μm
Inputs from PF
Inputs from CF
Inputs from IN1
Inputs from IN2
Outputs to IN
Outputs to PC
TPI
IN2
IN1
Interneurons (ranked)
6%
0%
6%
88%
PC
IN1
IN2
INx
Dendrite density
×10^-3
Axon complexity
×10^-4
d_IN (μm)
d_s (μm)

IN1 (74)

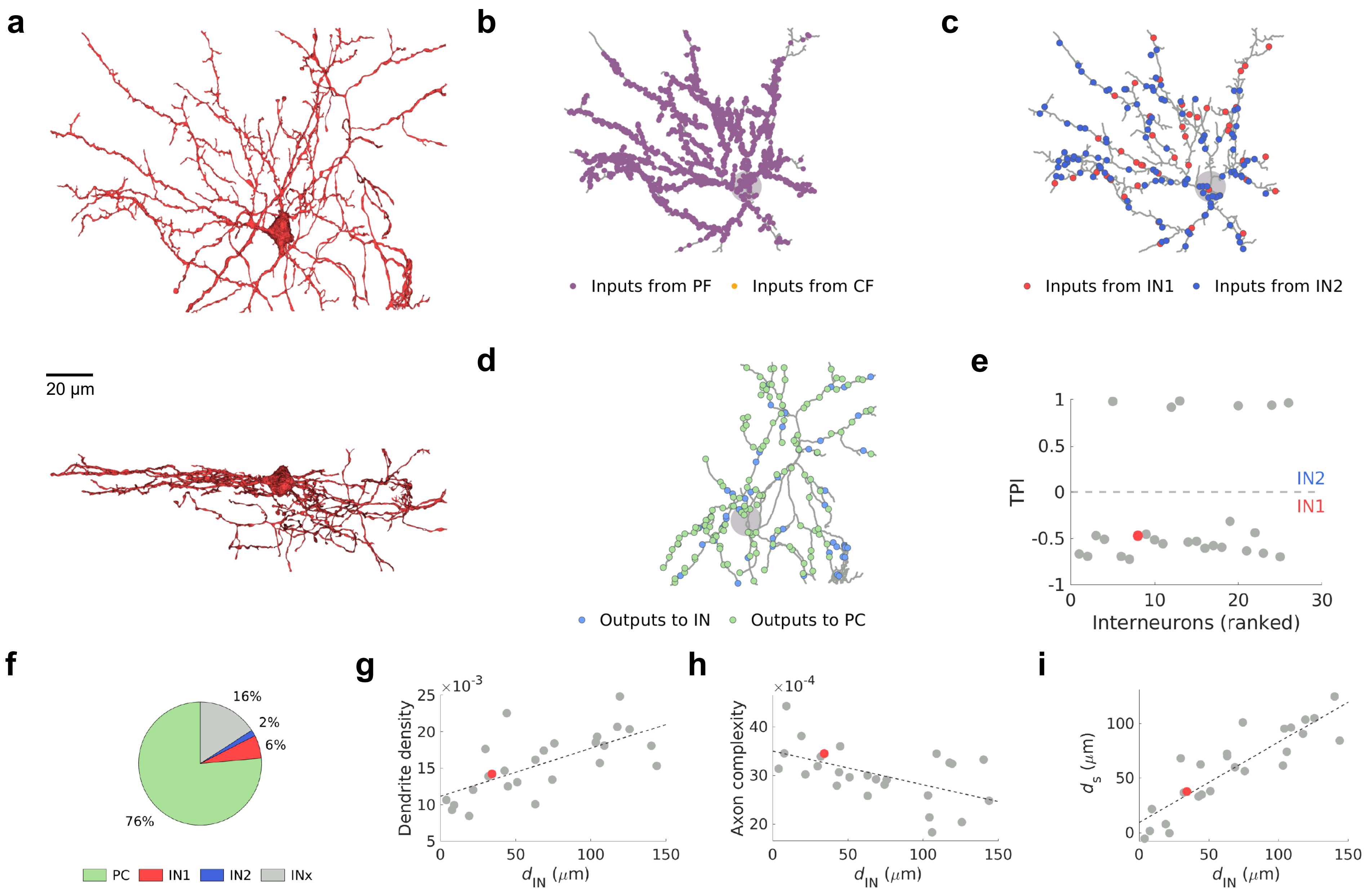
a
20 μm
b
Inputs from PF
Inputs from CF
c
Inputs from IN1
Inputs from IN2
d
Outputs to IN
Outputs to PC
e
TPI
IN2
IN1
Interneurons (ranked)
f
16%
2%
6%
76%
PC
IN1
IN2
INx
g
Dendrite density
$\times 10^{-3}$
$d_{IN}$ ($\mu$m)
h
Axon complexity
$\times 10^{-4}$
$d_{IN}$ ($\mu$m)
i
$d_s$ ($\mu$m)
$d_{IN}$ ($\mu$m)

IN1 (15)

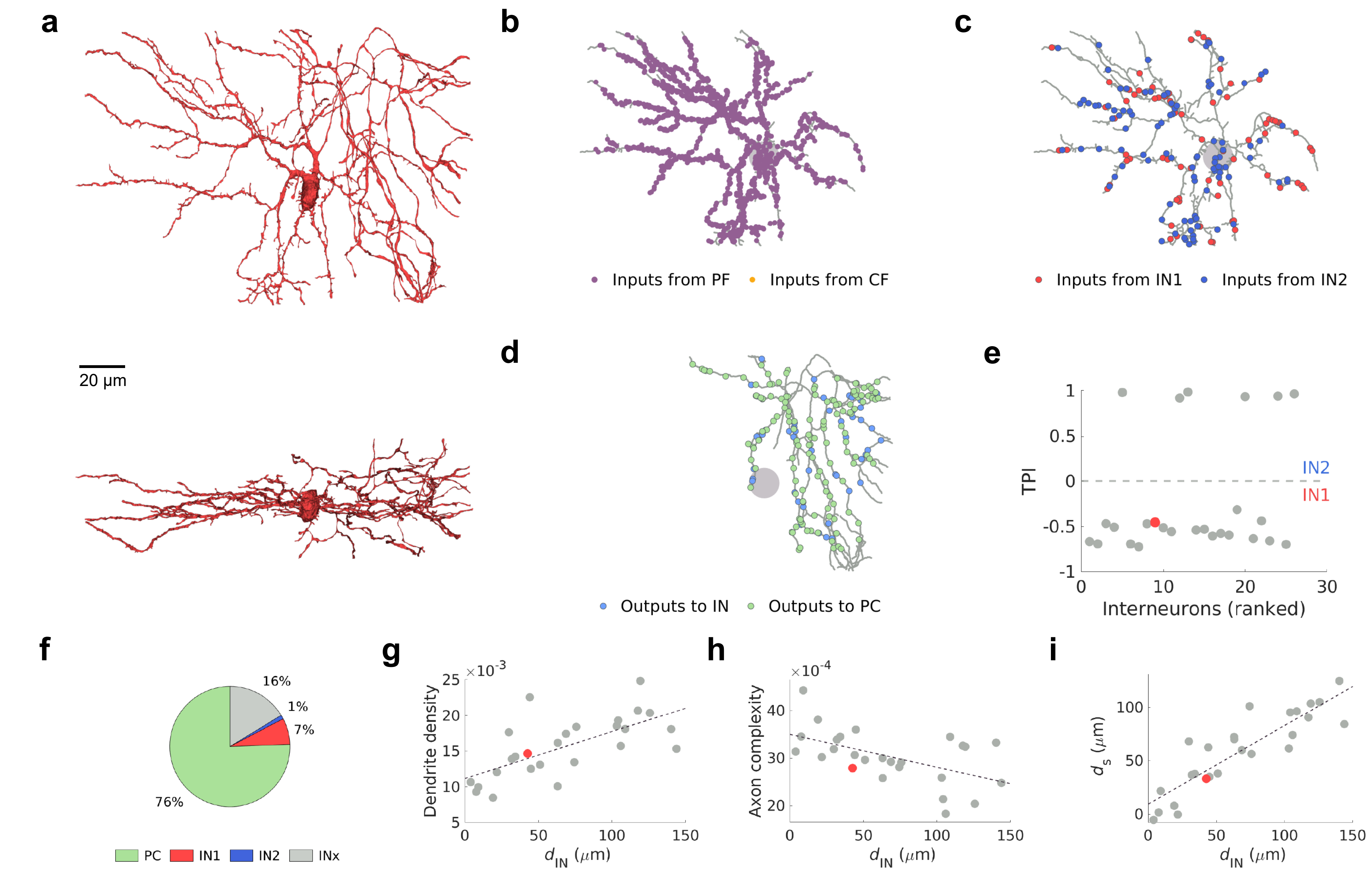
a
20 μm
b
Inputs from PF
Inputs from CF
c
Inputs from IN1
Inputs from IN2
d
Outputs to IN
Outputs to PC
e
TPI
IN2
IN1
1
0.5
0
-0.5
-1
0
10
20
30
Interneurons (ranked)
f
16%
1%
7%
76%
PC
IN1
IN2
INx
g
×10-3
Dendrite density
$d_{IN}$ (μm)
h
×10-4
Axon complexity
$d_{IN}$ (μm)
i
$d_s$ (μm)
$d_{IN}$ (μm)

IN1 (17)

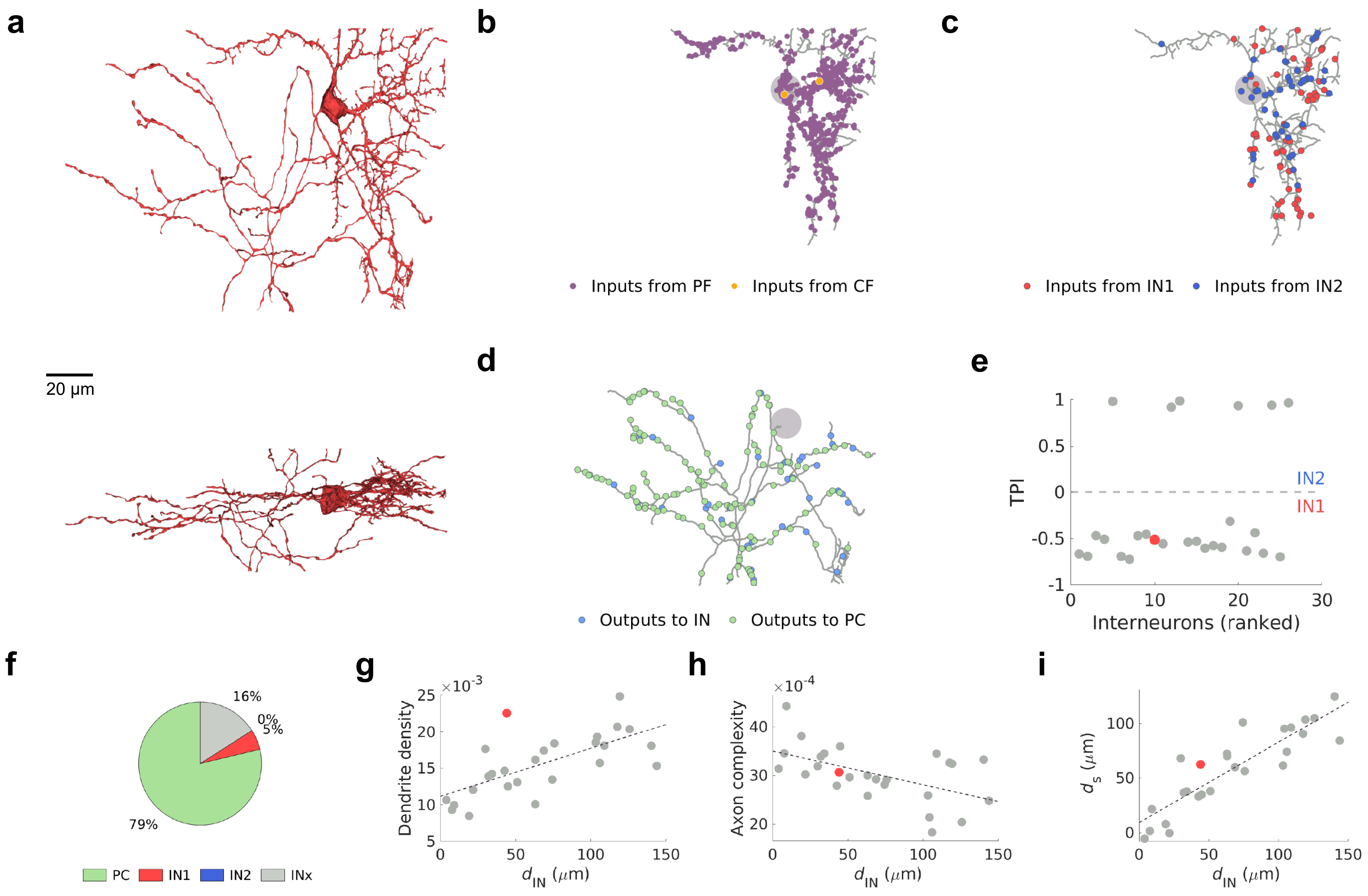
a
20 μm
b
Inputs from PF
Inputs from CF
c
Inputs from IN1
Inputs from IN2
d
Outputs to IN
Outputs to PC
e
TPI
IN2
IN1
Interneurons (ranked)
f
16%
0%
5%
79%
PC
IN1
IN2
INx
g
Dendrite density
$\times 10^{-3}$
$d_{IN}$ ($\mu$m)
h
Axon complexity
$\times 10^{-4}$
$d_{IN}$ ($\mu$m)
i
$d_s$ ($\mu$m)
$d_{IN}$ ($\mu$m)

IN1 (10)

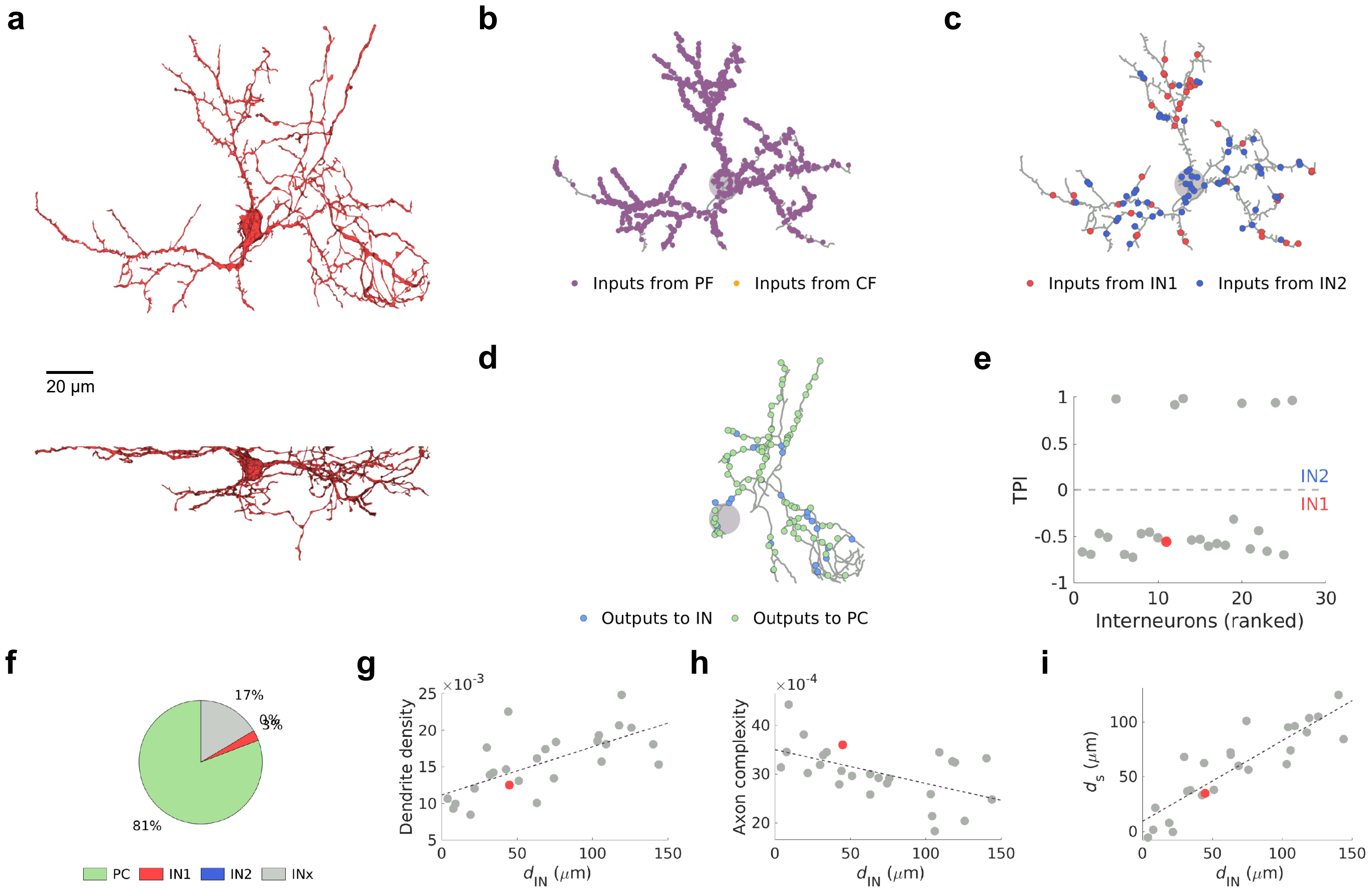
a
20 μm
b
Inputs from PF
Inputs from CF
c
Inputs from IN1
Inputs from IN2
d
Outputs to IN
Outputs to PC
e
TPI
IN2
IN1
Interneurons (ranked)
f
17%
0%
3%
81%
PC
IN1
IN2
INx
g
Dendrite density
$\times 10^{-3}$
$d_{IN}$ ($\mu$m)
h
Axon complexity
$\times 10^{-4}$
$d_{IN}$ ($\mu$m)
i
$d_s$ ($\mu$m)
$d_{IN}$ ($\mu$m)

IN1 (26)

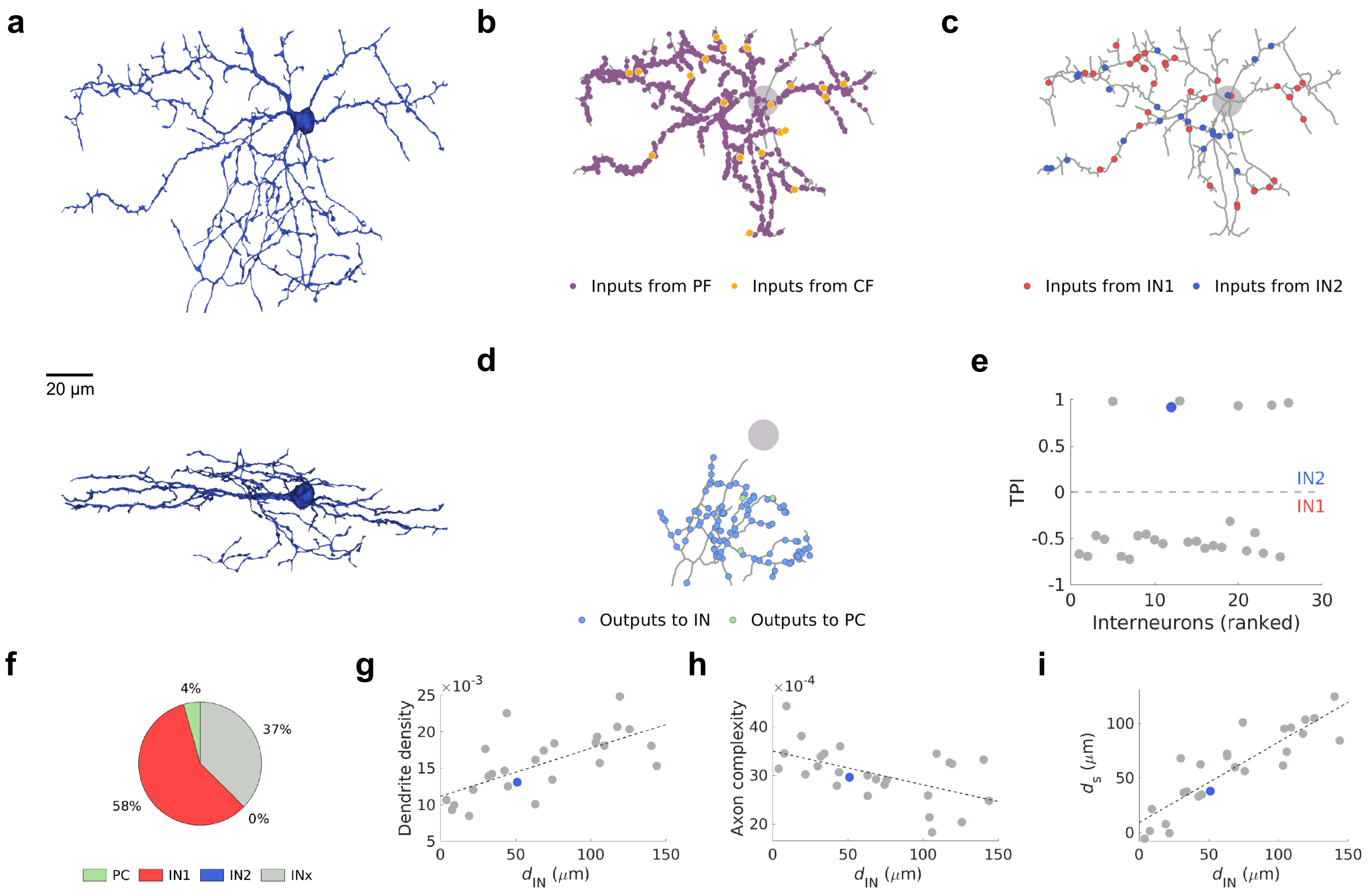

a
20 μm
b
Inputs from PF
Inputs from CF
c
Inputs from IN1
Inputs from IN2
d
Outputs to IN
Outputs to PC
e
TPI
IN2
IN1
Interneurons (ranked)
f
4%
37%
58%
0%
PC
IN1
IN2
INx
g
Dendrite density
×10^-3
$d_{IN}$ ($\mu$m)
h
Axon complexity
×10^-4
$d_{IN}$ ($\mu$m)
i
$d_s$ ($\mu$m)
$d_{IN}$ ($\mu$m)


IN2 (31)

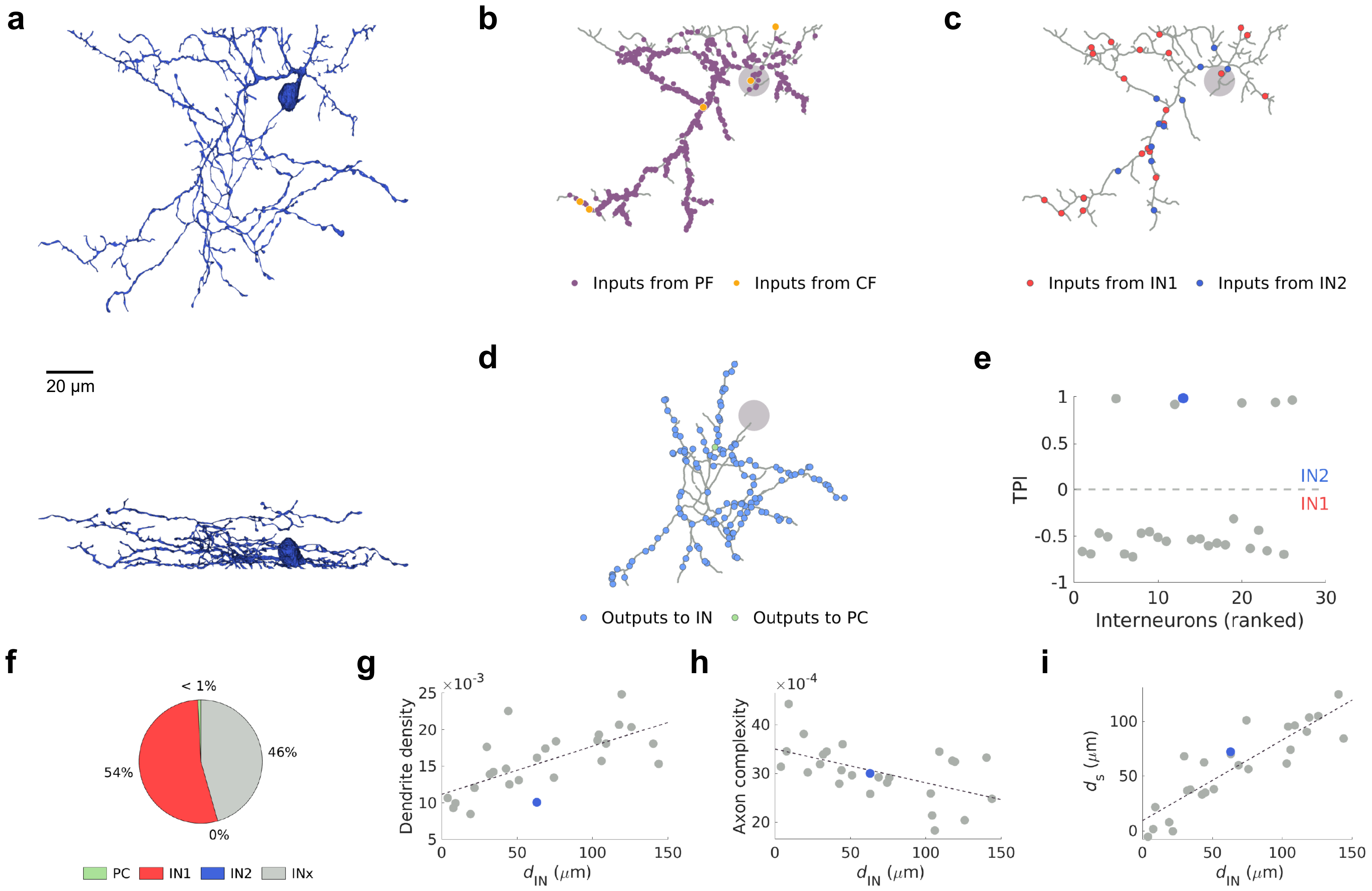
a
20 μm
b
Inputs from PF
Inputs from CF
c
Inputs from IN1
Inputs from IN2
d
Outputs to IN
Outputs to PC
e
TPI
IN2
IN1
Interneurons (ranked)
f
< 1%
46%
54%
0%
PC
IN1
IN2
INx
g
Dendrite density
×10-3
d_IN (μm)
h
Axon complexity
×10-4
d_IN (μm)
i
d_s (μm)
d_IN (μm)

IN2 (27)

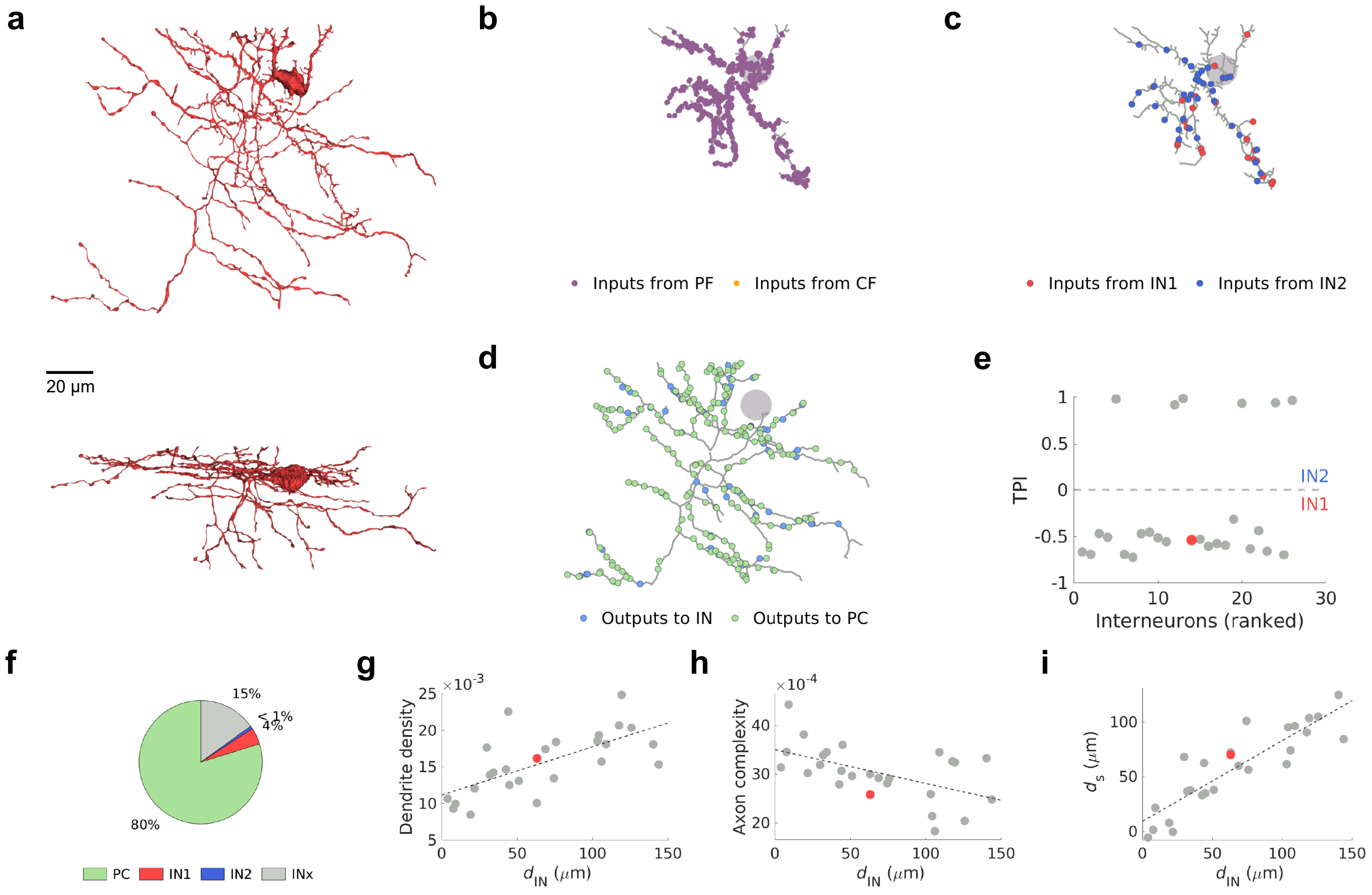
a
20 μm
b
Inputs from PF
Inputs from CF
c
Inputs from IN1
Inputs from IN2
d
Outputs to IN
Outputs to PC
e
TPI
IN2
IN1
Interneurons (ranked)
f
15%
< 1%
4%
80%
PC
IN1
IN2
INx
g
Dendrite density
×10-3
d_IN (μm)
h
Axon complexity
×10-4
d_IN (μm)
i
d_s (μm)
d_IN (μm)

IN1 (39)

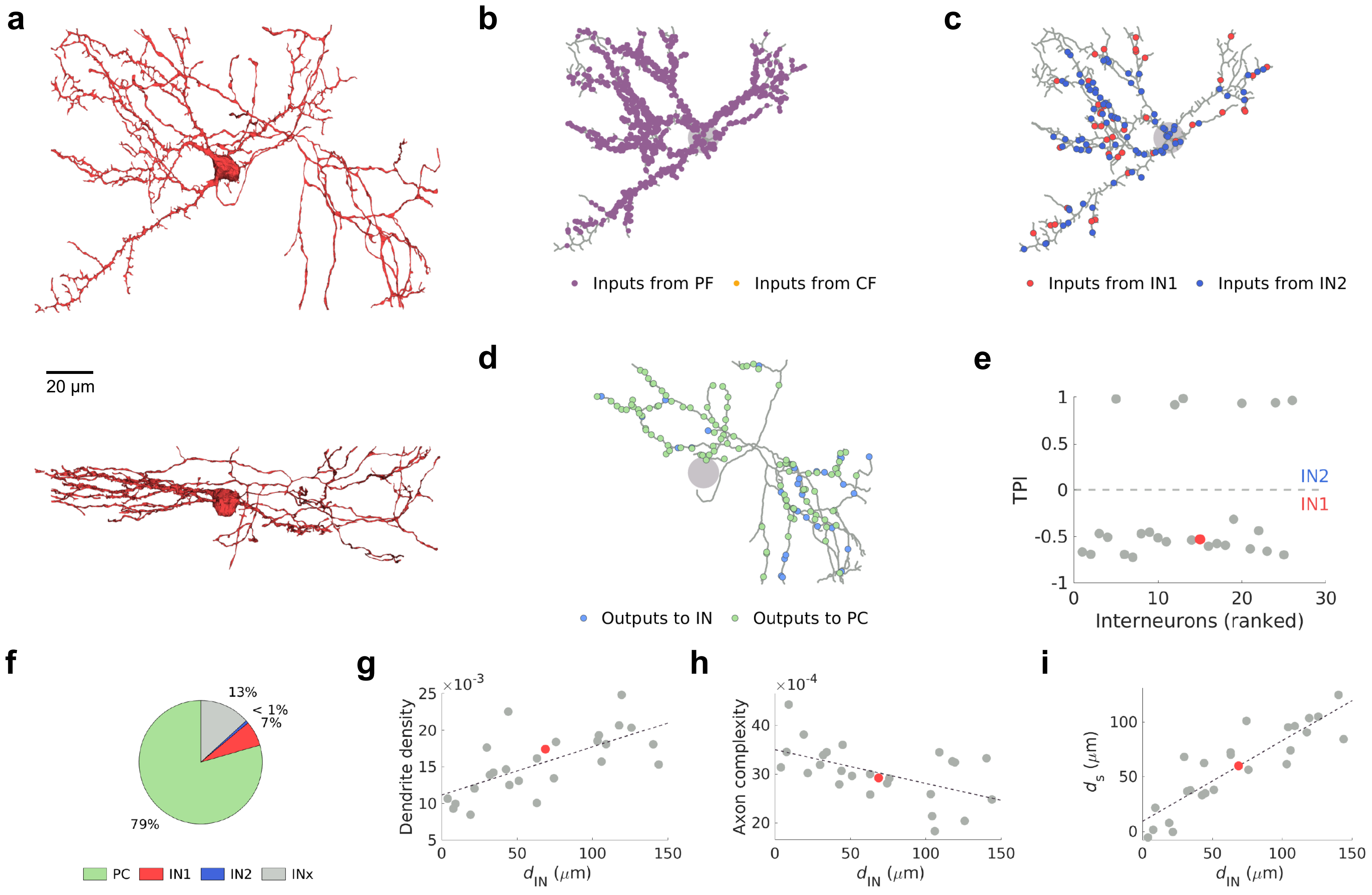
a
20 μm
b
Inputs from PF
Inputs from CF
c
Inputs from IN1
Inputs from IN2
d
Outputs to IN
Outputs to PC
e
TPI
IN2
IN1
Interneurons (ranked)
f
13%
< 1%
7%
79%
PC
IN1
IN2
INx
g
Dendrite density
×10^-3
h
Axon complexity
×10^-4
i
d_s (μm)
d_IN (μm)

IN1 (12)

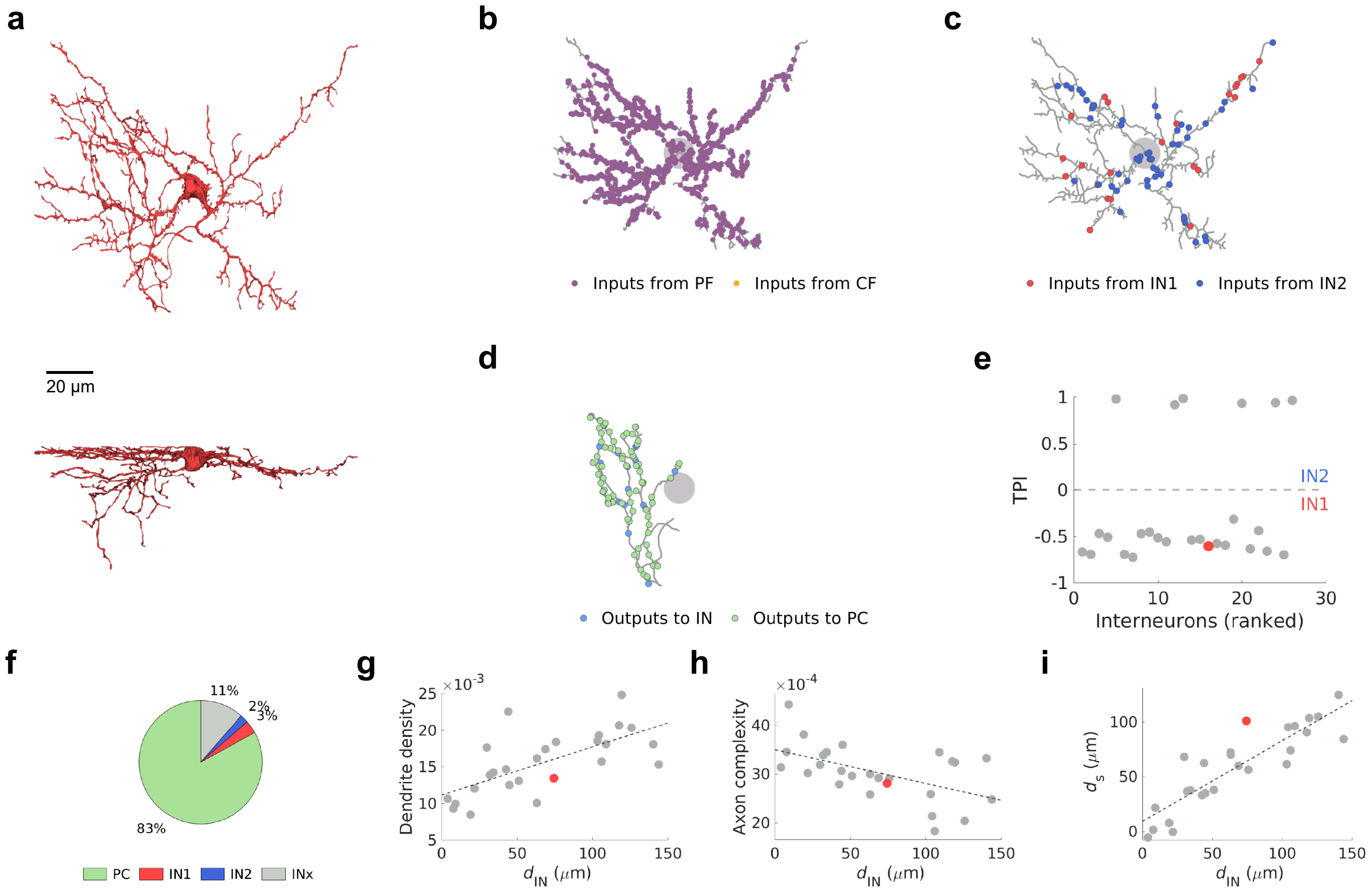
a
20 µm
b
Inputs from PF
Inputs from CF
c
Inputs from IN1
Inputs from IN2
d
Outputs to IN
Outputs to PC
e
TPI
IN2
IN1
Interneurons (ranked)
f
11%
2%
3%
83%
PC
IN1
IN2
INx
g
Dendrite density
×10^-3
d_IN (µm)
h
Axon complexity
×10^-4
d_IN (µm)
i
d_s (µm)
d_IN (µm)

IN1 (54)

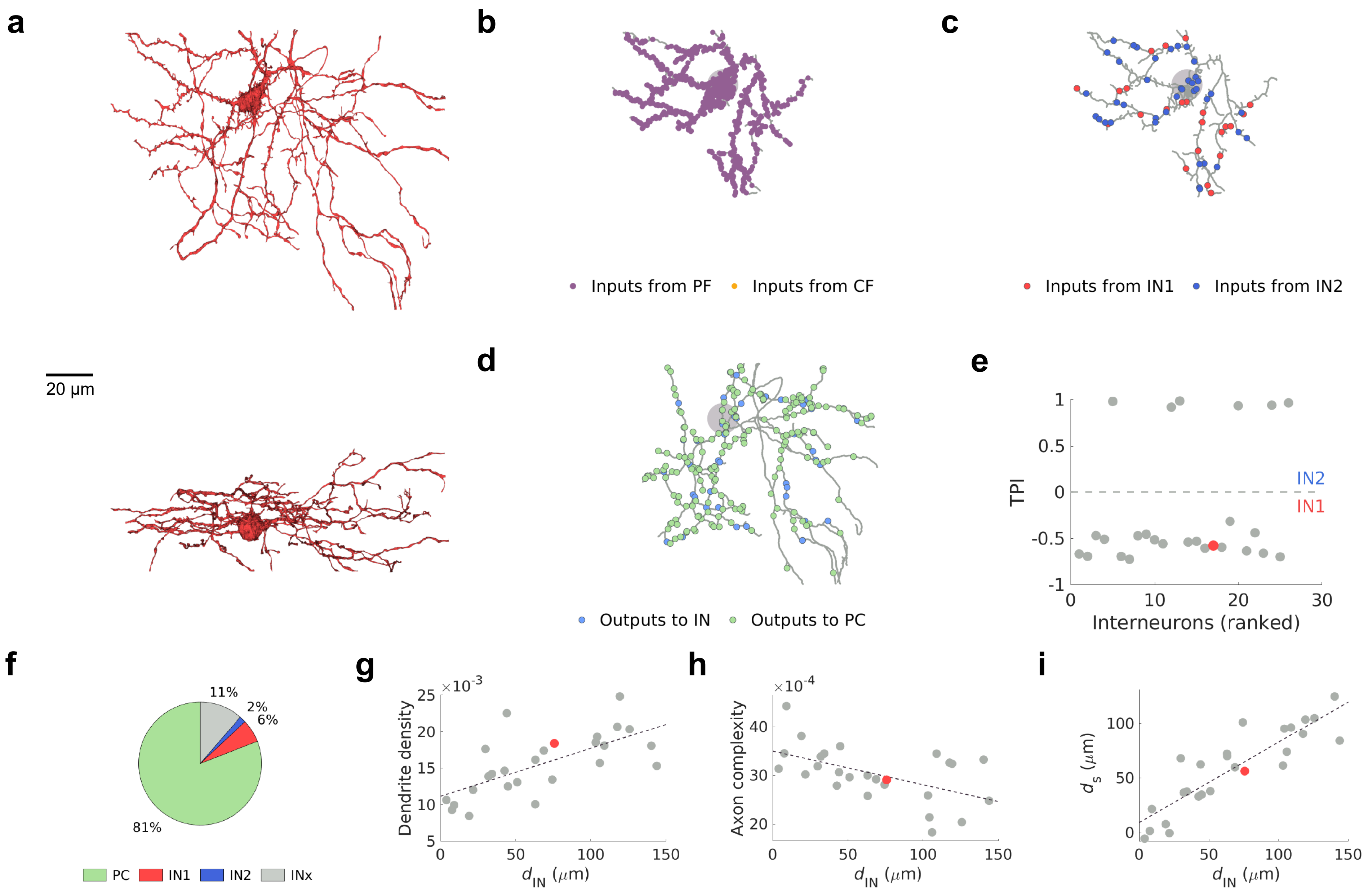
a
20 μm
b
Inputs from PF
Inputs from CF
c
Inputs from IN1
Inputs from IN2
d
Outputs to IN
Outputs to PC
e
TPI
IN2
IN1
Interneurons (ranked)
f
11%
2%
6%
81%
PC
IN1
IN2
INx
g
Dendrite density
×10-3
dIN (μm)
h
Axon complexity
×10-4
dIN (μm)
i
ds (μm)
dIN (μm)

IN1 (23)

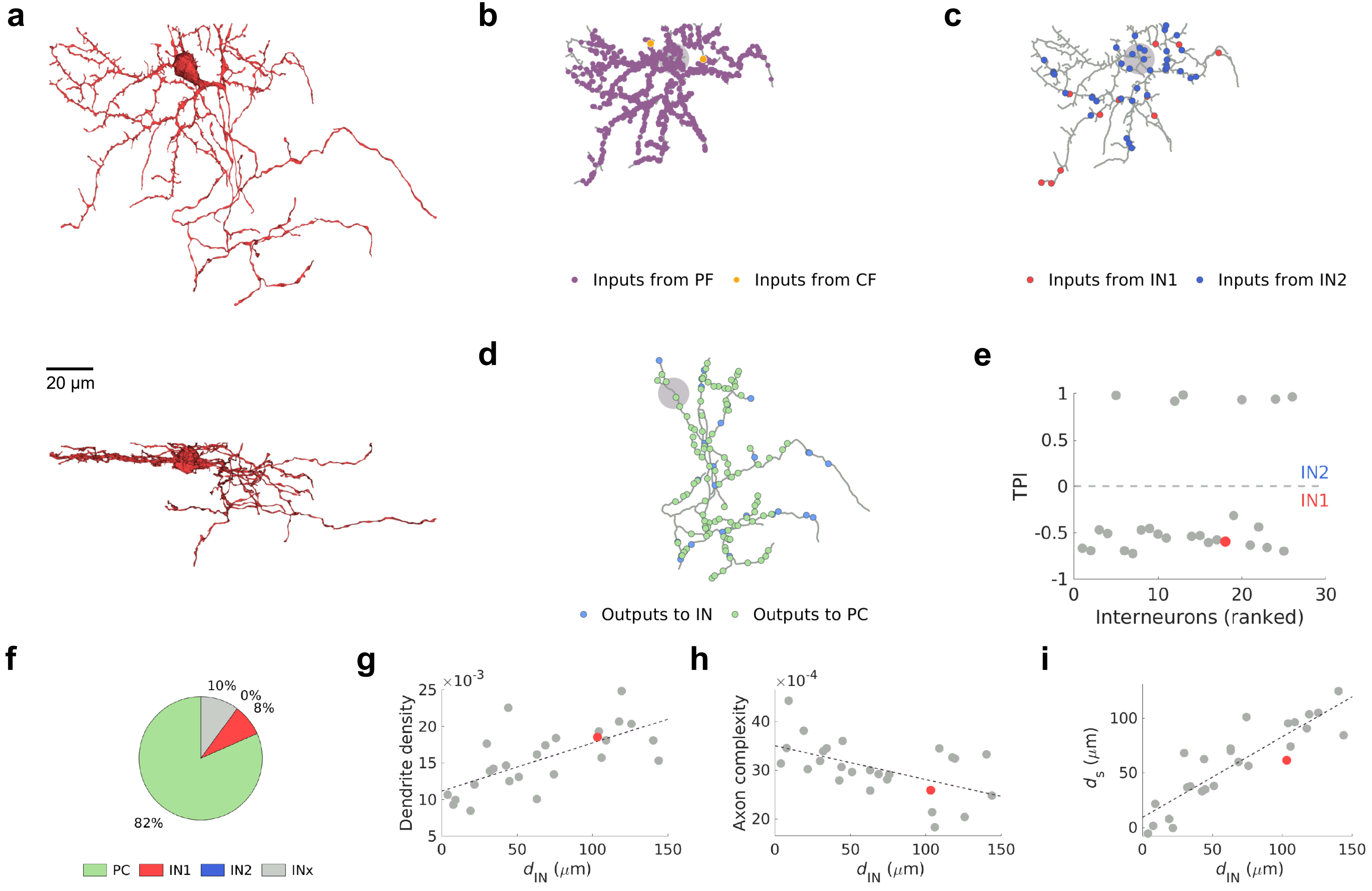
a
20 µm
b
Inputs from PF
Inputs from CF
c
Inputs from IN1
Inputs from IN2
d
Outputs to IN
Outputs to PC
e
TPI
IN2
IN1
Interneurons (ranked)
f
10%
0%
8%
82%
PC
IN1
IN2
INx
g
Dendrite density
×10^-3
d_IN (µm)
h
Axon complexity
×10^-4
d_IN (µm)
i
d_s (µm)
d_IN (µm)

IN1 (64)

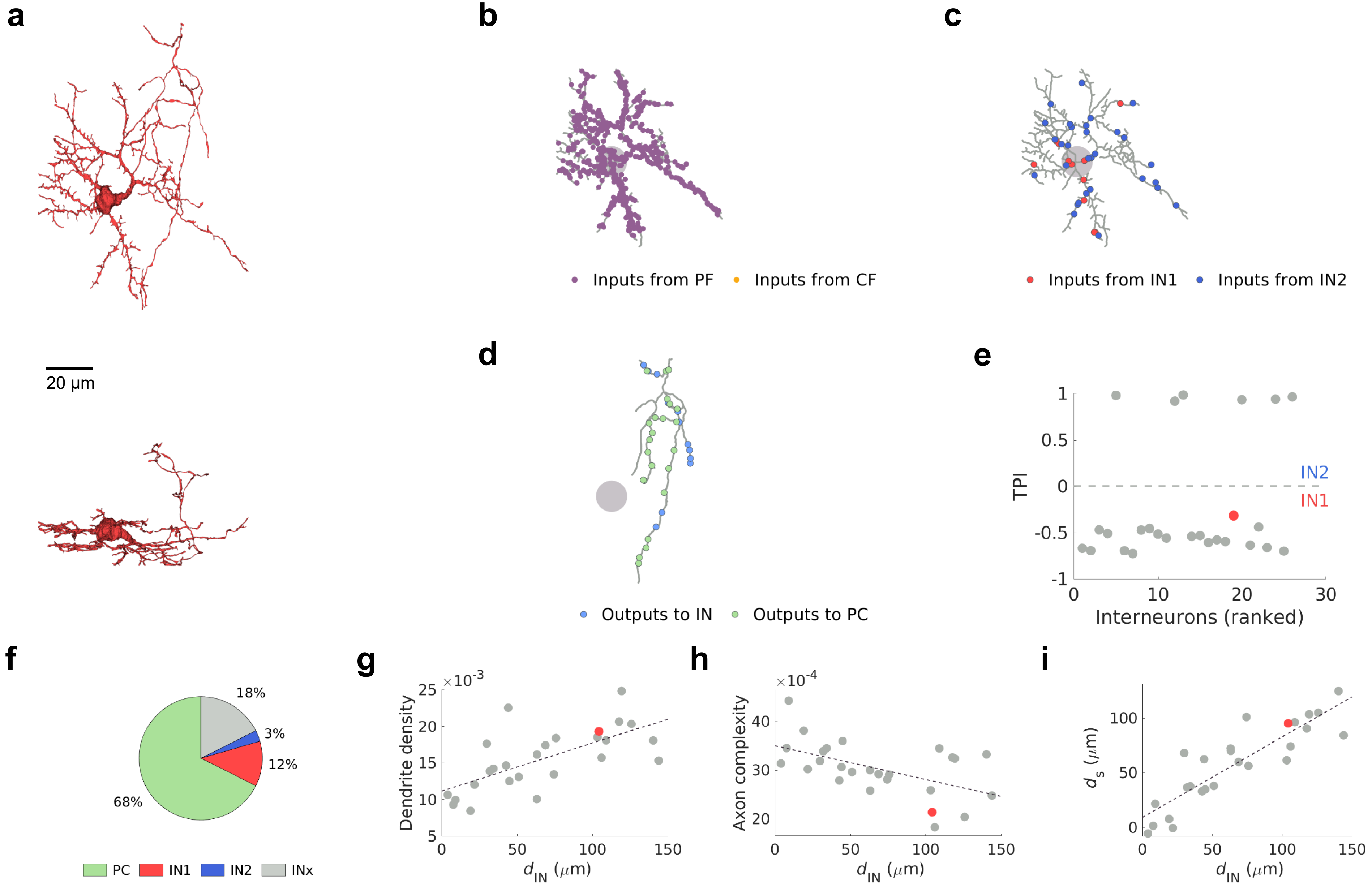

a
20 μm
b
Inputs from PF
Inputs from CF
c
Inputs from IN1
Inputs from IN2
d
Outputs to IN
Outputs to PC
e
TPI
IN2
IN1
Interneurons (ranked)
f
18%
3%
12%
68%
PC
IN1
IN2
INx
g
×10^-3
Dendrite density
d_IN (μm)
h
×10^-4
Axon complexity
d_IN (μm)
i
d_s (μm)
d_IN (μm)


IN1 (37)

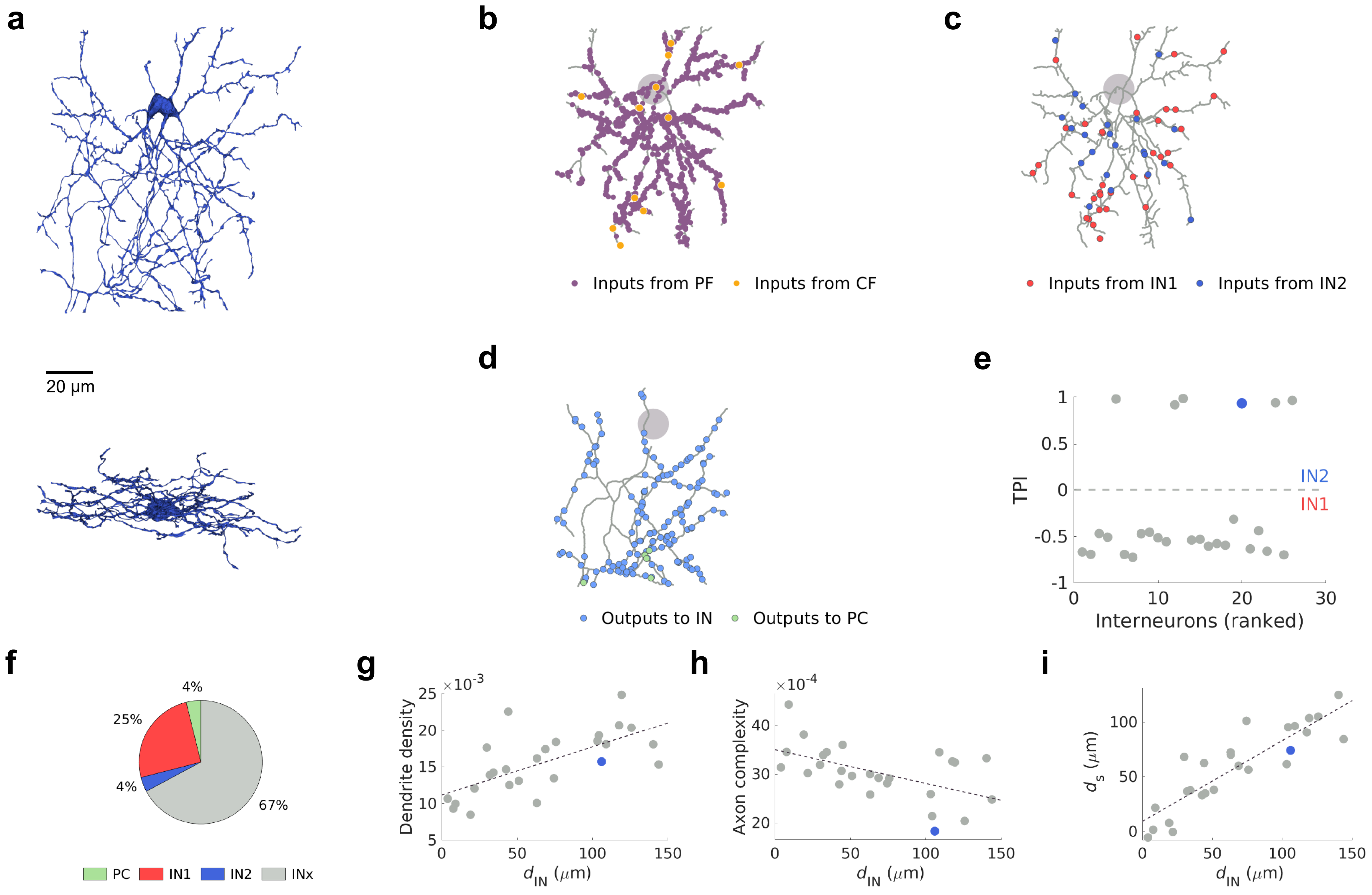
a
20 µm
b
Inputs from PF
Inputs from CF
c
Inputs from IN1
Inputs from IN2
d
Outputs to IN
Outputs to PC
e
TPI
IN2
IN1
Interneurons (ranked)
f
4%
25%
4%
67%
PC
IN1
IN2
INx
g
Dendrite density
×10-3
$d_{IN}$ ($\mu$m)
h
Axon complexity
×10-4
$d_{IN}$ ($\mu$m)
i
$d_s$ ($\mu$m)
$d_{IN}$ ($\mu$m)

IN2 (22)

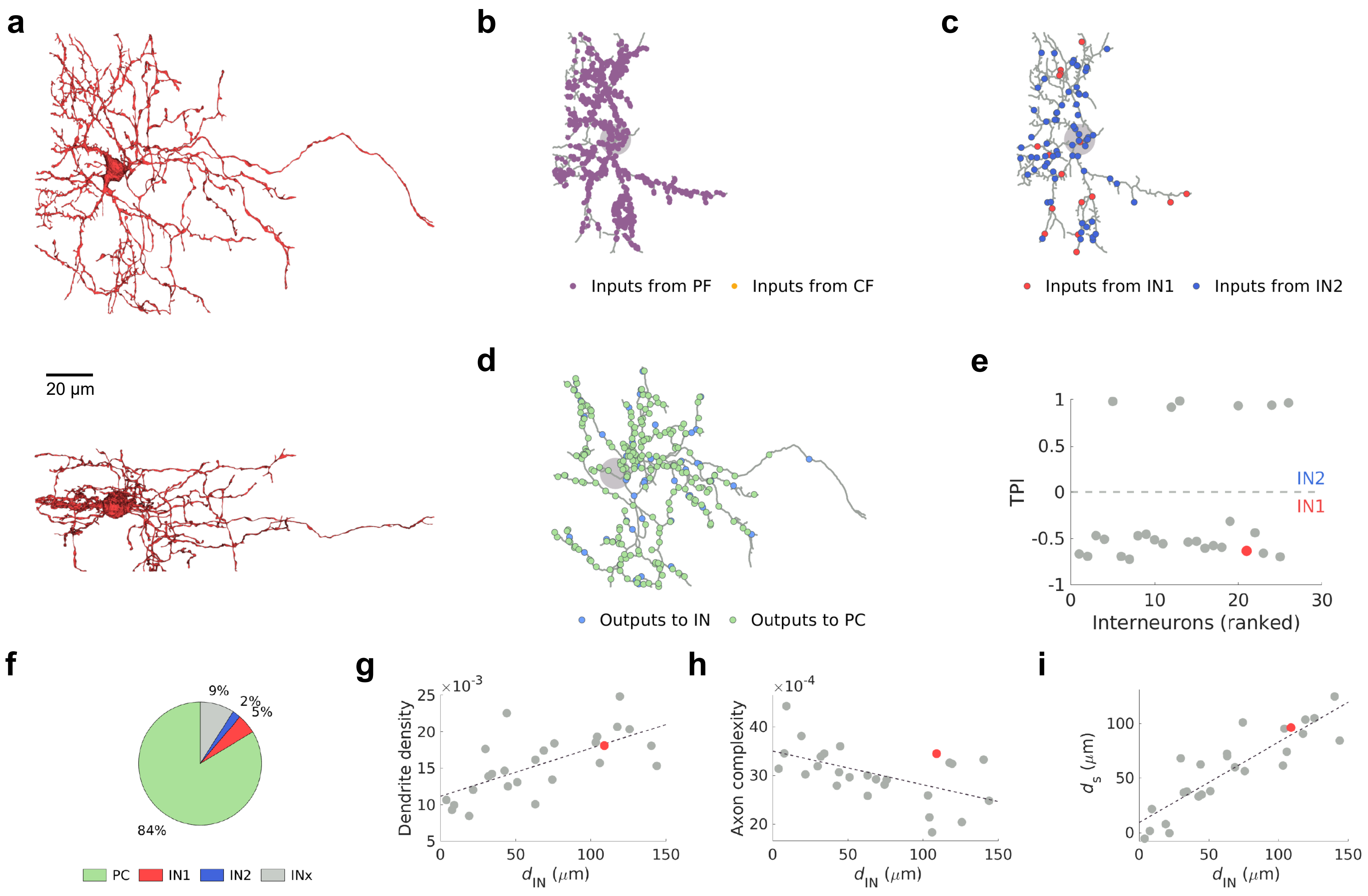

a
20 μm
b
Inputs from PF
Inputs from CF
c
Inputs from IN1
Inputs from IN2
d
Outputs to IN
Outputs to PC
e
TPI
IN2
IN1
Interneurons (ranked)
f
9%
2%
5%
84%
PC
IN1
IN2
INx
g
Dendrite density
$\times 10^{-3}$
$d_{IN}$ ($\mu$m)
h
Axon complexity
$\times 10^{-4}$
$d_{IN}$ ($\mu$m)
i
$d_s$ ($\mu$m)
$d_{IN}$ ($\mu$m)


IN1 (36)

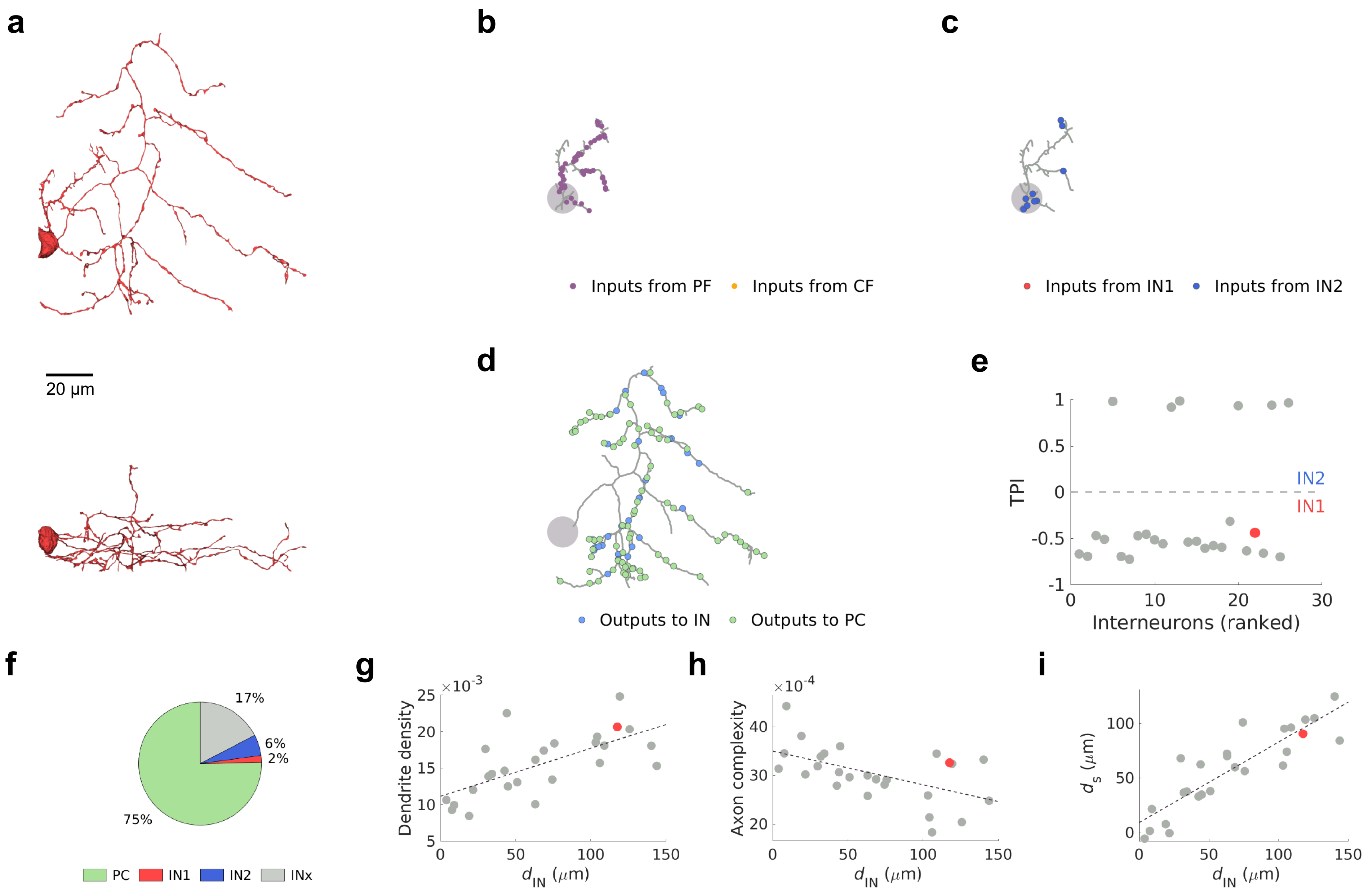
a
20 μm
b
Inputs from PF
Inputs from CF
c
Inputs from IN1
Inputs from IN2
d
Outputs to IN
Outputs to PC
e
TPI
IN2
IN1
Interneurons (ranked)
f
17%
6%
2%
75%
PC
IN1
IN2
INx
g
×10^-3
Dendrite density
$d_{IN}$ (μm)
h
×10^-4
Axon complexity
$d_{IN}$ (μm)
i
$d_s$ (μm)
$d_{IN}$ (μm)

IN1 (47)

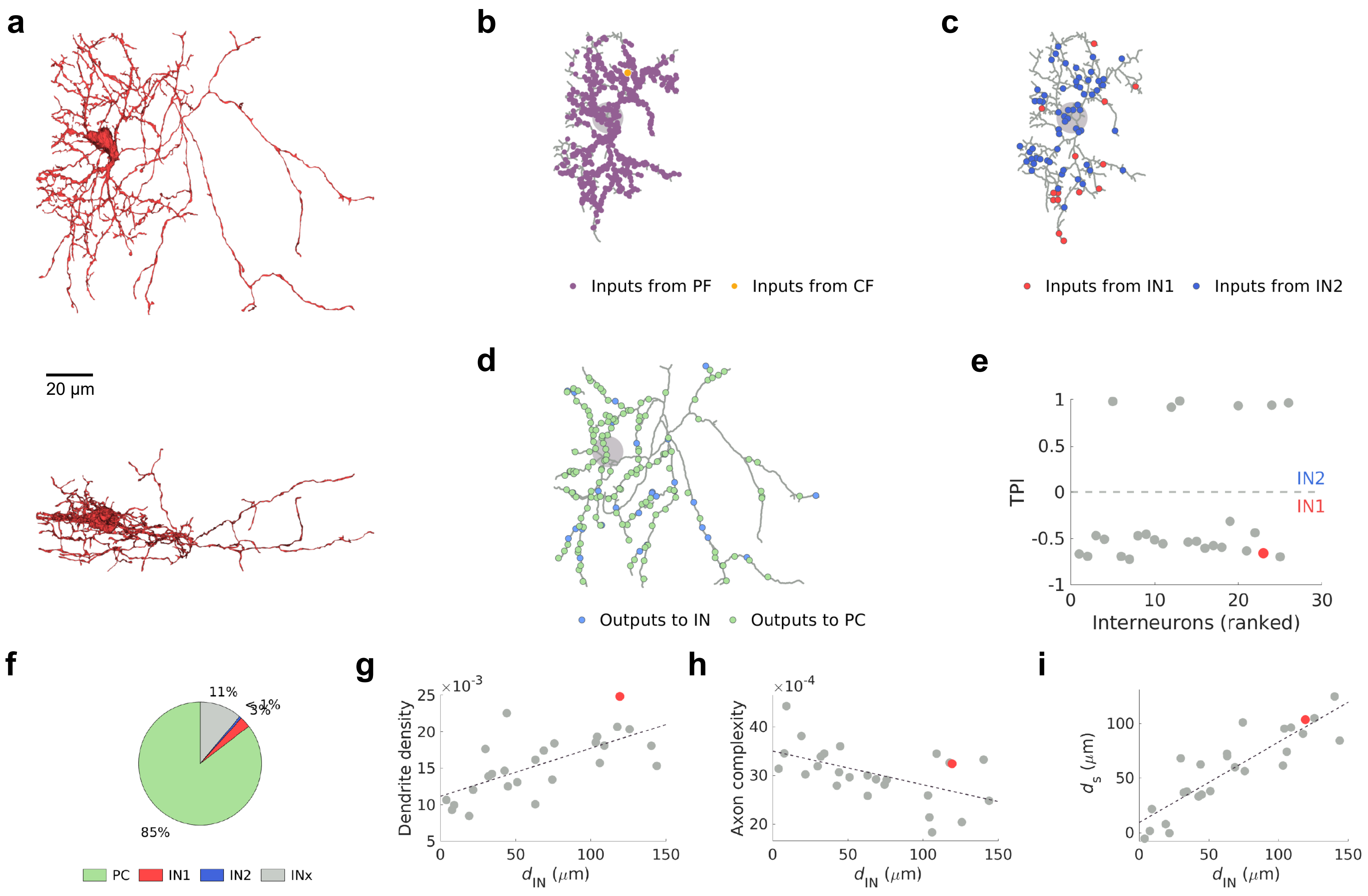
a
20 μm
b
Inputs from PF
Inputs from CF
c
Inputs from IN1
Inputs from IN2
d
Outputs to IN
Outputs to PC
e
TPI
IN2
IN1
Interneurons (ranked)
f
11%
85%
PC
IN1
IN2
INx
g
Dendrite density
$d_{IN}$ ($\mu$m)
h
Axon complexity
$d_{IN}$ ($\mu$m)
i
$d_s$ ($\mu$m)
$d_{IN}$ ($\mu$m)

IN1 (35)

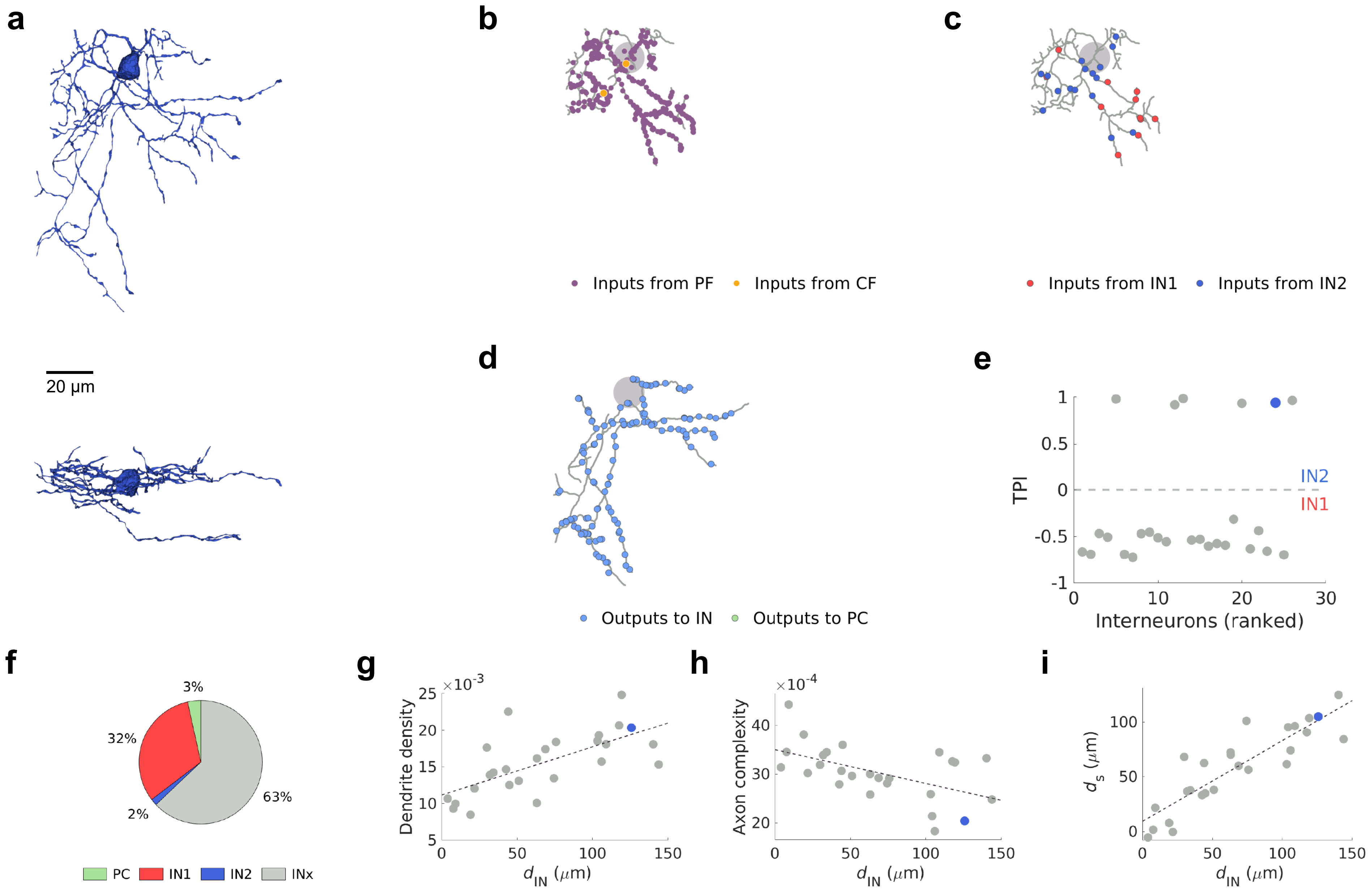

a
20 μm
b
Inputs from PF
Inputs from CF
c
Inputs from IN1
Inputs from IN2
d
Outputs to IN
Outputs to PC
e
TPI
IN2
IN1
Interneurons (ranked)
f
3%
32%
63%
2%
PC
IN1
IN2
INx
g
Dendrite density
$\times10^{-3}$
$d_{IN}$ ($\mu$m)
h
Axon complexity
$\times10^{-4}$
$d_{IN}$ ($\mu$m)
i
$d_s$ ($\mu$m)
$d_{IN}$ ($\mu$m)


IN2 (40)

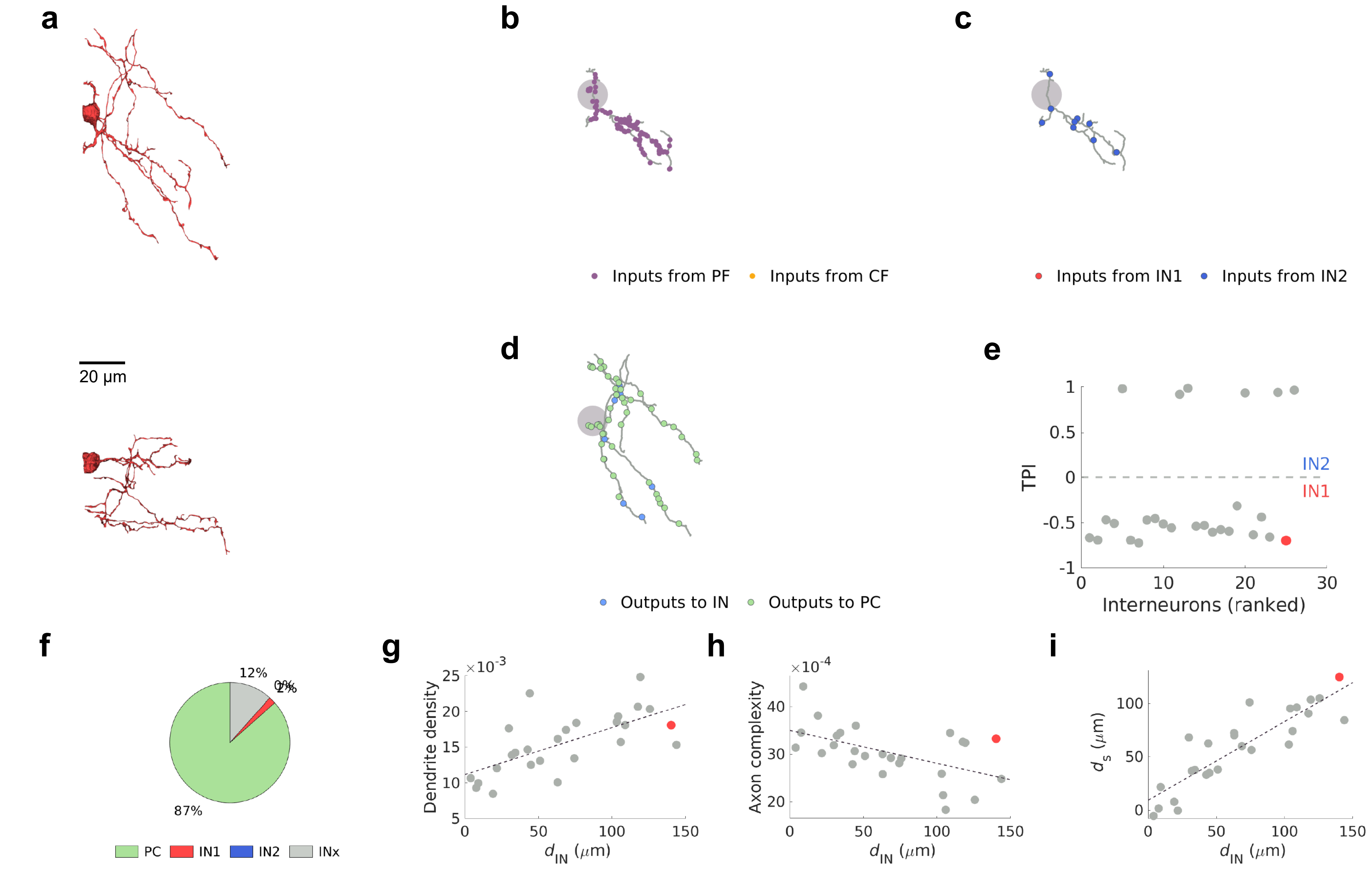
a
b
c
20 µm
Inputs from PF
Inputs from CF
Inputs from IN1
Inputs from IN2
d
e
Outputs to IN
Outputs to PC
TPI
IN2
IN1
Interneurons (ranked)
f
12%
0%
2%
87%
PC
IN1
IN2
INx
g
Dendrite density
×10^-3
d_IN (µm)
h
Axon complexity
×10^-4
d_IN (µm)
i
d_s (µm)
d_IN (µm)

IN1 (46)

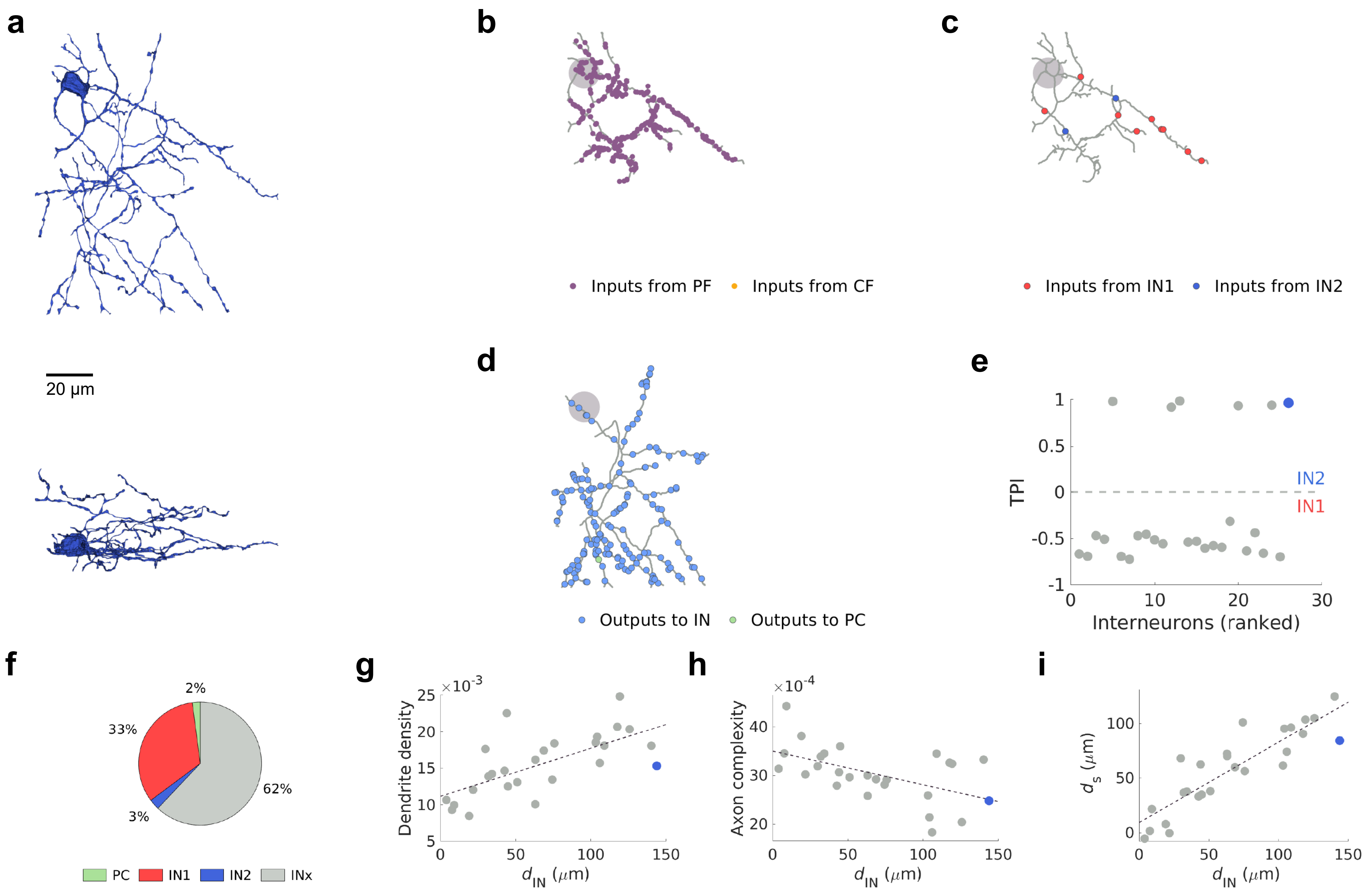

a
20 μm
b
Inputs from PF
Inputs from CF
c
Inputs from IN1
Inputs from IN2
d
Outputs to IN
Outputs to PC
e
TPI
IN2
IN1
Interneurons (ranked)
f
2%
33%
62%
3%
PC
IN1
IN2
INx
g
Dendrite density
×10^-3
d_IN (μm)
h
Axon complexity
×10^-4
d_IN (μm)
i
d_s (μm)
d_IN (μm)


IN2 (41)
