## Supplementary Data 5 for "A cerebellar disinhibitory circuit supports synaptic plasticity"

4308, 1062, 186

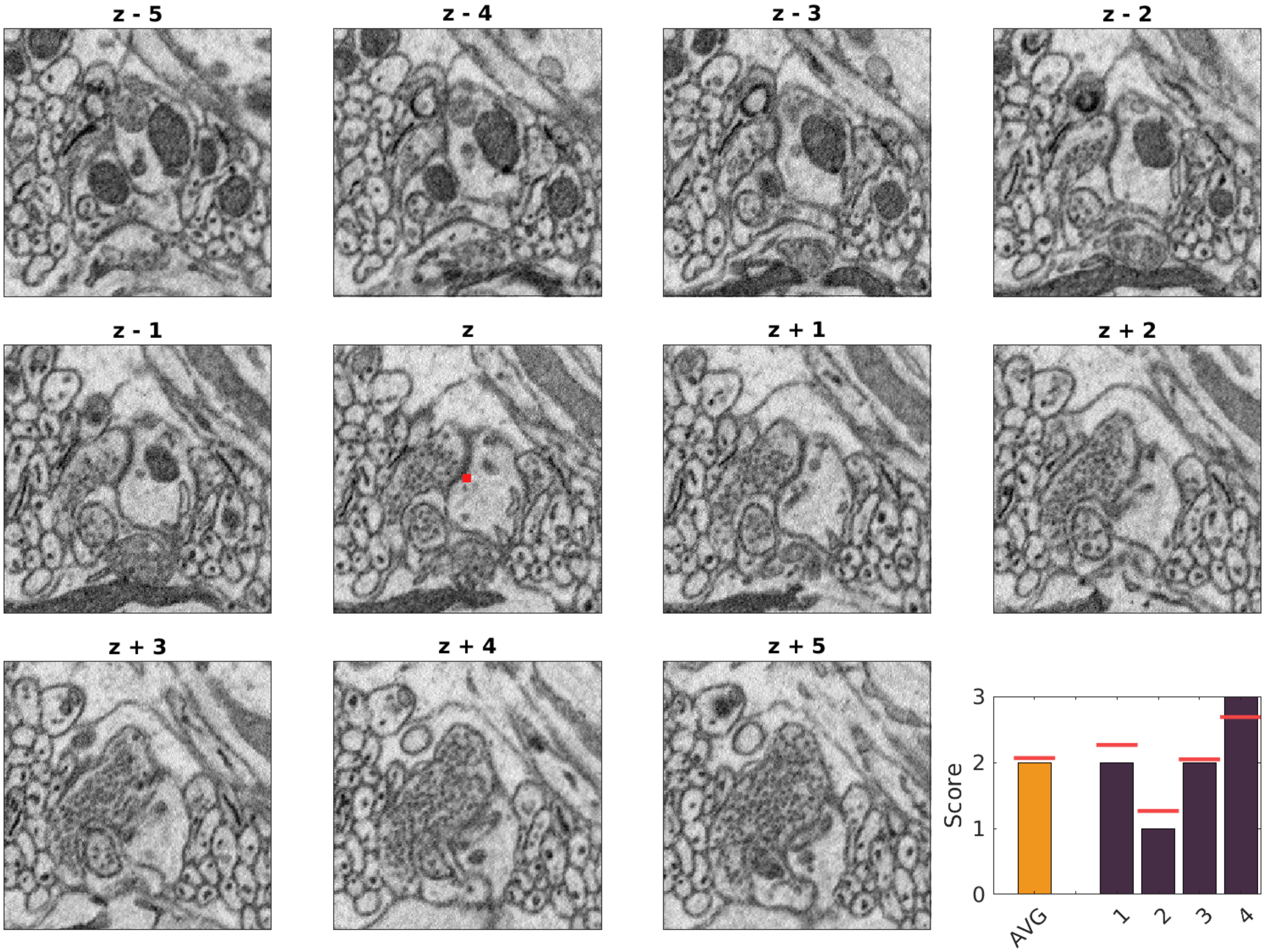

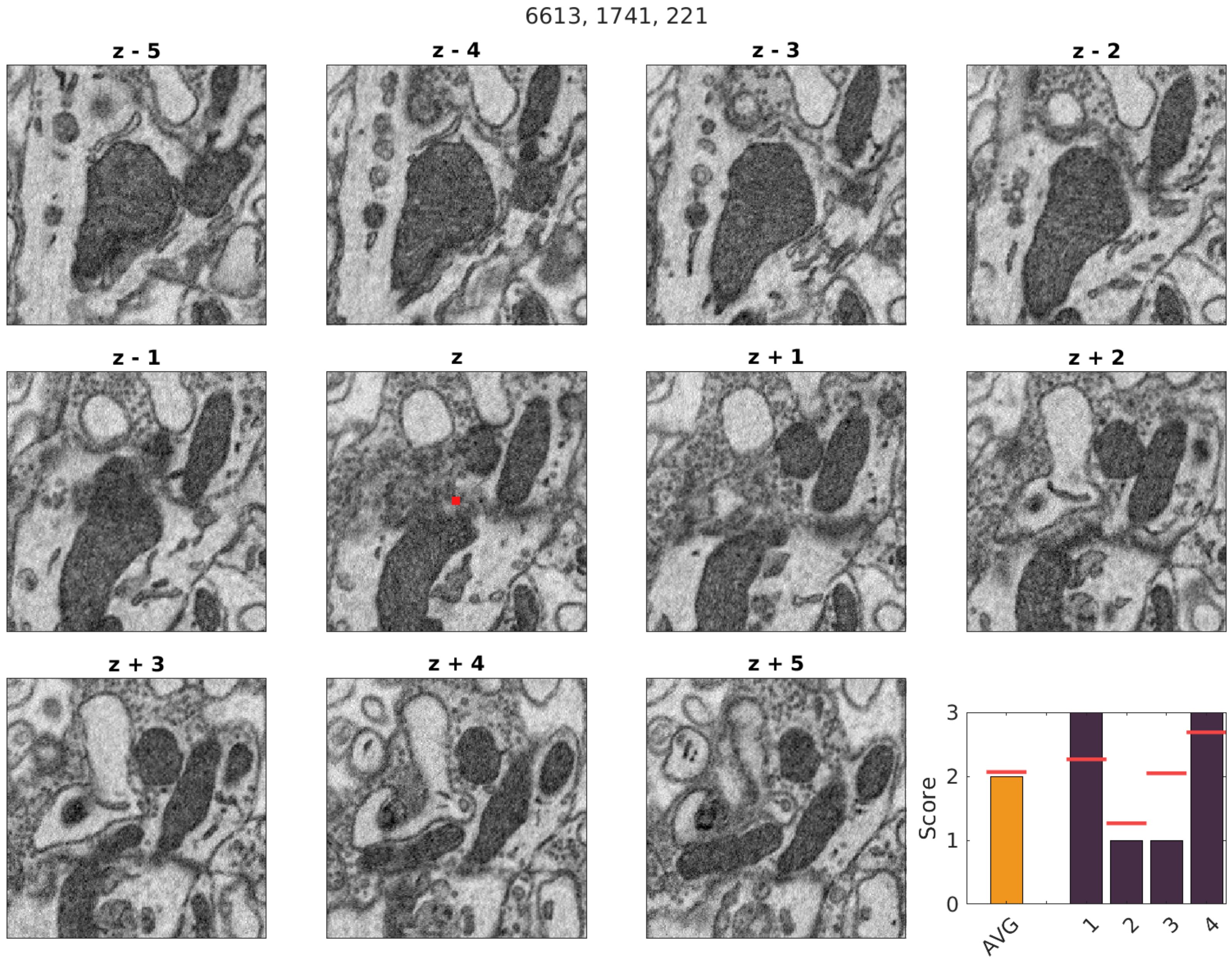

6613, 1741, 221
z - 5
z - 4
z - 3
z - 2
z - 1
z
z + 1
z + 2
z + 3
z + 4
z + 5
Score
0
1
2
3
AVG
1
2
3
4

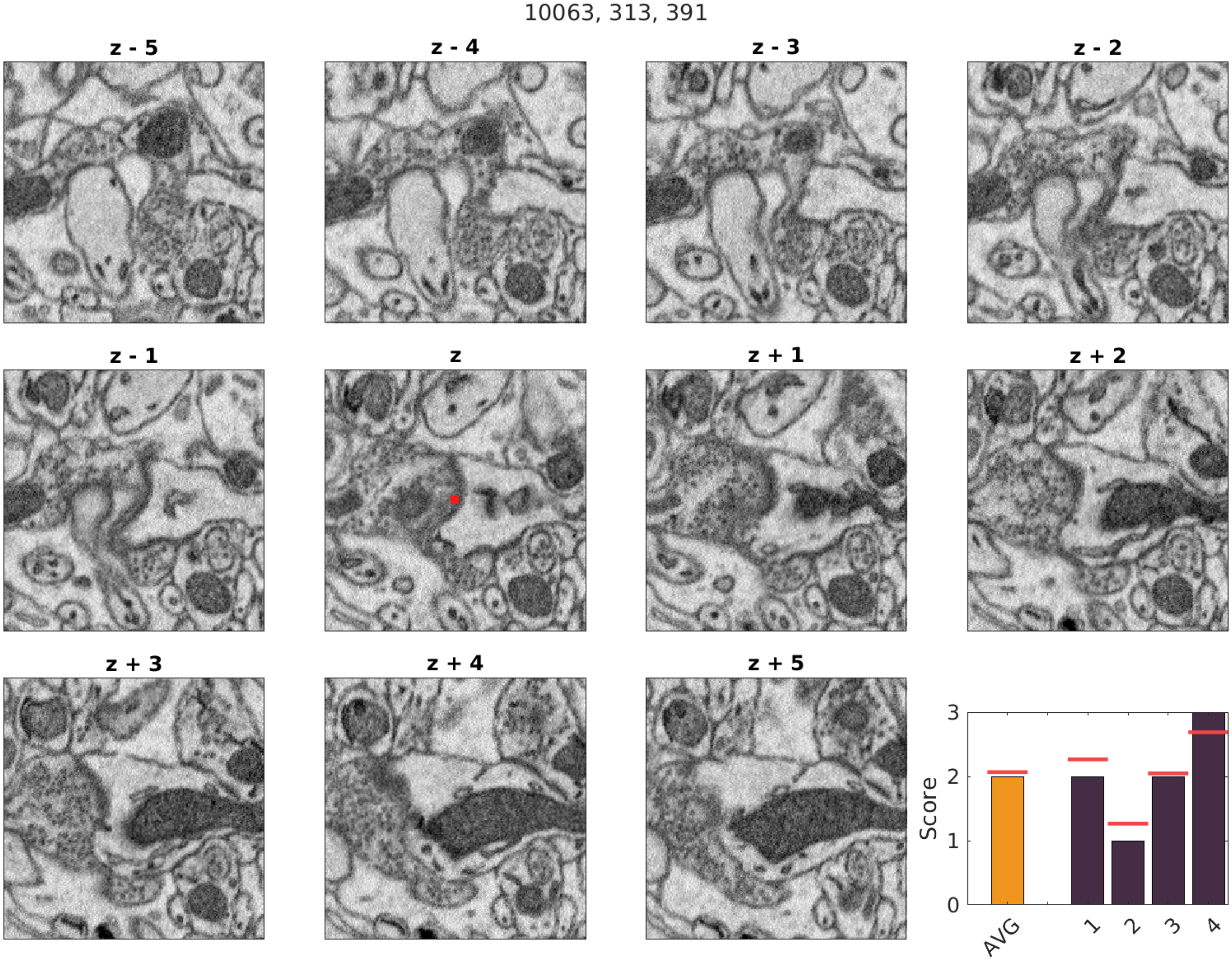

10063, 313, 391
z - 5
z - 4
z - 3
z - 2
z - 1
z
z + 1
z + 2
z + 3
z + 4
z + 5
Score
AVG
1
2
3
4

9669, 3465, 331

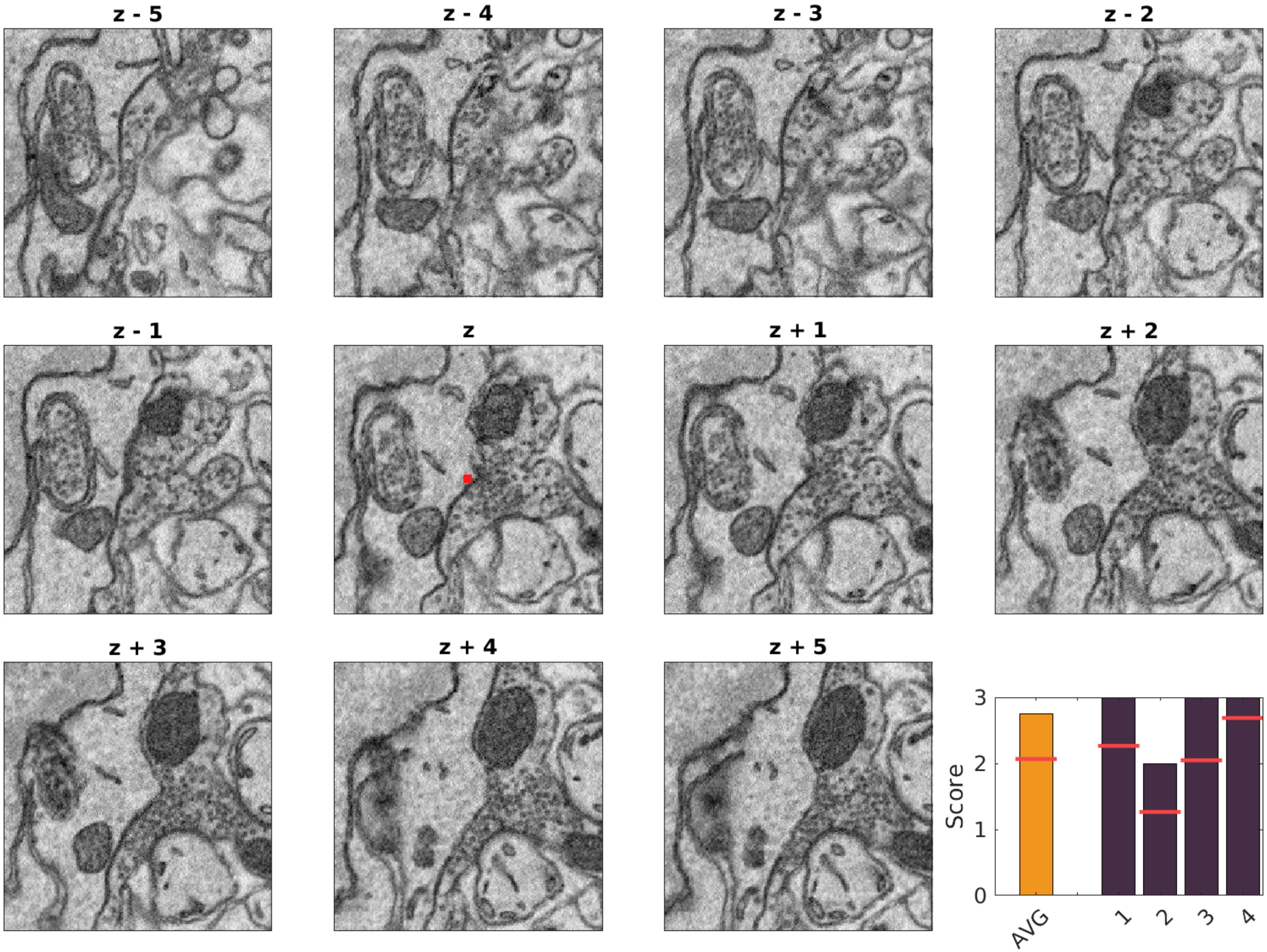

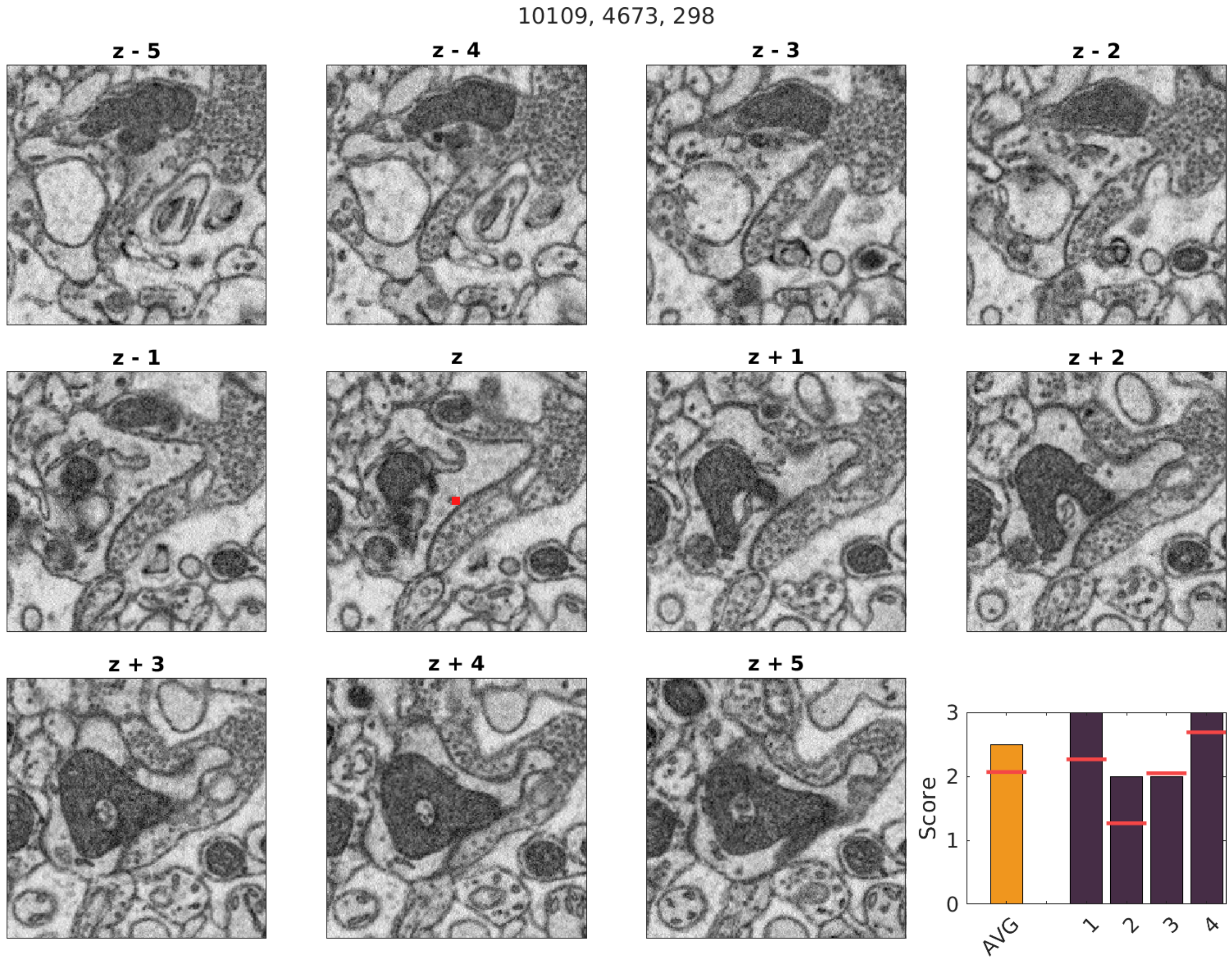
10109, 4673, 298
z - 5
z - 4
z - 3
z - 2
z - 1
z
z + 1
z + 2
z + 3
z + 4
z + 5
Score
AVG
1
2
3
4

10320, 4587, 310

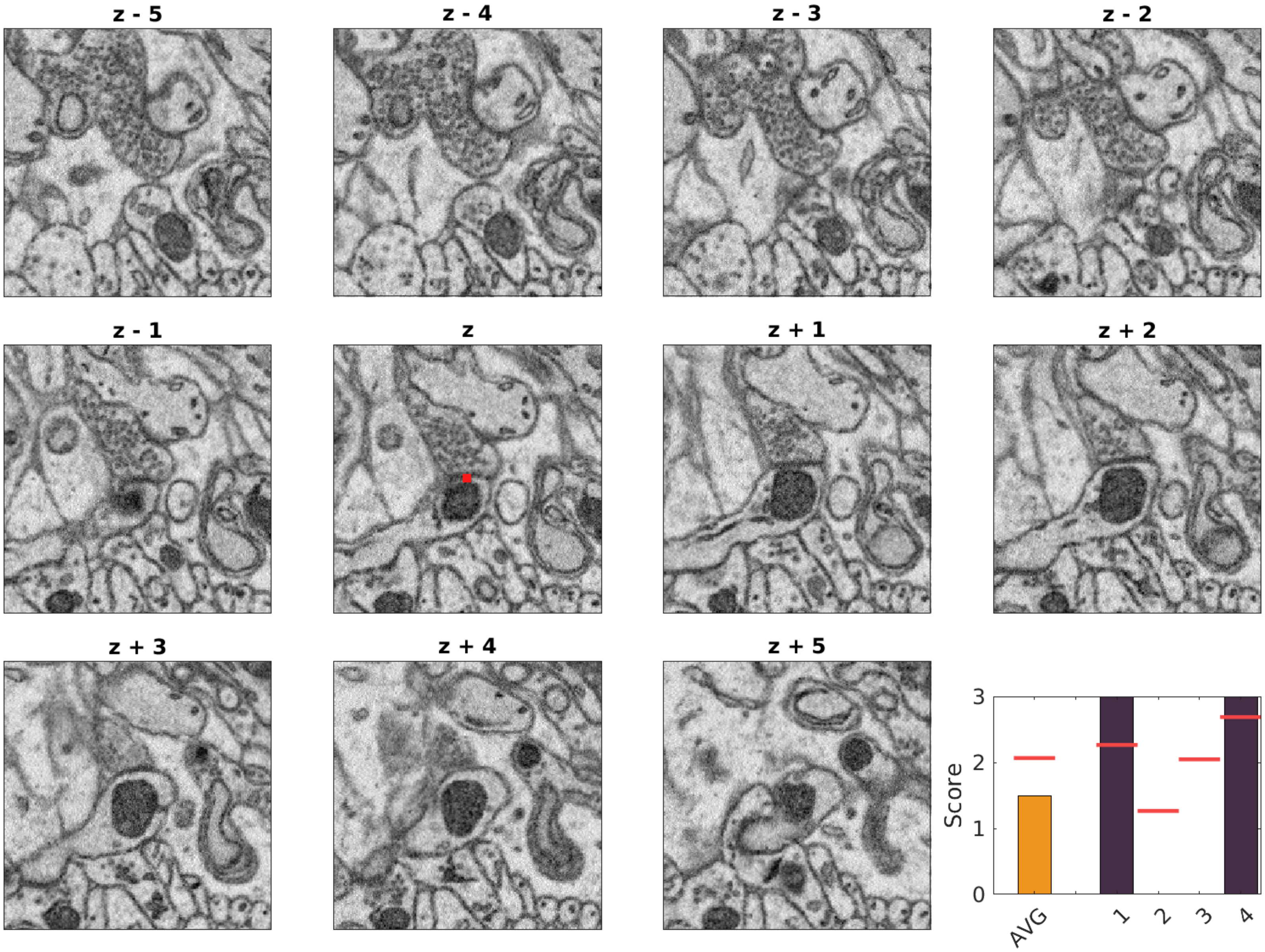

10886, 5440, 311

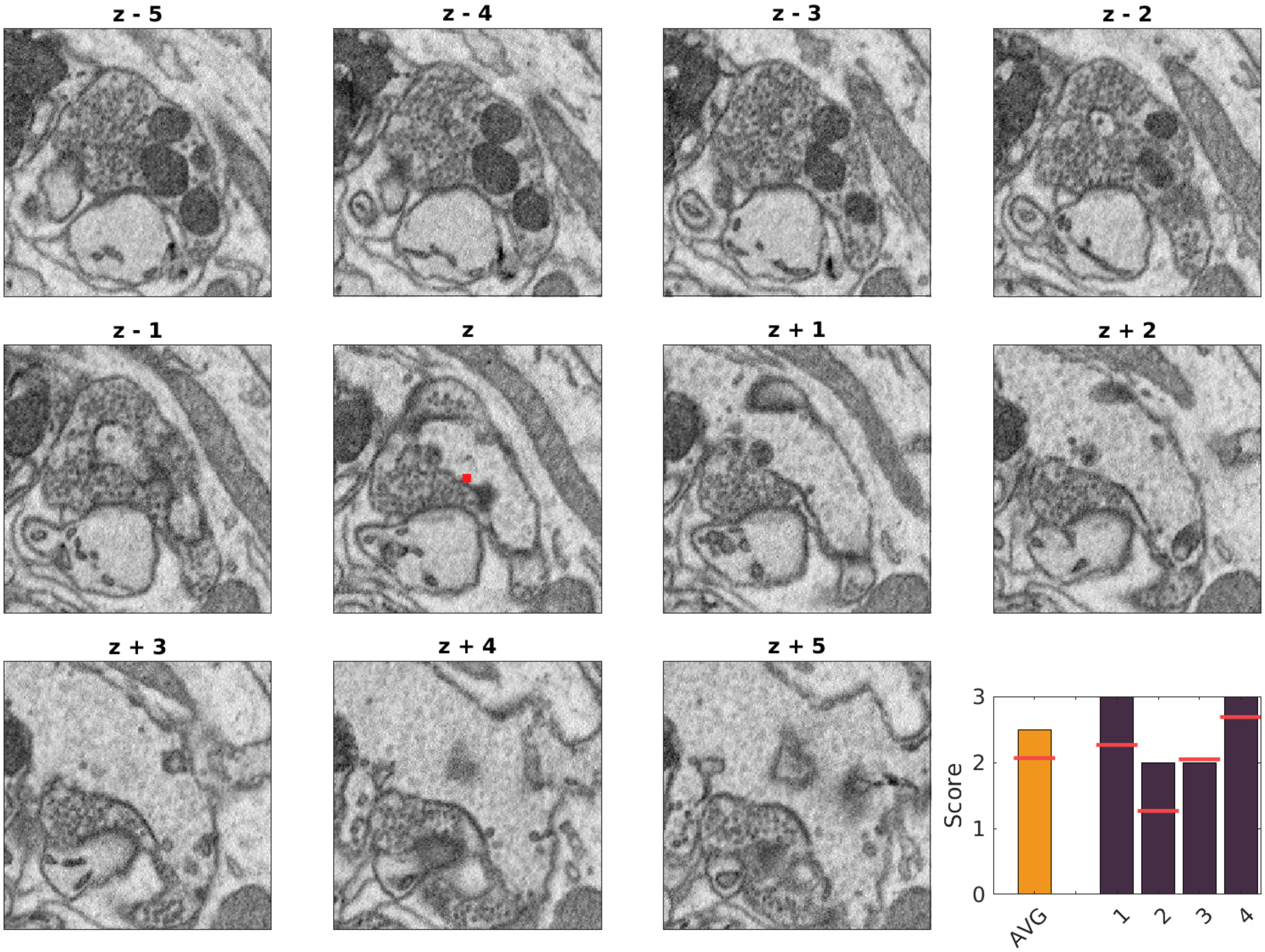

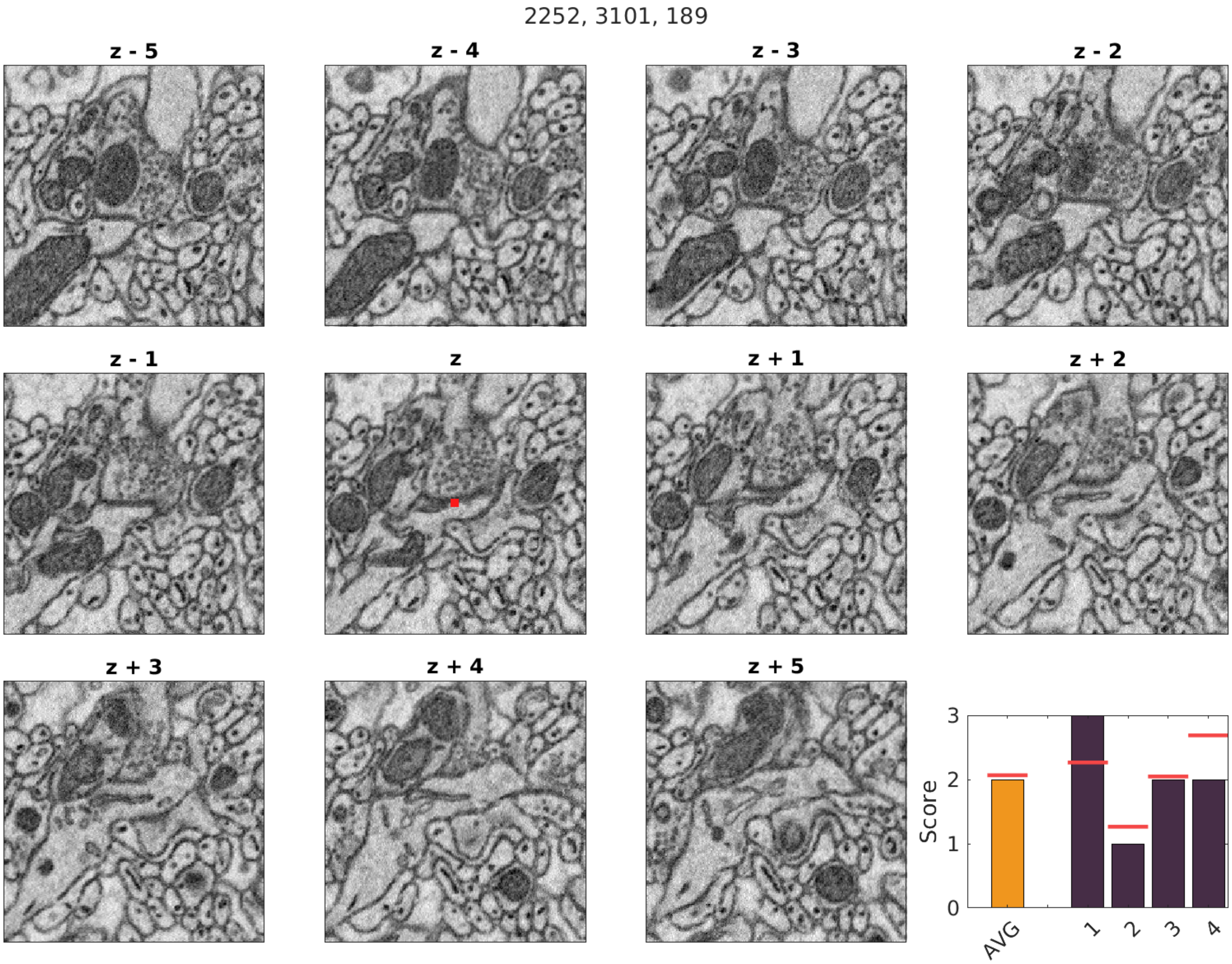
2252, 3101, 189
z - 5
z - 4
z - 3
z - 2
z - 1
z
z + 1
z + 2
z + 3
z + 4
z + 5
Score
0
1
2
3
AVG
1
2
3
4

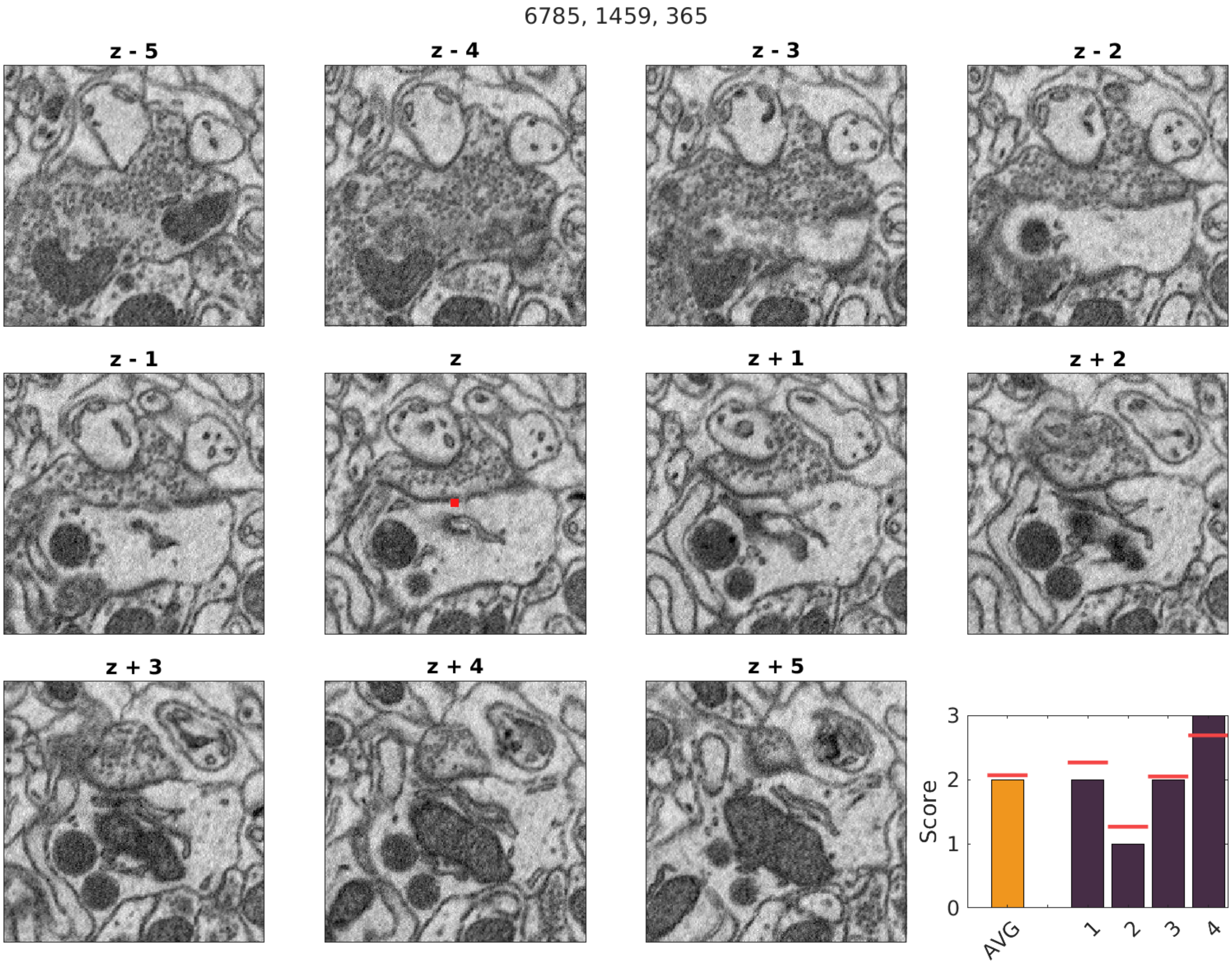

6785, 1459, 365
z - 5
z - 4
z - 3
z - 2
z - 1
z
z + 1
z + 2
z + 3
z + 4
z + 5
Score
0
1
2
3
AVG
1
2
3
4

7280, 494, 391

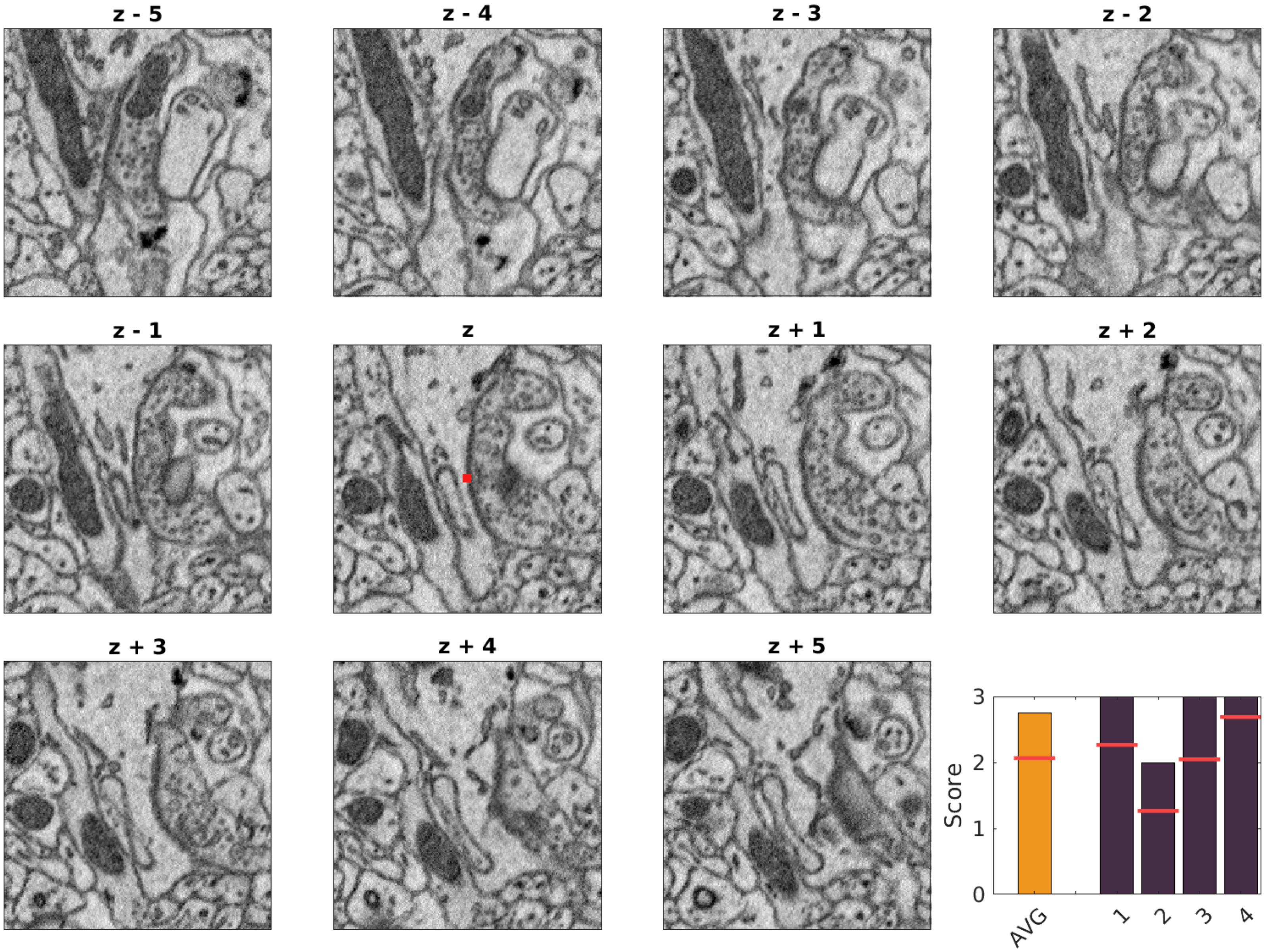

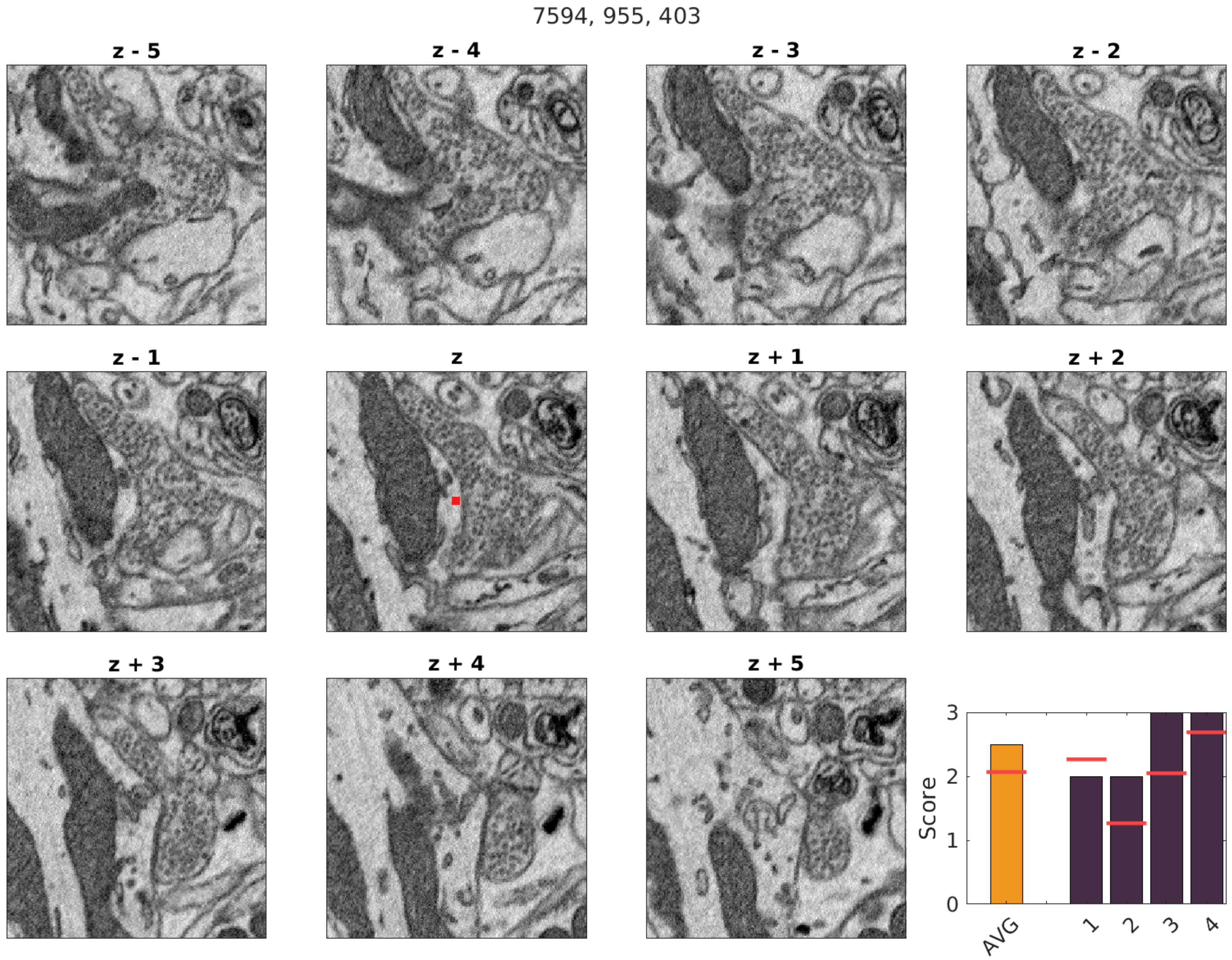

7594, 955, 403
z - 5
z - 4
z - 3
z - 2
z - 1
z
z + 1
z + 2
z + 3
z + 4
z + 5
Score
0
1
2
3
AVG
1
2
3
4

8611, 930, 402

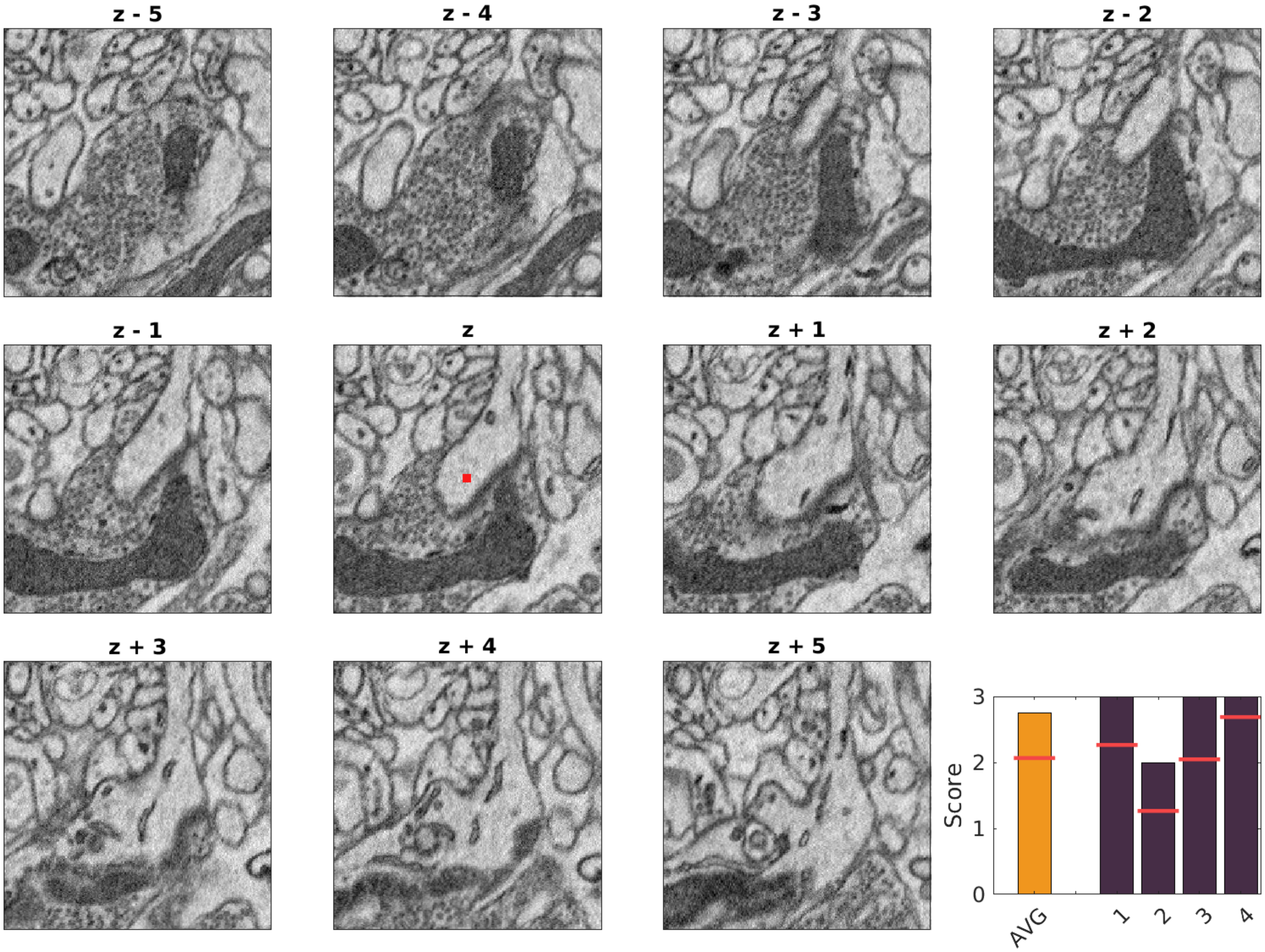

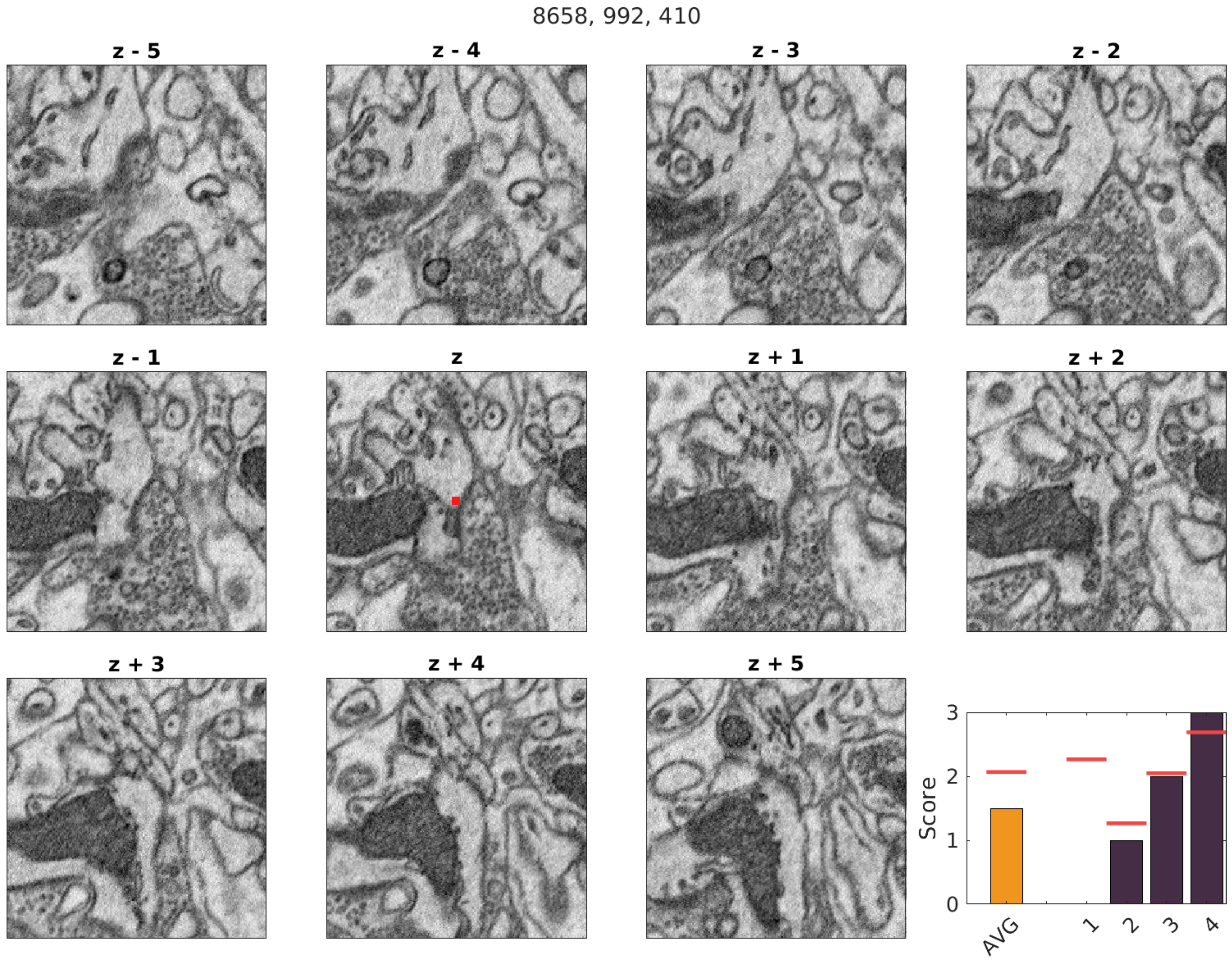

8658, 992, 410
z - 5
z - 4
z - 3
z - 2
z - 1
z
z + 1
z + 2
z + 3
z + 4
z + 5
Score
0
1
2
3
AVG
1
2
3
4

3390, 2045, 706

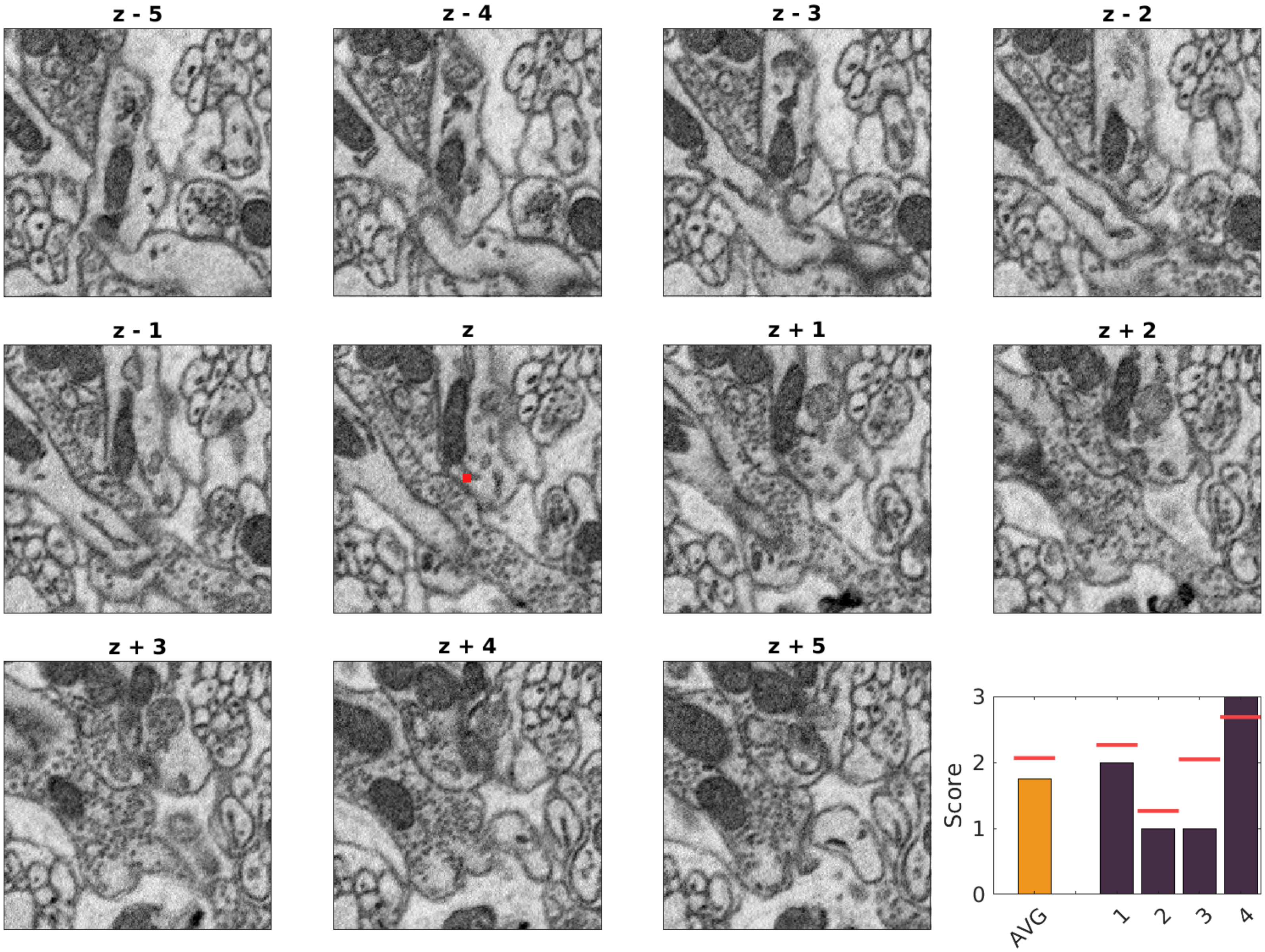

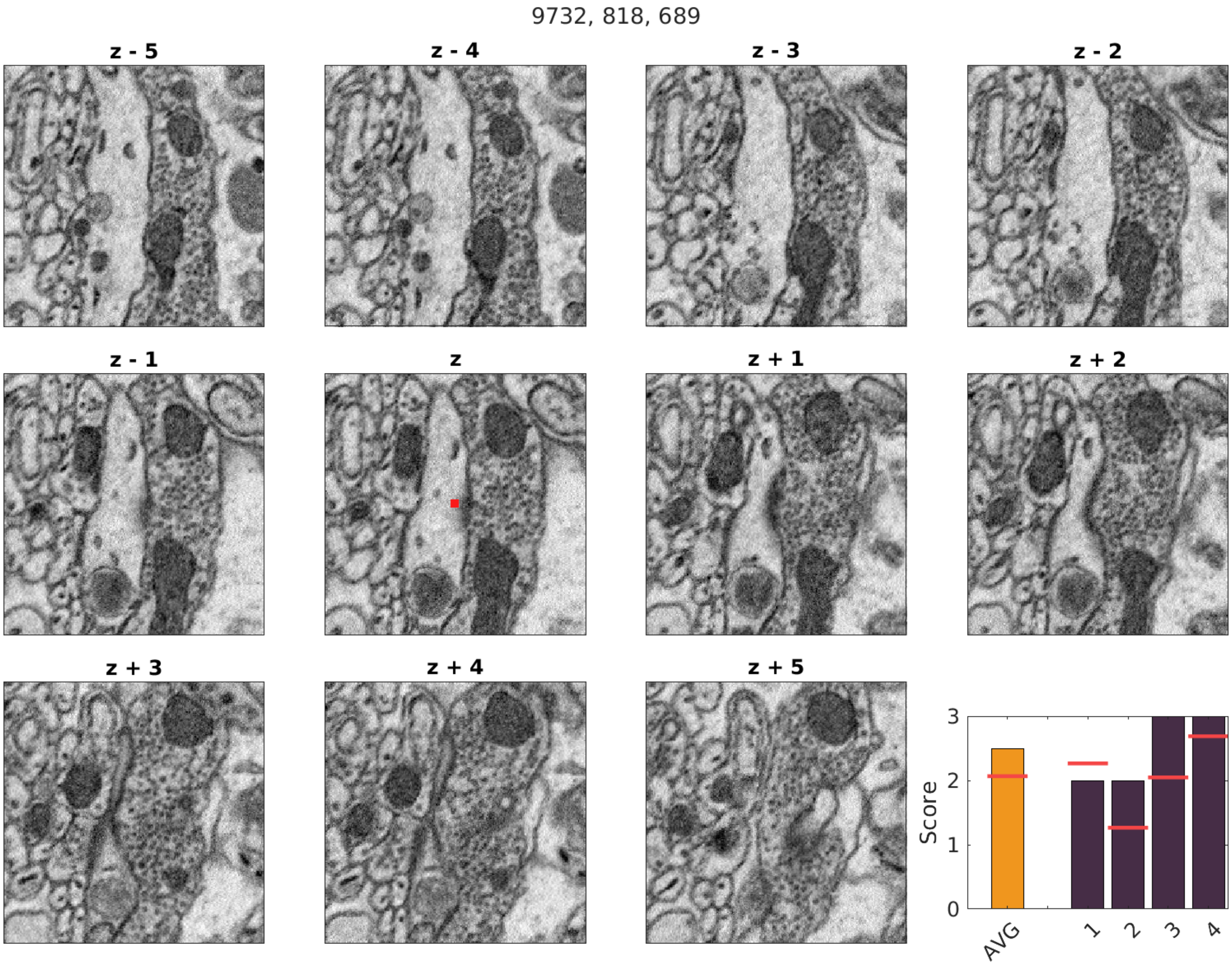
9732, 818, 689
z - 5
z - 4
z - 3
z - 2
z - 1
z
z + 1
z + 2
z + 3
z + 4
z + 5
Score
0
1
2
3
AVG
1
2
3
4

9414, 4607, 565

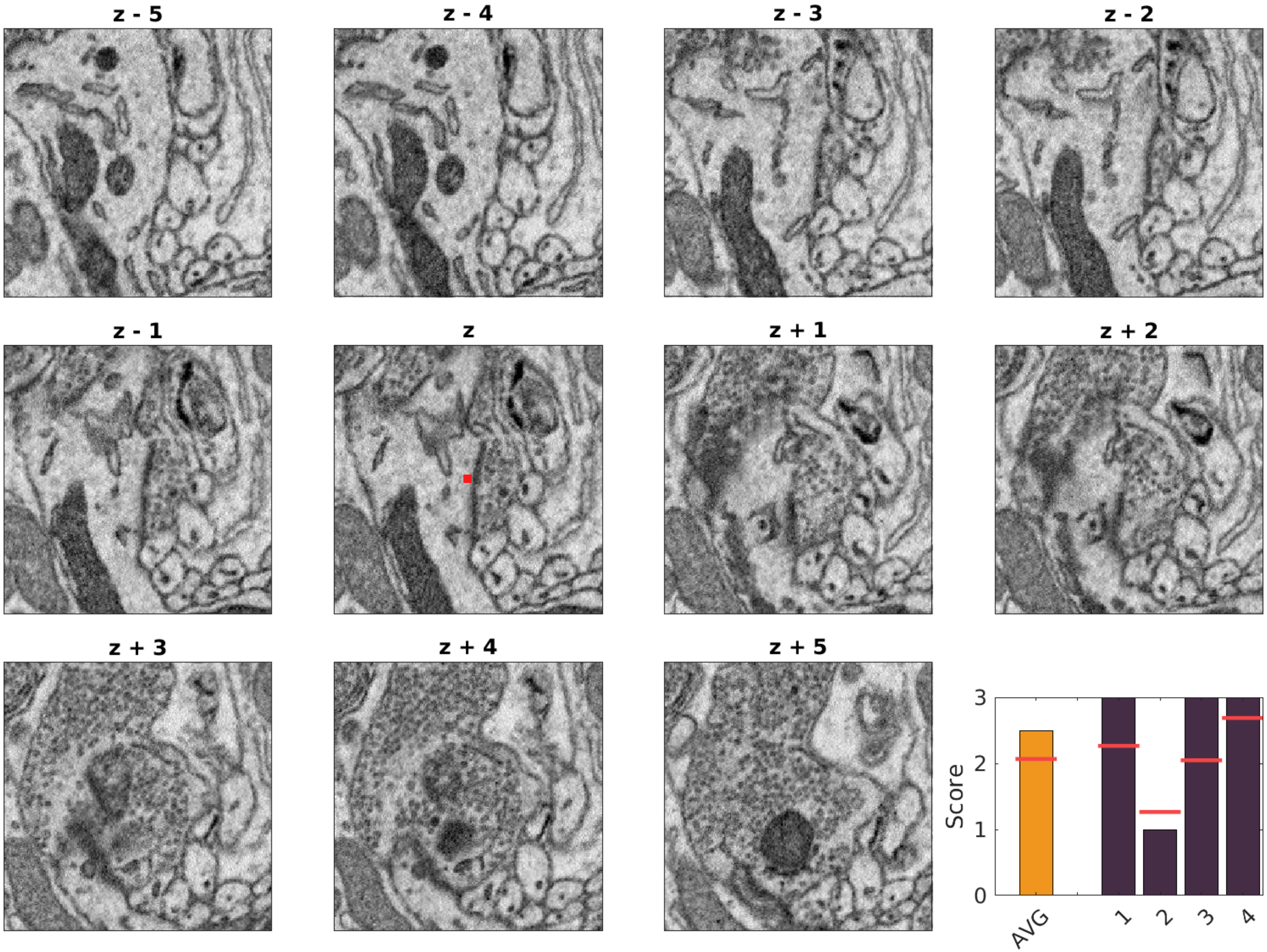

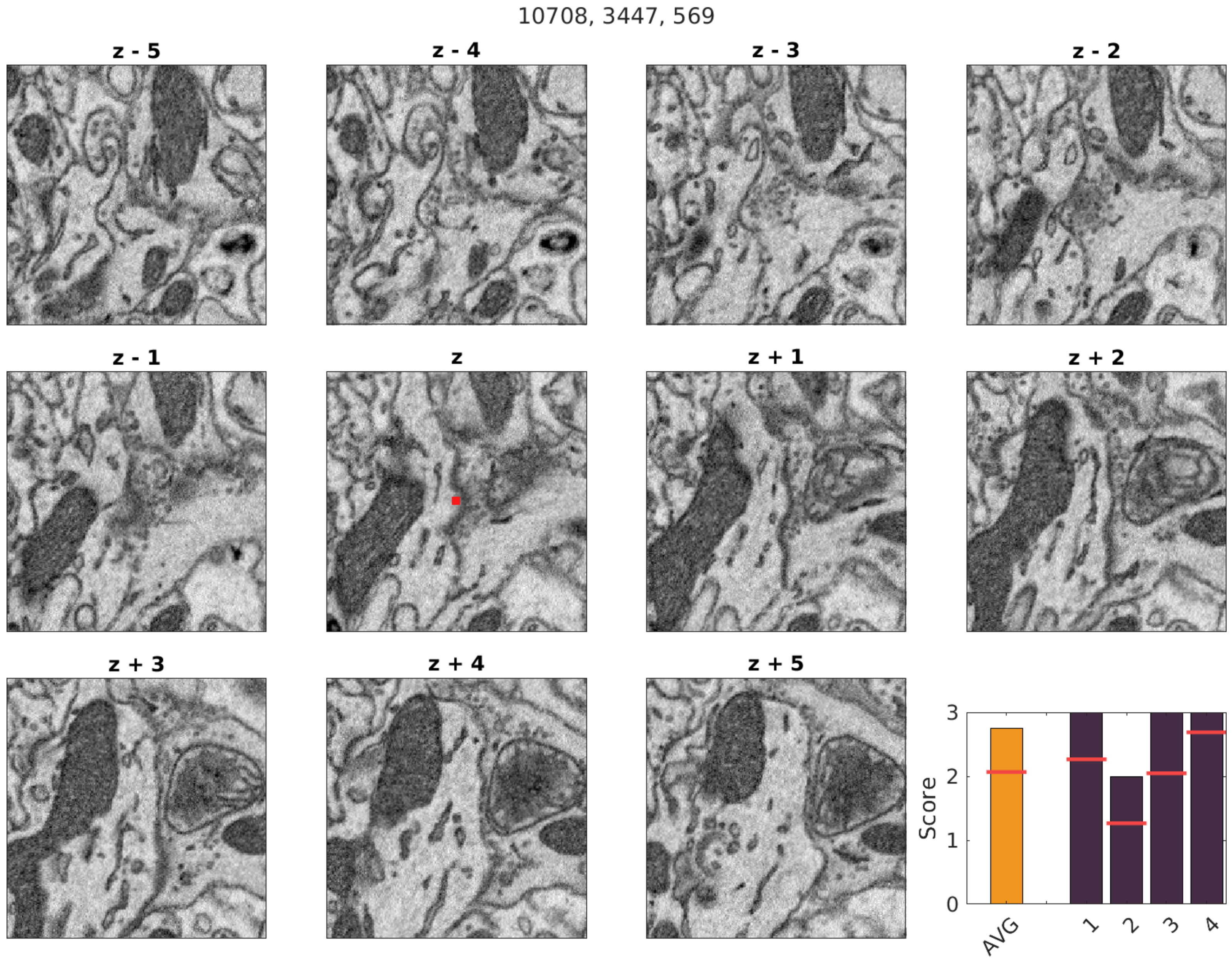
10708, 3447, 569
z - 5
z - 4
z - 3
z - 2
z - 1
z
z + 1
z + 2
z + 3
z + 4
z + 5
Score
AVG
1
2
3
4
0
1
2
3

11962, 2703, 619

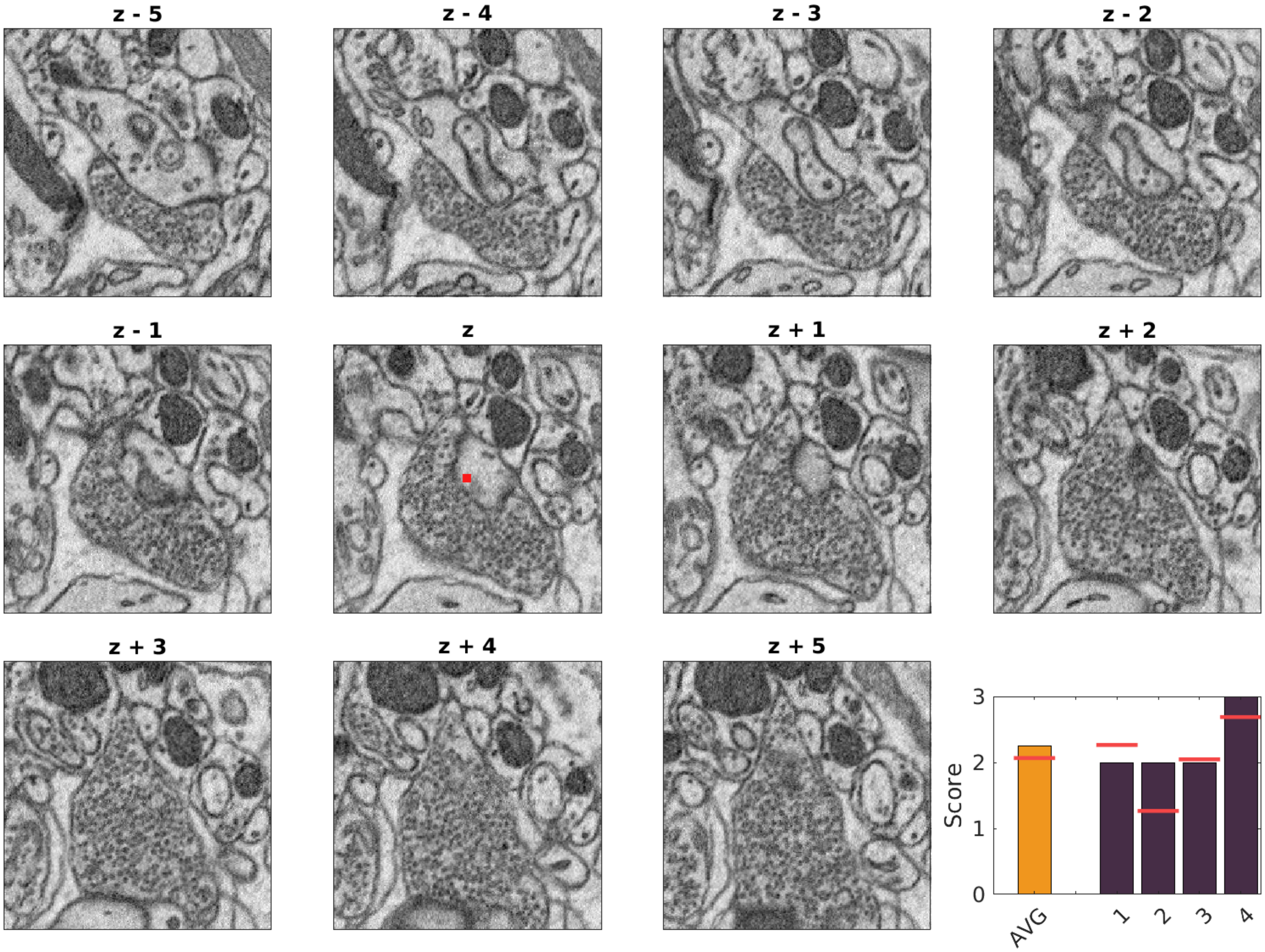

12075, 3113, 613

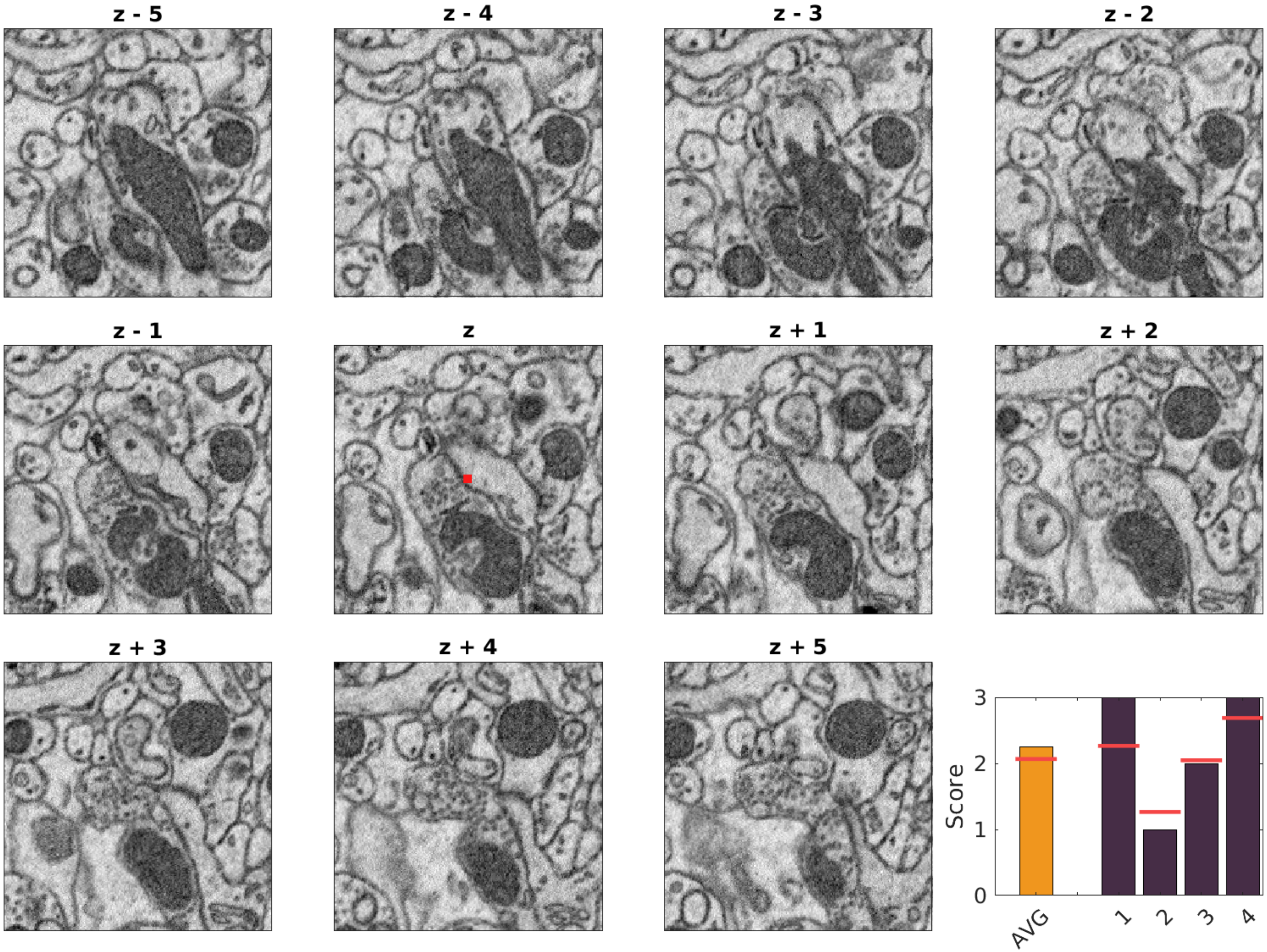

1288, 3244, 582

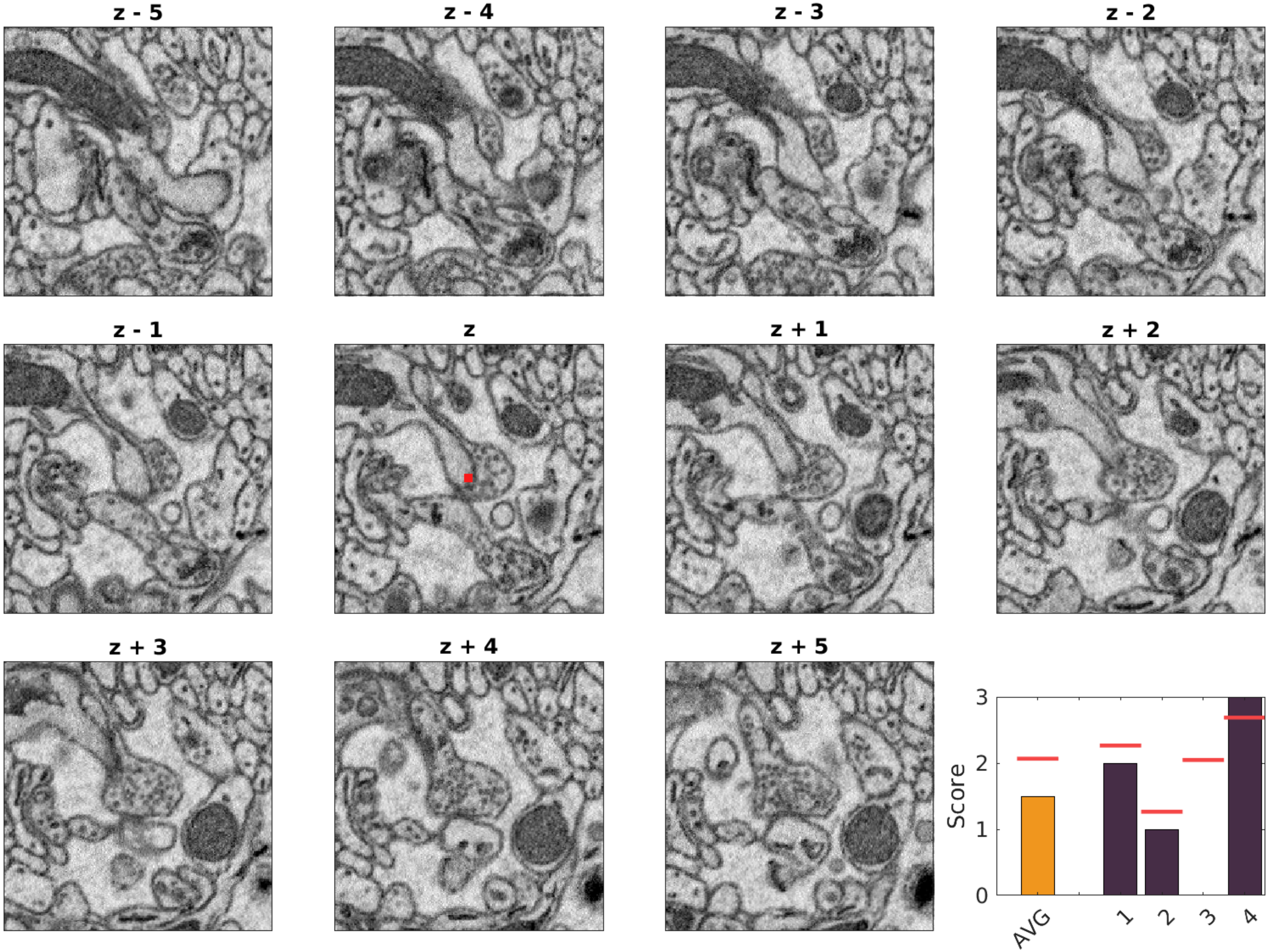

1330, 4423, 571

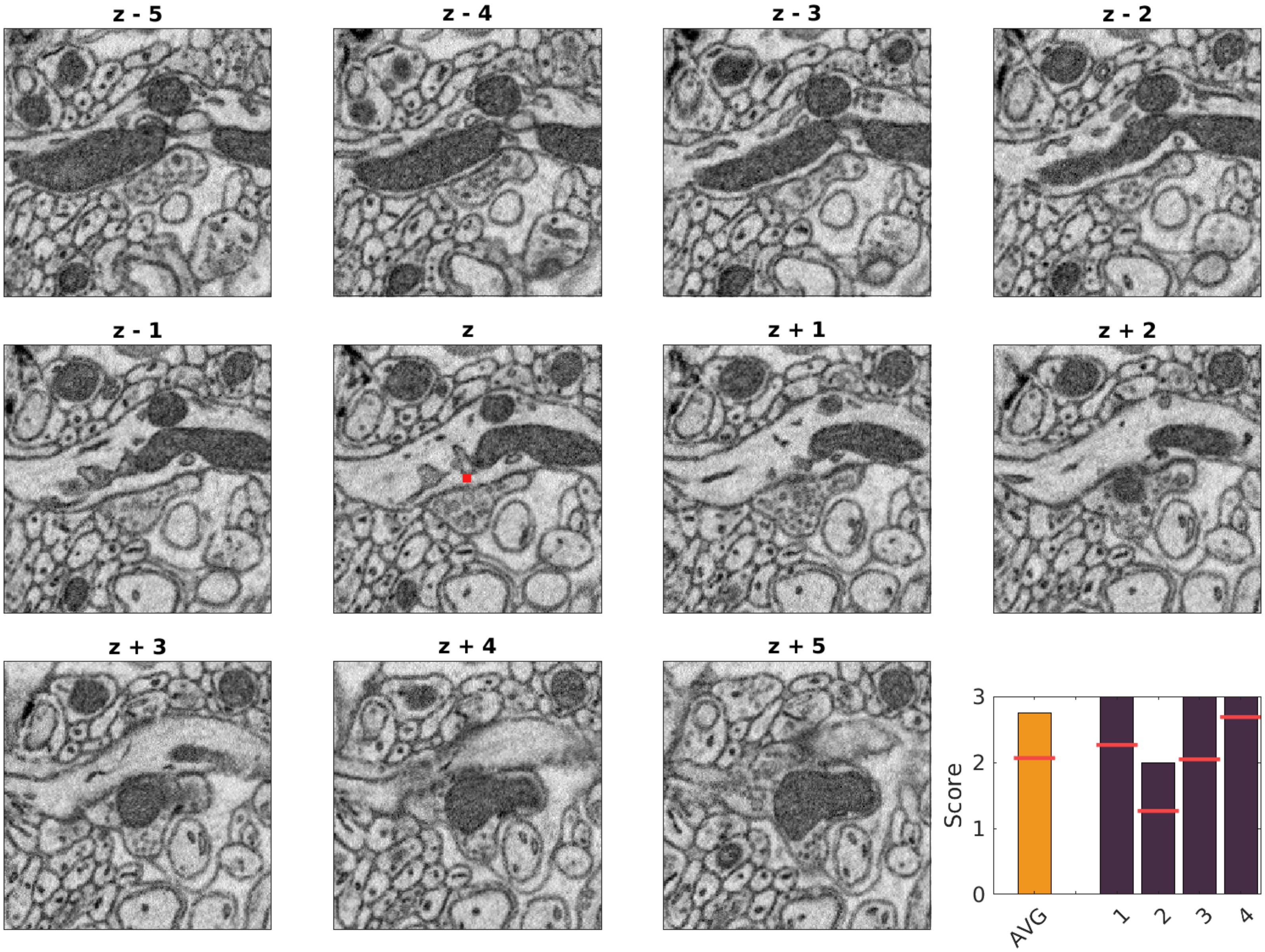

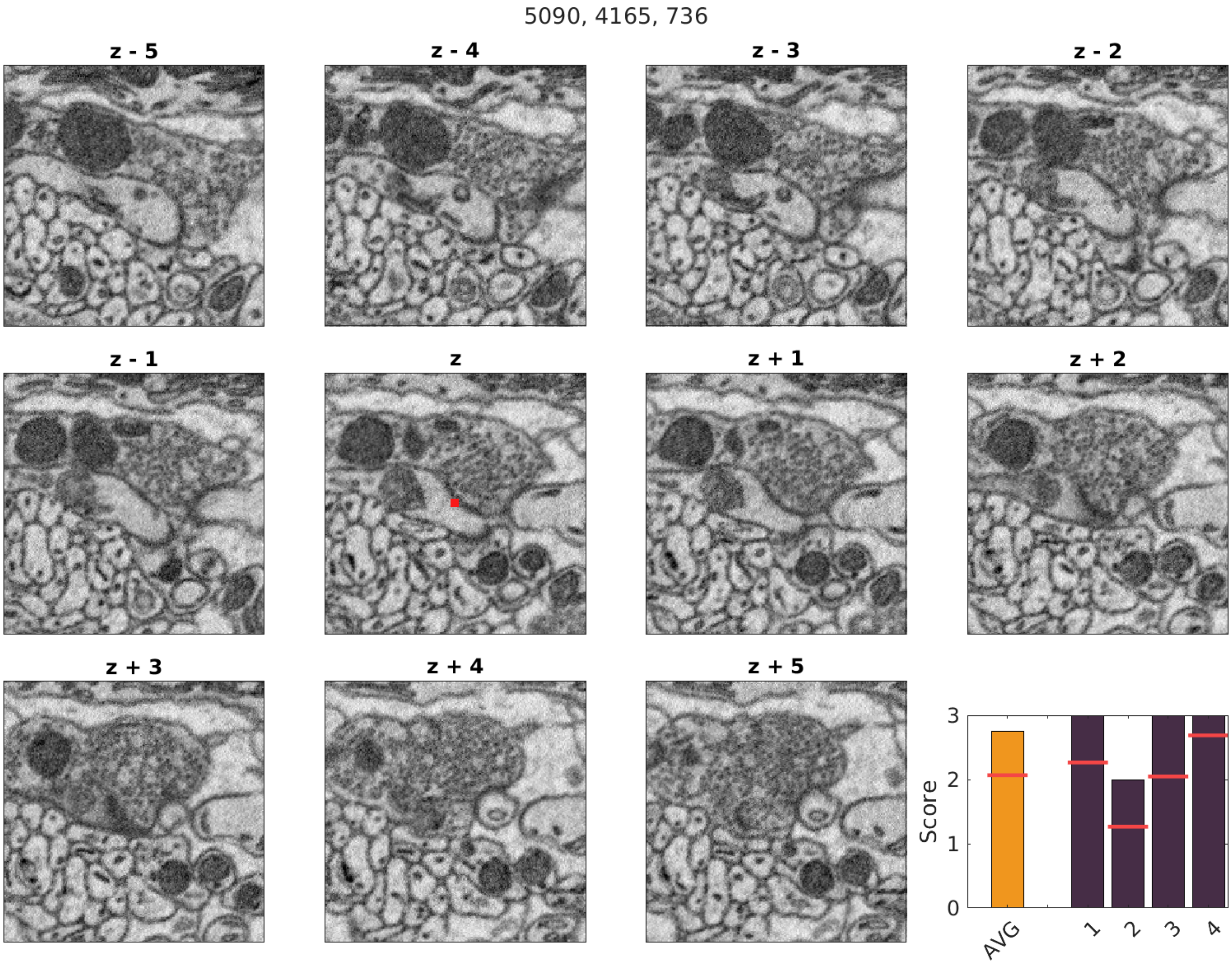

5090, 4165, 736
z - 5
z - 4
z - 3
z - 2
z - 1
z
z + 1
z + 2
z + 3
z + 4
z + 5
Score
3
2
1
0
AVG
1
2
3
4

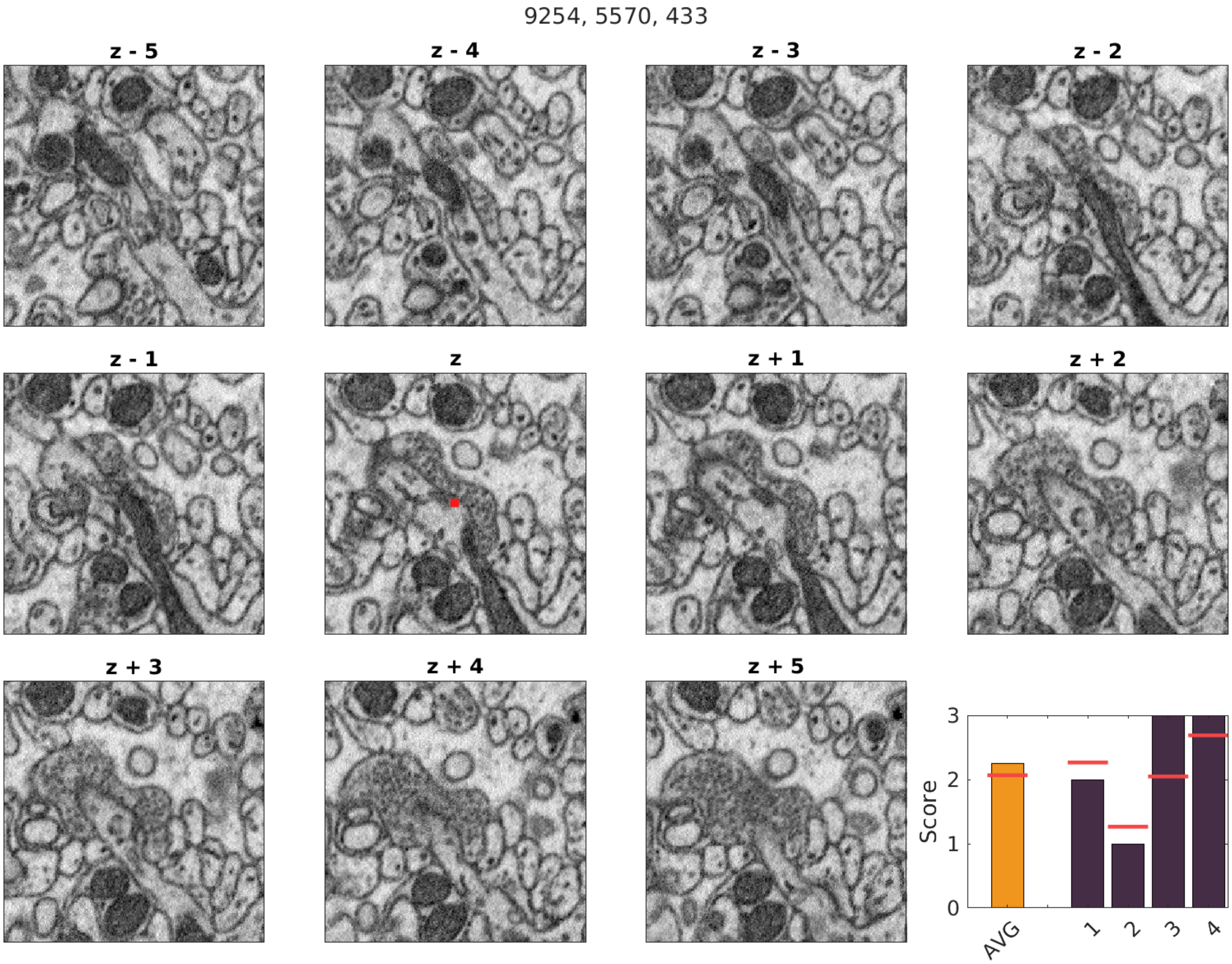
9254, 5570, 433
z - 5
z - 4
z - 3
z - 2
z - 1
z
z + 1
z + 2
z + 3
z + 4
z + 5
Score
0
1
2
3
AVG
1
2
3
4

10723, 7175, 454

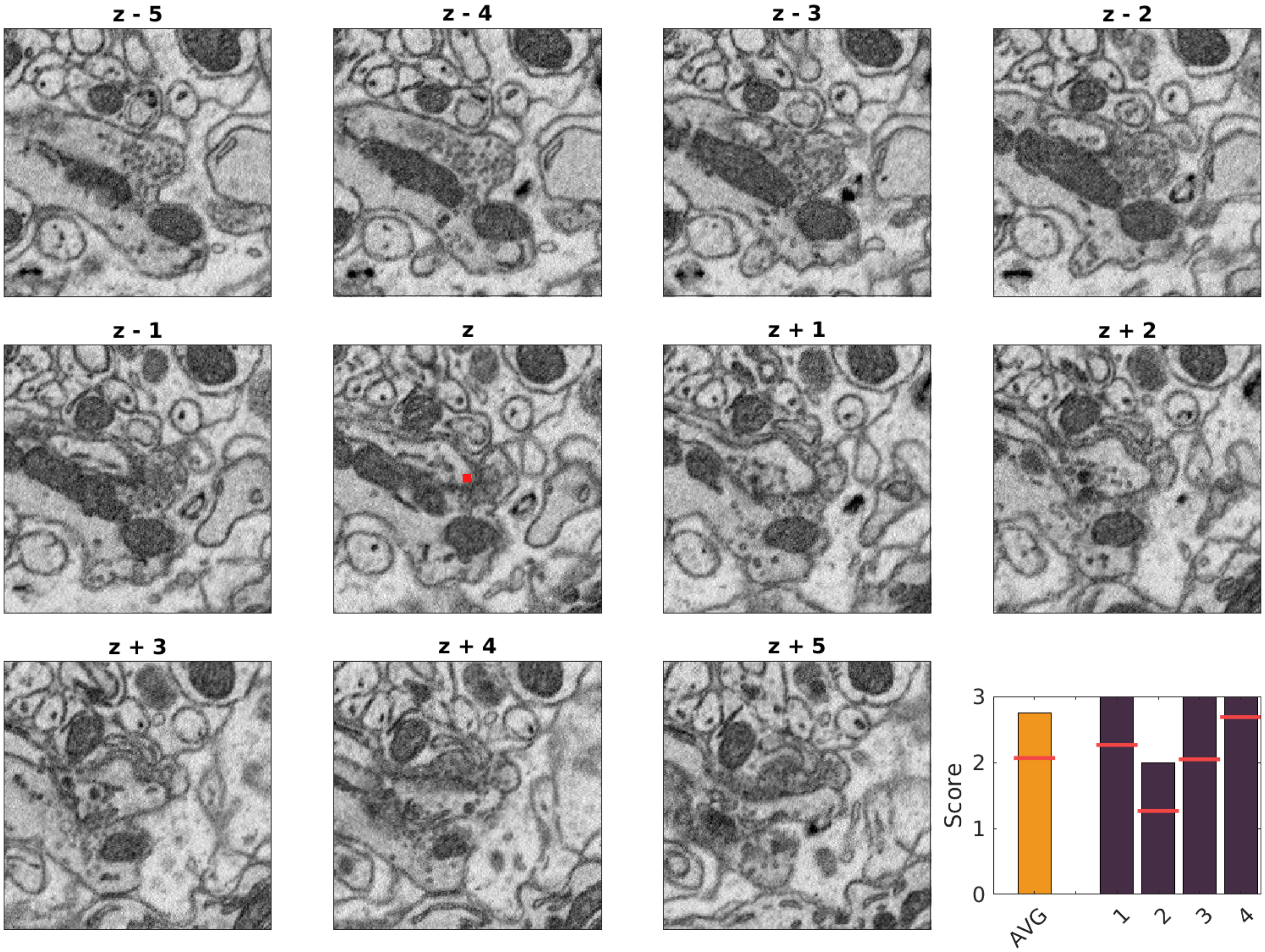

7601, 5587, 551

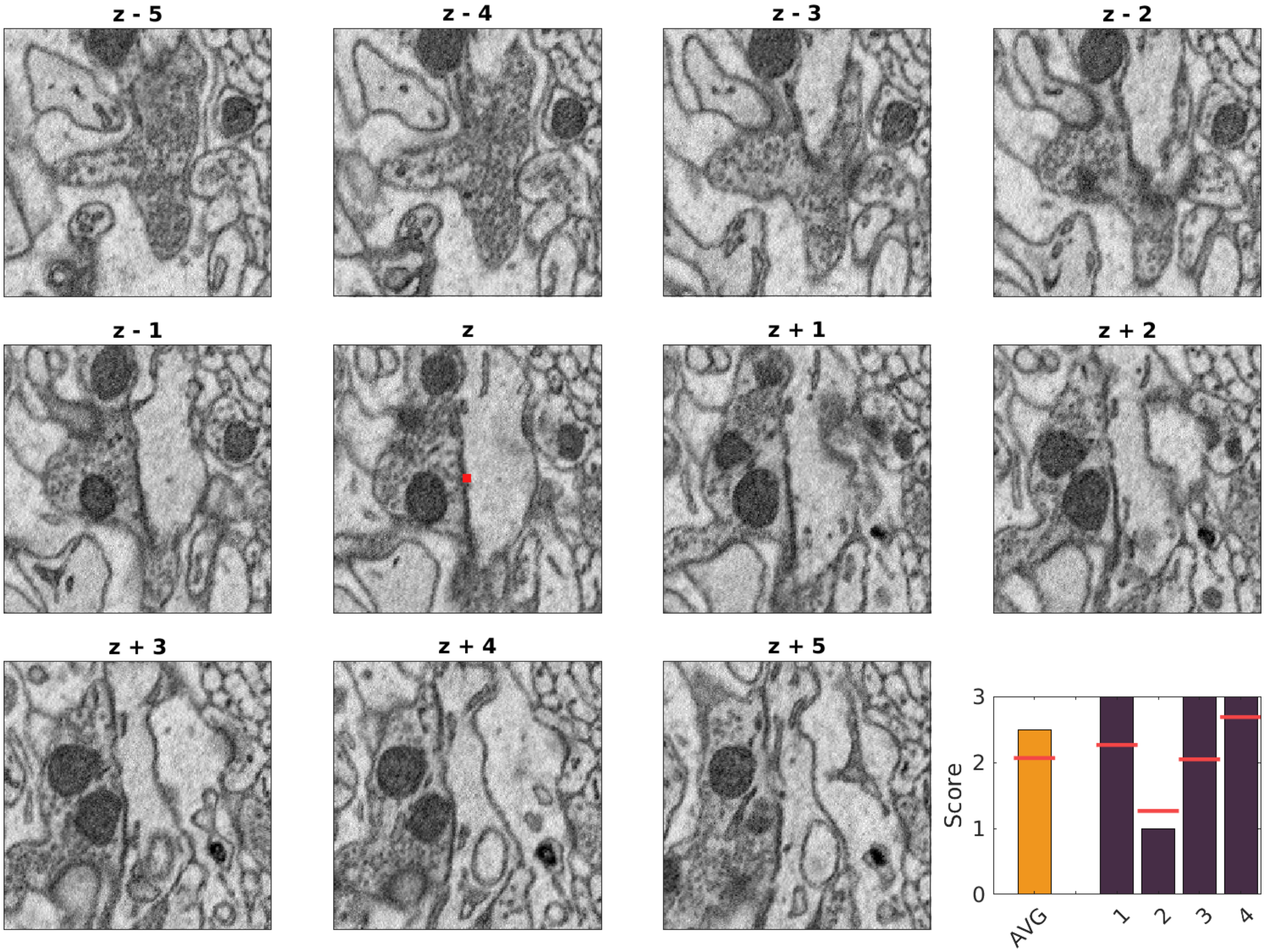

7404, 7108, 434

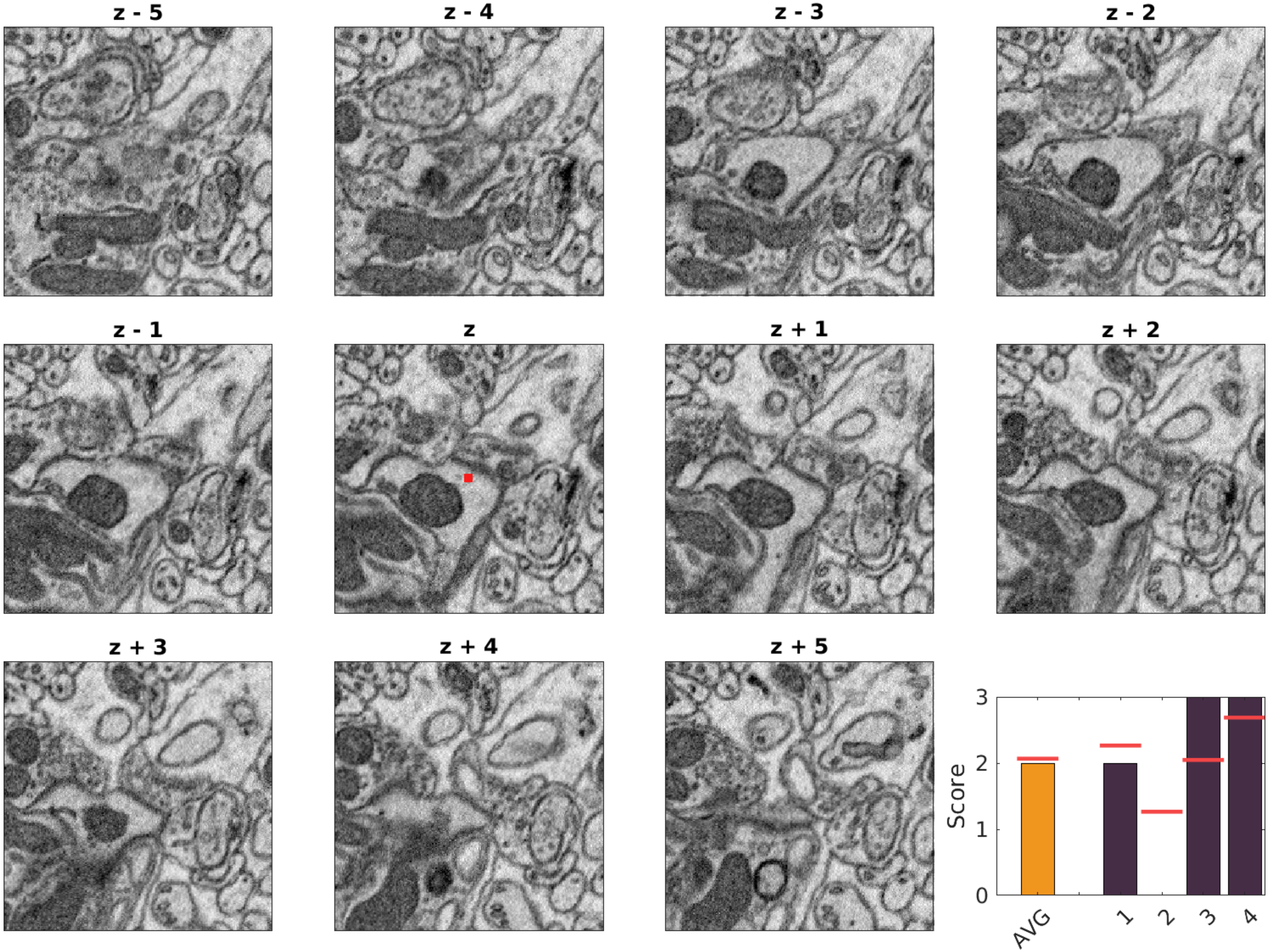

9785, 195, 982

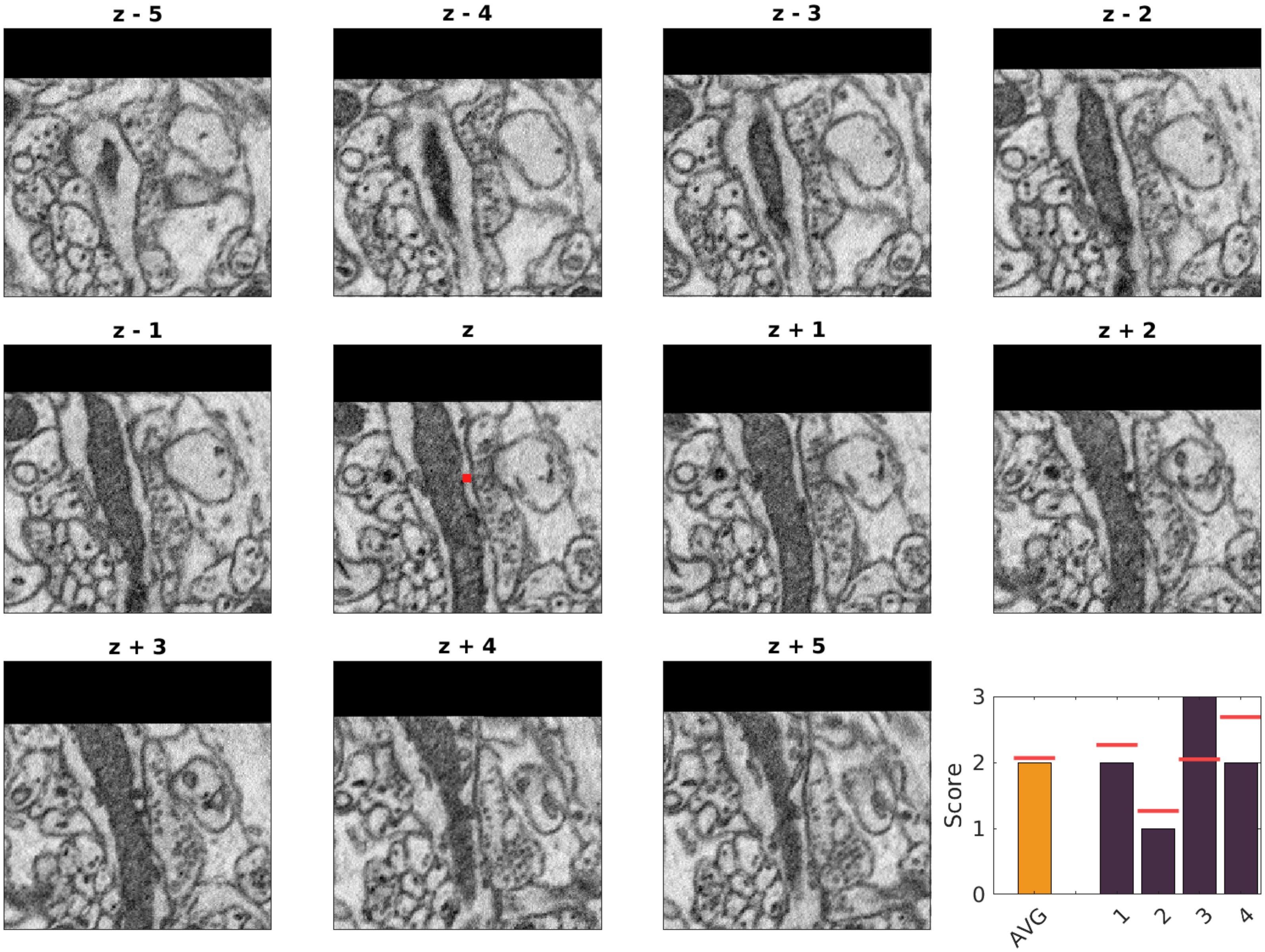

10277, 3013, 524

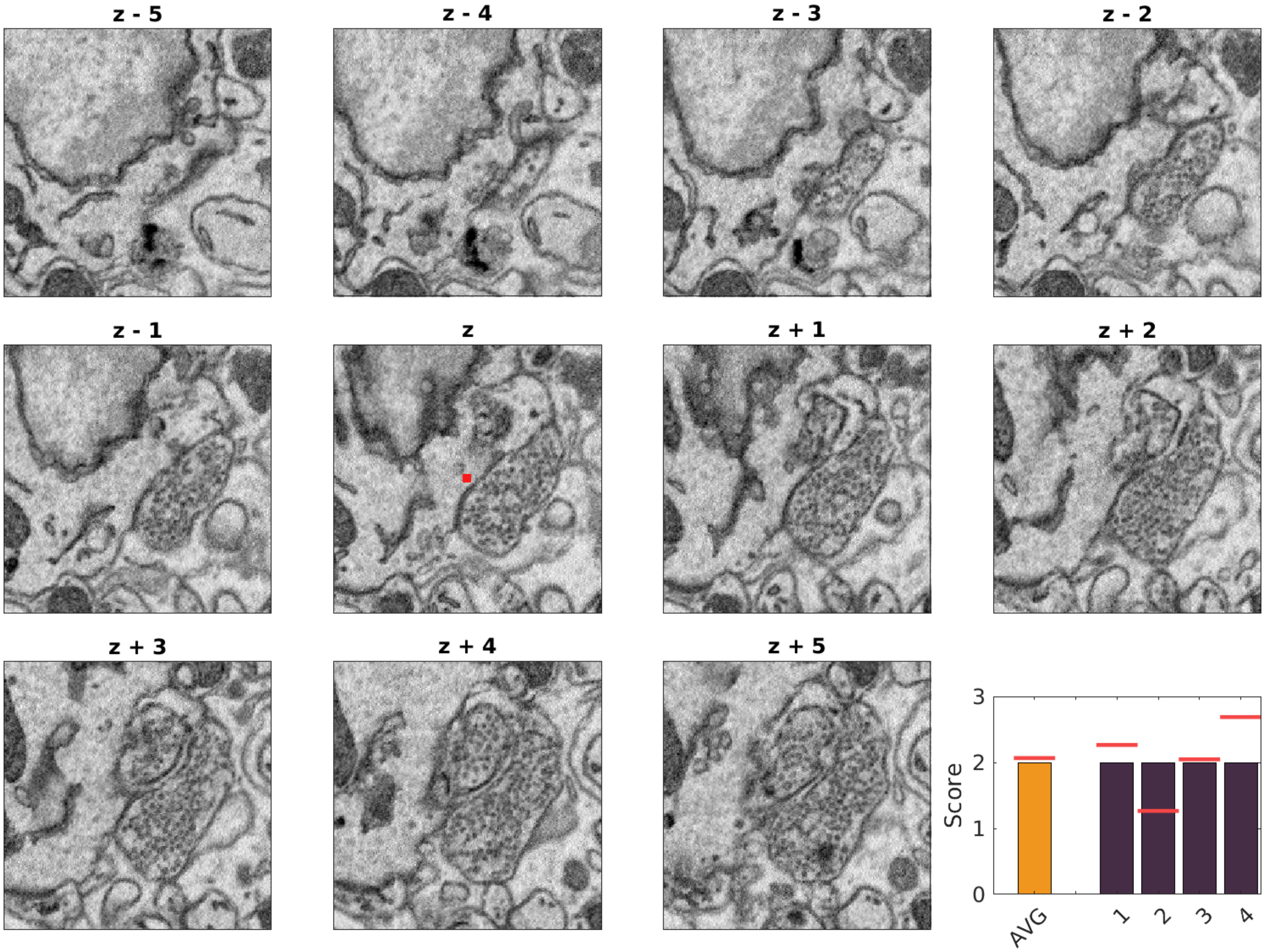

11804, 2417, 541

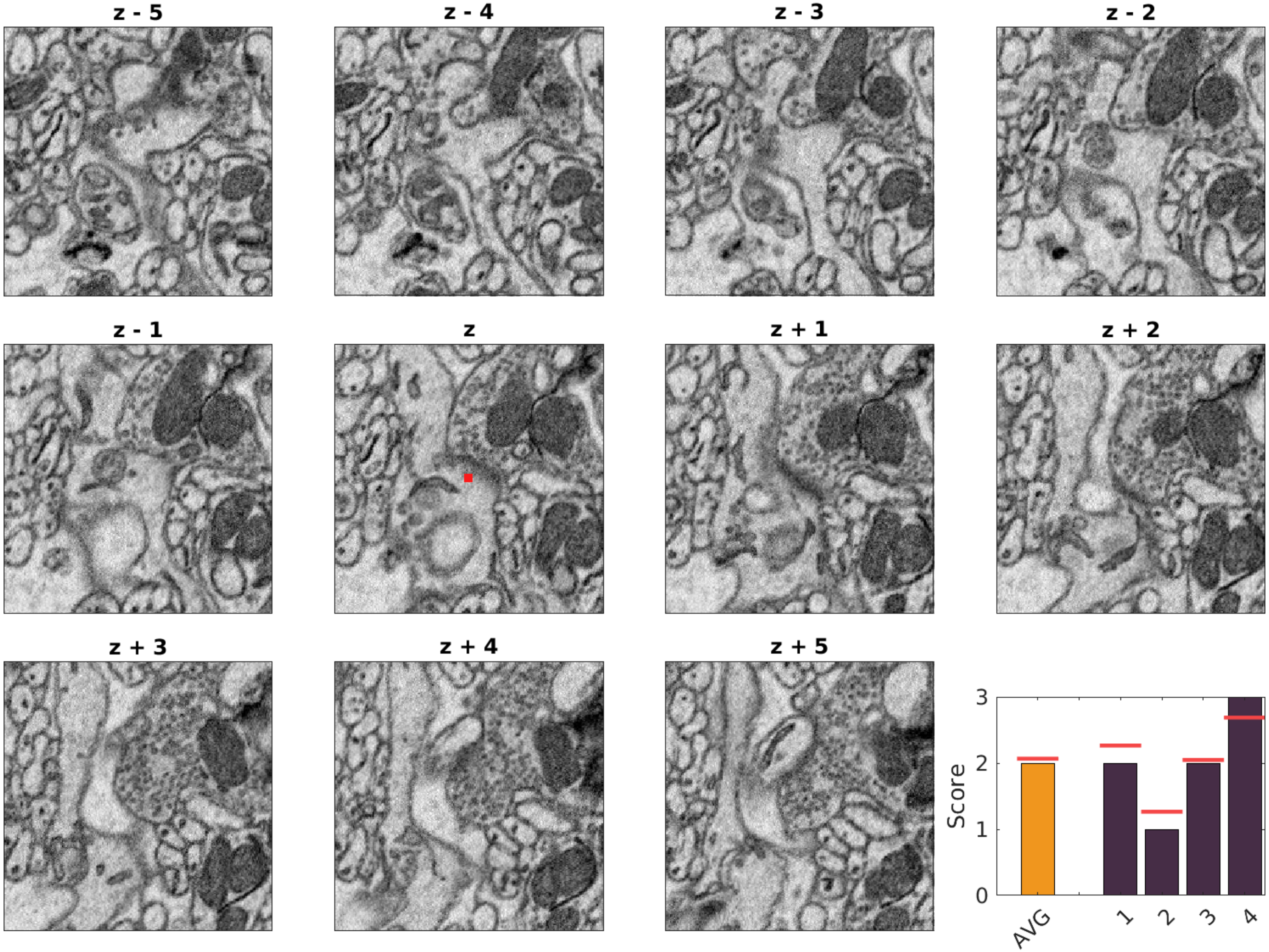

10568, 2631, 546

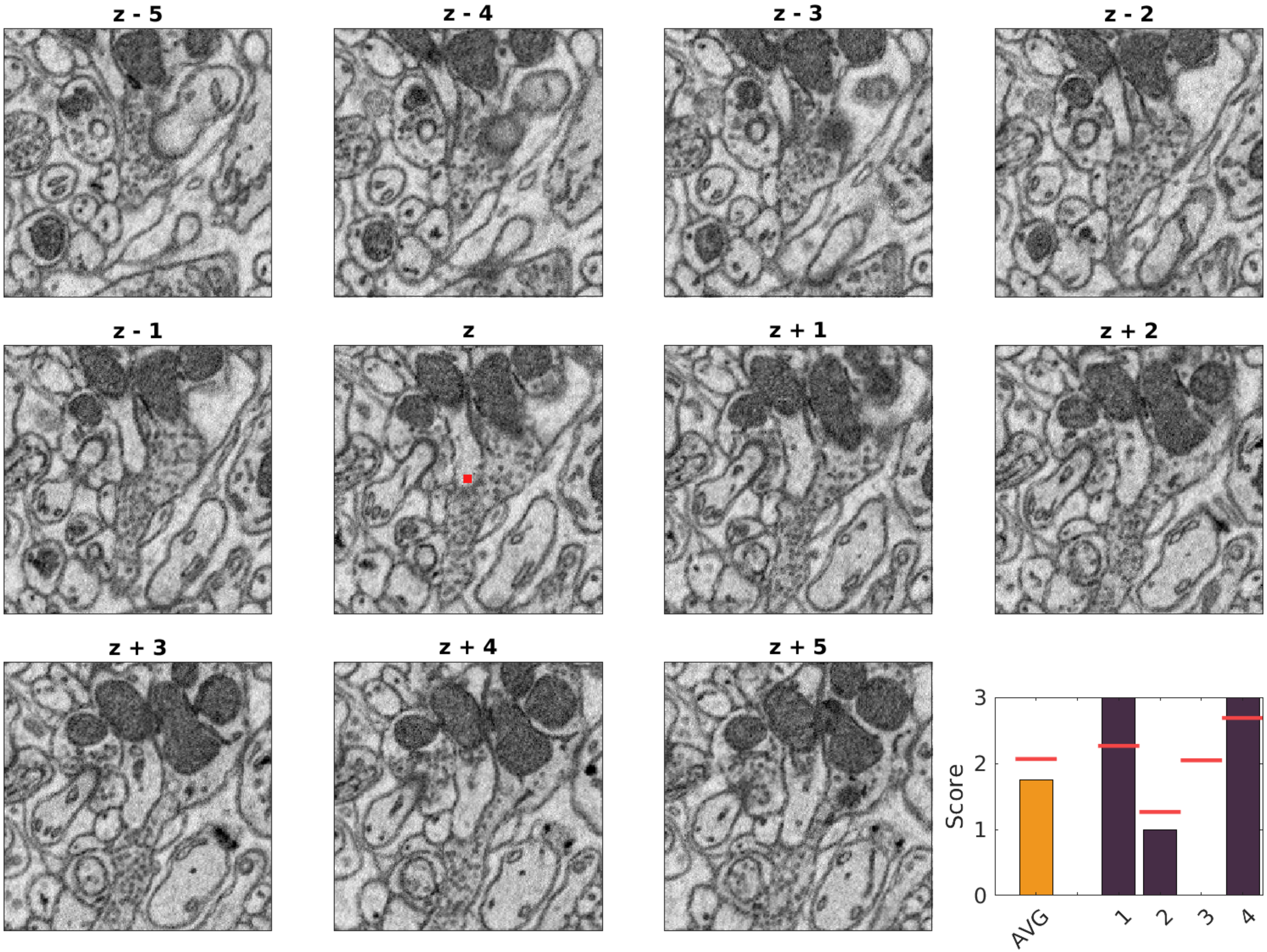

11981, 2725, 605

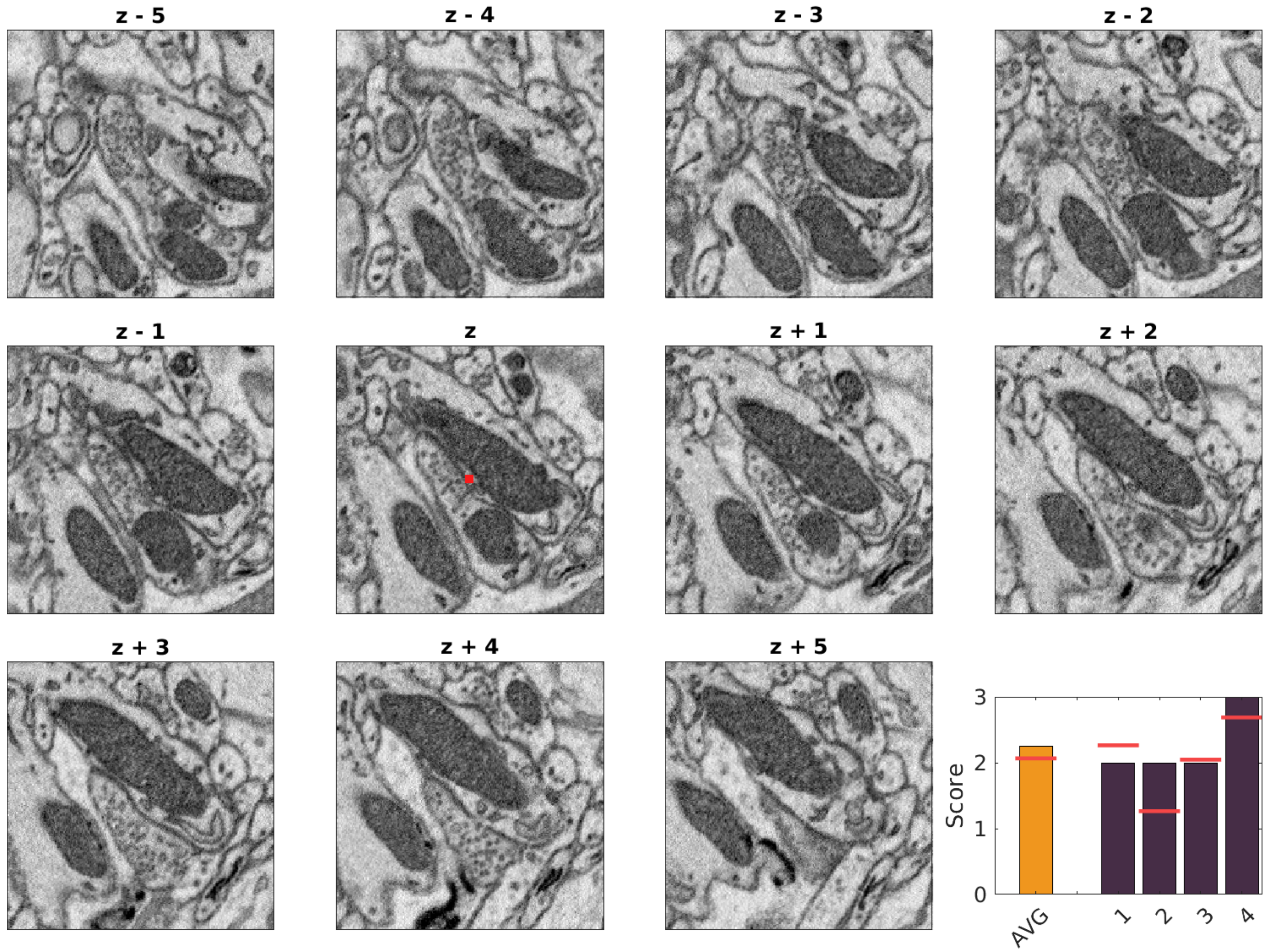

12872, 2530, 657

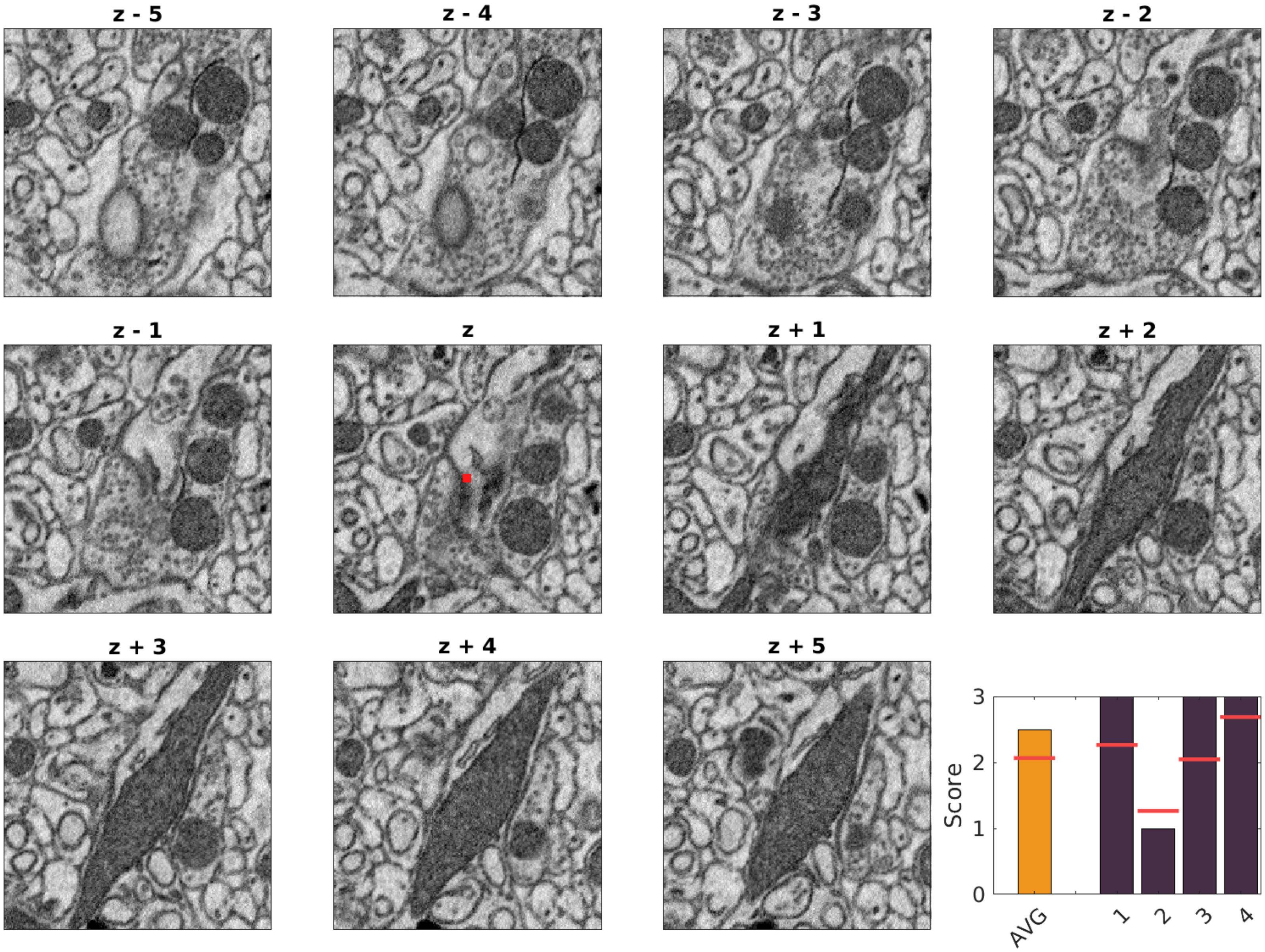

3454, 2057, 473

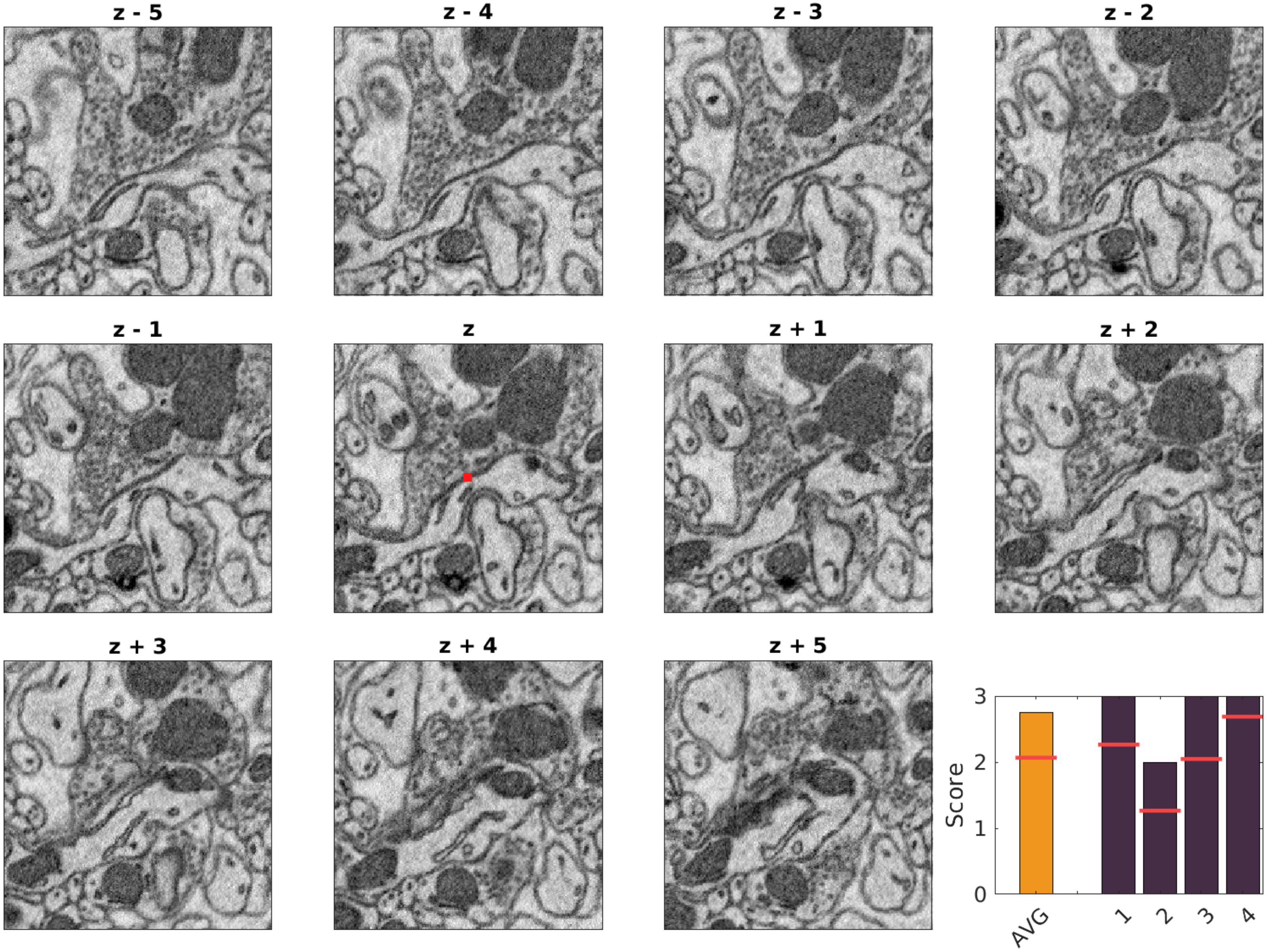

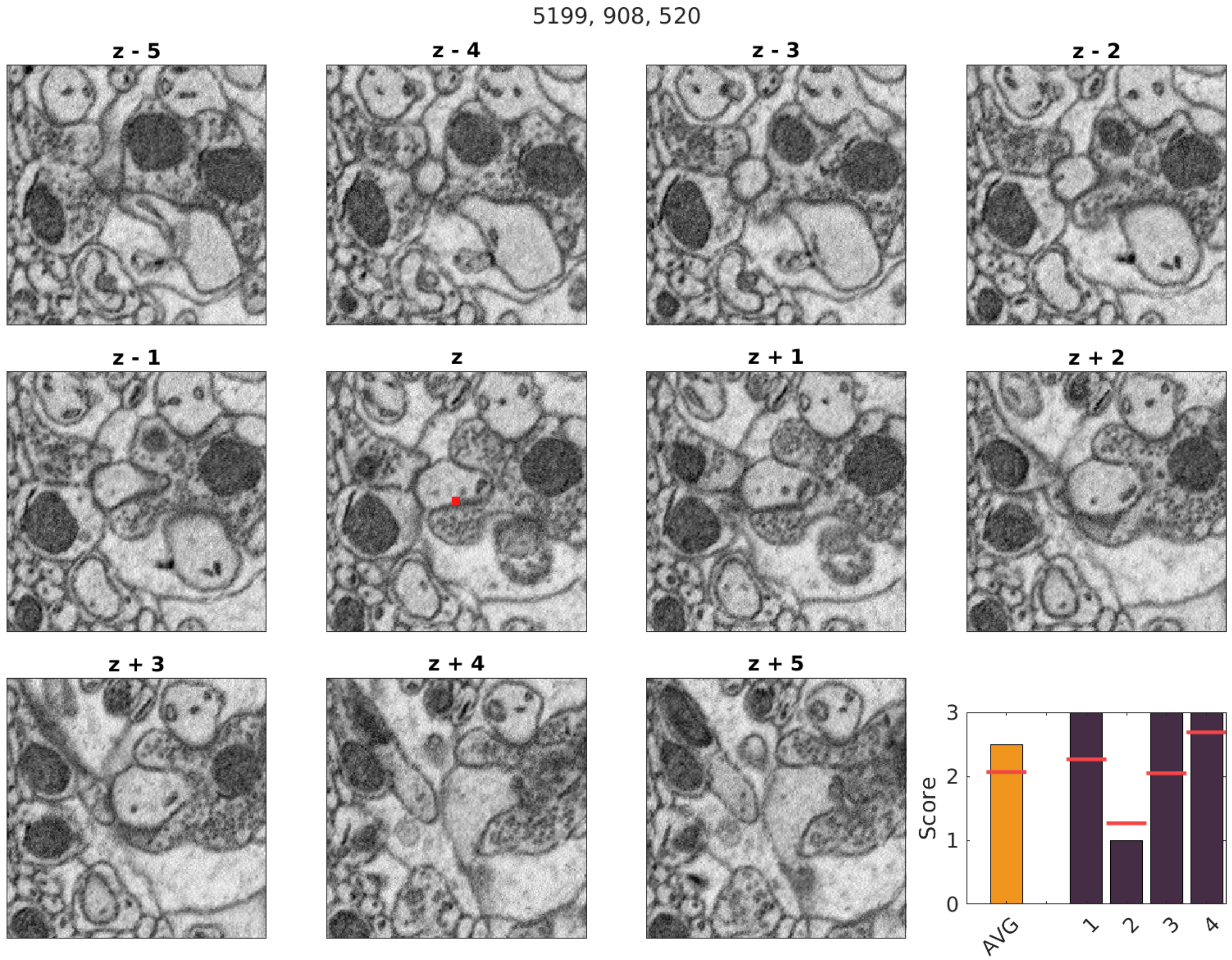

5199, 908, 520
z - 5
z - 4
z - 3
z - 2
z - 1
z
z + 1
z + 2
z + 3
z + 4
z + 5
Score
0
1
2
3
AVG
1
2
3
4

5090, 1430, 565

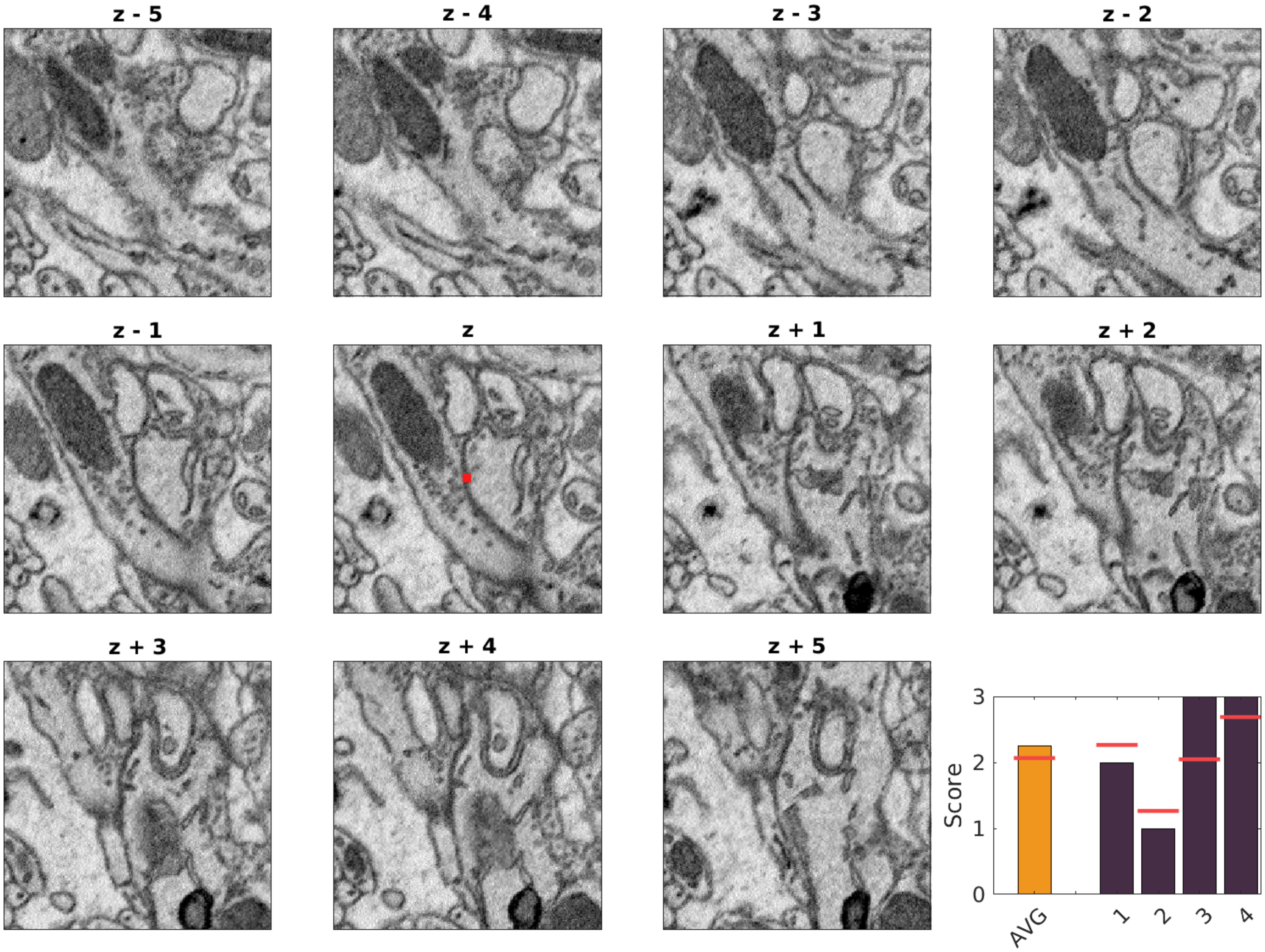

6136, 2164, 419

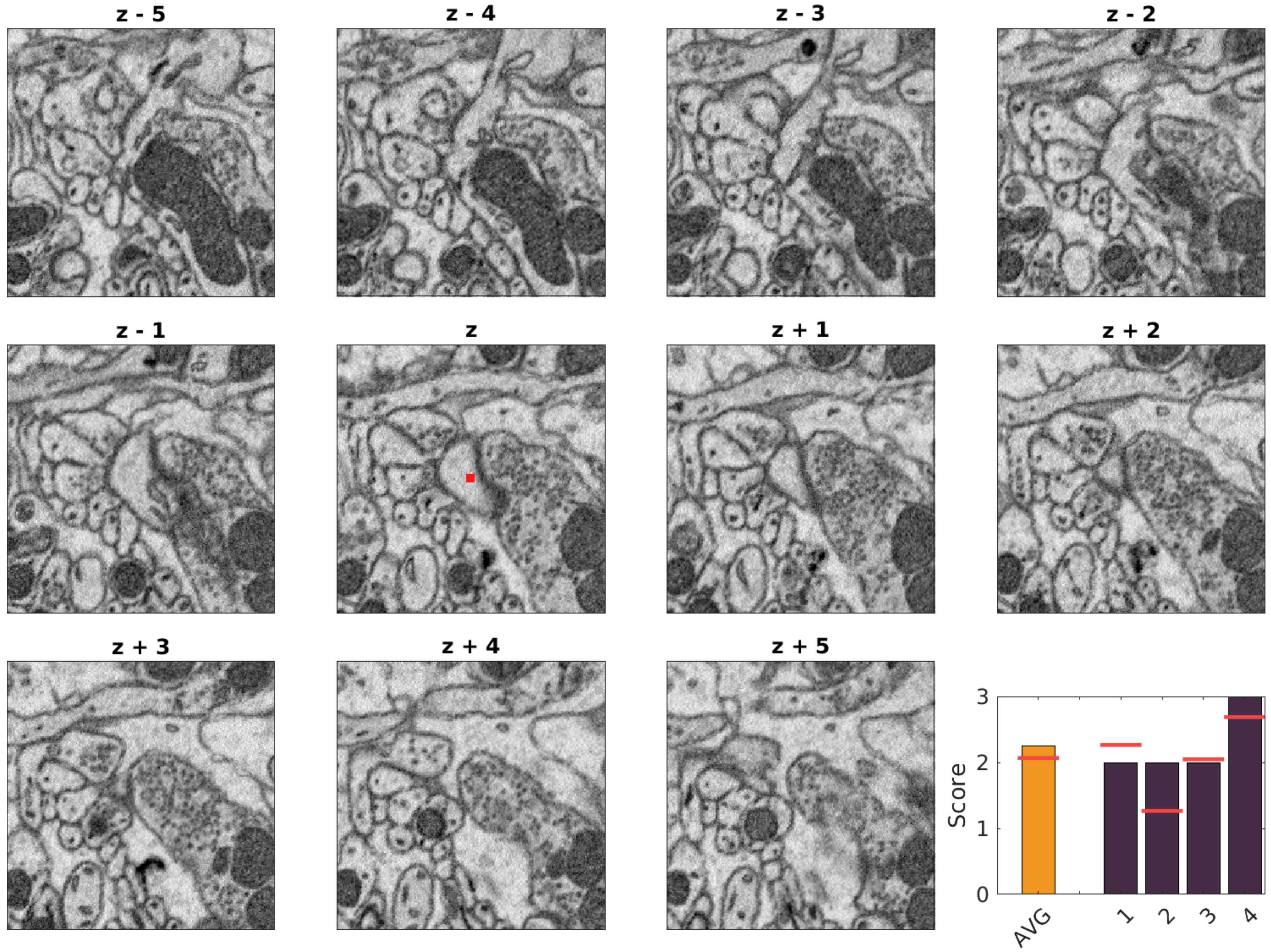

7609, 3364, 412

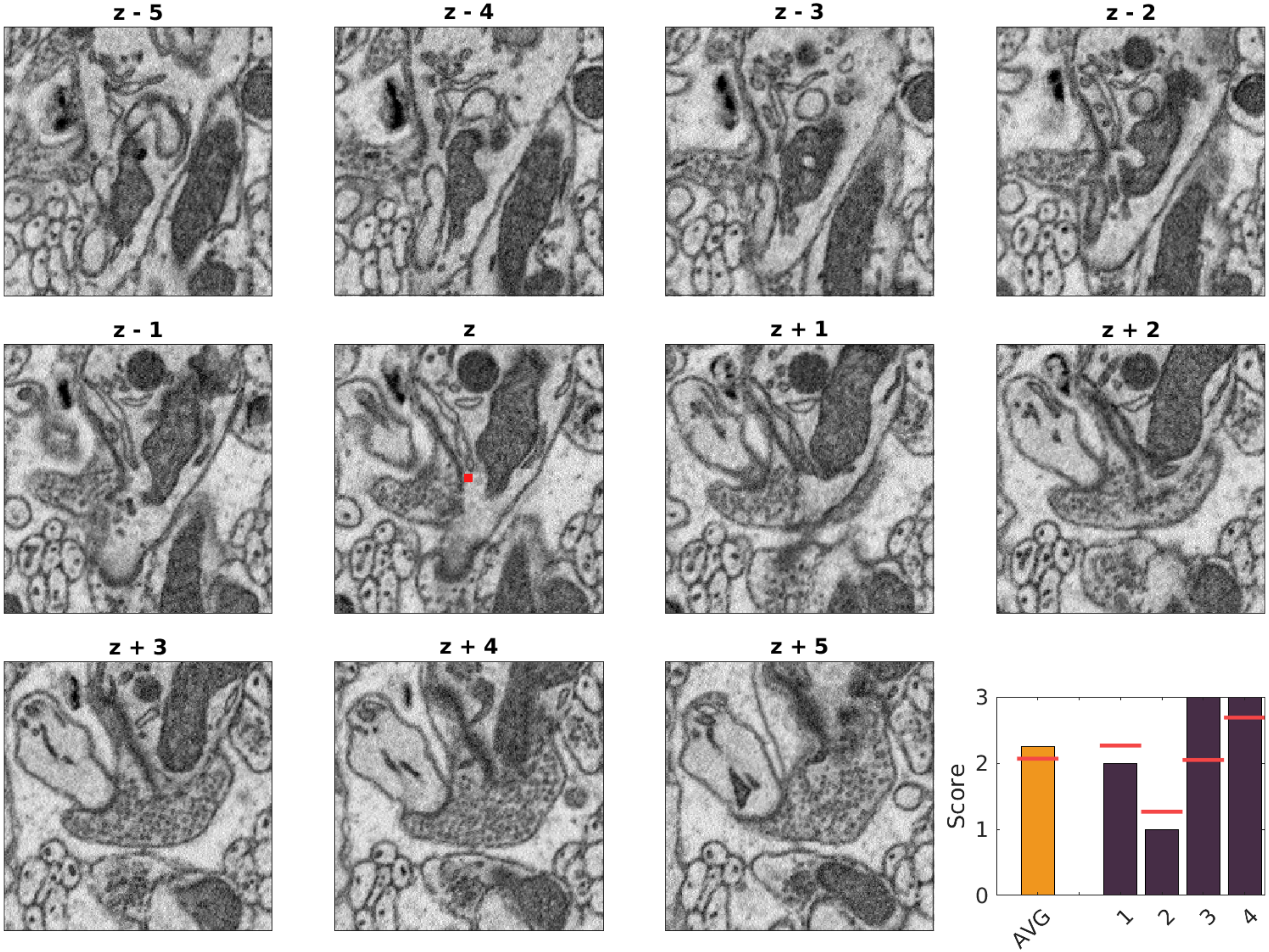

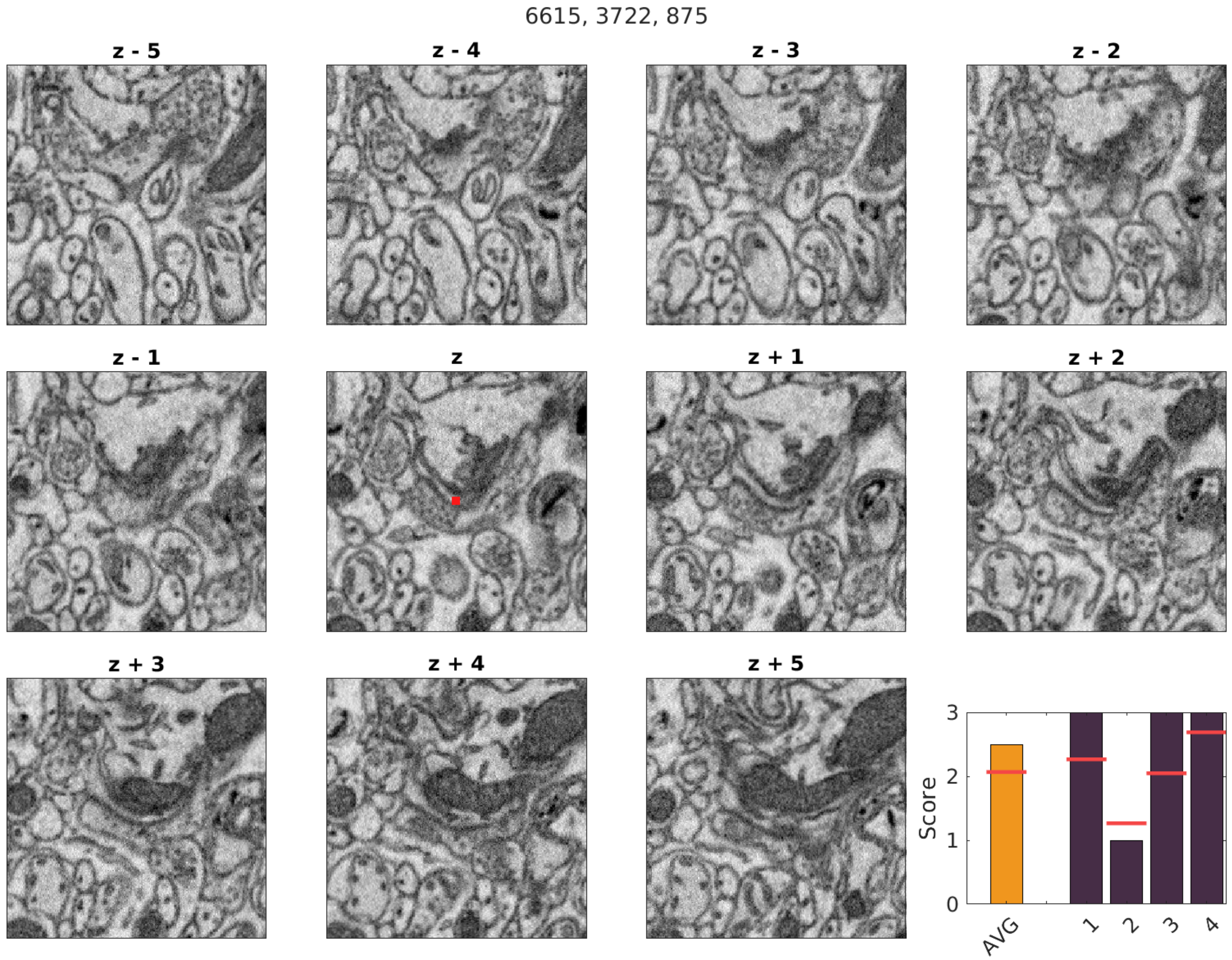
6615, 3722, 875
z - 5
z - 4
z - 3
z - 2
z - 1
z
z + 1
z + 2
z + 3
z + 4
z + 5
Score
0
1
2
3
AVG
1
2
3
4

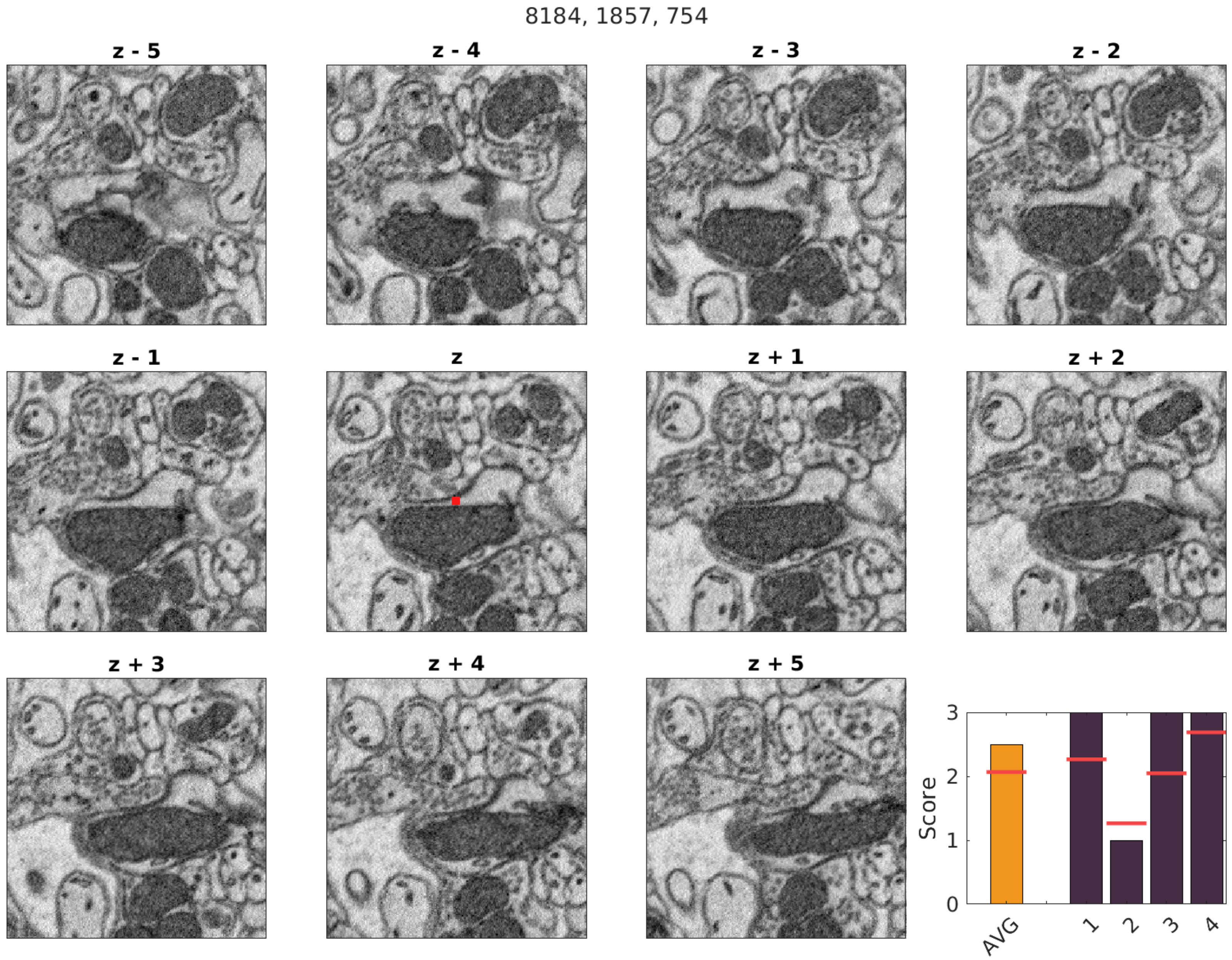
8184, 1857, 754
z - 5
z - 4
z - 3
z - 2
z - 1
z
z + 1
z + 2
z + 3
z + 4
z + 5
Score
0
1
2
3
AVG
1
2
3
4

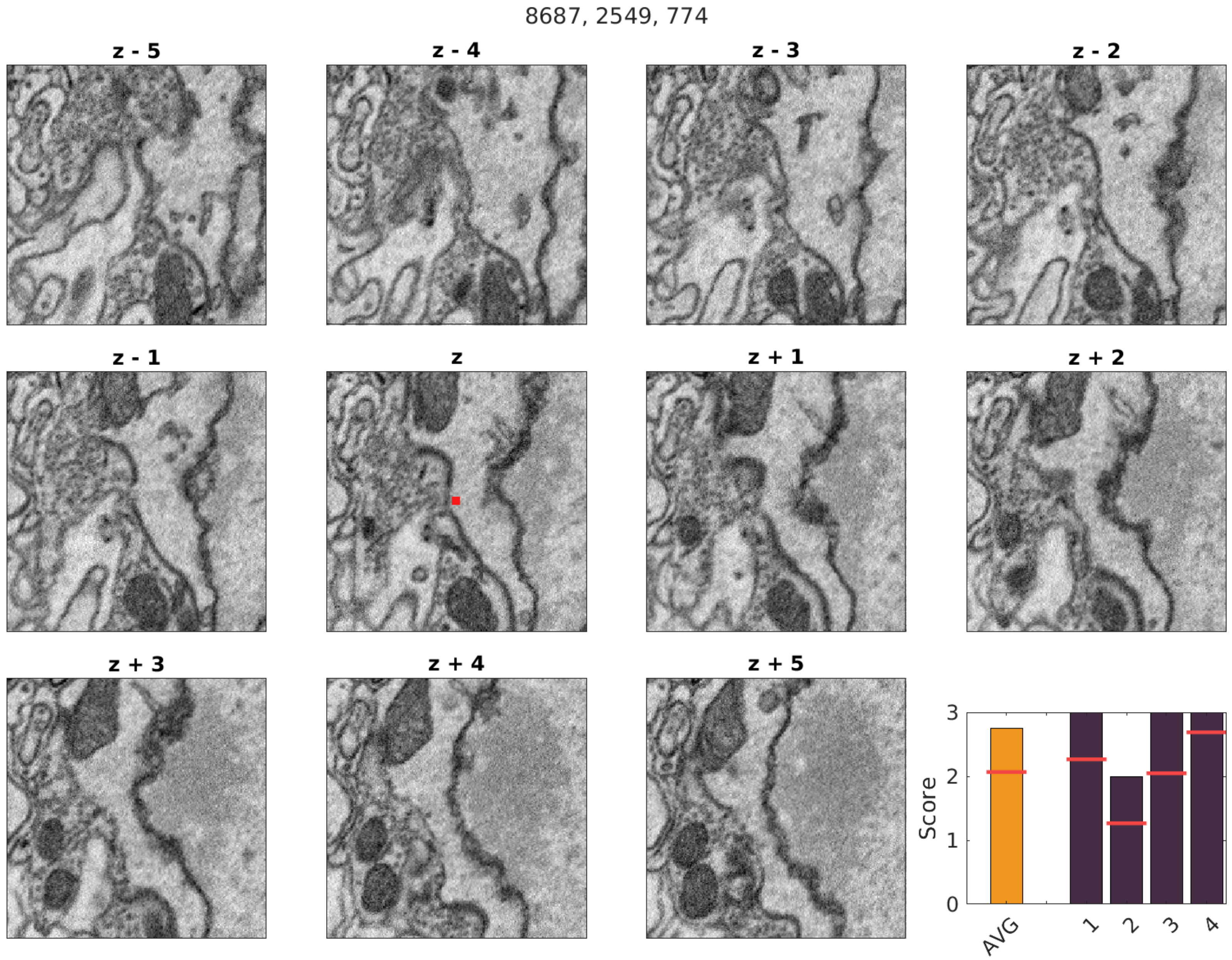
8687, 2549, 774
z - 5
z - 4
z - 3
z - 2
z - 1
z
z + 1
z + 2
z + 3
z + 4
z + 5
Score
0
1
2
3
AVG
1
2
3
4

3239, 1799, 363

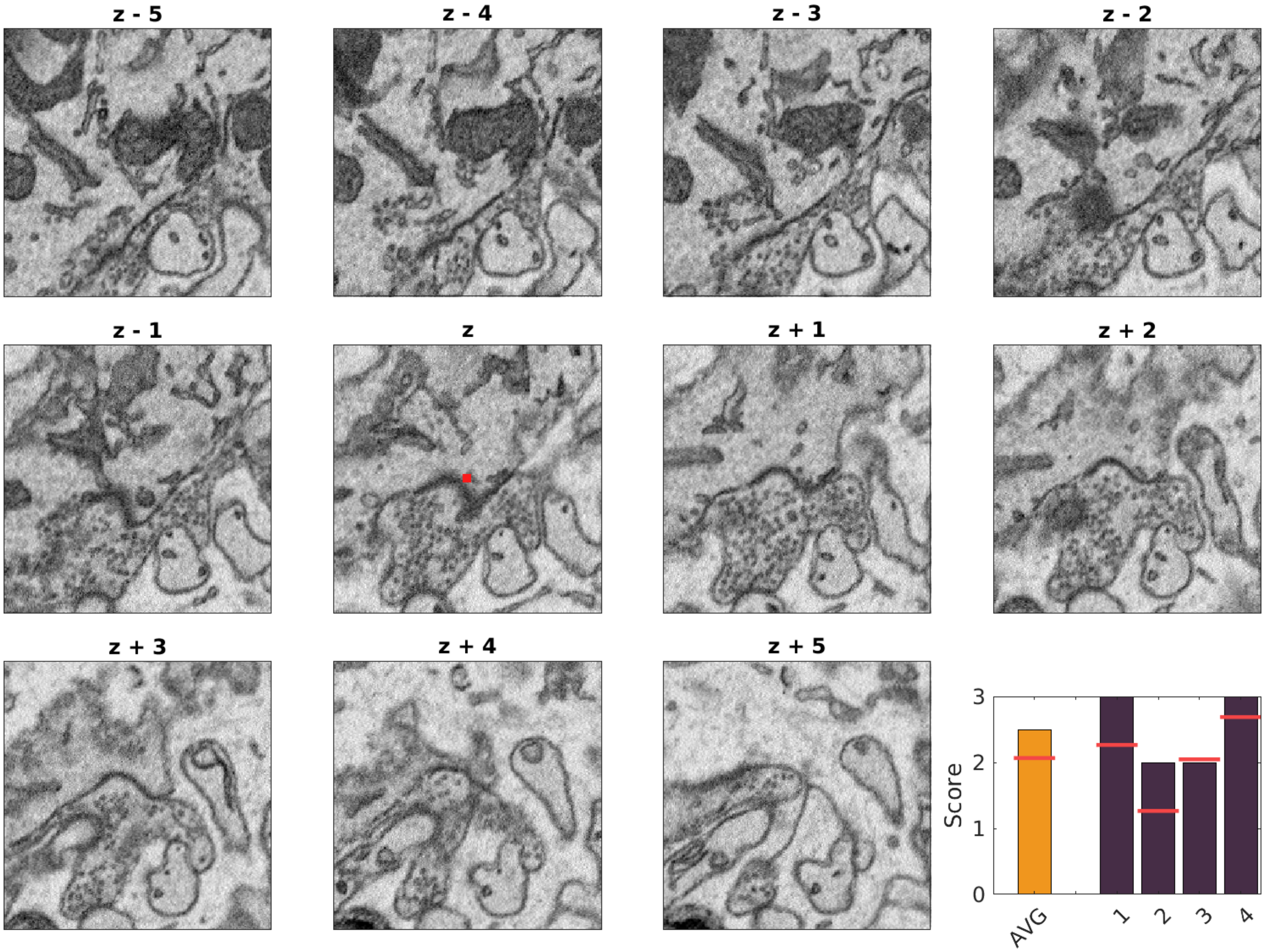

3862, 2002, 463

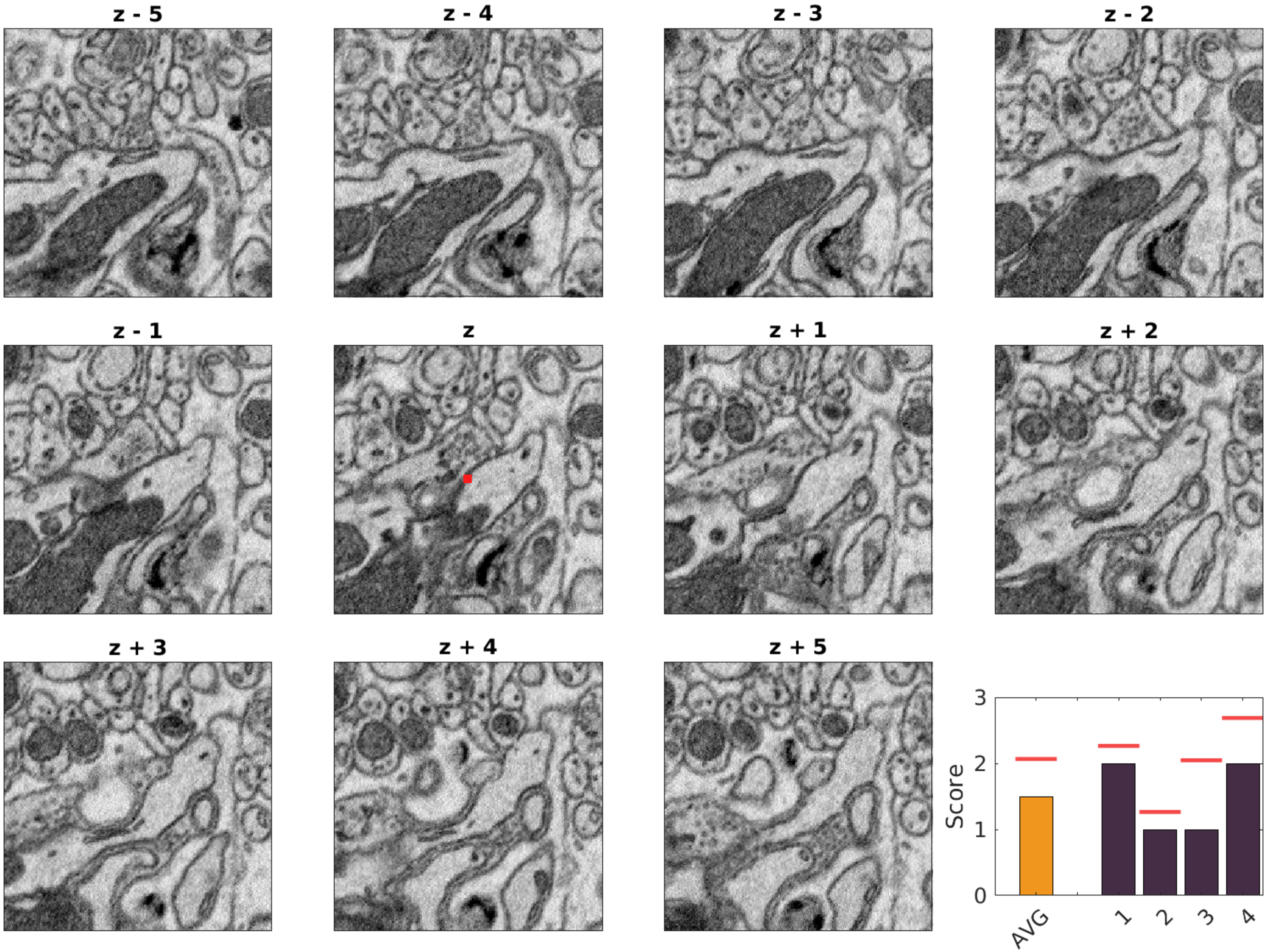

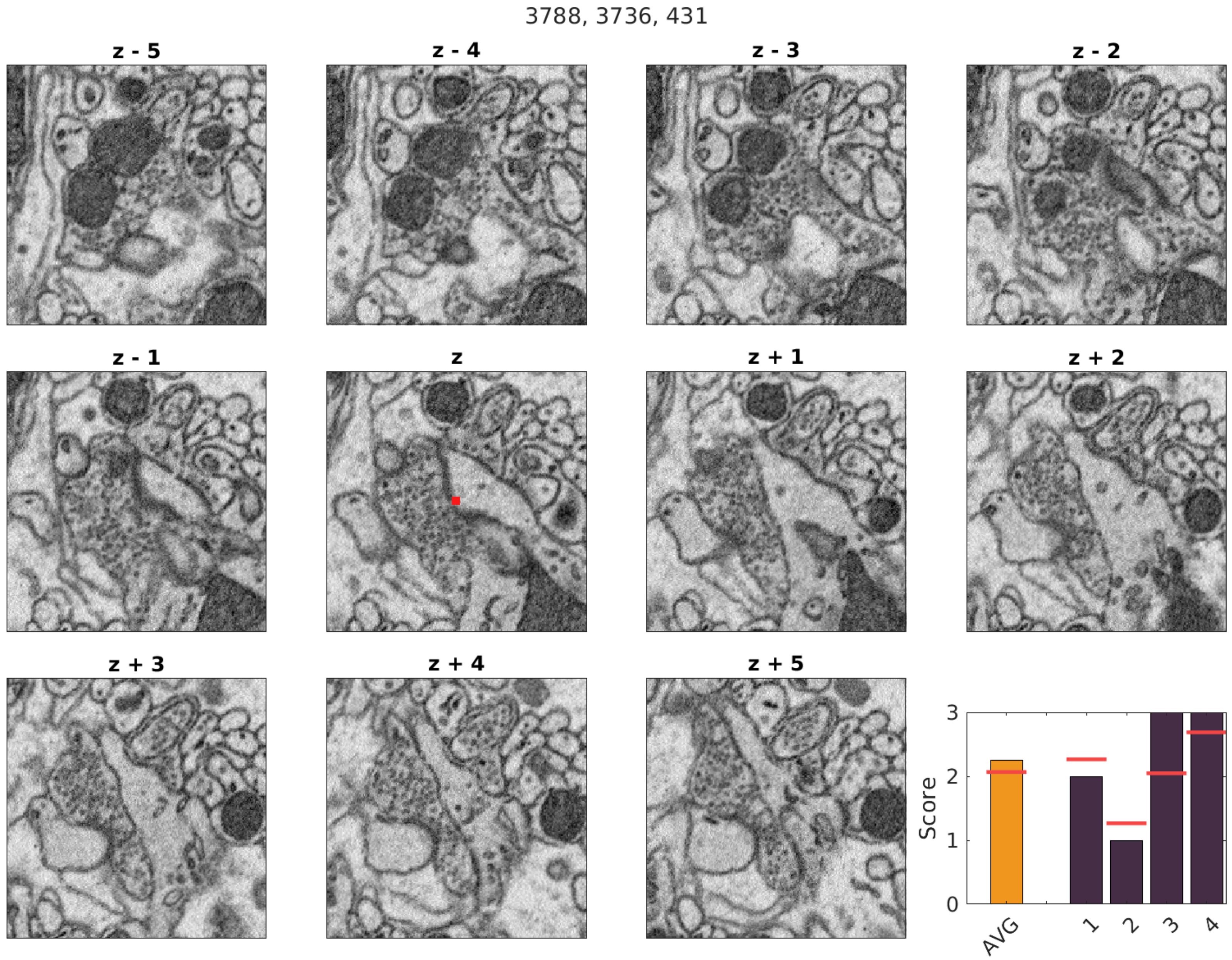
3788, 3736, 431
z - 5
z - 4
z - 3
z - 2
z - 1
z
z + 1
z + 2
z + 3
z + 4
z + 5
Score
0
1
2
3
AVG
1
2
3
4

3614, 7670, 295

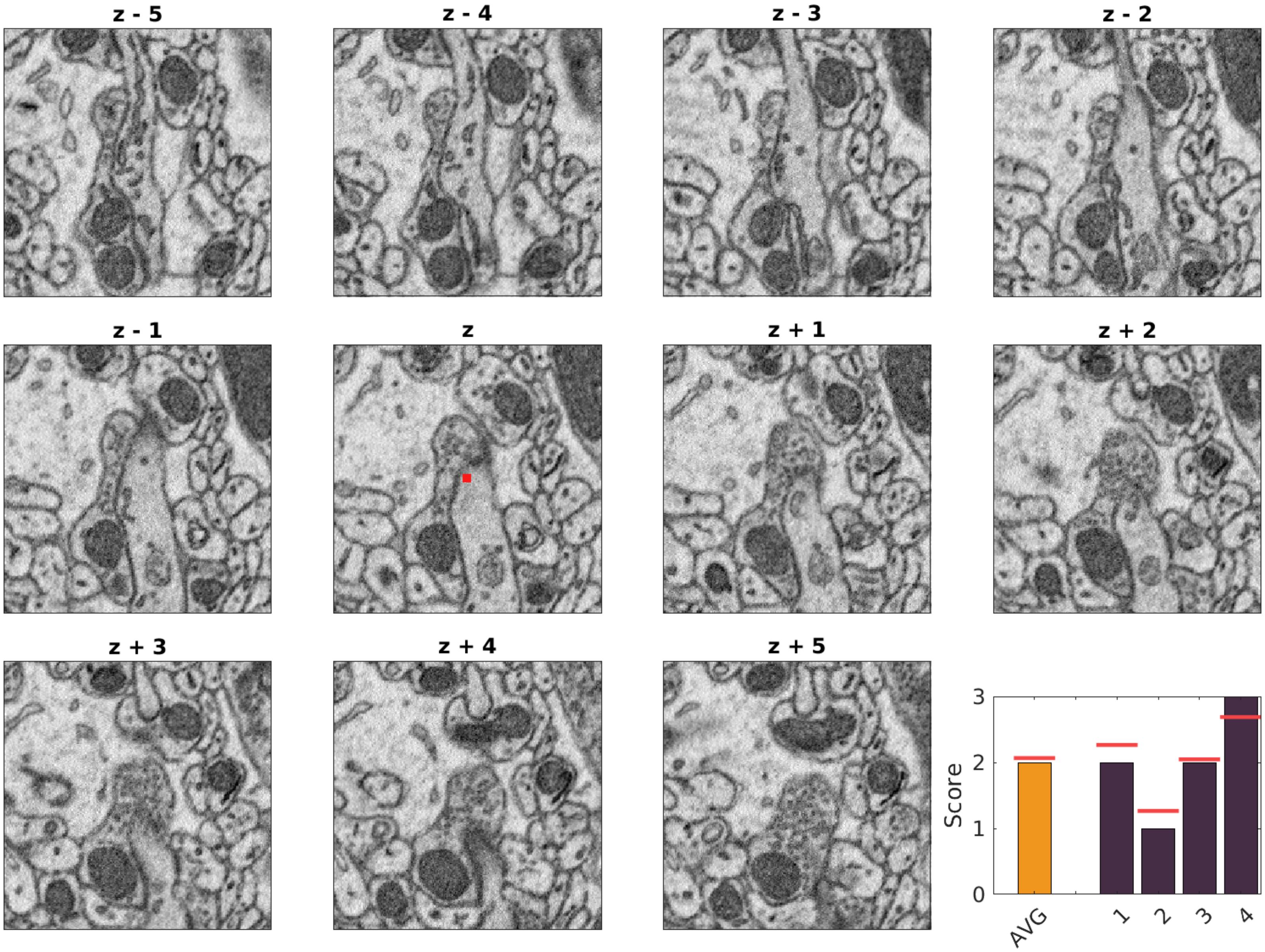

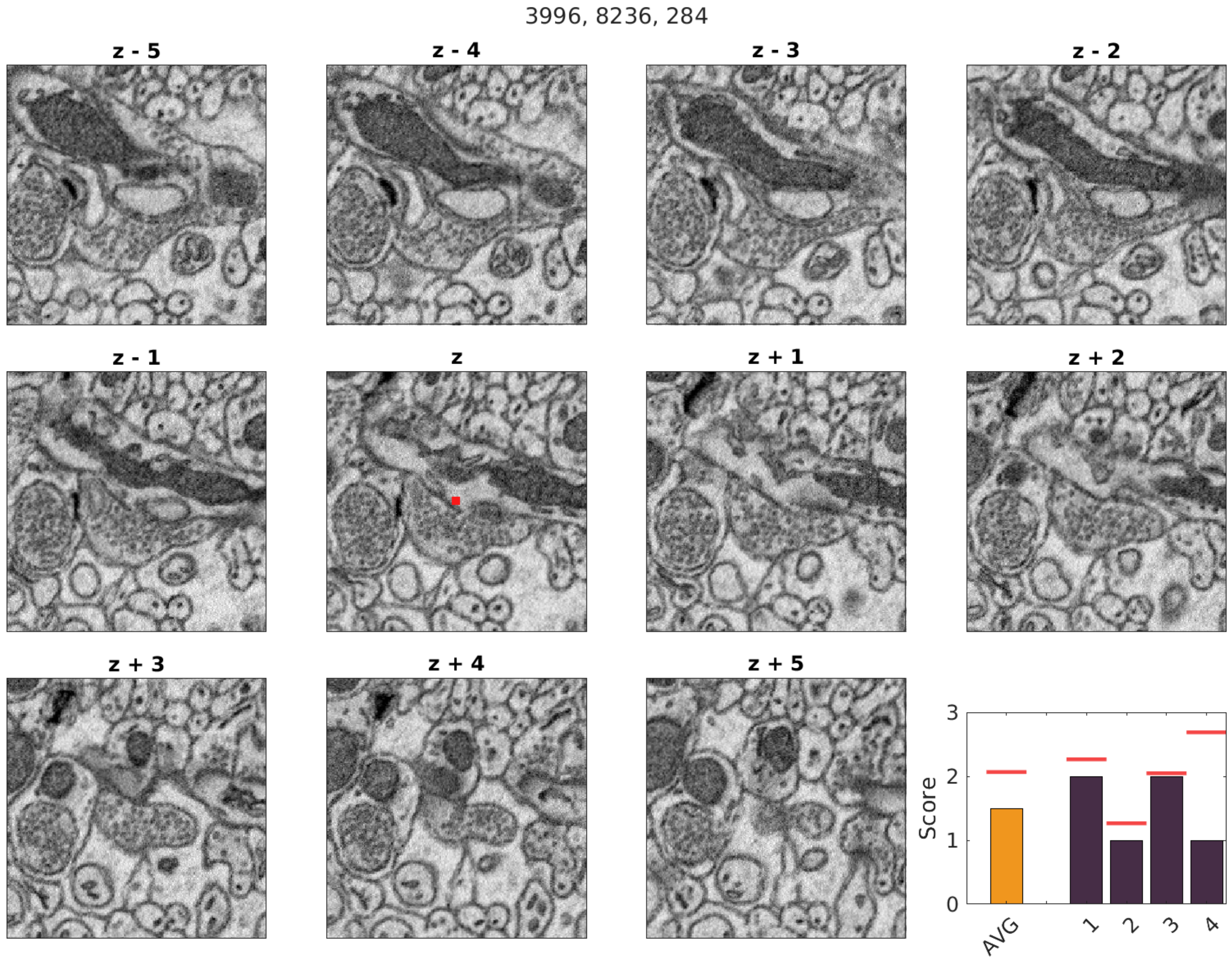
3996, 8236, 284
z - 5
z - 4
z - 3
z - 2
z - 1
z
z + 1
z + 2
z + 3
z + 4
z + 5
Score
0
1
2
3
AVG
1
2
3
4

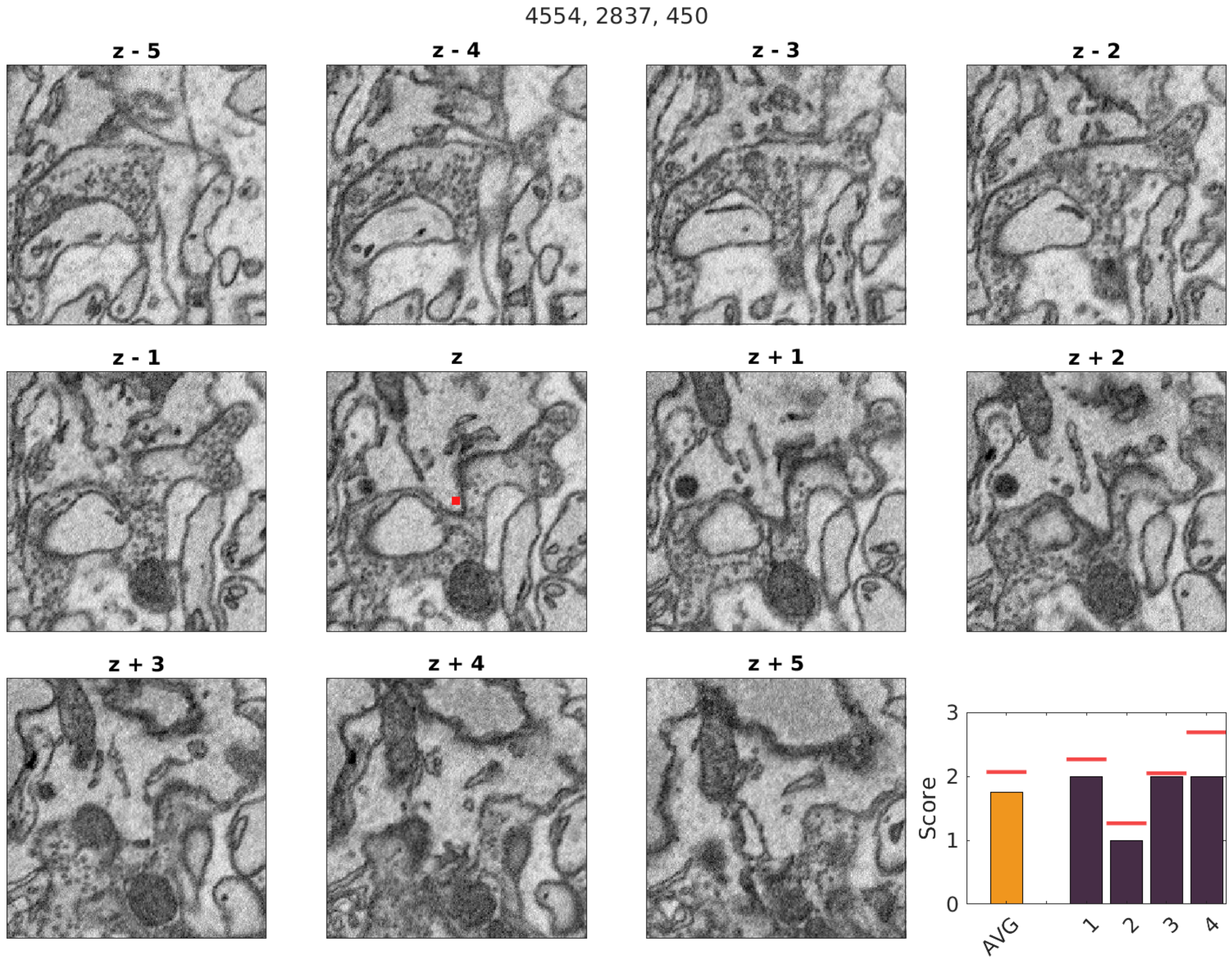

4554, 2837, 450
z - 5
z - 4
z - 3
z - 2
z - 1
z
z + 1
z + 2
z + 3
z + 4
z + 5
Score
0
1
2
3
AVG
1
2
3
4

4459, 5676, 326

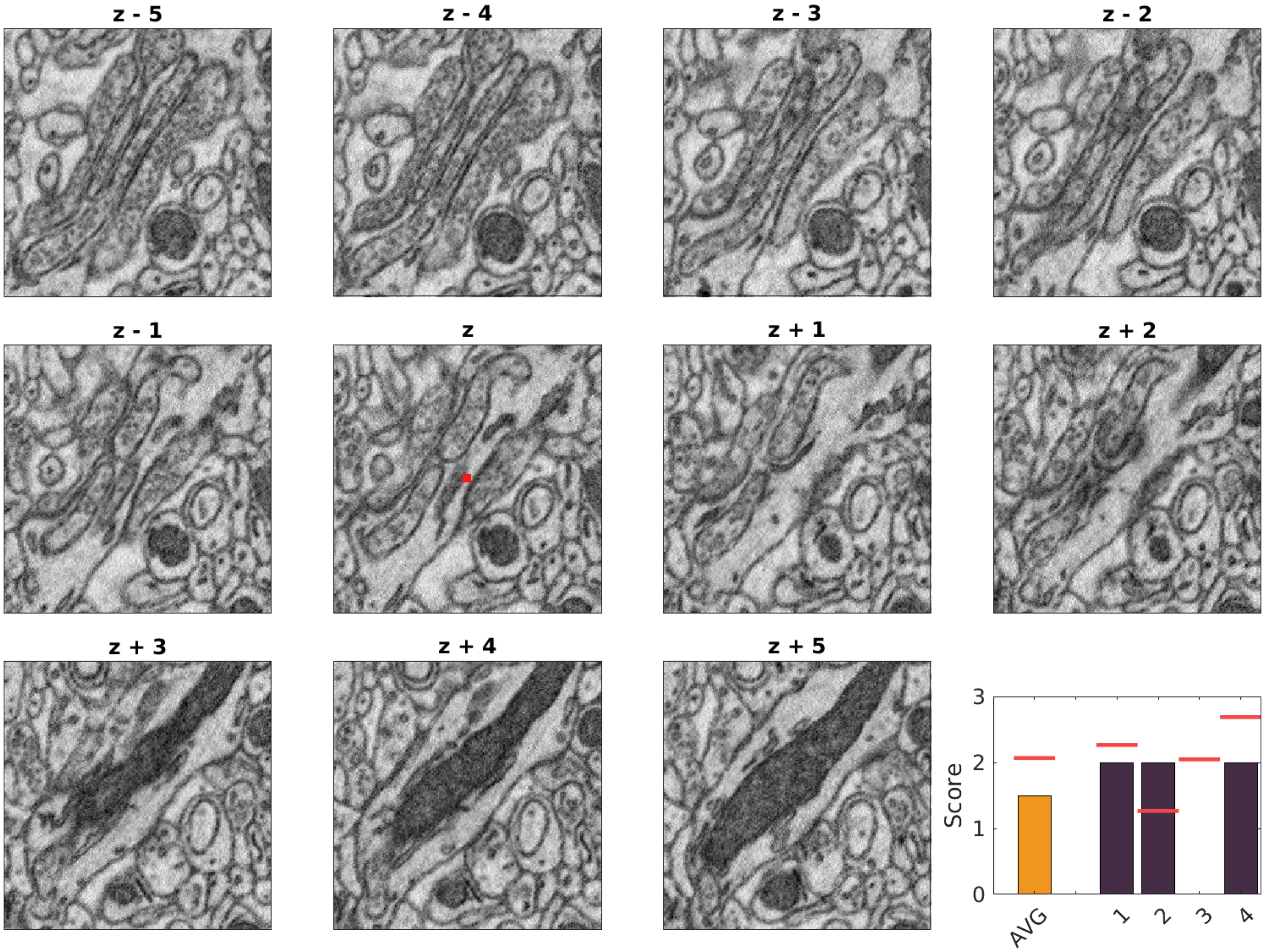

3833, 3744, 557

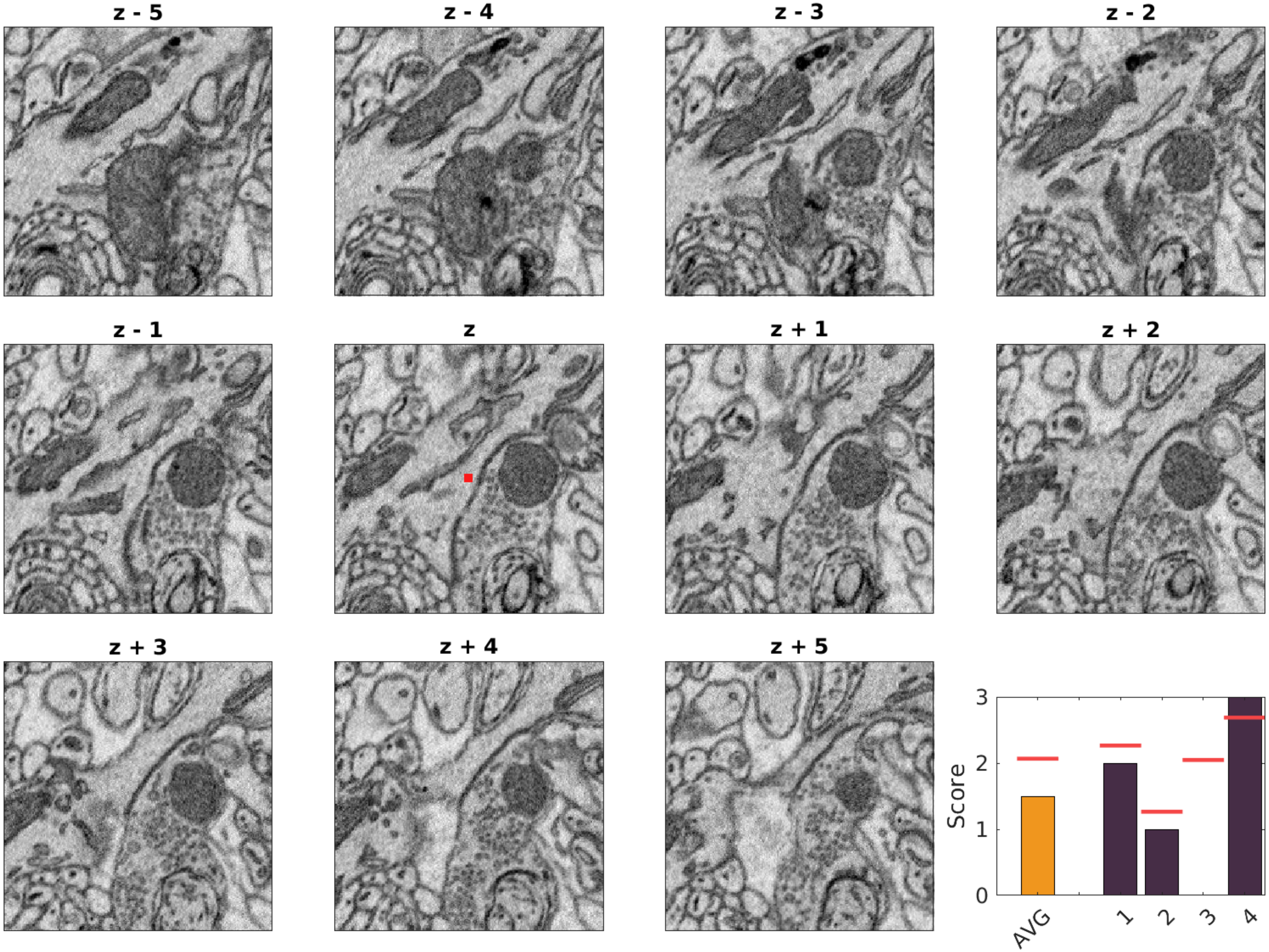

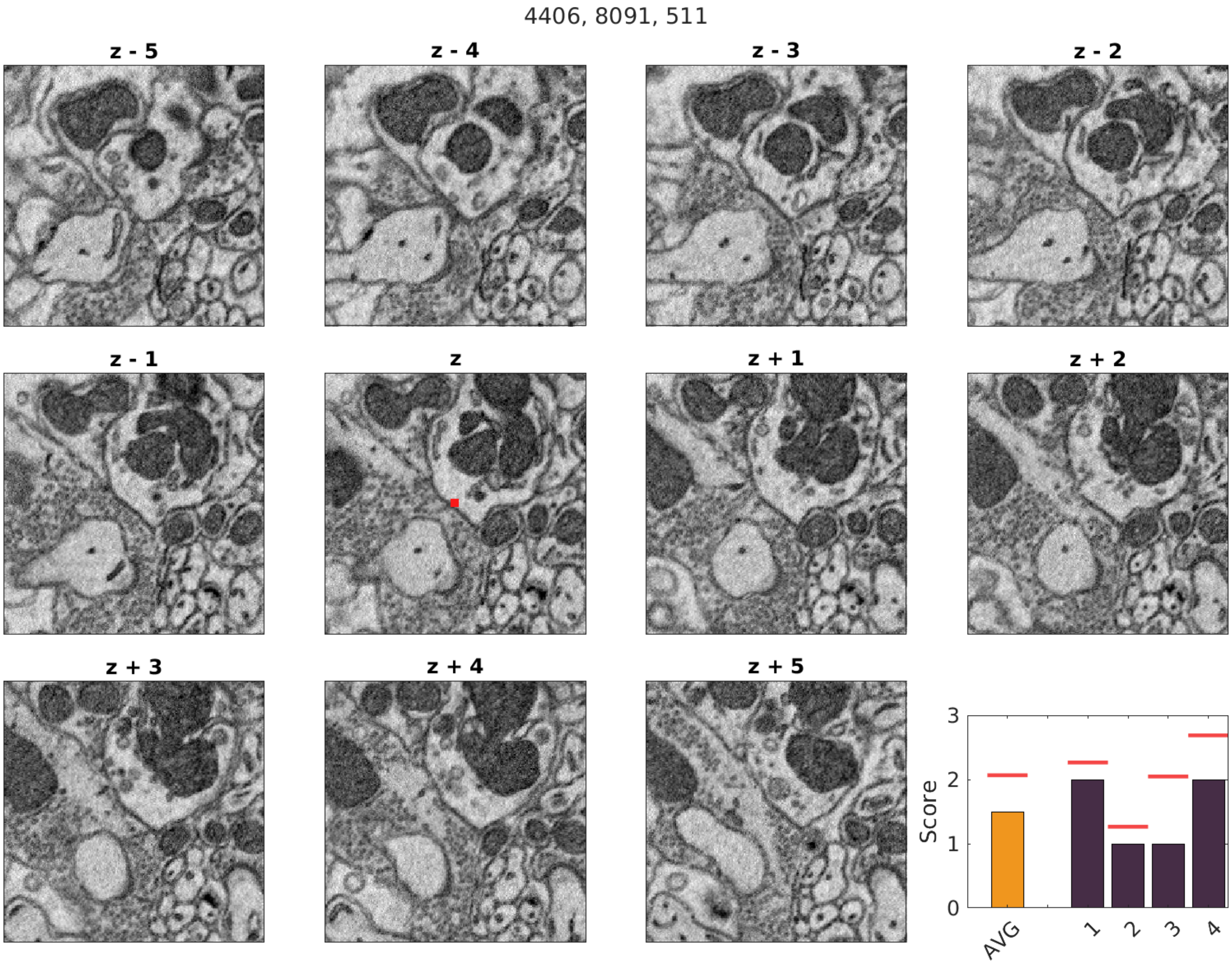
4406, 8091, 511
z - 5
z - 4
z - 3
z - 2
z - 1
z
z + 1
z + 2
z + 3
z + 4
z + 5
Score
3
2
1
0
AVG
1
2
3
4

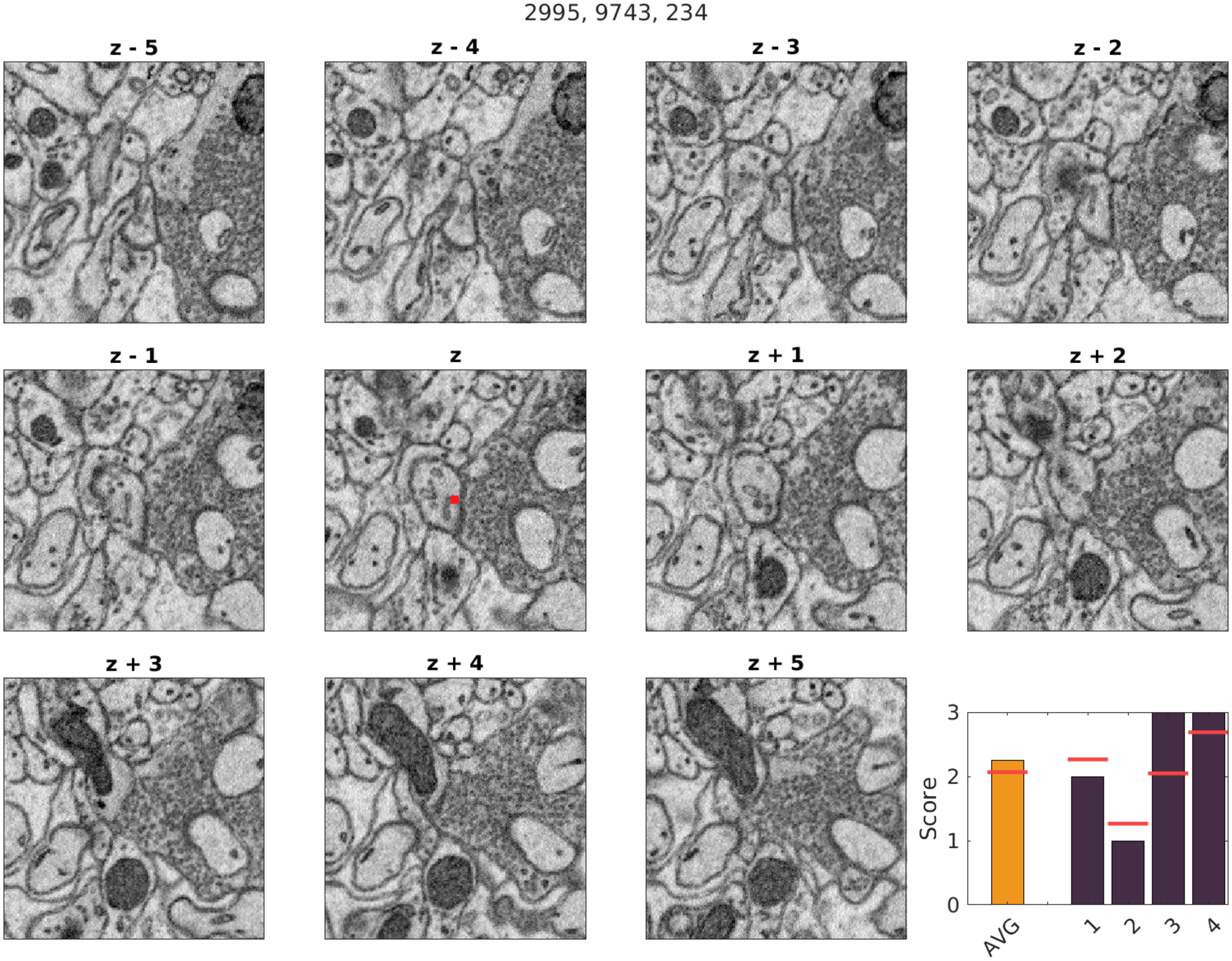
2995, 9743, 234
z - 5
z - 4
z - 3
z - 2
z - 1
z
z + 1
z + 2
z + 3
z + 4
z + 5
Score
0
1
2
3
AVG
1
2
3
4

2660, 9000, 258

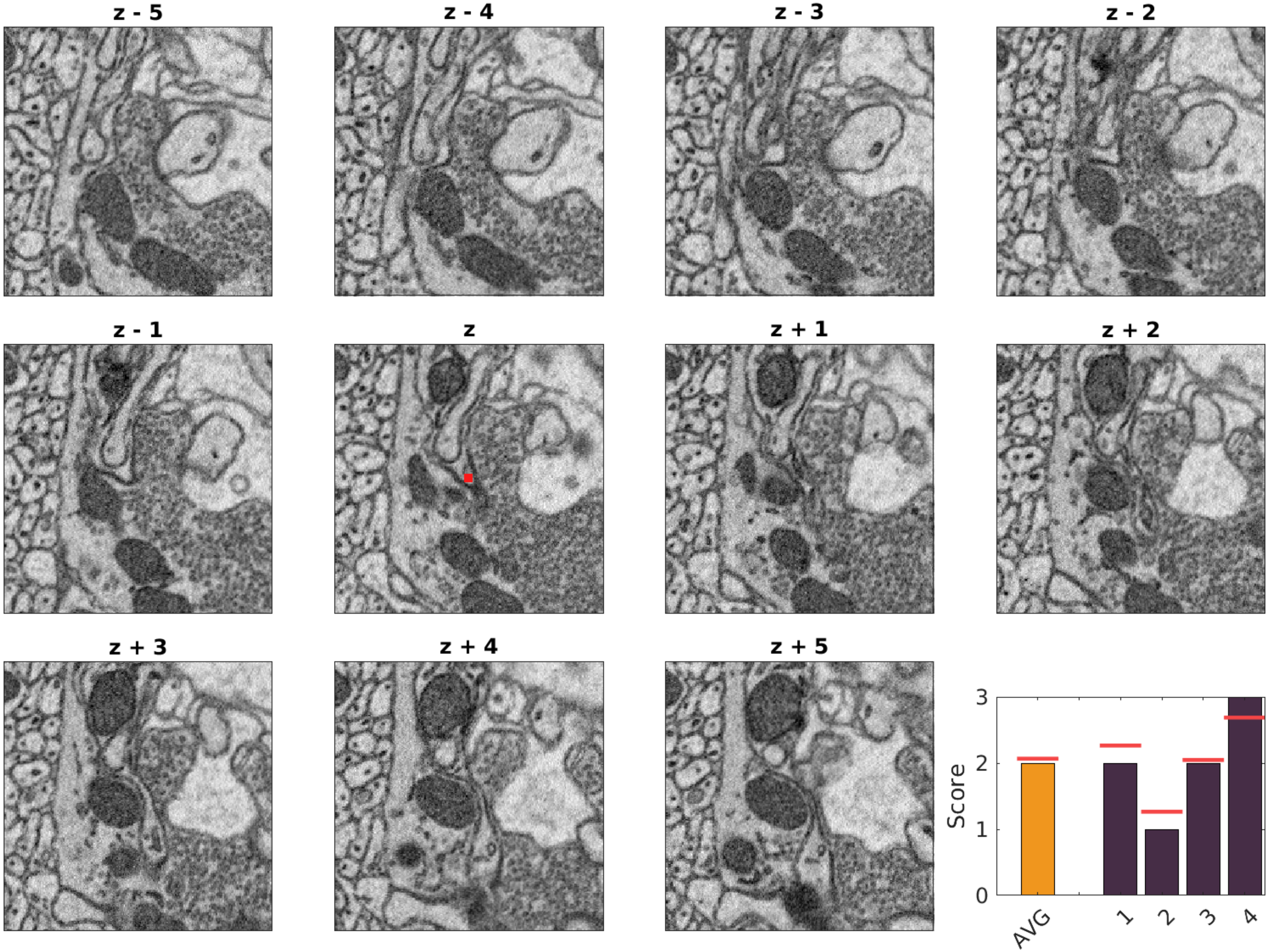

1222, 7840, 621

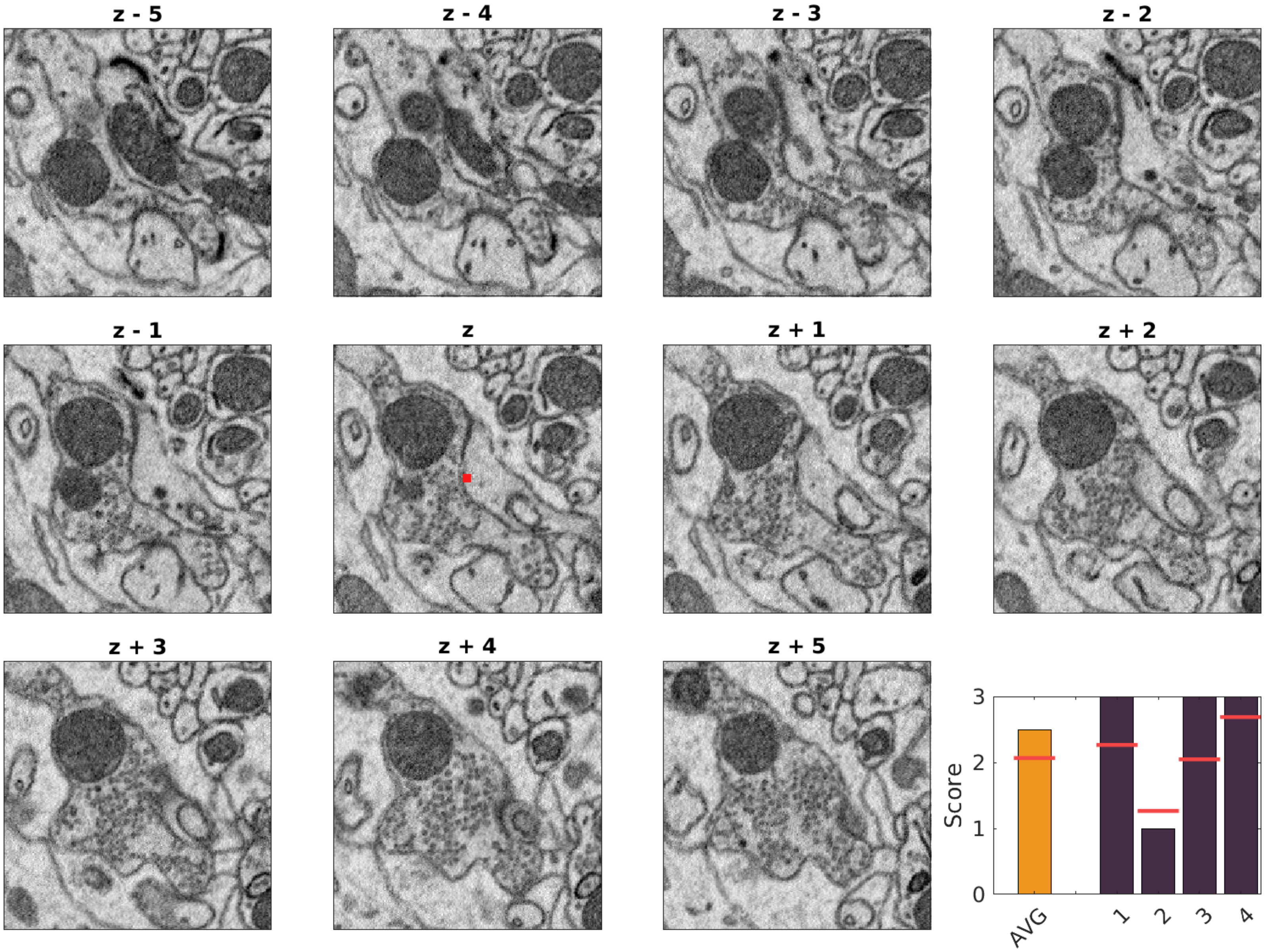

1634, 8177, 625

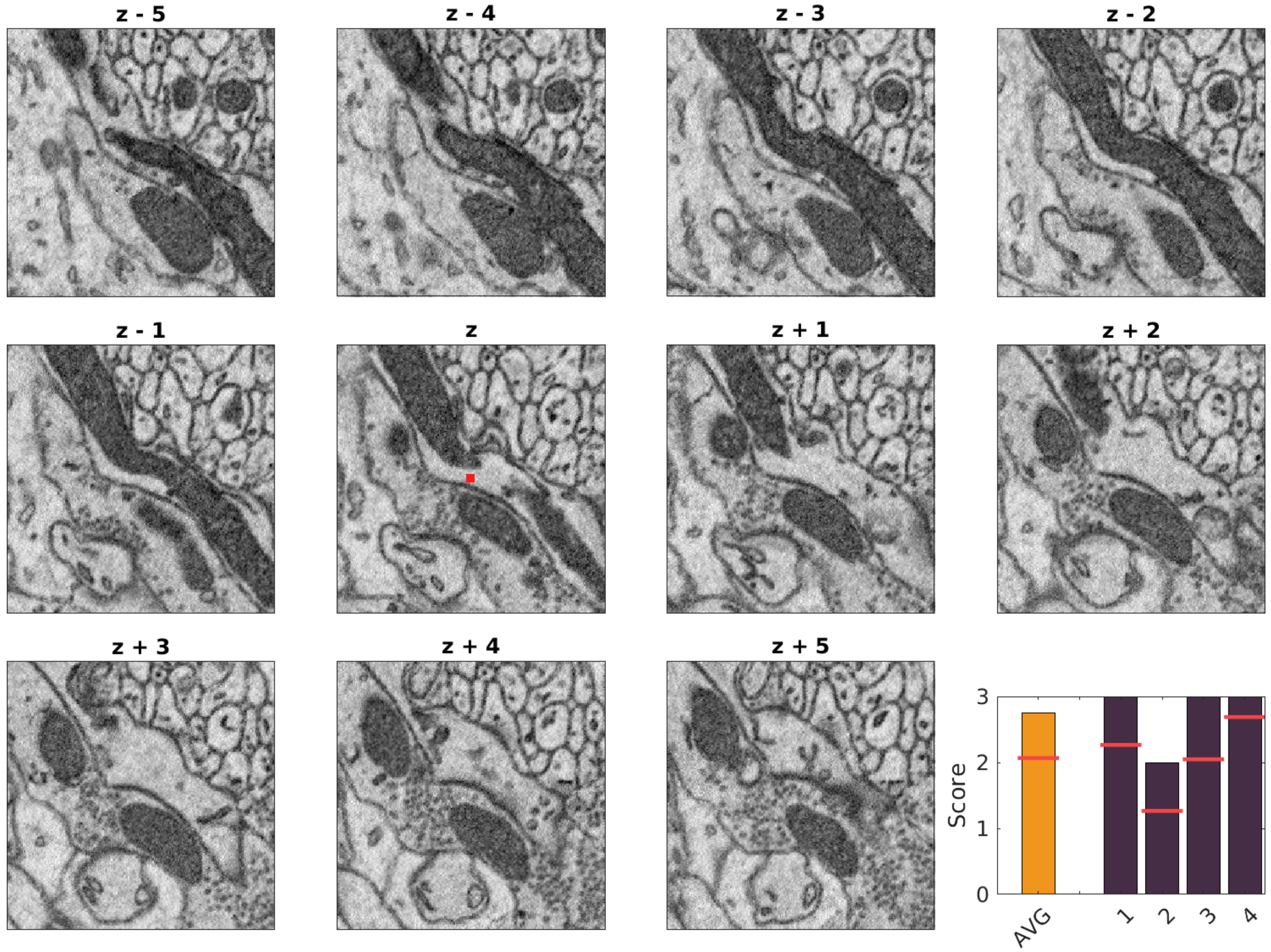
