## Supplementary Information for "A cerebellar disinhibitory circuit supports synaptic plasticity"

**Supplementary Data 1. Open-source data repository with annotation files.** The complete dataset of this study, shared in the open-source data repository https://bossdb.org/project/park2023 with the interactive data viewer, neuroglancer (https://github.com/google/neuroglancer). The data include the registered 3D EM images, the proofread segmentation of each segment representing one neuron, and the 3D rendering of the neurons. The accompanying ZIP file contains 4 CSV files for the list of segment numbers for respective cell types (PC, IN, CF, and PF) and 6 CSV files for the list of synapse annotations for respective synapse types (CF-IN, CF-PC, IN-IN, IN-PC, PF-IN, and PF-PC), which can be imported and displayed with neuroglancer.

**Supplementary Data 2. Gallery of the main Purkinje cells.** All the pages have the same panel organisation and each page contains the data for each of the 10 main PCs. The cell identification code in parentheses at the bottom left of the page is the same as that in Supplementary Data 1. **a**, The XY-view (top) and the XZ-view (bottom) of the PC. Scale bar: 20 µm. **b-c**, The locations of the synaptic inputs to the PC, displayed on the PC dendritic skeleton (green) and soma (grey circle). The inhibitory IN1 (red) and IN2 (blue) inputs **(b)** and the excitatory CF (orange) and PF (purple) inputs **(c)**. **d**, The PCL distance of the IN input synapses (*d*_s_) plotted for each IN on the *x*-axis, whose orders are ranked by the PCL distance of the somata (*d*_IN_). **e**, Disinhibition coverages, reflecting the proportion of IN1s presynaptic to a PC are disynaptically connected by the PC-innervating CF via IN2s. The definition can be varied by using the number of cells, the number of synapses, or the total area of synapses. **f-h**, Polar plots of the angular distributions of CF **(f)**, IN **(g)**, and PF **(h)** volumes around the PC dendrites. Figs. 3d–f were reproduced for each cell.

**Supplementary Data 3. Gallery of the main interneurons.** All the pages have the same panel organisation and each page contains the data for each of the 26 main INs. The cell identification code in parentheses at the bottom left of the page is the same as that in Supplementary Data 1. **a**, The XY-view (top) and the XZ-view (bottom) of the IN. Scale bar: 20 µm. **b-c**, The locations of the synaptic inputs to the PC, displayed on the IN dendritic skeleton and soma. The excitatory CF (orange) and PF (purple) inputs **(b)** and the inhibitory IN1 (red) and IN2 (blue) inputs **(c)**. **d**, The locations of the inhibitory synaptic outputs from this IN to PCs (green) and other INs (blue), displayed on the axonal skeleton and soma (grey). **e**, The TPI of this IN is shown in colour (red for IN1 or blue for IN2), while those of the other main INs are shown in grey. **f**, The proportions of the synaptic outputs to PC (green), IN1 (red), IN2 (blue), and INx (grey). **g-i**, The dendrite density **(f)**, axon complexity **(g)**, and IN-PC synapse distance **(h)** vs. the soma distance from the PCL (*d*_IN_) for this IN is depicted in colour, while those of the other main INs are shown in grey. Figs. 4g–i were redisplayed for each cell.

**Supplementary Data 4. Gallery of the main climbing fibres.** All the pages have the same panel organisation and each page contains the data for each of the 10 main CFs. The cell identification code in parentheses at the bottom left of the page is the same as that in Supplementary Data 1. The CF innervates the PC of Supplementary Data 2 on the same page. **a**, The XY-view (top) and the XZ-view (bottom) of the CF. Scale bar: 20 µm. **b**, The locations of the excitatory synaptic outputs to PCs (green), displayed on the skeleton (orange). **c**, The locations of the excitatory synaptic outputs to IN1s (red) and IN2s (blue), displayed on the skeleton (orange). The numbers are the cell identification code of the INs.

**Supplementary Data 5. Gallery of all the identified CF-IN synapses.** All the pages have the same panel organisation and each page contains the data for each of the 53 CF-IN synapses. **a**, Eleven consecutive EM section images showing a synapse. The coordinate of the central point of the synapse on section *z* (red dot) is shown on the top, which can be used for Supplementary Data 1. Scale bar: 500 nm. **b**, The synaptic score of the synapse, the average (orange), and the individual scores by experts (blue). The red bars indicate the average score given by the particular human expert for all the synapses, showing the tendency of the person as a reference.
